# abx_amr_simulator: A simulation environment for antibiotic prescribing policy optimization under antimicrobial resistance

**DOI:** 10.64898/2026.03.12.711398

**Authors:** Joyce Lee, Seth Blumberg

## Abstract

Antimicrobial resistance (AMR) threatens antibiotic effectiveness, but quantitatively evaluating stewardship strategies under partial observability and delayed feedback remains difficult in real-world data. We developed abx_amr_simulator, a Gymnasium-compatible simulation framework, and used it to benchmark reinforcement learning (RL) prescribing policies against value-iteration (VI) benchmarks and fixed prescribing rules across four experiment sets of increasing complexity, with varying levels of information degradation.

Across scenarios, temporal abstraction was consistently important: flat PPO was competitive only in simpler settings, whereas hierarchical PPO was generally needed when prescribing decisions had delayed, coupled effects on future resistance. We found that adding recurrent memory did not uniformly improve performance; its value was context-dependent. In some degraded-information settings, memoryless policies performed better by adopting conservative update-responsive behavior, while in more complex, multi-signal partially observable settings, recurrent memory provided modest advantages.

Patient heterogeneity and risk-stratification signals were major determinants of policy quality. When agents could differentiate higher-from lower-risk patients, they more reliably learned selective treatment behavior, stabilized AMR, and improved clinical outcomes. Exaggerated risk stratification modestly outperformed accurate stratification, while compressed stratification produced moderate degradation. In more realistic settings combining noisy patient observations, delayed AMR surveillance, and multi-patient decisions, hierarchical agents outperformed fixed prescribing rules across both stewardship and clinical metrics, converging to conservative low-AMR equilibria with reduced cross-seed variance.

Across experiments, results support the utility of hierarchical RL as a best-case policy-analysis tool for stewardship under uncertainty, while also highlighting that performance estimates are sensitive to observation structure and training horizon design. The framework provides a controlled environment for hypothesis generation and for stress-testing prescribing strategies before translation to policy-relevant settings.

## 1 Introduction

Rising antimicrobial resistance (AMR) is widely recognized as a major threat to global public health, adversely affecting the clinical care and outcomes of millions of patients worldwide by reducing the effectiveness of currently available antibiotics (WHO 2015). Recent estimates suggest that 4.95 million deaths globally in 2019 were associated with antibiotic-resistant bacterial infections, with a disproportionate burden borne by those living in resource-limited settings (Murray et al. 2022). In response, a range of interventions have been proposed and implemented to curb the emergence and spread of AMR (America (IDSA) 2011). In the United States, one of the primary strategies has been the establishment of antibiotic stewardship programs (ASPs) (Sutton and Ashley 2024), which are multidisciplinary efforts encompassing clinician education, dissemination of prescribing guidelines, compilation of antibiograms from microbiological data, and prospective review and optimization of antibiotic regimens, particularly in inpatient settings (Barlam et al. 2016).

Despite the widespread adoption of such interventions, the quantitative evidence base evaluating their population-level impact remains limited. Many studies assessing the effectiveness of ASPs or related strategies have been constrained in geographic scope, duration, or both (Bertollo, Lutkemeyer, and Levin 2018). A fundamental challenge to performing quantitative impact evaluation of these programs is the pervasive partial observability of key system components (Laxminarayan et al. 2013). Even in resource-rich settings, where antibiotic prescription records are often available, true selection pressure on pathogen populations is incompletely observed due to unmeasured sources of antibiotic exposure, including agricultural use and environmental contamination (Van Boeckel et al. 2015). Measurement of AMR itself presents an even greater challenge: while initiatives such as the WHO’s Global Antimicrobial Resistance and Use Surveillance System (GLASS) have sought to standardize surveillance, participation remains voluntary and coverage incomplete, and no universally adopted scalar metric of resistance at the community level yet exists (Organization 2022; Leth and Schultsz 2023). One commonly used proxy, the antibiogram, is subject to substantial biases, as it usually reflects only cultured clinical isolates from patients seeking care and is typically updated infrequently, often on an annual basis (Truong et al. 2021). Consequently, antibiograms provide an incomplete and temporally lagged view of underlying resistance dynamics.

Given these challenges, it is difficult to directly quantify the long-term effects of antibiotic stewardship program interventions using observational or interventional studies alone (Bertollo, Lutkemeyer, and Levin 2018; Schweitzer et al. 2019). Simulation-based approaches offer a complementary framework in which key mechanisms can be explicitly specified and systematically varied, enabling controlled investigation of trade-offs that are difficult or impossible to observe in real-world settings.

Prior simulation-based work in this domain has largely operated at the pathogen level, modeling evolutionary dynamics of resistance under drug pressure (King et al. 2025; Weaver et al. 2024). While these models have yielded important insights into resistance evolution, they do not capture the patient-level clinical dynamics — including treatment decisions, outcomes, and feedback between prescribing behavior and population-level resistance — that are central to evaluating antibiotic stewardship interventions. Separately, there has been growing interest in applying machine learning to AMR-related problems, including resistance prediction and empirical antibiotic selection (Haredasht et al. 2025; Harandi et al. 2025). However, these approaches predominantly utilize supervised learning to focus on prediction rather than optimization, and typically operate on static snapshots of patient data rather than modeling the dynamic, longitudinal consequences of prescribing decisions at the population level. To our knowledge, the only prior application of reinforcement learning to an AMR-related problem is Weaver et al. (Weaver et al. 2024), which uses RL to identify optimal treatment cycling strategies at the pathogen level. Our work differs fundamentally in that we apply RL to optimize clinical prescribing policy within a patient-level simulator, with the explicit goal of evaluating trade-offs between individual clinical outcomes and population-level resistance.

In this study, we conduct a series of reinforcement learning experiments using a simulation framework to investigate antibiotic prescribing strategies under varying levels of observability and uncertainty. The experiments were performed using abx_amr_simulator, a Python-based simulation environment designed to model the relationship between prescribing decisions and the evolution of AMR (Lee 2026). A detailed description of the simulator’s architecture and functionality is provided in a separate manuscript (Lee and Blumberg 2026), while here we focus on its application to experimental settings.

Using the simulator, we explored the trade-off between immediate clinical benefit and long-term resistance management. In simplified, fully observable settings, we find that fixed prescribing rules that emulate how real-world prescribers make clinical decisions about antibiotic treatment perform fairly well; however, as uncertainty and delayed effects are introduced, the performance of these fixed prescribing rules degrade. We hypothesize that RL algorithms are capable of optimizing antibiotic prescribing policies even under conditions of partial observability. By training different types of RL agents across a range of simulated scenarios, we perform comparative policy analyses to characterize the magnitude and structure of potential gains achievable through adaptive prescribing strategies relative to clinically interpretable fixed prescribing rules.

In the following sections, we describe our experimental design, including how we simulate synthetic patient populations, define the reward function, and vary the quality and timeliness of information available to the RL prescribing agent.

## 2 Materials and Methods

### 2.1 Reinforcement Learning Framework

Optimizing antibiotic prescribing is a sequential decision-making problem under uncertainty: prescribing decisions affect both immediate patient outcomes and longer-term resistance dynamics, and the true underlying state of the system is rarely fully observable. We address this using reinforcement learning (RL), a class of machine learning methods designed to maximize cumulative rewards in such settings. The experiments were conducted using abx_amr_simulator, a Python-based simulation framework designed to model antibiotic prescribing decisions and antimicrobial resistance (AMR) dynamics, built on the standard Gymnasium API. A detailed description of the simulator’s architecture, functionality, and configuration options is available in the abx_amr_simulator GitHub documentation (Lee 2026) and companion preprint (Lee and Blumberg 2026). This section provides a brief overview of key components and experimental settings relevant to the study.

### 2.2 Agent-environment interaction loop

In reinforcement learning, the way that agents learn to optimize sequential decision making is by interacting with the environment. In this work, the environment is created by abx_amr_simulator according to the user’s specifications in order to construct a specific simulated scenario. The environment consists of three main subcomponents: a PatientGenerator, an array of AMR_LeakyBalloon models that keep track of each antibiotic’s resistance level, and a RewardCalculator which computes the reward for the agent’s action at each timestep based on patient outcomes. (For more detailed information on the abx_amr_simulator package and the subcomponents of the environment class, see S2 in Supplementary Materials, also documentation in Github repository.)

For the reinforcement learning agents used in our experiments, we used PPO (proximal policy optimization) agent implementations from the Python package stable-baselines3. Each of the subcomponents of the environment is discussed further below, as are the various types of RL agents used in our experiments.

A schematic overview of the simulator components and the agent–environment interaction loop is shown in Figure 1.

**Figure 1:**
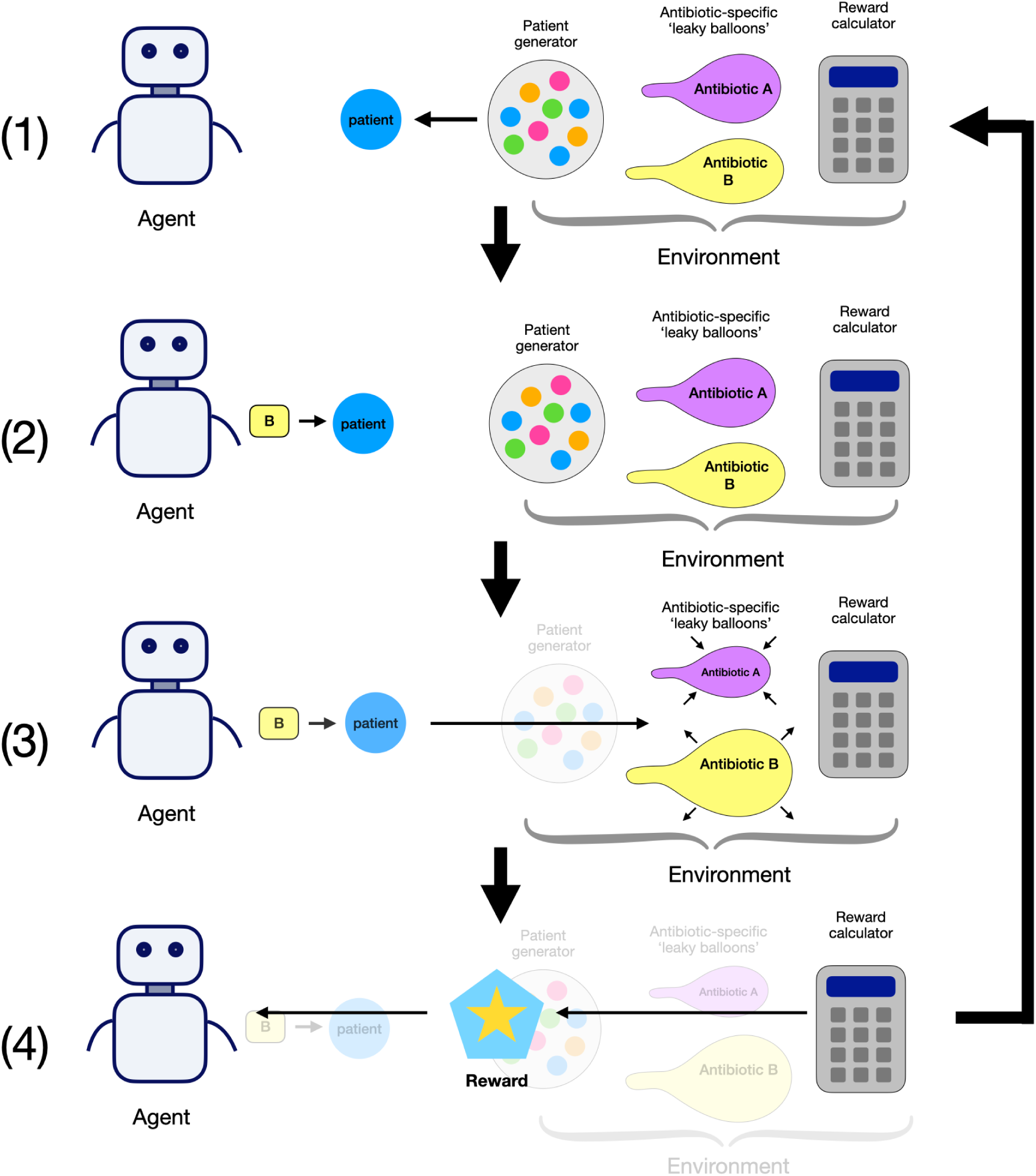
Schematic overview of the environment and its subcomponents (patient generator, antibiotic leaky balloons, and reward calculator), and the interaction loop between the agent and environment. 1) The environment uses the PatientGenerator to create a patient and present that patient to the agent. 2) The agent decides to prescribe Antibiotic B for the patient. 3) The agent informs the environment that it decided to prescribe Antibiotic B for the current patient. 4) The environment uses the RewardCalculator to compute the reward for the current action, updates each antibiotic’s AMR LeakyBalloon, and advances to the next timestep and next patient.

Note: this figure is reproduced from the companion software manuscript (Lee and Blumberg 2026).

### 2.3 Environment subcomponents

#### 2.3.1 Patient Population and Observability

At each timestep, the environment uses the PatientGenerator subcomponent to generate a cohort of synthetic patients, which are characterized by attributes that determine their clinical profile and treatment outcomes. These attributes include infection probability, clinical benefit and failure multipliers, and spontaneous recovery probability. Attribute values can be set to constant values (creating homogenous patient populations), or drawn from a probability distribution (creating heterogeneous populations). Observations available to the RL agent can be manipulated by introducing noise or bias, or by concealing certain attributes altogether, therefore enabling exploration of prescribing strategies under varying levels of partial observability.

#### 2.3.2 Antibiotic Resistance Dynamics

Antimicrobial resistance dynamics are modeled using the AMR_LeakyBalloon class, which tracks resistance to each antibiotic as a function of cumulative prescribing pressure. The model operates as a soft-bounded accumulator: prescribing a given antibiotic increases the internal latent ‘pressure’, while in the absence of prescribing, this internal pressure decays over time as selection pressure is relieved — analogous to a balloon that inflates as air is pumped in, but slowly deflates when left alone. The observable resistance level is derived from this latent pressure via mapping through a sigmoid function, and represents the probability that a new infection at the current time is resistant to the given antibiotic. The simulator also supports cross-resistance, allowing users to specify how prescribing one antibiotic influences resistance to others.

As with patient attributes, the fidelity of resistance level observations can be manipulated by introducing noise, bias, or delay, facilitating systematic investigation of prescribing strategies under imperfect surveillance.

#### 2.3.3 Treatment Decisions

At each timestep, the agent makes an independent treatment decision for each patient: prescribe antibiotic A, prescribe antibiotic B,…, or withhold treatment. The agent cannot prescribe multiple antibiotics to the same patient simultaneously, reflecting standard clinical practice of monotherapy (single-drug treatment) for straightforward infections. This constraint simplifies the decision space and aligns with real-world prescribing norms.

#### 2.3.4 Reward Function

The reward function as calculated by the RewardCalculator balances immediate clinical outcomes against long-term antimicrobial resistance considerations. It consists of two components: the individual reward, which reflects the clinical benefit or failure of treating patients, and the community reward, which penalizes high levels of antimicrobial resistance (AMR) across all antibiotics. These components are combined into a single scalar reward using a tunable weight parameter *λ* ∈ [0, 1], allowing the user to control the relative importance of individual versus community objectives. In all experiments reported in this manuscript, we fixed *λ* = 0, so agents were optimized using only individual clinical reward. This design allows us to test whether AMR-preserving behavior can emerge from long-horizon environment dynamics and policy architecture without explicit AMR reward shaping.

The individual reward is calculated based on patient-specific attributes, antibiotic-specific properties, and discrete clinical outcomes (e.g., treatment success, failure, or adverse effects). The community reward, on the other hand, is based on the observed AMR levels of all antibiotics, ensuring consistency with realistic surveillance data rather than ground truth resistance levels. This design encourages agents to learn prescribing strategies that work within realistic informational constraints.

For mathematical definitions of the reward function, including equations, parameter descriptions, and details on stochastic modeling of clinical outcomes, see Supplementary Materials S2.

### 2.4 Reinforcement learning agents

To evaluate how policy architecture affects antibiotic prescribing under uncertainty, we implemented four PPO-based agent classes using the stable-baselines3 Python library: flat memoryless PPO, flat recurrent PPO, hierarchical memoryless PPO, and hierarchical recurrent PPO.

The memoryless agents make each decision from the current observation only. By contrast, recurrent agents carry a short internal memory across timesteps, so recent observation history can influence the current decision. This distinction is important when AMR information is delayed or intermittently updated: agents that retain recent history may better approximate the latent current resistance state.

We also compared flat and hierarchical agent architectures. Flat agents choose a treatment action directly at each timestep. Hierarchical agents instead choose among higher-level options (also called workers), where each option encodes a clinically interpretable prescribing strategy over one or more steps. In our experiments, we used a ‘heuristic-worker’ option type, which uses fixed risk-based rules to choose treatment for each patient.

Heuristic workers use the observable patient attributes to compute an expected ‘reward’ for each available action, then uses rule-based strategies to select which action to take (e.g., “treat high-risk patients with antibiotic A,” “withhold from low-risk patients”). The manager then learns to select among these context-appropriate options based on patient characteristics and AMR state. Unlike the fixed prescribing rules described in Section 3.6, heuristic workers serve as options within a learning architecture—the manager learns to dynamically select which worker to deploy based on AMR state and population context. Heuristic workers themselves encode deterministic clinical logic (similar to the fixed rules), but the key difference is that the hierarchical agent learns when to switch between them, enabling adaptive cycling strategies that fixed rules cannot achieve.

We created a variety of heuristic-workers by using different risk thresholds to choose antibiotic treatment for patients, and also by varying the antibiotic ‘preference’ of each heuristic worker. Creating a range of options for the manaer agent to choose from allows it more flexibility in optimizing its antibiotic prescribing strategy. Our use of hierarchical agents was motivated by the long-horizon nature of AMR dynamics, in which current prescribing can affect resistance and treatment utility many timesteps later. (For more information, please see Supplementary Materials S2: Options and Option Libraries, which discusses the option types and instances that were used in these experiments.)

### 2.5 Experimental Pipeline

Effective reinforcement learning requires careful hyperparameter tuning. The abx_amr_simulator package uses the Optuna library (a hyperparameter optimization framework) for automated hyperparameter tuning (Akiba et al. 2019). This allows us to build modular end-to-end pipelines with three phases:

#### Tuning Phase

Optuna explores the hyperparameter search space, evaluating candidate configurations on preliminary environments. For each trial, the agent undergoes truncated training across 5 random seeds, with the study ultimately selecting the hyperparameter set that maximized a stability-adjusted objective (mean reward minus a modest variance penalty across seeds).

#### Training Phase

Multiple agent copies (20 per experiment) are initialized with optimized hyperparameters and trained on target environments, each using a distinct random seed. This approach accounts for the empirical sensitivity of learned policies to initialization: we observed for certain scenarios and agent architectures, environmental stochasticity (particularly patient infection and resistance status across seeds) substantially influences policy acquisition, even with identical hyperparameters.

#### Evaluation Phase

Following training, each seed-deployed agent executes 3 evaluative rollouts with logging enabled. During evaluation, we record three categories of cumulative metrics:

- *Clinical Outcomes*: patients experiencing benefit, failure, or adverse effects
- *Patient Status*: counts by true infection status and treatment received
- *Antibiotic Response*: prescriptions per antibiotic, and sensitive vs. resistant infections treated

These metrics are agnostic to the decision algorithm—applicable to RL agents, fixed prescribing rules, or any strategy—enabling quantitative policy comparison. When the reward function is held constant while evaluating different policies, raw reward provides a valid shorthand for relative performance. However, examining cumulative count metrics remain essential for interpretation, since they show which outcome components (for example, clinical successes, failures, and adverse events) are driving those reward differences. All experiments in this study used the same reward function with fixed RewardCalculator hyperparameter values.

Figure 2 shows a schematic of the full experimental pipeline.

**Figure 2:**
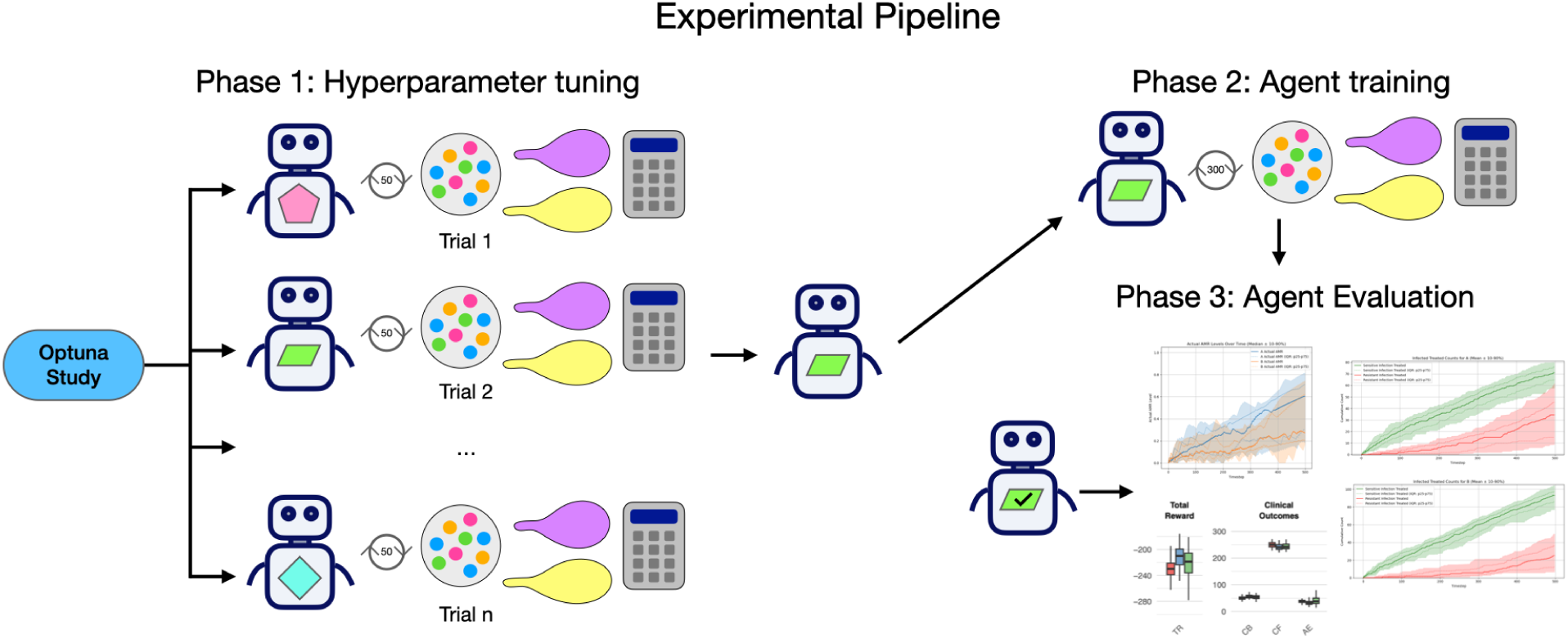
The experimental pipeline consists of three phases. Phase 1 (Tuning): An Optuna hyperparameter search evaluates candidate configurations via truncated training across 5 random seeds, selecting the configuration that maximizes a stability-adjusted objective (mean reward minus a modest variance penalty across seeds). Phase 2 (Training): Twenty agent copies are initialized with the tuned hyperparameters and undergo prolonged training, each using a distinct random seed. Phase 3 (Evaluation): Each trained agent executes three evaluative rollouts in the environment across unseen random seeds; clinical outcome, patient status, and antibiotic response metrics are logged and aggregated for analysis.

### 2.6 Fixed Prescribing Rule Baseline

To evaluate whether learned policies provide value beyond static decision rules, we compared trained RL agents against a fixed prescribing baseline that perform no learning or adaptation.

#### Expected-reward (greedy)

For each patient, compute the expected net clinical benefit (expected reward) of treating with each available antibiotic based on observed patient attributes and known scenario parameters. If no antibiotic has positive expected reward, withhold treatment. Otherwise, prescribe the antibiotic with the highest expected reward.

This fixed prescribing rule is intended to imitate how providers actually make decisions about antibiotic treatment, akin to static practice guidelines: they use available patient information and visible AMR data, but do not update behavior through experience. This makes them suitable non-learning comparators for isolating the incremental value of RL-based adaptation.

For fair comparison, fixed-rule policies are evaluated under the same rollout framework used for trained agents (multiple seeds and repeated evaluation episodes per seed), so differences reflect policy behavior rather than evaluation protocol artifacts.

### 2.7 Experiments: varying partial observability of patient attributes and AMR levels

Using the abx_amr_simulator package, we evaluated policies across three different antibiotic scenarios of increasing complexity: (1) a single-antibiotic scenario, (2) a two-antibiotic scenario without cross-resistance, and (3) a two-antibiotic scenario with moderate asymmetric cross-resistance.

For each of these antibiotic scenarios, we varied patient-population structure and information quality across four sets of experiments (Table 1). Populations were either homogeneous (fixed patient attribute values) or heterogeneous (attribute values sampled from Gaussian distributions), including mixed high-risk/low-risk cohorts in the heterogeneous settings. Importantly, the reward function was held constant across all experiment sets and antibiotic scenarios (with fixed RewardCalculator hyperparameter values), so performance differences reflect policy behavior under different observability and population conditions rather than changes in reward design.

**Table 1:**
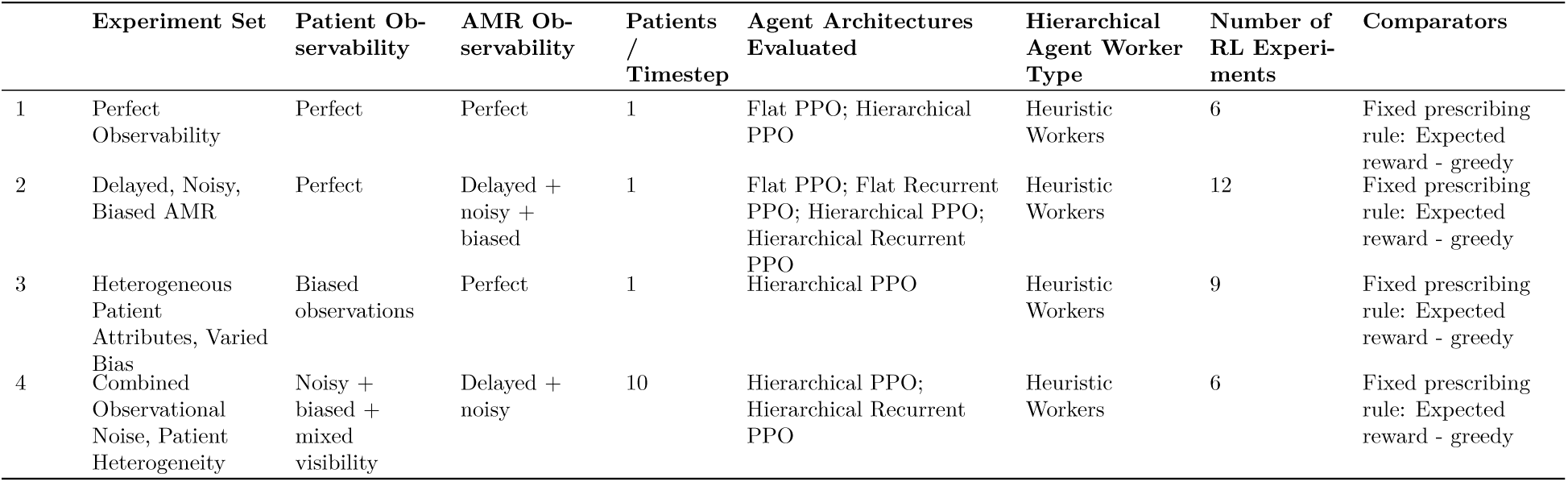
Overview of experimental conditions across four experiment sets. Observability, agent architectures, and comparators were systematically varied. All sets evaluated three antibiotic scenarios: single-antibiotic, two-antibiotic without cross-resistance, and two-antibiotic with cross-resistance.

We now discuss the benchmark scenario construction, and describe the four experiment sets that progressively degrade observability and increase heterogeneity.

#### Benchmark scenario design and reference policies derived from fixed prescribing rules

In designing the three different antibiotic scenarios, we selected environment parameter values in order to produce nontrivial stewardship dynamics (i.e., situations that were required policies with meaningful trade-offs, rather than trivial always-treat or never-treat behavior). In particular, for the homogeneous benchmark population we fixed baseline infection probability at *p* = 0.7, so treatment withholding is not trivially optimal and agents must balance near-term clinical benefit against longer-term resistance consequences.

In order to compute a sensible baseline to use as a comparator when evaluating the performance of the reinforcement learning agents in these scenarios, we rolled out the fixed prescribing rule ‘Expected Reward - Greedy’ as described in Section 3.6 in each benchmark scenario, under each of the conditions described in Table 1.

#### Experiment Set 1: Perfect Observability

We first conducted a set of experiments in which the agent had full observability of the environment: all patient attributes and antibiotic-specific AMR levels were presented to the agent without noise, bias, or delay, and all patient attributes are constant values and homogenous across all patients. These experiments serve as a best-case baseline, establishing the level of performance achievable by reinforcement learning agents and fixed prescribing rules when the true state of the system is fully known. We tested both flat and hierarchical PPO architectures; the rationale for the hierarchical architecture and the option designs used are described in Section 3.4.

#### Experiment Set 2: Delayed, Noisy, Biased AMR

In the second set of experiments, we introduced partial observability in the form of noise, bias, and temporal delay in the AMR levels visible to the agent, while maintaining perfect observability of patient attributes, where the patient population is again constant and homogeneous. This type of degraded information regarding AMR levels does resemble real-world clinical practice: as discussed in the Introduction, antibiograms provide an incomplete and typically delayed proxy for community resistance levels, requiring clinicians to make prescribing decisions using outdated and potentially biased information.

Introducing delayed updates to AMR observations renders the environment partially observable, as the agent now only has periodic access to updated resistance levels; otherwise, during inter-update periods, this information is stale. For this reason, in addition to memoryless PPO agents, we evaluated recurrent flat and hierarchical PPO agents with LSTM-based policies, which are commonly used to handle temporal partial observability by allowing agents to integrate information over time. Intuitively, such agents may learn to maintain an internal estimate of latent resistance dynamics based on prior observations and prescribing outcomes, analogous to a clinician forming informal beliefs from accumulated experience.

#### Experiment Set 3: Heterogeneous Patient Attributes, Varied Bias

In Experiment Set 3, we introduced heterogeneous patient populations with true differences in infection risk and treatment response while maintaining population-level averages matching the homogeneous baseline (Experiment Set 1). Patients were drawn from two subpopulations (high-risk and low-risk) with equal probability, where infection probability followed Gaussian distributions (*μ* = 0.84 vs. *μ* = 0.56, *σ* = 0.1) rather than constant values. This creates ground-truth risk stratification that optimal policies should exploit.

We then manipulated observational bias in how agents perceive these patient attributes, testing three conditions for each antibiotic scenario (see Table 2): (1) Accurate: true risk levels correctly observed, (2) Exaggerated: high-risk patients appear even higher risk, low-risk appear even lower, and (3) Compressed: risk differences appear smaller than they truly are. This isolates the cost of misperceiving patient heterogeneity when AMR information remains perfect and up-to-date.

**Table 2:**
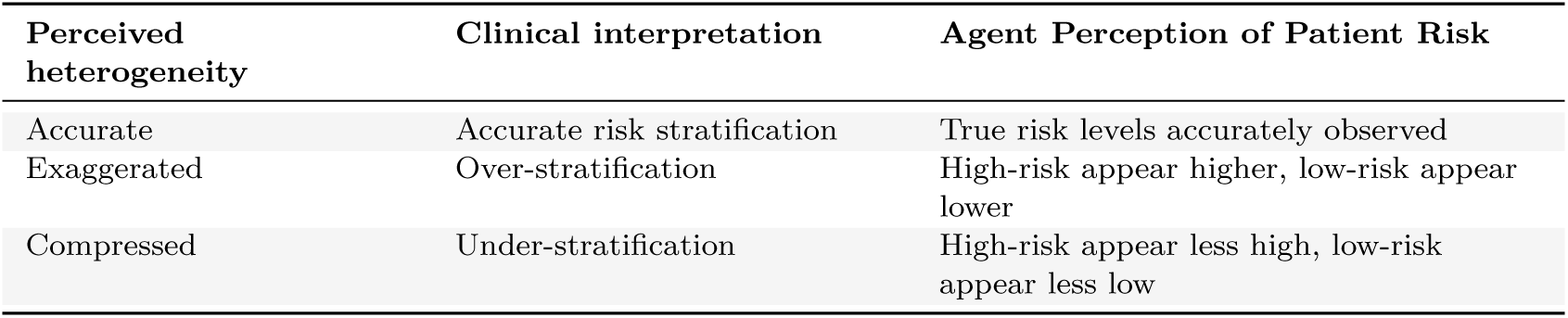
Experimental conditions for perceived vs. true patient heterogeneity in Experiment Set 3. Each of these three risk stratification conditions was tested across all three antibiotic scenarios (single-antibiotic, two-antibiotic without cross-resistance, two-antibiotic with cross-resistance). All experiments use mixed low-risk and high-risk heterogeneous populations as ground truth.

| Perceived heterogeneity | Clinical interpretation | Agent Perception of Patient Risk |
| --- | --- | --- |
| Accurate | Accurate risk stratification | True risk levels accurately observed |
| Exaggerated | Over-stratification | High-risk appear higher, low-risk appear lower |
| Compressed | Under-stratification | High-risk appear less high, low-risk appear less low |

#### Experiment Set 4: Combined Observational Noise, Patient Heterogeneity

In the fourth and final set of experiments, we combine all sources of uncertainty from Experiment Sets 2 and 3: agents face noisy/biased patient observations alongside noisy/biased/delayed AMR surveillance. We also substantially expand patient heterogeneity beyond Experiment Set 3. Whereas Experiment Set 3 introduced Gaussian variation only in infection probability (with other treatment response parameters held constant within subpopulations), Experiment Set 4 samples all available patient attributes from Gaussian distributions. This creates both between-subpopulation differences (high-risk vs. low-risk mean parameter values) and within-subpopulation variation (*σ* = 0.1 for all attributes), more closely approximating real-world patient diversity.

Two additional design changes increase realism. First, we increase patient volume to 10 patients per timestep. Second, we implement differential observability: low-risk patients present with minimal workup (only 2 observable attributes: infection probability and spontaneous recovery probability), while high-risk patients undergo comprehensive assessment (all patient attributes observable). This mirrors real-world clinical triage, where different patients have differential observability based on risk assessment.

Because delayed AMR observations render the environment partially observable over time, we evaluated both hierarchical PPO and hierarchical recurrent PPO agents to assess the impact of adding memory to the agent. This experiment set allowed us to evaluate how increasing noise, bias in both information streams and delay in resistance information affected learned prescribing policies. We also then compared the performance of these learned policies to fixed prescribing rules that were operating under the same conditions of increased information degradation.

## 3 Results

### 3.1 Fixed Prescribing Rule: Expected Reward - Greedy Policy

We used the fixed prescribing rule “expected reward - greedy” as a benchmark for all three antibiotic scenarios under conditions of perfect observability. The fixed prescribing rule was rolled across 60 seed per scenario, as described previously. AMR trajectories (top panels) and cumulative clinical count metrics (bottom panels) are summarized in Figure 3. The cumulative clinical response plots allow us to evaluate antibiotic efficacy throughout the episode by plotting counts of sensitive vs. resistant infections treated per antibiotic; sustained separation, with sensitive counts above resistant counts, indicates preserved clinical utility over the episode.

- Single-antibiotic scenario: ‘Expected reward - greedy’ policy induces threshold-like prescribing that stabilizes AMR near an equilibrium of approximately 0.327 (Figure 3, left column).
- Two-antibiotic scenario (no cross-resistance): ‘Expected reward - greedy’ policy converges to essentially the same equilibria for each antibiotic: 0.327 for antibiotic A and 0.326 for antibiotic B (Figure 3, middle column).
- Two-antibiotic scenario (with cross-resistance): The ‘Expected reward - greedy’ policy again converges to essentially the same equilibria for each antibiotic: 0.326 for antibiotic A and 0.327 for antibiotic B (Figure 3, right column).

**Figure 3:**
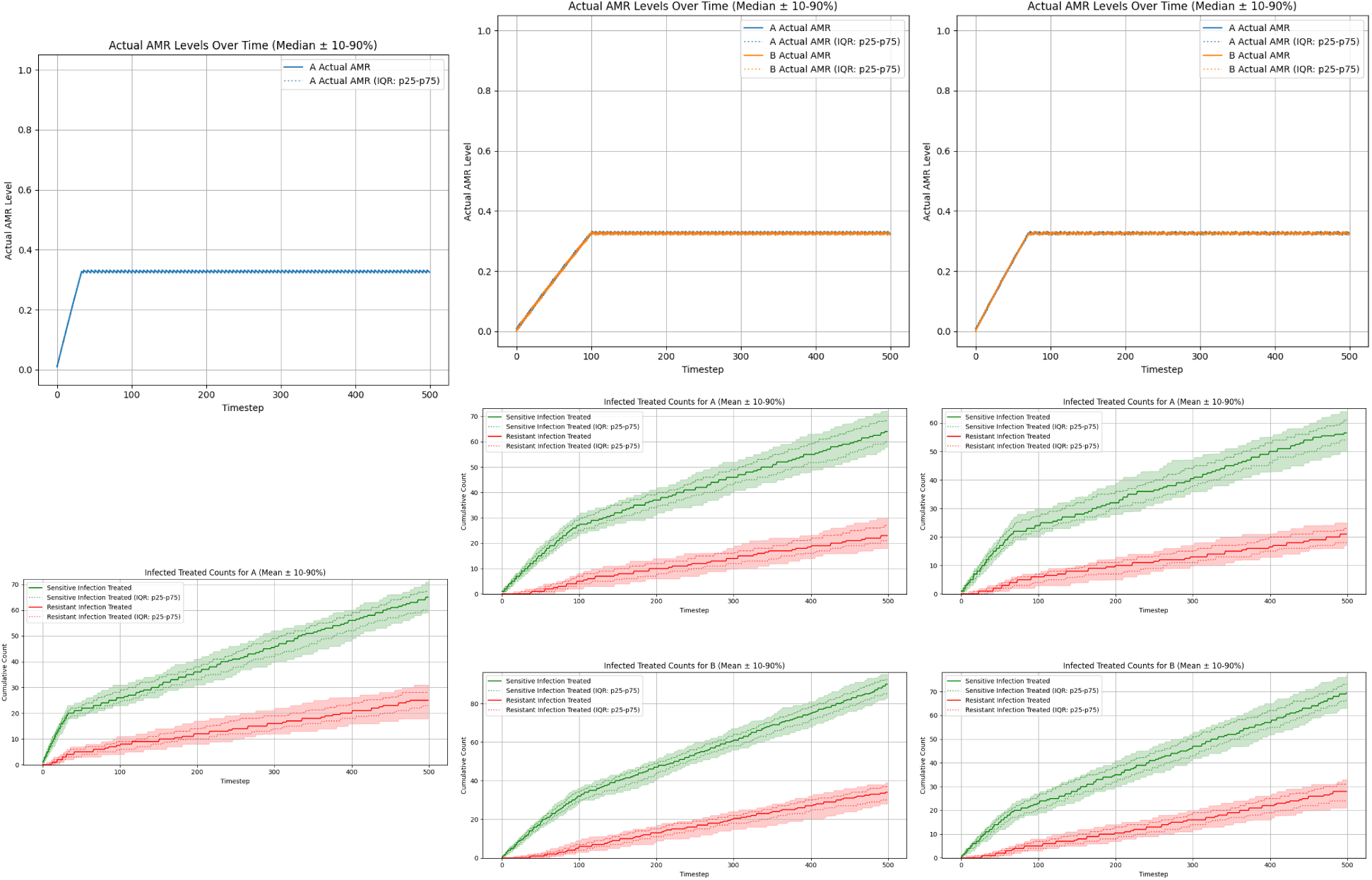
AMR levels evolution and cumulative count of sensitive vs. resistant infections treated per antibiotic, when ‘Expected reward - greedy’ policy is followed (where policy is computed using perfectly observed environments). Left: single-antibiotic scenario. Middle: two-antibiotic scenario without cross-resistance. Right: two-antibiotic scenario with cross-resistance. Each column shows evolution of true AMR levels (top) and cumulative infected/treated counts per antibiotic over time (bottom rows).

### 3.2 Experiment Set #1: Perfect Observability

Under perfect observability, we first asked a basic feasibility question: can reinforcement learning recover reasonable, near-optimal prescribing policies when all patient attributes and AMR levels are fully and accurately observed at every timestep? This best-case setting removes information degradation and therefore isolates the core learning challenge in these environments—long-horizon credit assignment.

Flat PPO showed only limited success in this setting. In each of the three different scenarios, it learned policies that stabilized AMR levels at steady-steate equilibria, but the learned equilibrium levels varied widely across seeds and did not consistently match the equilibria levels discovered by the ‘Expected Reward - Greedy’ fixed prescribing rule. In both two-antibiotic scenarios, the flat PPO performed particularly poorly: equilibrium AMR levels were highly variable across seeds, and cumulative sensitive-versus-resistant treated counts showed weaker separation, indicating poorer preservation of antibiotic efficacy over the episode (Figure 4, middle and right columns).

**Figure 4:**
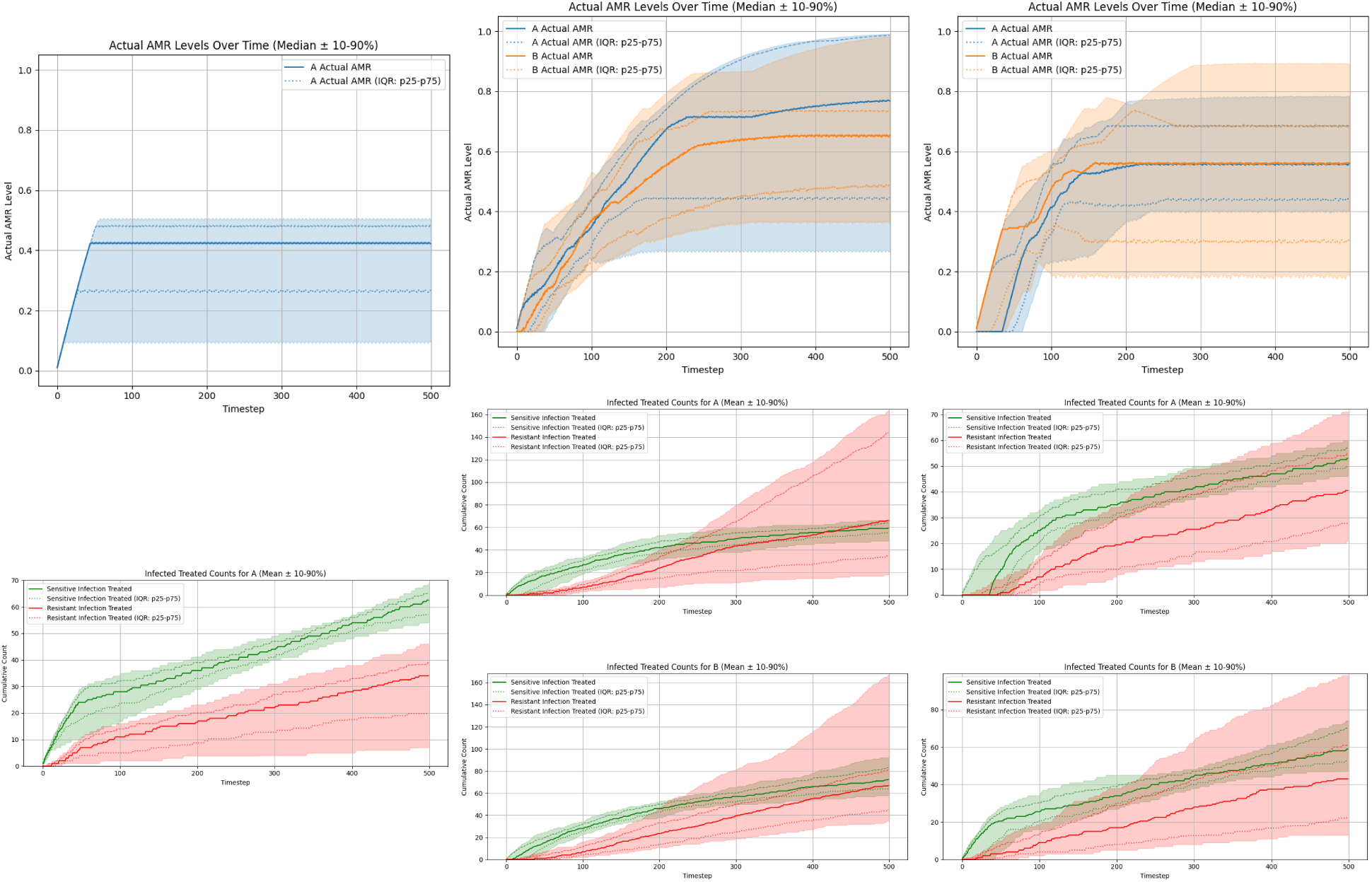
AMR levels evolution and cumulative count of sensitive vs. resistant infections treated per antibiotic, when tuned Flat PPO agent is trained in perfectly observed environment and then has evaluative rollouts performed. Left: single-antibiotic scenario. Middle: two-antibiotic scenario without cross-resistance. Right: two-antibiotic scenario with cross-resistance. Each column shows evolution of true AMR levels (top) and cumulative infected/treated counts per antibiotic over time (bottom rows).

**Figure 5:**
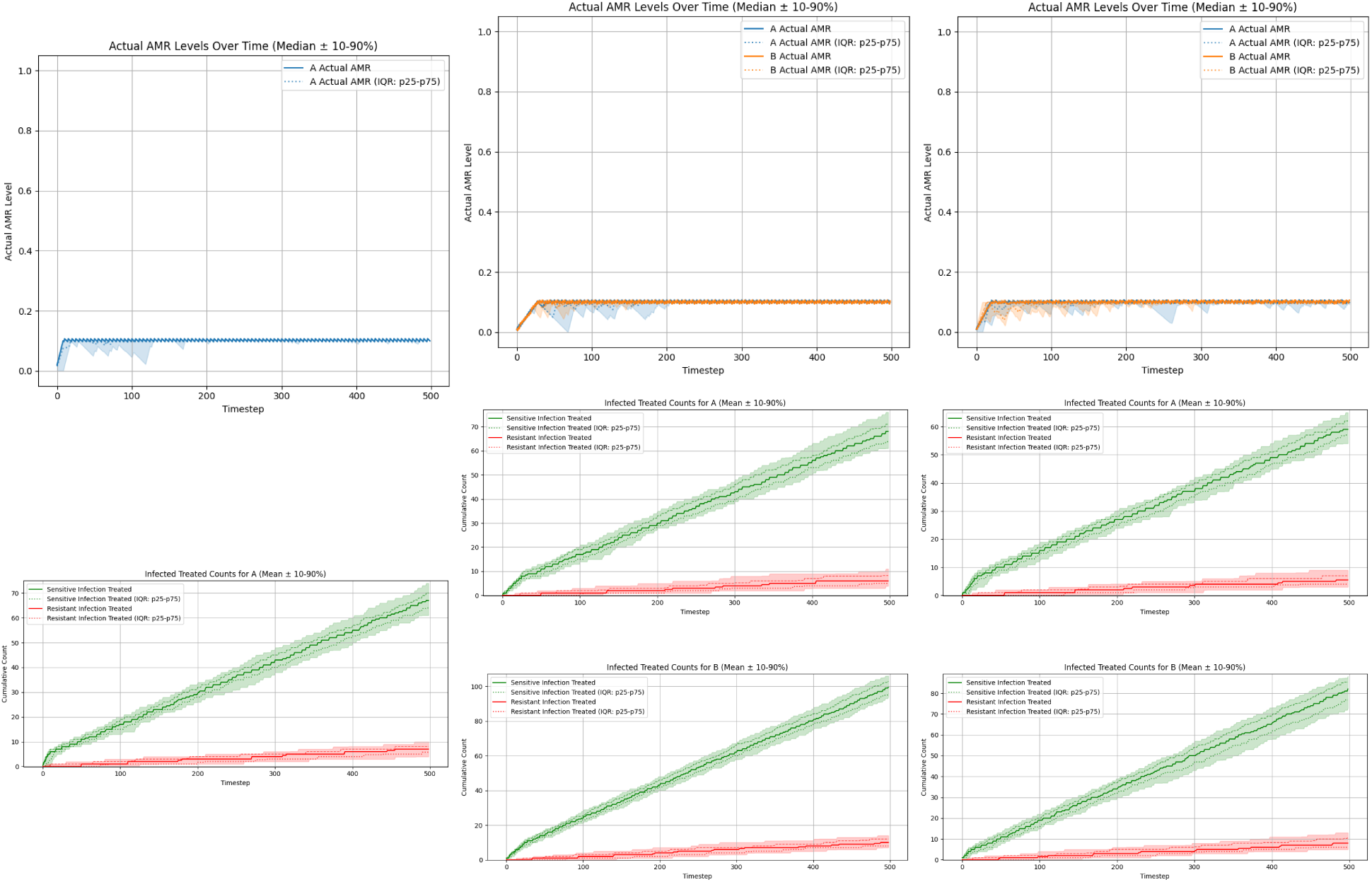
AMR levels evolution and cumulative count of sensitive vs. resistant infections treated per antibiotic, when tuned Hierarchical PPO agent is trained in perfectly observed environment and then has evaluative rollouts performed. Left: single-antibiotic scenario. Middle: two-antibiotic scenario without cross-resistance. Right: two-antibiotic scenario with cross-resistance. Each column shows evolution of true AMR levels (top) and cumulative infected/treated counts per antibiotic over time (bottom rows).

Hierarchical PPO demonstrated much more robust learning across all three scenarios, with limited cross-seed variance. Interestingly, the policies discovered by hierarchical PPO discover significantly *lower* AMR equilibria levels compared to the ‘Expected Reward - Greedy’ fixed prescribing rule. We can see that the policies discovered by hierarchical PPO do slightly outperform the fixed prescribing rule, although the advantage is relatively small, particularly for the single antibiotic scenario (Figure 6).

**Figure 6:**
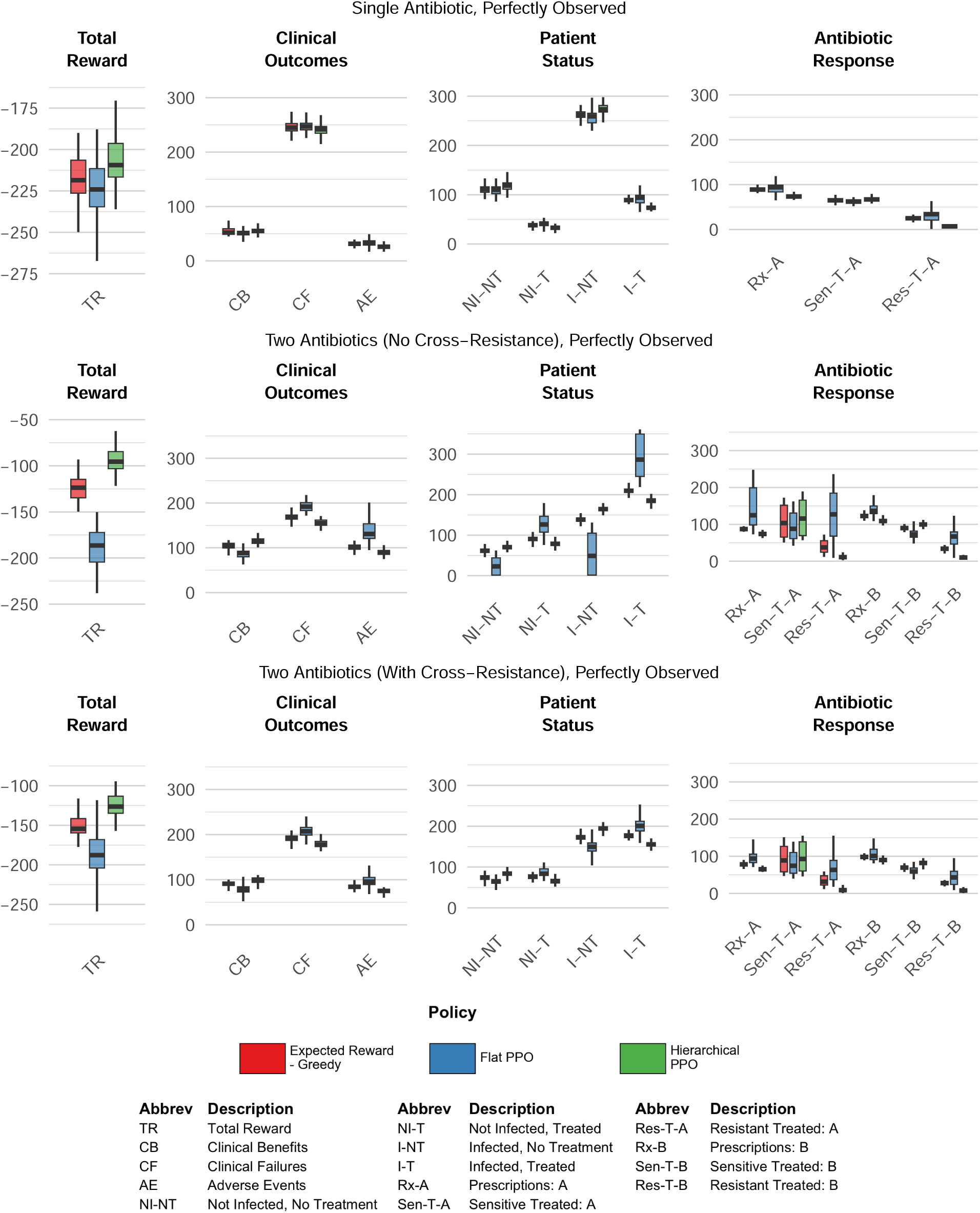
Comparison of cumulative performance metrics of Fixed Prescribing Rule (Expected Reward-Greedy) vs. Flat PPO vs. Hierarchical PPO in perfectly observed environment.

### 3.3 Experiment Set #2: Delayed, Noisy, Biased AMR

In Experiment Set 2, policy performance is evaluated when AMR surveillance is partially observable: AMR levels are updated every 90 timesteps, and the visible AMR signal is both noisy and biased (noise = 0.2, bias = −0.2). Figure 7 compares four PPO-based agents (flat memoryless, flat recurrent, hierarchical memoryless, hierarchical recurrent) against the ‘Expected Reward - Greedy’ fixed prescribing rule benchmark across all three antibiotic scenarios.

**Figure 7:**
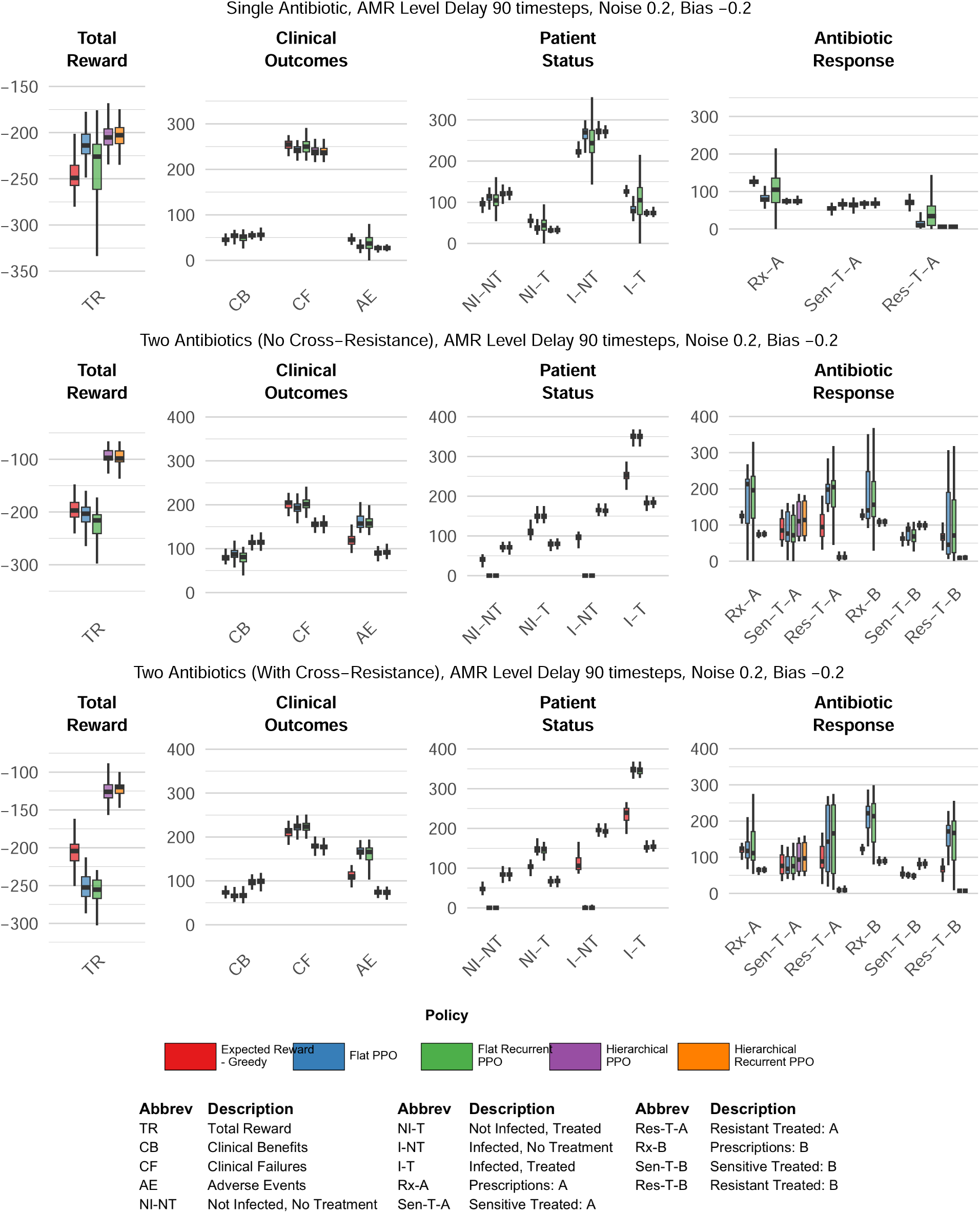
Comparison of cumulative performance metrics of Fixed Prescribing Rule (Expected Reward-Greedy) vs. Flat PPO vs. Flat Recurrent PPO vs. Hierarchical PPO vs. Hierarchical Recurrent PPO, in environment with degraded AMR level information. AMR levels affected by update delay = 90 timesteps, noise = 0.2, bias = −0.2.

Among the flat agents, adding recurrent memory did not improve performance. The memoryless flat PPO agent outperformed the recurrent flat PPO agent in the single-antibiotic and two-antibiotic no cross-resistance scenarios, with similar performance in the cross-resistance setting. These differences were accompanied by distinct prescribing patterns. In the single-antibiotic scenario, the memoryless flat agent concentrated prescribing shortly after each AMR observation update and then largely ceased prescribing until the next update, whereas the recurrent flat agent reduced prescribing during stale-information periods but did not stop as completely. In the two-antibiotic scenario without cross-resistance, this same pattern appeared as relatively clean on/off alternation between antibiotics A and B for the memoryless flat agent, whereas the recurrent flat agent reduced use of the disfavored antibiotic without fully discontinuing it. When cross-resistance was introduced, the distinction between flat memoryless and recurrent prescribing patterns became less pronounced. Detailed prescribing time series and AMR trajectories for these agents are provided in Supplementary Materials S4.

The hierarchical agents demonstrated qualitatively different behavior; even in the setting of degraded information regarding AMR levels, they were able to essentially recapitulate their performance from the perfectly observable versions of the three different scenarios. Adding memory to the hierarchical PPO does yield moderately improved performance, mostly due to the HRL recurrent PPO agents driving more stable trajectories compared to the HRL memoryless PPO agents.

Figure 7 illustrates that under conditions of degraded information for AMR levels, the hierarchical PPO agents show a clear performance advantage over both the flat PPO agents and the ‘Expected Reward-Greedy’ fixed prescribing rule.

### 3.4 Experiment Set #3: Heterogeneous Patient Attributes, Varied Bias

In Experiment Set 3, we isolate the effect of patient-level perception error by introducing bias in observed patient attributes while keeping AMR observations perfect and up to date. We modeled two subpopulations (high-risk and low-risk) sampled with equal probability, with subpopulation parameters chosen so that population-average attribute values match the homogeneous baseline used in Experiment Set 1, allowing direct comparison to the VI benchmark.

Figure 8 shows that introducing true risk heterogeneity to the patient population allows the hierarchial PPO agents to substantially improves both clinical outcomes and antibiotic stewardship relative to the homogeneous baseline even in the perfectly observed environment (Experiment Set 1). The ‘Expected Reward-Greedy’ fixed prescribing rule does not show the same universal boost in performance; in some scenarios under certain risk stratification conditions, the ‘Expected Reward - Greedy’ fixed prescribing rule outperformed the corresponding scenario in Experiment Set 1, in others it underperformed. (See Supplementary Materials for full performance metrics and tables.)

**Figure 8:**
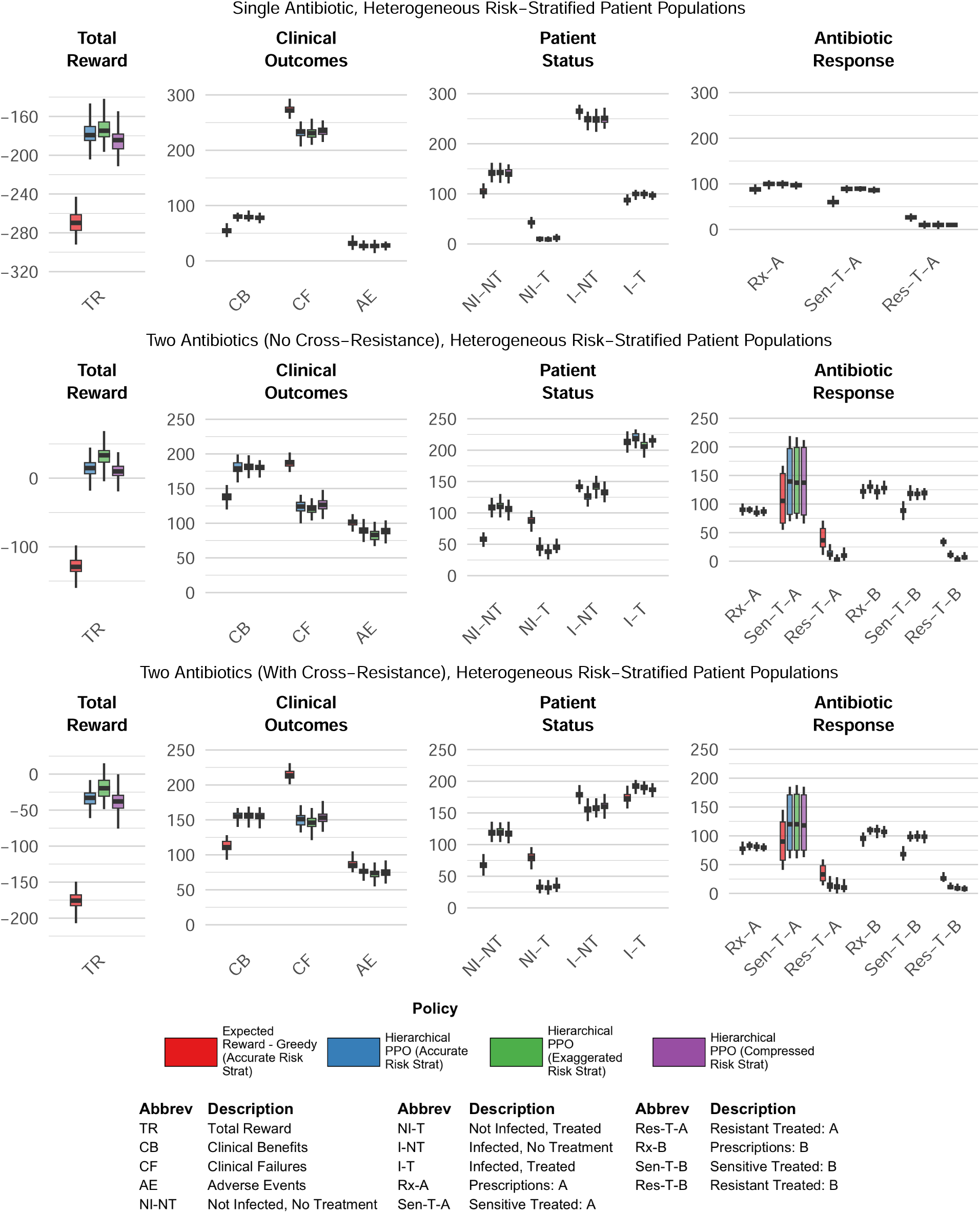
Comparison of cumulative performance metrics of Expected Reward - Greedy (Accurate Risk Stratification) vs. Hierarchical PPO (Accurate Risk Stratification Environment) vs. Hierarchical PPO (Exaggerated Risk Stratification Environment) vs. Hierarchical PPO (Compressed Risk Stratification Environment), in environments with heterogeneous risk-stratified patient population with varying levels of bias.

Figure 8 compares the performance of the ‘Expected Reward - Greedy’ fixed prescribing rule under ‘accurate risk stratification’ conditions, vs. HRL PPO agents trained under ‘accurate risk stratification’, ‘exaggerated risk stratification’, and ‘compressed risk stratification’ conditions respectively. Interestingly, for the HRL PPO agents, performance under exaggerated risk stratification was modestly better than accurate stratification across all three scenarios. Performance under compressed risk stratification was slightly worse than accurate stratification, though substantially better than the homogeneous population VI policy baseline.

The largest performance gains across all stratification conditions appeared in the two-antibiotic scenario without cross-resistance (Figure 8, middle panels). In the cross-resistance scenario, these gains were attenuated relative to the no cross-resistance case (Figure 8, right panels).

### 3.5 Experiment Set #4: Combined Observational Noise, Patient Heterogeneity

In Experiment Set 4, we combined both sources of uncertainty from Experiment Sets 2 and 3 and also increase patient volume: noisy and biased patient observations, delayed and noisy AMR surveillance, and ten patients per timestep. We then compared the performance of hierarchical PPO and hierarchical recurrent PPO against the fixed prescribing rule described in Section 3.6.

Both hierarchical architectures adopted markedly more conservative prescribing strategies than the “Expected Reward - Greedy” fixed prescribing rule benchmark across all three antibiotic scenarios (Figure 10). Under the conditions of combined information degradation and increased patient volume, the “Expected Reward - Greedy” fixed prescribing rule showed it was no longer able to maintain steady-state equilibria for AMR levels, but instead exhibited monotonically increasing AMR levels over the episode, leading to depleted clinical efficacy of existing antibiotics, as demonstrated in Figure 9.

**Figure 9:**
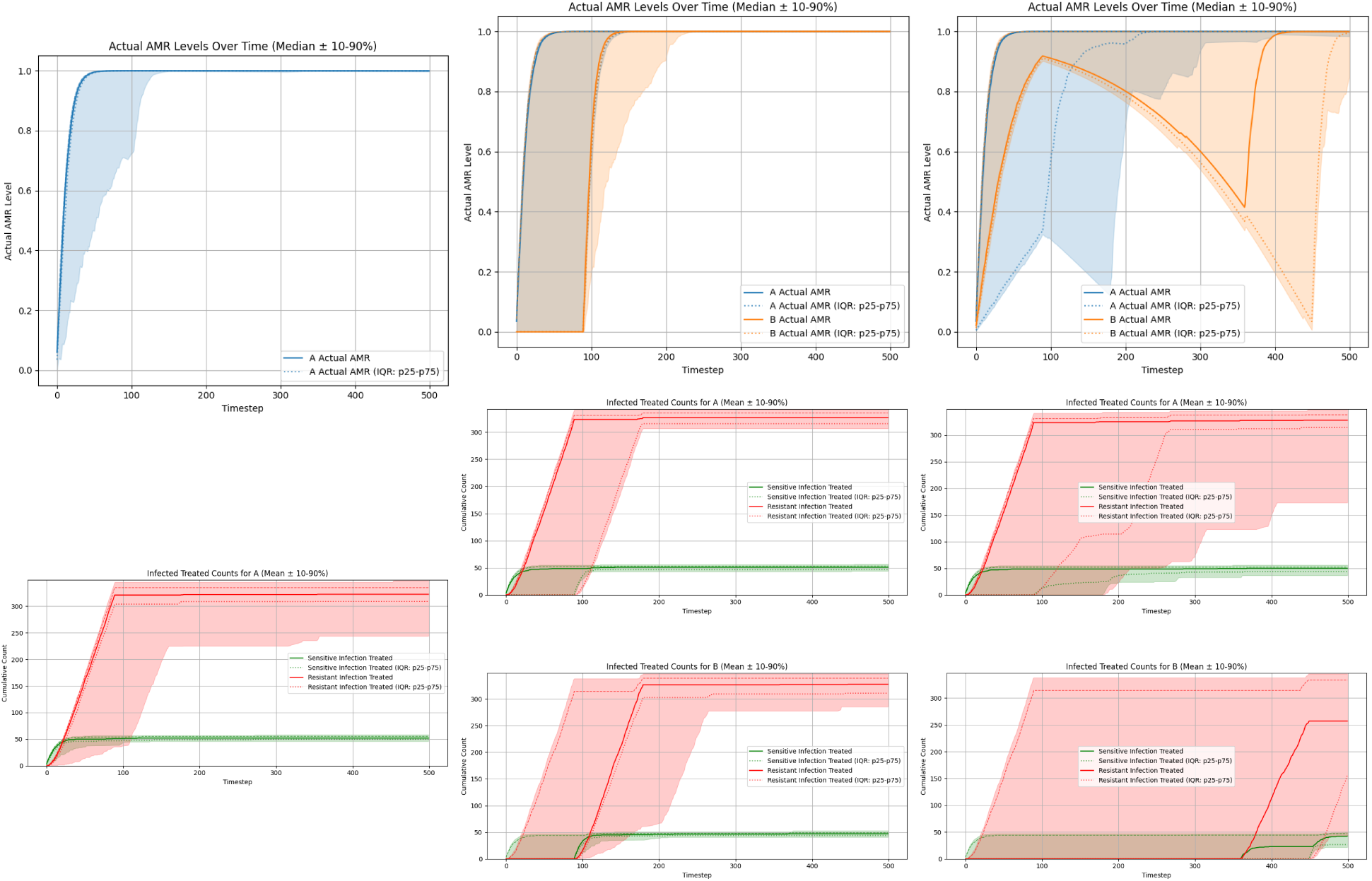
AMR levels evolution and cumulative count of sensitive vs. resistant infections treated per antibiotic, when ‘Expected reward - greedy’ policy is followed under experiment set 4 conditions (heterogeneous noisy risk-stratified mixed visibility patients, noisy and delayed AMR levels, 10 patients per timestep). Left: single-antibiotic scenario. Middle: two-antibiotic scenario without cross-resistance. Right: two-antibiotic scenario with cross-resistance. Each column shows evolution of true AMR levels (top) and cumulative infected/treated counts per antibiotic over time (bottom rows).

**Figure 10:**
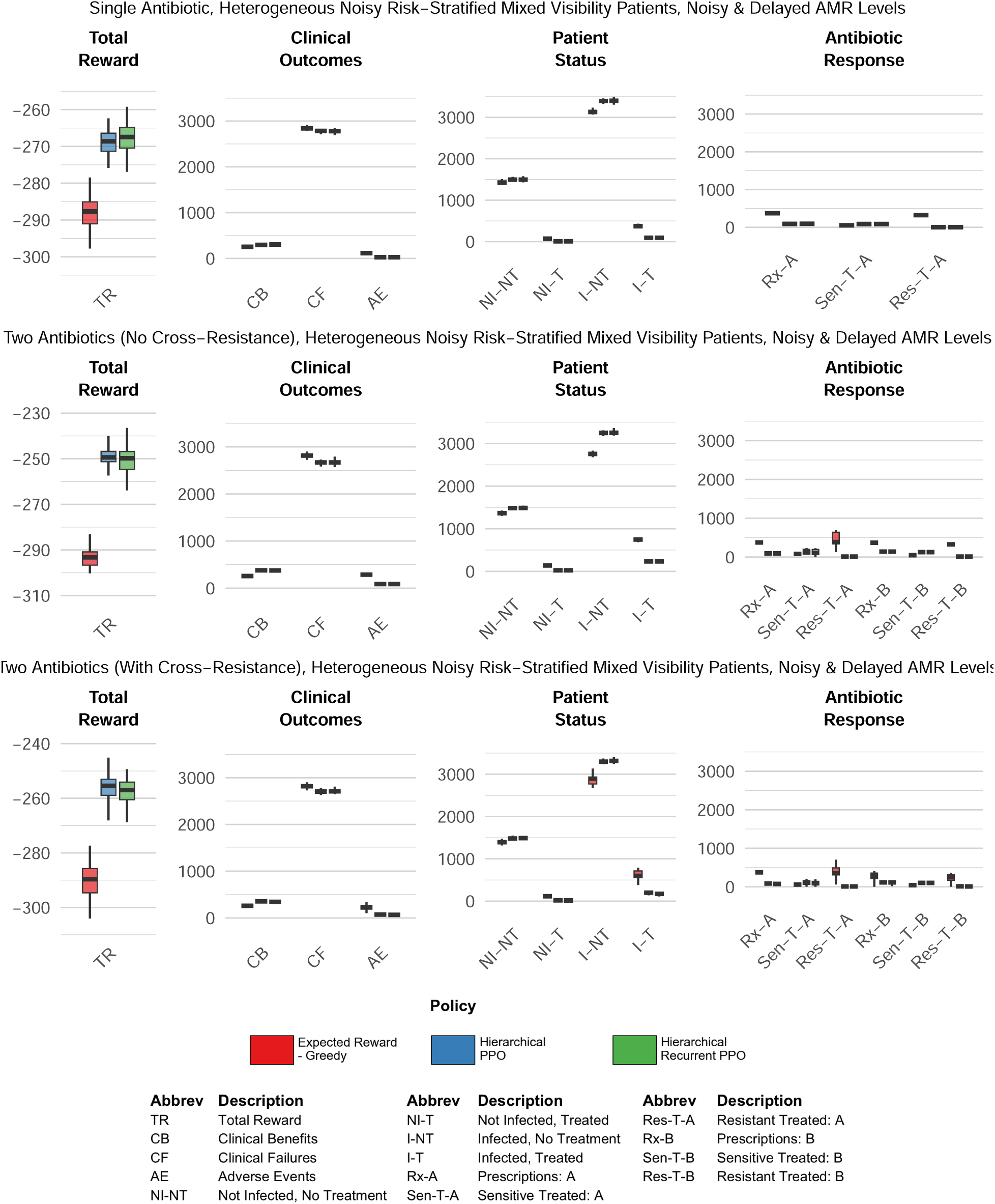
Comparison of cumulative performance metrics of Fixed Prescribing Rule (Expected Reward-Greedy) vs. Hierarchical PPO vs. Hierarchical Recurrent PPO, in environments with heterogeneous noisy risk-stratified mixed visibility patient populations (noise=0.1), noisy and delayed AMR Levels (delay = 90 timesteps, noise=0.2).

In this set of experiments, the performance advantage of the hierarchical RL agents over the ‘Expected Reward - Greedy’ fixed prescribing rule is even more pronounced than in the prior sets of experiments. The hierarchical policies were not merely trading off short-term clinical outcomes for long-term stewardship: they achieved more clinical successes, fewer clinical failures, and significantly lower long-term AMR levels compared to the “Expected Reward - Greedy” fixed rule comparator.

Figure 10 compares the cumulative performance metrics of the “Expected Reward - Greedy” fixed prescribing rule to the hierarchical PPO agents, both memoryless and recurrent. The hierarchical memoryless PPO appears to slightly outperform the hierarchical current PPO in both two-antibiotic scenarios, although the advantage is slight.

## 4 Discussion

Across four experiment sets with varying kinds and degrees of information degradation, we found that reinforcement learning agents can discover good antibiotic prescribing policies that dominate/outperform a clinically-realistic fixed prescribing rule, and that the learned policies by the RL agents are fairly robust against various types of noise.

### 4.1 Individual Experiment Set Findings

#### Experiment Set 1 (Perfect Observability)

Under conditions of perfect observability and one patient per time step, the performance of the ‘Expected Reward - Greedy’ fixed prescribing rule and the HRL PPO agent were largely comparable, although the HRL PPO agent was able to modestly outperform the fixed prescribing rule in the two-antibiotic scenarios. In contrast, the flat PPO agent performed adequately in the single-antibiotic scenario, but distinctly failed to learn effective policies in both two-antibiotic settings, even with complete state information. This suggests the failure mode is not informational but architectural: flat agents struggle with the long-horizon credit assignment required when prescribing decisions have delayed, coupled effects on future resistance levels. Switching to a hierarchical architecture — which decomposes the long-horizon problem into sequences of macro-actions — proved sufficient to recover competitive performance across all three scenarios.

The comparable performance metrics between the fixed prescribing rule policy and the HRL PPO policy in this set of experiments suggest that when providers have perfect information, non-learning fixed rules can actually perform fairly well, and that the marginal advantage of adaptive learning offered by reinforcement learning agents is fairly small.

#### Experiment Set 2 (Delayed, Noisy, Biased AMR)

Under delayed and noisy AMR surveillance, the performance of the ‘Expected Reward - Greedy’ fixed prescribing rule and the flat PPO now degrade significantly, although again the fixed prescribing rule actually outperforms the flat PPO in both of the two-antibiotic scenarios. In contrast, the HRL PPO agents are able to essentially recapitulate their performance from the perfectly observable environment, proving that they are robust to the level of AMR information degradation tested in this set of experiments. Interestingly, adding memory to the HRL PPO architecture did not appear to have a significant effect, as the performance of the HRL memoryless PPO and HRL recurrent PPO agents were largely similar across all three different antibiotic scenarios. Our initial hypothesis had been that under conditions of delayed information about AMR levels, that adding memory would allow for a performance boost due to the agent being able to persist an internal belief state about the true latent dynamics of the environment; however, this was not borne out by the results of the experiments run under the conditions of this particular experiment set.

#### Experiment Set 3 (Heterogeneous Patient Attributes, Varied Bias)

In Experiment Set 3, we can now observe that the marginal advantage of RL agents over the fixed prescribing rule even more pronounced compared to the results of Experiment Set 2. Observable patient heterogeneity now provides reliable triage signals, enabling selective treatment of high-risk patients while confidently withholding treatment from low-risk ones. This is a fundamental contrast to the homogeneous populations in Experiment Sets 1 and 2, where all patients shared identical observable attributes and the agent had no basis for differential treatment decisions. Interestingly, it appears that the HRL PPO agents are able to better exploit the patient heterogeneity to greater advantage compared to the fixed prescribing rules. The performance metrics for the fixed prescribing rule in this set of experiments are either worse or similar to the performance metrics for the fixed prescribing rule in Experiment Set 1 under conditions of perfect observability; this is in contrast to the performance achieved by the HRL PPO agents, which are able to significantly improve on the performance that they achieved in Experiment Set 1, and notably in the two-antibiotic cross-resistance scenario, the HRL agents are actually able to achieve an average positive overall total reward. It appears that this difference in performance is driven by the HRL agents being more selective about triaging patients by risk stratification in order to inform their decision to treat, leading to the HRL agents’ policies resulting in lower equilibrium AMR levels and higher long-term clinical efficacy for existing antibiotics.

The finding that exaggerated risk stratification modestly outperformed accurate stratification is somewhat surprising, but upon further reflection this finding even further emphasize the importance of triage. We think the likely mechanism is that stronger perceived separation between risk groups increases the agent’s confidence in withholding treatment from low-risk patients, further reducing unnecessary prescribing and preserving antibiotic efficacy. Compressed risk stratification produced only modest degradation in performance, suggesting that the gains from heterogeneity are fairly robust to miscalibration as long as relative risk ordering is preserved — though the direction of miscalibration matters, with over-stratification preferable to under-stratification in the scenarios tested here.

The largest gains across stratification conditions appeared in the two-antibiotic scenario with no cross-resistance. Here, risk-stratified treatment can be combined with antibiotic rotation: resting one antibiotic allows its resistance level to decay while the other is used, compounding the efficacy-preservation benefit of selective treatment. In the cross-resistance scenario these gains are attenuated, since prescribing one antibiotic also drives up resistance to the other, reducing the benefit of rotation.

#### Experiment Set 4 (Combined Observational Noise, Patient Heterogeneity)

In the most complex scenarios tested in this study, hierarchical PPO agents even more strongly dominate the fixed prescribing rule benchmark, outperforming the benchmark across both clinical and stewardship metrics. Under conditions of combined information degradation and increased patient volume, the “Expected Reward - Greedy” fixed prescribing rule was no longer able to maintain lower steady-state equilibria for AMR levels, but instead ‘saturated’ them by driving the AMR levels up towards 100%, leading to depleted clinical efficacy of the existing antibiotics. We think a likely mechanism for this catastrophic failure of the fixed prescribing rule under these conditions is due to the increased patient volume at each timestep; in the prior experiment sets, the fixed prescribing rule only had to treat one patient per timestep, meaning that its ability to drive up AMR levels too high before feeling the consequences was limited. In this set of experiments with 10 patients per timesteps, this natural bottleneck no longer exists, leading to the failure of the fixed prescribing rule to maintain long-term clinical efficacy of the antibiotics.

In contrast, both the HRL memoryless PPO and HRL recurrent PPO agents converted to conservative low-AMR equilibria that preserved efficacy of existing antibiotics throughout the episode. Again, adding memory did not appear to offer a clear performance advantage, and in fact the HRL memoryless PPO agent slightly outperformed the HRL recurrent PPO agent in the two-antibiotic scenarios. We suspect that under the conditions we tested in this set of experiments, the ability to maintain an internal belief state about the world was essentially balanced by the cost of imperfect memory-based inference, but that there are likely other experimental conditions where adding memory would offer a clearer performance advantage.

### 4.2 Cross-Experiment Set Findings

Across all four experiment sets, the results confirm our central hypothesis: reinforcement learning agents can successfully learn antibiotic prescribing policies that perform comparably or dominate static benchmark polices, and furthermore the learned policies are robust to various types of information degradation. However, the architecture of the RL agents matters: flat agents were insufficient for all but the simplest single-antibiotic scenario; hierarchical RL with temporal abstraction was necessary to address the long-horizon credit assignment structure inherent in the antibiotic prescribing problem. We make no claim that hierarchical RL is the only viable strategy for these environments, nor that it would scale without modification to substantially more complex simulations, i.e. more antibiotics, larger patient cohorts, or richer resistance dynamics. Nevertheless, even within these relatively controlled scenarios, the experiments surfaced several non-obvious insights: notably, that over-stratification of patient risk may yield better aggregate outcomes than accurate risk calibration, and that clinically-realistic prescribing practices can lead to catastrophic failure by either completely or almost completely depleting the clinical efficacy of existing antibiotics.

We note here that the RL agents in these experiments achieved these outcomes despite being optimized using only individual clinical reward (*λ* = 0, with no explicit penalty for increasing community-level antimicrobial resistance). This indicates that when the environment itself creates long-term consequences for short-sighted actions — as antimicrobial resistance dynamics do — appropriately designed agent architectures can learn farsighted prescribing strategies without requiring explicit stewardship reward shaping. The consistent performance advantage of hierarchical over flat agents underscores the importance of policy architecture in settings where treatment decisions have delayed, coupled effects on future treatment efficacy.

### 4.3 Limitations

#### 4.3.1 Abstraction of pathogen identity

Our simulation environment necessarily represents a simplified and idealized abstraction of real-world antibiotic prescribing and antimicrobial resistance dynamics. The most significant simplification involves our treatment of pathogens: we deliberately abstract away individual pathogen identity, even though in clinical practice pathogens serve as the natural unit for measuring antimicrobial resistance. Real-world antibiograms stratify resistance by pathogen species, tallying sensitive and resistant isolates separately for each organism; our simulator omits this pathogen-level granularity entirely.

We made this choice deliberately in order to distill clinical decision-making regarding antibiotic therapy to its most essential elements: (1) the probability that a given patient has a bacterial infection requiring antibiotic treatment, (2) the likelihood that a specific antibiotic will effectively treat that infection if present, and (3) the clinical trade-offs associated with prescribing. By reducing the decision to these core questions, we gain interpretability — both in how patients and AMR levels are represented, and in how the policies learned by RL agents can be understood. We also hypothesize that this simplification likely represents a theoretical upper bound on achievable clinical benefit; in a more stochastic, multi-pathogen environment, policy performance would likely degrade further.

#### 4.3.2 Stationarity assumptions

In this study, we model both the patient population and the antimicrobial resistance response curve for each antibiotic as stationary entities — that is, patient attribute distributions and resistance dynamics remain constant throughout each episode and across episodes. In reality, neither is static: patient populations shift over time due to demographic change, seasonal variation in infection prevalence, and emergence of high-risk subpopulations, while resistance mechanisms can evolve in response to sustained selection pressure in ways that may be partially or fully irreversible. Policies learned under stationary assumptions may therefore be poorly calibrated to real-world settings where the underlying dynamics are drifting, and our results should be interpreted accordingly as characterizing performance under idealized stable conditions. Extensions to non-stationary dynamics are discussed in the Future Work section.

#### 4.3.3 Single centralized prescriber

The simulator models a single, centralized prescriber with complete information about all patients and system-level AMR dynamics. This is unrealistic: real-world antibiotic prescribing involves multiple independent clinicians with limited visibility into their peers’ prescribing patterns and heterogeneous access to surveillance data. We plan to address this limitation in future work by extending the framework to support multi-agent, multi-locale environments where individual prescribers optimize locally with incomplete information (see Section 5.4).

#### 4.3.4 Reinforcement learning algorithm selection

PPO was selected primarily for pragmatic reasons: it natively supports multidiscrete action spaces, as required for our multi-patient, multi-antibiotic setup, and is computationally tractable for the experimental scope of this study. Other RL algorithms or agent architectures may yield different performance characteristics. The modular design of the abx_amr_simulator package — which conforms to the Gymnasium API — allows interested users to straightforwardly implement and evaluate alternative agent classes beyond those tested here.

#### 4.3.5 AMR resistance dynamic assumptions

The AMR_LeakyBalloon model makes specific assumptions about the shape of the antibiotic resistance response curve: latent resistance pressure is mapped to an observable AMR level through a sigmoid function, yielding a smooth, bounded accumulation and decay dynamic. While we believe this is a reasonable abstraction of resistance emergence and relief, the true functional form of resistance dynamics in real bacterial populations is not well characterized at the community level, and alternative response curve shapes could meaningfully alter the prescribing strategies learned by RL agents.

Additionally, our current implementation models cross-resistance as strictly positive — prescribing antibiotic A increases resistance to antibiotic B — whereas certain antibiotic combinations in fact exhibit collateral sensitivity, where use of one agent decreases resistance to another (Wang et al. 2025). Incorporating negative cross-resistance could meaningfully alter optimal prescribing strategies, potentially favoring cycling or mixing regimens that exploit these collateral effects.

Both of these limitations are mitigated by the modular architecture of the simulator. AMR dynamics are implemented through an abstract interface (AMRDynamicsBase), with AMR_LeakyBalloon as the concrete implementation used in this study. Users can provide alternative implementations of AMRDynamicsBase to specify different response curve shapes or cross-resistance structures, enabling systematic investigation of how resistance dynamic assumptions affect learned prescribing policies without requiring modification to the broader simulation framework.

#### 4.3.6 Scope of findings and generalizability

Because the simulator is intentionally abstract, findings should not be interpreted as direct clinical policy recommendations. The framework is most useful for directional analysis under controlled conditions — quantifying the effect of delayed, noisy, or biased antibiogram updates, biased patient-risk perception, and varying levels of patient-level information on prescribing decisions and the clinical-stewardship trade-off. Findings are hypothesis-generating and intended to inform the design of stewardship interventions and surveillance investments, not to prescribe specific clinical protocols.

### 4.4 Future Work

Several natural extensions of the abx_amr_simulator framework could substantially broaden its applicability and scientific scope.

#### 4.4.1 Multi-agent, multi-locale environments

We plan to extend the framework to support multiple agents operating across geographically distinct locales. In this configuration, each locale would represent a separate community with its own AMR dynamics, and patients could migrate between locales, potentially spreading resistant infections. Individual agents would represent clinicians or healthcare facilities with limited visibility into peers’ prescribing practices — observing only their own patient encounters and local antibiogram data. This extension would enable investigation of emergent dynamics in decentralized prescribing ecosystems, including how local optimization by individual agents affects community-level resistance, whether coordination mechanisms could improve collective outcomes, and how informational asymmetries across locales shape optimal strategies.

#### 4.4.2 Non-stationary dynamics

Real-world patient populations and resistance mechanisms evolve over time, yet our current model treats both as stationary. Extending the simulator to support non-stationary dynamics would enable investigation of clinically important questions: How frequently should prescribing guidelines be updated to remain effective? What signals indicate that a guideline has become outdated? How much avoidable harm accumulates when policies fail to adapt to shifting patient demographics or pathogen ecology? We plan to address this by implementing time-varying patient attribute distributions to model demographic shifts, seasonal variation in infection prevalence, and emergence of high-risk subpopulations. We also plan to extend the leaky-balloon resistance dynamics to support regime-dependent behavior — for instance, modeling scenarios where sustained intensive antibiotic use creates irreversible changes in resistance mechanisms, such that the balloon accumulates a permanent baseline pressure that cannot be fully relieved even with extended periods of antibiotic restriction.

#### 4.4.3 Upper bounds on personalized antibiogram prediction

Recent literature has explored machine learning approaches to predict patient-specific antibiotic susceptibilities from electronic medical record data, with the goal of improving empiric therapy selection (Al Mazrouei et al. 2025). While this body of work has grown substantially and spans a range of clinical settings, predictive models have generally achieved moderate discriminative performance (Sakagianni et al. 2023), leaving open a more fundamental question: even with perfect patient-level susceptibility prediction, how much collective clinical benefit could actually be achieved at the population level?

Our simulation framework is well positioned to address this question. By extending the patient model to include individual-level antibiotic susceptibility profiles, we can compare outcomes when RL agents have access to population-level antibiogram data alone versus perfect oracle knowledge of each patient’s true susceptibility — directly quantifying the theoretical ceiling on collective clinical benefit from personalized prediction under realistic conditions, including noisy observations, delayed resistance data, and heterogeneous patient populations. If this ceiling is high, it would justify continued investment in improving model performance despite currently modest discriminative metrics. If even perfect personalized prediction yields only marginal improvement over population-level antibiograms, it would suggest that resources might be more productively directed toward other stewardship interventions. We could further examine how this theoretical benefit varies across clinical contexts — including baseline resistance prevalence, population heterogeneity, and available antibiotic options — to identify settings where personalized prediction tools would provide the greatest value.

### 4.5 Conclusions

This study demonstrates how the abx_amr_simulator library can serve as a quantitative testbed for examining how information deficiencies and structural uncertainty shape antibiotic prescribing strategy performance under controlled conditions. Across four experiment sets of increasing complexity and worsening conditions of information degradation, we showed that hierarchical reinforcement learning agents can learn robust and effective prescribing policies that preserve antibiotic efficacy — outperforming a clinically-realistic fixed prescribing rule across clinical and stewardship metrics — without requiring explicit antimicrobial resistance penalties in the reward function. The framework supports directional analysis of factors including delayed or noisy antibiogram data, biased patient-risk perception, and differential access to patient-level clinical information, and their effects on the clinical-stewardship trade-off inherent in antibiotic prescribing. While findings are not intended as direct clinical guidance, they provide empirically grounded, hypothesis-generating insights that can inform stewardship program design, surveillance investment priorities, and the value of improving clinical risk assessment tools.

## Acknowledgments

The authors thank Daniel De la Rosa Martinez and Paula Weidemuller for their helpful feedback on earlier versions of this work.

## 5 Author Contributions

J.L.: Conceptualization, Methodology, Software, Formal Analysis, Writing – Original Draft, Writing – Review & Editing. S.B.: Funding Acquisition, Supervision, Writing – Review & Editing.

