## Supplementary material for "abx_amr_simulator: A simulation environment for antibiotic prescribing policy optimization under antimicrobial resistance": S1 Supplementary Material

### Supplementary Materials: Reinforcement Learning for Antibiotic Stewardship

Joyce Lee

2026-03-24

**S4 RL Agents & Fixed Experimental Rules: AMR Trajectories, Cumulative Clinical Outcomes, and Summary Statistics 7**

#### S1 Environment Subcomponents: `PatientGenerator`, `AMR_LeakyBalloon`, and `RewardCalculator`

The `abx_amr_simulator` package constructs the environment using the `ABXAMREnv` class, which serves as an umbrella class containing three primary subcomponents. At each timestep, `ABXAMREnv` samples patients via `PatientGenerator`, receives agent actions, updates antibiotic resistance state via `AMR_LeakyBalloon`, and computes scalar reward via `RewardCalculator`.

- `PatientGenerator`: This subcomponent creates synthetic patients with configurable attributes, where each attribute can be set to a constant value or sampled from a probability distribution, and where attributes can be marked as observable or hidden. It also generates a fixed number of patients at each timestep for agent evaluation.

- `RewardCalculator`: This subcomponent calculates the scalar reward that the agent receives for the decisions it makes at each timestep. See Section S2 for a full discussion of the reward function.
- `AMR_LeakyBalloon`: The `AMR_LeakyBalloon` model tracks how resistance levels for a given antibiotic evolve over time in response to prescribing patterns. The `ABXAMREnv` class contains an array of `AMR_LeakyBalloon` instances, one for each antibiotic.

Further details of the `abx_amr_simulator` package are discussed in the companion paper (see manuscript for citation), with full documentation available at [GitHub repository link].

#### S1.1 `PatientGenerator`: patient attributes

In the experiments presented in this study, each synthetic patient is represented by six attributes that jointly determine clinical profile and expected treatment outcomes:

**Probability of infection ($\pi_i$):** The likelihood that the patient has a bacterial infection requiring antimicrobial treatment. This serves as the primary risk stratification metric.

**Benefit value multiplier ($\phi_{b,i}$):** Scales the magnitude of clinical benefit reward if the patient is successfully treated with an effective antibiotic. Values greater than 1.0 represent patients who derive greater benefit from successful treatment (e.g., immunocompromised individuals).

**Benefit probability multiplier ($\omega_{b,i}$):** Scales the likelihood of achieving clinical benefit when an effective antibiotic is administered. Values less than 1.0 represent patients with comorbidities or factors that reduce treatment efficacy.

**Failure value multiplier ($\phi_{f,i}$):** Scales the magnitude of clinical failure penalty when the patient is inadequately treated (ineffective antibiotic or no treatment when infected). Higher values represent patients who would incur worse outcomes than baseline from treatment failure.

**Failure probability multiplier ($\omega_{f,i}$):** Scales the likelihood of experiencing clinical failure when inadequately treated.

**Recovery without treatment probability ($\rho_i$):** The probability that an infected patient will spontaneously recover without antimicrobial intervention, representing self-limiting infections.

These attributes can be configured to create either homogeneous populations (all patients share identical values) or heterogeneous populations (values drawn from Gaussian distributions). At each time step, the agent receives an observation vector for each patient containing some or all of these attributes. Thus, each attribute has a true underlying value, and an observed value; the user can choose to manipulate the observed value of any attribute by introducing a specific quantity of noise (random variance around true values) or bias (systematic over- or under-estimation), enabling controlled investigation of partial observability. At minimum, the agent observes an estimated probability of infection for each patient, though the true infection status remains latent.

#### S1.2 `AMR_LeakyBalloon`: antibiotic resistance dynamics

Antimicrobial resistance for each antibiotic is modeled using a soft-bounded accumulator with decay dynamics, implemented via a "leaky balloon" abstraction. Prescribing an antibiotic increases latent resistance pressure, while resistance decays over time in the absence of continued selection pressure. Latent pressure is mapped to an observable AMR level through a sigmoid function, yielding a value in [0,1] representing the probability that a newly occurring infection is resistant to that antibiotic at the current timestep. We refer to this observable AMR level as the "volume" of the leaky balloon and denote it by $\sigma_a$; specifically, $\sigma_a(t)$ is the volume for antibiotic $a$ at time $t$, equivalent to the probability that a current infection is resistant to antibiotic $a$.

Each antibiotic is characterized by its own response-curve parameters, controlling both the steepness of resistance emergence and the rate of resistance decay. The agent observes antibiotic-specific AMR levels as part of its observation state. As with patient attributes, users can modify observation fidelity by choosing how much noise, bias, or delay to introduce into the AMR levels available to the agent, enabling systematic exploration of partial observability in resistance surveillance.

Users can also specify whether cross-resistance exists between different antibiotics. In real-world settings, prescribing one antibiotic can increase resistance not only to that drug, but also to other related antibiotics. When creating an `abx_amr_simulator` environment, users can define cross-resistance strength between antibiotic pairs (e.g., how much prescribing antibiotic A increases resistance to antibiotic B).

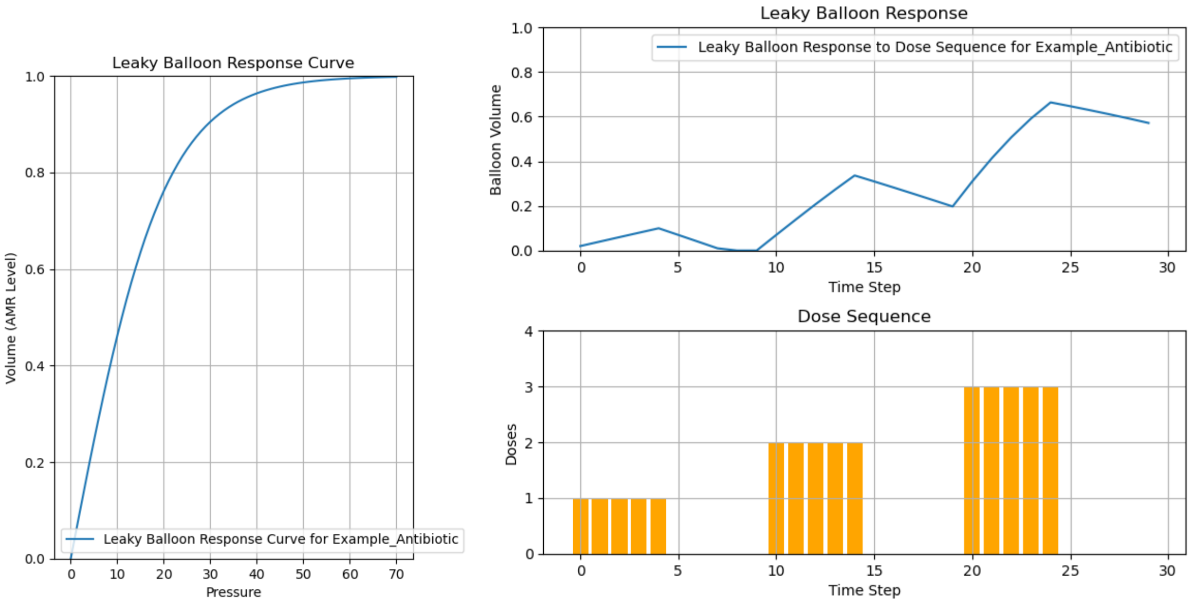

Figure S1: Pressure-volume response curve vs. dose sequence-volume response curve for example antibiotic.

##### S1.3 `RewardCalculator`: compute rewards for agent actions

`RewardCalculator` is the `ABXAMREnv` subcomponent responsible for computing the scalar reward the agent receives after each action. This reward signal drives learning by providing feedback on whether decisions improve or worsen outcomes. `RewardCalculator` implements the reward function discussed in detail in Section S2.

#### S2 Reward function

The reward function balances immediate clinical outcomes against long-term antimicrobial resistance considerations. We accomplish this by explicitly splitting the reward function into two parts, the 'individual reward' and the 'community reward'; the individual reward calculates the overall net clinical benefit to each patient 'treated' by the agent, while the 'community reward' encourages the agent to keep overall AMR levels for all existing antibiotics low. The overall reward function has the following form, where $\lambda$ is a tuneable weight $\in [0, 1]$ that allows the user to adjust how much to weight the individual reward vs. the community reward. The individual reward is averaged across patients and the community reward is averaged across antibiotics, creating comparable scales. This enables $\lambda$ to intuitively weight individual vs. community objectives.

$$R_{\text{overall}}(t) = (1 - \lambda) \cdot \langle \tilde{R}_{\text{individual}}(t) \rangle + \lambda \cdot \langle R_{\text{community}}(t) \rangle \tag{S1}$$

#### S2.1 Community reward

The community reward is calculated as the negative sum of the visible AMR levels for all the antibiotics in the environment (where $\mathcal{A}$ represents the set of existing antibiotics).

$$R_{\text{community}}(t) = -\sum_{a \in \mathcal{A}} \sigma_{a,\text{visible}}(t) \tag{S2}$$

The community reward uses **visible (observed) AMR levels**, maintaining consistency with the agent's observations. This design reflects clinical reality: clinicians receive feedback only from imperfect surveillance data (antibiograms, resistance patterns), not ground-truth AMR. By basing the community reward signal on observed AMR rather than true AMR, we ensure agents learn strategies that work with realistic information constraints. This POMDP-consistent approach is more clinically authentic than rewarding based on hidden ground truth.

Note that individual patient outcome rewards (clinical benefit/failure) still use ground-truth infection sensitivity, because patients either recover or don't based on reality. Only the AMR feedback channel uses observed data, reflecting how clinicians actually receive resistance information.

Given that $\sigma_a(t)$ has a maximum range of $[0, 1]$, $\sigma_a$ is not rescaled; we instead divide $R_{\text{community}}$ by the total number of antibiotics in the environment $|\mathcal{A}|$:

$$\langle R_{\text{community}} \rangle = \frac{R_{\text{community}}}{|\mathcal{A}|} \tag{S3}$$

#### S2.2 Individual reward

Individual rewards combine antibiotic and patient-specific parameters with discrete clinical outcomes. The environment samples Bernoulli random variables to generate clinical indicators (infection status, treatment success, adverse effects), which determine the realized reward for each decision.

##### S2.2.1 Prescribing antibiotic

The realized net individual reward for patient $i$ at time $t$ when the agent prescribes antibiotic $a$ (where $a \in \mathcal{A}$) is:

$$\begin{aligned} r_{i,a\in\mathcal{A}}(t) = \mathbf{1}\{\text{infected}\}\Big[&(1 - \mathbf{1}\{\text{resistant infection to } a\})\mathbf{1}\{\text{achieve clinical benefit reward}\}(\phi_{b,i}B) \\ &+ \mathbf{1}\{\text{resistant infection to } a\}\mathbf{1}\{\text{incur clinical failure penalty}\}(\phi_{f,i}F)\Big] \\ &+ \mathbf{1}\{\text{adverse effect from } a\}\, A_a \end{aligned} \tag{S4}$$

See the indicator variables table below for complete definitions of all parameters and their scope.

##### S2.2.2 No treatment

When the agent chooses no treatment, the individual reward depends on whether the patient spontaneously recovers:

$$
\begin{aligned}
r_{i,a=\text{no rx}}(t) = \mathbf{1}\{\text{infected}\}\Big[& \\
&\mathbf{1}\{\text{recovery without treatment}\}(\phi_{b,i}B) \\
&+ (1-\mathbf{1}\{\text{recovery without treatment}\})\mathbf{1}\{\text{incur clinical failure penalty}\}(\phi_{f,i}F)\Big]
\end{aligned} \tag{S5}
$$

where $\rho_i$ is the individual's probability of recovery without treatment. When a patient spontaneously recovers despite not receiving treatment, they receive the clinical benefit reward. When they do not recover spontaneously, they may incur a clinical failure penalty if the failure outcome occurs. We model spontaneous recovery as a separate stochastic event because it reflects clinical reality: even without treatment, some infections resolve naturally, and clinicians account for this baseline probability when deciding whether to prescribe.

#### S2.3 Overview of indicator variables for reward function

All terms of the form $\mathbf{1}\{\cdot\}$ denote indicator variables taking values in $\{0,1\}$, sampled from Bernoulli distributions. The table below summarizes each indicator and the parameters that determine its probability:

Table S1: Indicator variables in the reward function

| **Indicator** | **Distribution** | **Probability inputs** | **Meaning** | **Scope** |
|---|---|---|---|---|
| $\mathbf{1}\{\text{infected}\}$ | Bernoulli($\pi_i$) | $\pi_i$ | Patient is infected | Patient-specific |
| $\mathbf{1}\{\text{resistant infection to } a\}$ | Bernoulli($\sigma_a(t)$) | $\sigma_a(t)$ | Infection is resistant to $a$ | Environment state |
| $\mathbf{1}\{\text{achieve clinical benefit reward}\}$ | Bernoulli($\omega_{b,i} \cdot P_{\text{benefit}}$) | $\omega_{b,i}$, $P_{\text{benefit}}$ | Clinical benefit is realized | Patient-specific + global |
| $\mathbf{1}\{\text{incur clinical failure penalty}\}$ | Bernoulli($\omega_{f,i} \cdot P_{\text{failure}}$) | $\omega_{f,i}$, $P_{\text{failure}}$ | Clinical failure penalty is realized | Patient-specific + global |
| $\mathbf{1}\{\text{adverse effect from } a\}$ | Bernoulli($\alpha_a$) | $\alpha_a$ | Adverse effect occurs | Antibiotic-specific |
| $\mathbf{1}\{\text{recovery without treatment}\}$ | Bernoulli($\rho_i$) | $\rho_i$ | Spontaneous recovery without treatment | Patient-specific |

**Parameter definitions by scope:** $B$, $F$, $P_{\text{benefit}}$, and $P_{\text{failure}}$ are global baseline parameters set at environment initialization. Each antibiotic in $\mathcal{A}$ has its own distinct adverse effect penalty and probability: antibiotic $a$ has parameters $A_a$ and $\alpha_a$, antibiotic $b$ has parameters $A_b$ and $\alpha_b$, and so on. These antibiotic-specific parameters distinguish the safety and toxicity profiles of different drugs. The patient-specific multipliers $\phi_{b,i}$ and $\phi_{f,i}$ scale the baseline rewards to reflect individual variation in treatment response (see Section S1.1 for details on patient heterogeneity).

#### S2.4 Range compression of individual reward & averaging

To ensure that $\lambda$ preserves intuitive meaning - where $\lambda = 0.5$ implies equal weighting between the individual reward and the community reward - we rescale individual rewards by a fixed design constant $\Theta$, which we define as the max absolute value of $B$, $F$, and all $A_{a\in\mathcal{A}}$. This bounds individual rewards to approximately within [-1, 1], therefore more closely matching the scale of community rewards:

$$\tilde{r}_{i,a,t} = \frac{r_{i,a,t}}{\Theta} \tag{S6}$$

where $\Theta = \max\left( |B|, |F|, |A_a| \text{ for all } a \in \mathcal{A}\right)$.

Finally, in order to calculate the combined individual reward for the patient cohort seen at time step $t$, we take the mean net realized individual reward:

$$\langle \tilde{R}_{\mathcal{I}}(t)\rangle = \sum_{i\in\mathcal{I}_t} \frac{\tilde{r}_{i,a,t}}{|\mathcal{I}_t|} \tag{S7}$$

The following table summarizes the reward components that apply under each combination of patient infection status and treatment decision:

Table S2: Reward components by patient infection status and treatment decision

| **Infected?** | **Treatment?** | **Indicator Behavior** | **Applicable Reward Terms** | **Notes** |
|---|---|---|---|---|
| No | No treatment | $\mathbf{1}\{\text{infected}\} = 0$ | 0 | No reward components apply |
| No | Prescribe $a$ | $\mathbf{1}\{\text{infected}\} = 0$ | $\mathbf{1}\{\text{adverse effect from } a\}\, A_a$ | Only adverse effects possible (independent of infection status) |
| Yes | No treatment | $\mathbf{1}\{\text{infected}\} = 1$ | $\mathbf{1}\{\text{recovery}\}(\phi_{b,i}B)$ or $(1 - \mathbf{1}\{\text{recovery}\})\mathbf{1}\{\text{failure}\}(\phi_{f,i}F)$ | Benefit if spontaneous recovery occurs, otherwise potential failure penalty |
| Yes | Prescribe $a$ (sensitive) | $\mathbf{1}\{\text{infected}\} = 1$, $\mathbf{1}\{\text{resistant to } a\} = 0$ | $\mathbf{1}\{\text{achieve benefit}\}(\phi_{b,i}B) + \mathbf{1}\{\text{adverse}\}A_a$ | Potential benefit (if success occurs) + possible adverse effects |
| Yes | Prescribe $a$ (resistant) | $\mathbf{1}\{\text{infected}\} = 1$, $\mathbf{1}\{\text{resistant to } a\} = 1$ | $\mathbf{1}\{\text{incur failure}\}(\phi_{f,i}F) + \mathbf{1}\{\text{adverse}\}A_a$ | Potential clinical failure penalty (if outcome occurs) + possible adverse effects |

### S3 Options and Option Libraries

#### S3.1 Options and Option Types

**Hierarchical reinforcement learning (HRL)** provides an alternative to standard RL by having the agent select among high-level strategies (called *options*) rather than raw actions at each timestep. Each option encapsulates a prescribing policy that runs for multiple timesteps before returning control to the agent (the "manager"), enabling learning of temporal strategies like cycling or risk-stratified protocols that are difficult to discover through action-by-action learning.

An **option** is a reusable decision-making module that takes environment state (patient attributes, AMR levels) as input and outputs prescribing actions for a fixed duration. The `abx_amr_simulator` package includes three standard option types:

- **Block options**: Prescribe a single antibiotic (or no treatment) for a fixed number of timesteps. *Example*: `A_10` prescribes antibiotic A for 10 consecutive timesteps.

- **Alternation options**: Prescribe a deterministic sequence of antibiotics that repeats as needed.
*Example*: `ALT_AABBA` prescribes A, A, B, B, A in sequence, then cycles back to the start.
- **Heuristic options** (also called **heuristic workers**): Apply clinical decision rules based on observable patient attributes and current AMR levels to choose which antibiotic to prescribe at each timestep within the option's duration.
*Example*: A conservative heuristic worker might prescribe only when infection probability exceeds 0.7 and expected clinical benefit is positive; an aggressive worker might prescribe whenever infection probability exceeds 0.3.

The terms **"option"** and **"worker"** are used interchangeably; "worker" emphasizes that heuristic options actively compute decisions rather than following fixed sequences. All options (including heuristic workers) implement the same interface, making them drop-in replaceable components.

An **option library** is a curated collection of options provided to the manager agent as its action space. During training, the manager learns to select options from this library based on current AMR levels and patient population characteristics. The library defines which strategies the agent can compose: a library with only block options enables learning of simple prescribing-or-withholding policies, while a library with block, alternation, and heuristic options enables discovery of cycling strategies and risk-stratified protocols that adapt to patient heterogeneity. Option libraries are specified in YAML configuration files that reference individual option definitions; full specifications are available in the GitHub repository documentation [GitHub repository link].

##### S3.2 Relationship Between Fixed Prescribing Rules and Heuristic Workers

The 'Expected Reward' fixed prescribing rules (as described in Section 3.6 of the main manuscript, and the heuristic workers used in hierarchical RL agents (as described above) compute the expected reward using the same underlying function that requires the same data: observed patient attributes and prespecified clinical scenario parameters. However, the fixed prescribing rules and the heuristic workers serve fundamentally different roles. Fixed prescribing rules apply this calculation once per patient and select an action deterministically — they never adapt or learn. Heuristic workers encode similar clinical logic but operate as temporal options within a hierarchical RL framework: the manager agent learns to select among multiple workers (e.g., "aggressive treatment," "conservative treatment," "withhold") based on AMR state evolution. This learned switching behavior enables discovery of cycling strategies and risk-stratified protocols that fixed rules cannot achieve.

#### S4 RL Agents & Fixed Experimental Rules: AMR Trajectories, Cumulative Clinical Outcomes, and Summary Statistics

See additional document, `Supplementary Figures: RL Agents & Fixed Experimental Rules - AMR Trajectories, Cumulative Clinical Outcomes, and Summary Statistics`
