## Supplementary material for "abx_amr_simulator: A simulation environment for antibiotic prescribing policy optimization under antimicrobial resistance": S2 Supplementary Figures

### Supplementary Figures: RL Agents & Fixed prescribing rule (Expected Reward - Greedy) - AMR Trajectories, Cumulative Clinical Outcomes, and Summary Statistics

Two Antibiotics No Cross-Resistance, Accurate Risk Stratification, Expected Reward Greedy 73
Two Antibiotics No Cross-Resistance, Accurate Risk Stratification, Expected Reward Lowest AMR . . . . . . . . . . . . . . . . . . . . . . . . . . . . . . . . . . . . . . . . . . 74

Two Antibiotics No Cross-Resistance, Exaggerated Risk Stratification, Expected Reward Greedy 75
Two Antibiotics No Cross-Resistance, Exaggerated Risk Stratification, Expected Reward Lowest AMR . . . 77
Two Antibiotics No Cross-Resistance, Compressed Risk Stratification, Expected Reward Greedy 78
Two Antibiotics No Cross-Resistance, Compressed Risk Stratification, Expected Reward Lowest AMR . . . 79
Two Antibiotics With Cross-Resistance, Accurate Risk Stratification, Expected Reward Greedy 81
Two Antibiotics With Cross-Resistance, Accurate Risk Stratification, Expected Reward Lowest AMR . . . 82
Two Antibiotics With Cross-Resistance, Exaggerated Risk Stratification, Expected Reward Greedy . . . 83
Two Antibiotics With Cross-Resistance, Exaggerated Risk Stratification, Expected Reward Lowest AMR . . . 85
Two Antibiotics With Cross-Resistance, Compressed Risk Stratification, Expected Reward Greedy . . . 86
Two Antibiotics With Cross-Resistance, Compressed Risk Stratification, Expected Reward Lowest AMR . . . 87
Experiment Set 4 . . . 89
Single Antibiotic, Expected Reward Greedy . . . 89
Single Antibiotic, Expected Reward Lowest AMR . . . 90
Two Antibiotics No Cross-Resistance, Expected Reward Greedy . . . 92
Two Antibiotics No Cross-Resistance, Expected Reward Lowest AMR . . . 93
Two Antibiotics With Cross-Resistance, Expected Reward Greedy . . . 94
Two Antibiotics With Cross-Resistance, Expected Reward Lowest AMR . . . 96

### RL Agents

#### Experiment Set 1

##### Single Antibiotic, Flat PPO

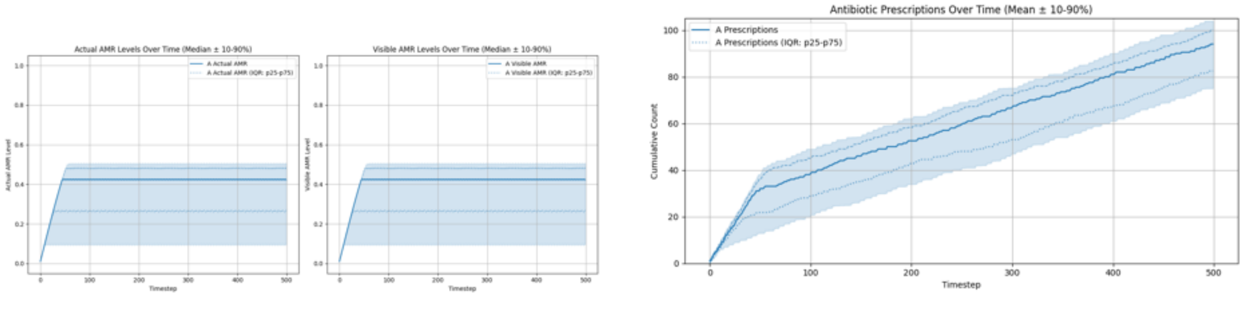

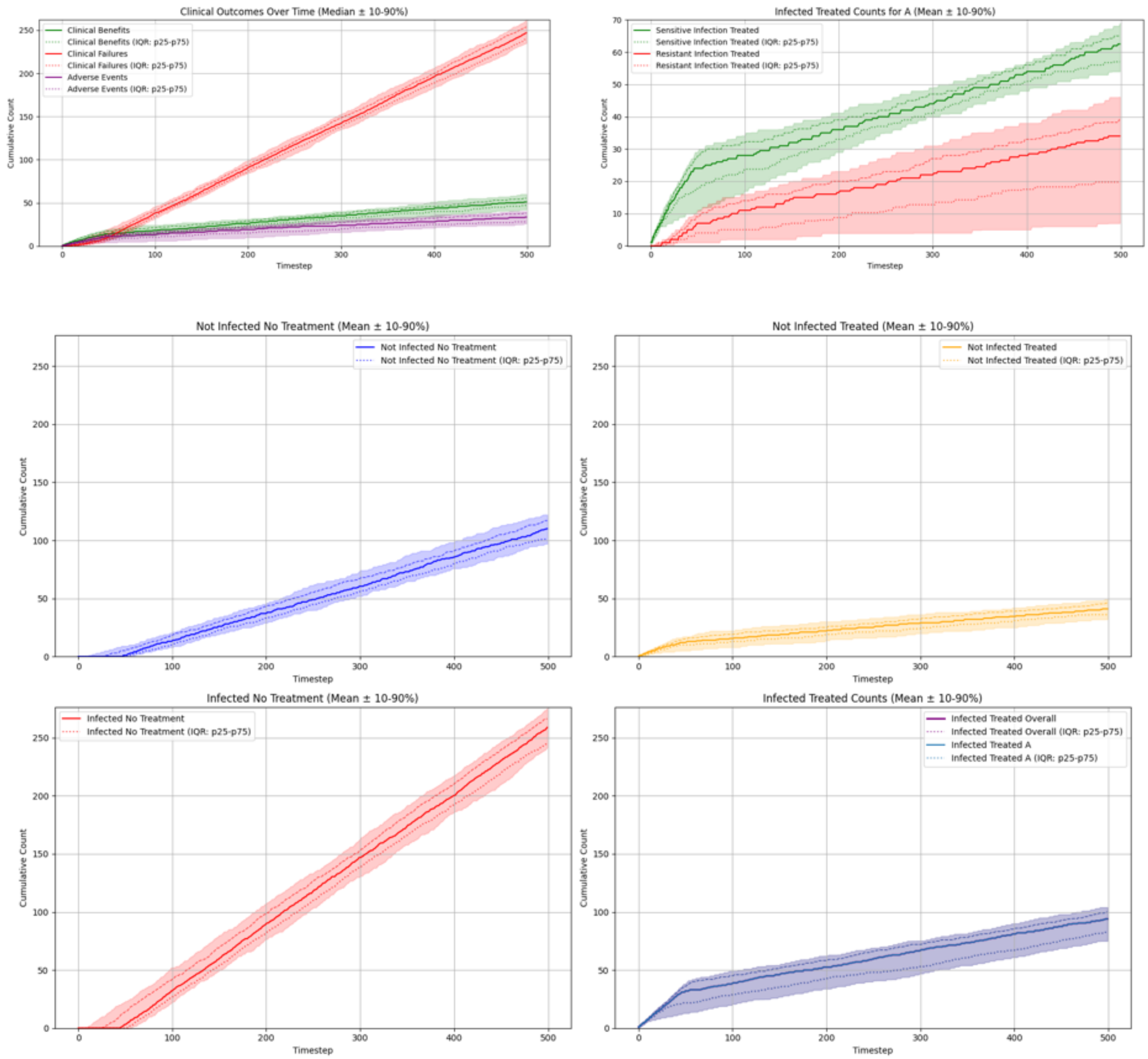

#### Summary Statistics (Single Antibiotic, Flat PPO)

| metric | p10 | p25 | p50 | p75 | p90 |
|---|---|---|---|---|---|
| overall_total_reward | -240.113 | -234.921 | -224.072 | -211.167 | -203.210 |
| overall_count_clinical_benefits | 42.000 | 47.000 | 51.000 | 55.250 | 60.000 |
| overall_count_clinical_failures | 234.900 | 240.000 | 247.000 | 254.000 | 261.100 |
| overall_count_adverse_events | 26.000 | 28.000 | 33.500 | 38.000 | 40.000 |
| overall_not_infected_no_treatment_count | 97.000 | 101.000 | 110.000 | 117.000 | 122.100 |
| overall_not_infected_treated_count | 32.000 | 36.000 | 41.000 | 46.000 | 49.000 |
| overall_infected_no_treatment_count | 241.600 | 245.000 | 259.000 | 266.500 | 275.500 |
| overall_infected_treated_count | 75.000 | 82.500 | 94.000 | 100.000 | 104.000 |
| overall_sensitive_infection_treated_count_per_abx_dict_A | 54.000 | 57.000 | 62.500 | 65.000 | 69.000 |
| overall_resistant_infection_treated_count_per_abx_dict_A | 7.000 | 19.750 | 34.000 | 39.000 | 46.100 |
| overall_sensitive_infection_treated_count | 54.000 | 57.000 | 62.500 | 65.000 | 69.000 |
| overall_resistant_infection_treated_count | 7.000 | 19.750 | 34.000 | 39.000 | 46.100 |
| overall_abx_prescriptions_count_per_abx_A | 75.000 | 82.500 | 94.000 | 100.000 | 104.000 |
| overall_abx_prescriptions_count | 75.000 | 82.500 | 94.000 | 100.000 | 104.000 |
| final_amr_actual_A | 0.095 | 0.265 | 0.424 | 0.481 | 0.504 |
| final_amr_visible_A | 0.095 | 0.265 | 0.424 | 0.481 | 0.504 |

#### Single Antibiotic, Hierarchical PPO

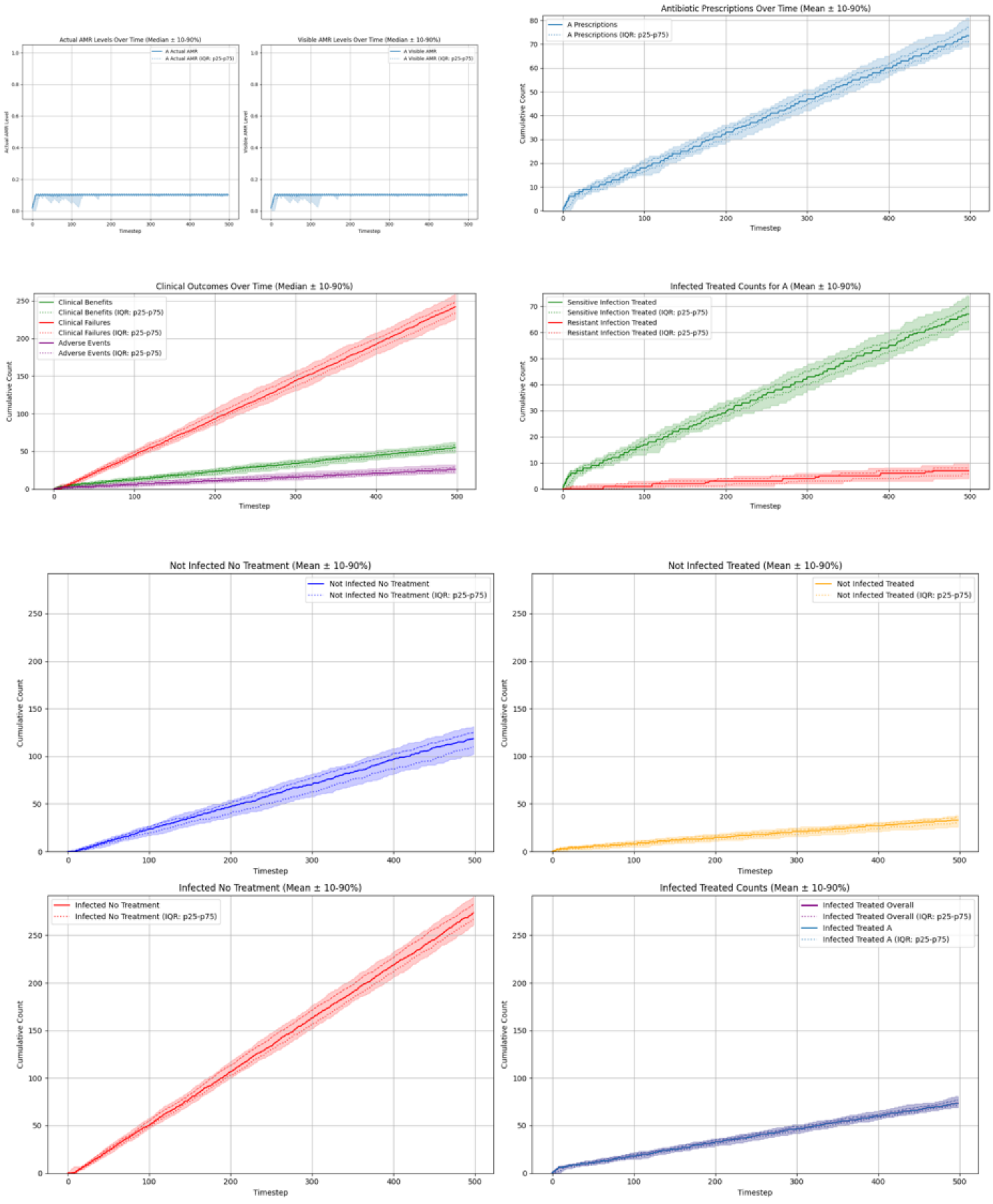

#### Summary Statistics (Single Antibiotic, Hierarchical PPO)

| metric | p10 | p25 | p50 | p75 | p90 |
|---|---|---|---|---|---|
| overall__total__reward | -224.286 | -216.966 | -209.366 | -195.966 | -192.406 |
| overall__count__clinical__benefits | 48.000 | 51.000 | 55.000 | 58.250 | 62.100 |
| overall__count__clinical__failures | 225.900 | 233.750 | 242.000 | 248.250 | 259.000 |
| overall__count__adverse__events | 21.000 | 23.000 | 26.000 | 29.000 | 32.200 |
| overall__not__infected__no__treatment__count | 102.000 | 110.000 | 118.500 | 125.250 | 131.200 |
| overall__not__infected__treated__count | 26.000 | 30.000 | 33.000 | 36.000 | 38.000 |
| overall__infected__no__treatment__count | 260.900 | 266.750 | 273.500 | 282.250 | 291.000 |
| overall__infected__treated__count | 69.000 | 71.000 | 73.500 | 77.000 | 81.000 |
| overall__sensitive__infection__treated__count__per__abx__dict__A | 61.000 | 64.000 | 67.000 | 70.000 | 74.000 |
| overall__resistant__infection__treated__count__per__abx__dict__A | 4.000 | 5.750 | 7.000 | 8.000 | 10.000 |
| overall__sensitive__infection__treated__count | 61.000 | 64.000 | 67.000 | 70.000 | 74.000 |
| overall__resistant__infection__treated__count | 4.000 | 5.750 | 7.000 | 8.000 | 10.000 |
| overall__abx__prescriptions__count__per__abx__A | 69.000 | 71.000 | 73.500 | 77.000 | 81.000 |
| overall__abx__prescriptions__count | 69.000 | 71.000 | 73.500 | 77.000 | 81.000 |
| final__amr__actual__A | 0.102 | 0.102 | 0.102 | 0.102 | 0.102 |
| final__amr__visible__A | 0.102 | 0.102 | 0.102 | 0.102 | 0.102 |

##### Two Antibiotics No Cross-Resistance, Flat PPO

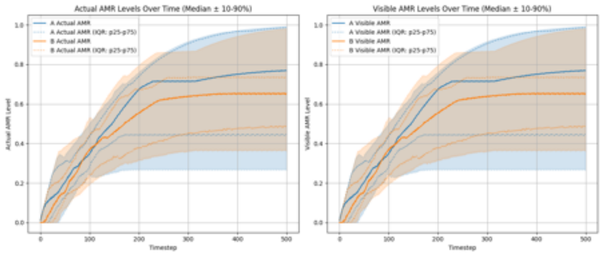

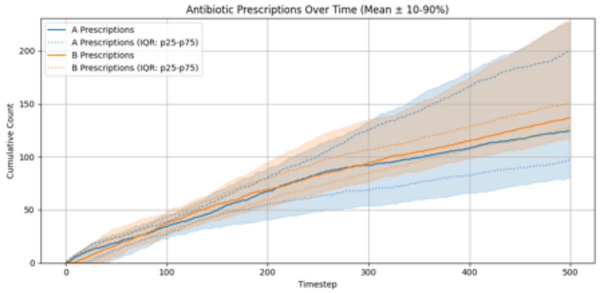

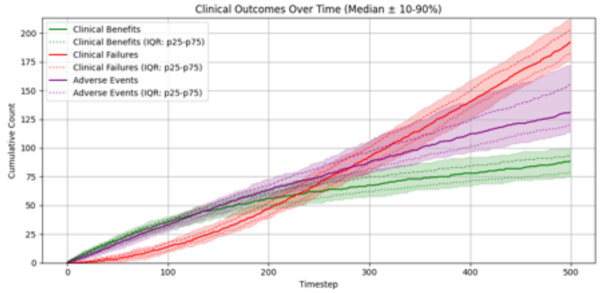

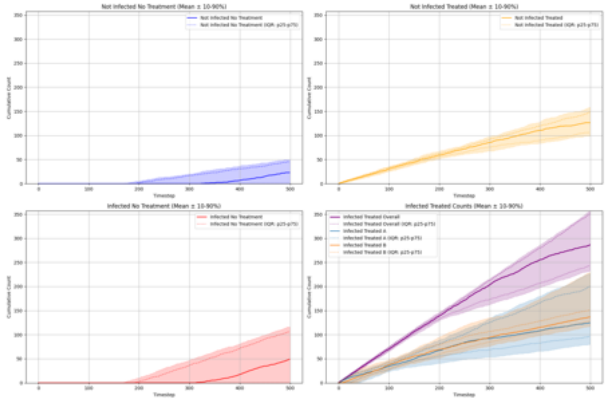

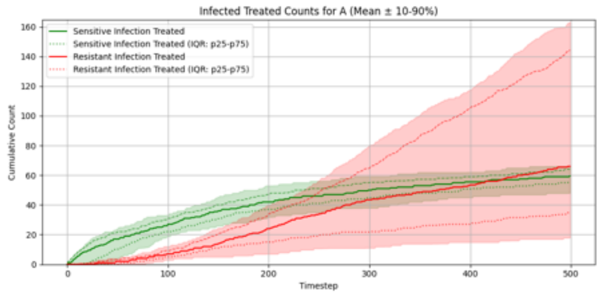

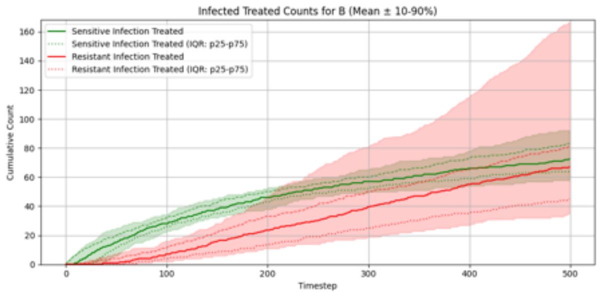

##### Summary Statistics (Two Antibiotics No Cross-Resistance, Flat PPO)

| metric | p10 | p25 | p50 | p75 | p90 |
|---|---|---|---|---|---|
| overall_total_reward | -226.544 | -204.980 | -186.425 | -171.697 | -162.257 |
| overall_count_clinical_benefits | 74.900 | 78.750 | 88.500 | 93.000 | 99.300 |
| overall_count_clinical_failures | 176.900 | 182.000 | 192.000 | 202.250 | 212.200 |
| overall_count_adverse_events | 113.800 | 119.750 | 131.000 | 155.000 | 172.000 |
| overall_not_infected_no_treatment_count | 0.000 | 0.000 | 23.000 | 45.250 | 50.200 |
| overall_not_infected_treated_count | 98.900 | 106.000 | 126.500 | 148.000 | 158.400 |
| overall_infected_no_treatment_count | 0.000 | 0.000 | 49.000 | 106.250 | 116.100 |
| overall_infected_treated_count | 232.900 | 243.750 | 286.500 | 351.000 | 356.100 |
| overall_sensitive_infection_treated_count_per_abx_dict_A | 48.000 | 56.000 | 59.500 | 64.000 | 66.100 |
| overall_sensitive_infection_treated_count_per_abx_dict_B | 57.800 | 63.750 | 72.500 | 83.000 | 92.100 |
| overall_resistant_infection_treated_count_per_abx_dict_A | 17.900 | 34.750 | 66.000 | 144.500 | 163.600 |
| overall_resistant_infection_treated_count_per_abx_dict_B | 34.700 | 45.000 | 67.000 | 81.000 | 167.100 |
| overall_sensitive_infection_treated_count | 118.900 | 123.750 | 132.000 | 143.000 | 151.400 |
| overall_resistant_infection_treated_count | 103.900 | 117.500 | 156.500 | 203.500 | 222.300 |
| overall_abx_prescriptions_count_per_abx_A | 79.900 | 96.750 | 124.500 | 200.500 | 226.500 |
| overall_abx_prescriptions_count_per_abx_B | 117.800 | 127.750 | 136.500 | 151.250 | 229.500 |
| overall_abx_prescriptions_count | 232.900 | 243.750 | 286.500 | 351.000 | 356.100 |
| final_amr_actual_A | 0.269 | 0.444 | 0.769 | 0.986 | 0.991 |
| final_amr_visible_A | 0.269 | 0.444 | 0.769 | 0.986 | 0.991 |
| final_amr_actual_B | 0.366 | 0.485 | 0.652 | 0.734 | 0.981 |
| final_amr_visible_B | 0.366 | 0.485 | 0.652 | 0.734 | 0.981 |

##### Two Antibiotics No Cross-Resistance, Hierarchical PPO

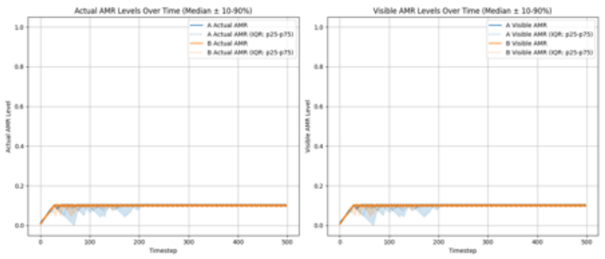

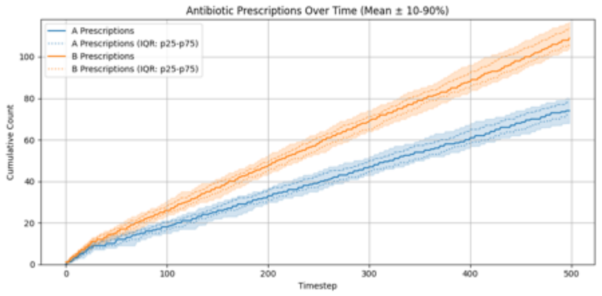

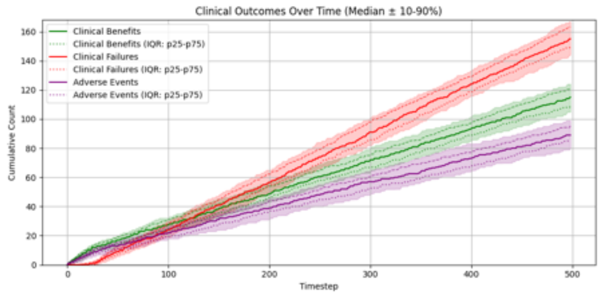

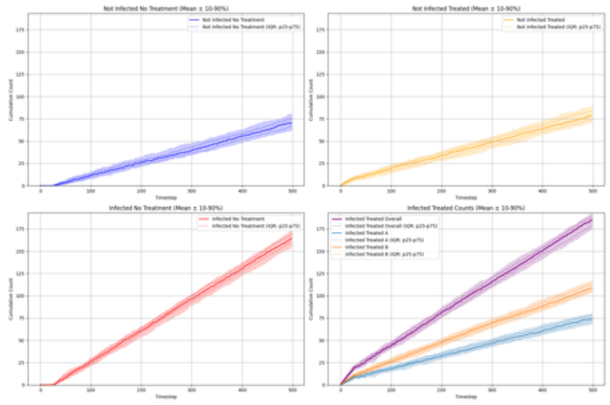

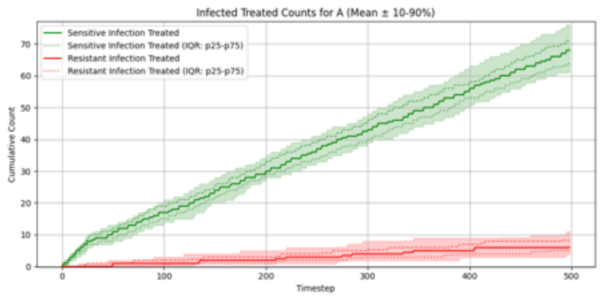

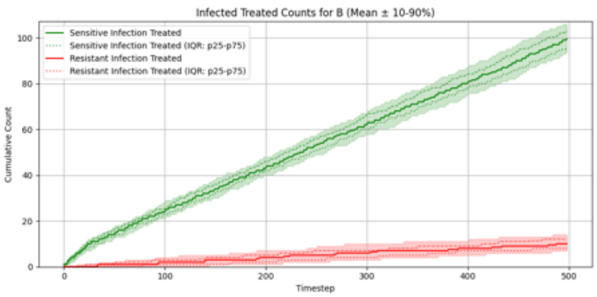

#### Summary Statistics (Two Antibiotics No Cross-Resistance, Hierarchical PPO)

| metric | p10 | p25 | p50 | p75 | p90 |
|---|---|---|---|---|---|
| overall_total_reward | -109.507 | -103.701 | -95.526 | -84.001 | -76.027 |
| overall_count_clinical_benefits | 104.900 | 108.750 | 115.000 | 121.000 | 124.100 |
| overall_count_clinical_failures | 144.000 | 149.000 | 155.000 | 163.250 | 167.000 |
| overall_count_adverse_events | 80.000 | 85.000 | 89.000 | 95.250 | 99.000 |
| overall_not_infected_no_treatment_count | 62.000 | 66.750 | 70.500 | 75.000 | 80.100 |
| overall_not_infected_treated_count | 71.900 | 74.750 | 78.500 | 83.500 | 89.000 |
| overall_infected_no_treatment_count | 154.900 | 160.000 | 164.500 | 169.000 | 173.100 |
| overall_infected_treated_count | 175.000 | 179.000 | 185.500 | 189.000 | 192.100 |
| overall_sensitive_infection_treated_count_per_abx_dict_A | 61.000 | 63.750 | 68.000 | 71.000 | 76.000 |
| overall_sensitive_infection_treated_count_per_abx_dict_B | 94.000 | 96.000 | 99.500 | 103.000 | 106.100 |
| overall_resistant_infection_treated_count_per_abx_dict_A | 3.900 | 5.000 | 6.000 | 8.250 | 11.000 |
| overall_resistant_infection_treated_count_per_abx_dict_B | 7.000 | 8.000 | 10.000 | 12.000 | 14.000 |
| overall_sensitive_infection_treated_count | 158.000 | 162.750 | 167.000 | 173.250 | 176.100 |
| overall_resistant_infection_treated_count | 12.000 | 14.000 | 16.000 | 20.000 | 22.100 |
| overall_abx_prescriptions_count_per_abx_A | 68.000 | 72.000 | 74.000 | 78.000 | 80.100 |
| overall_abx_prescriptions_count_per_abx_B | 104.000 | 106.000 | 109.000 | 114.000 | 117.000 |
| overall_abx_prescriptions_count | 175.000 | 179.000 | 185.500 | 189.000 | 192.100 |
| final_amr_actual_A | 0.102 | 0.102 | 0.102 | 0.102 | 0.102 |
| final_amr_visible_A | 0.102 | 0.102 | 0.102 | 0.102 | 0.102 |
| final_amr_actual_B | 0.100 | 0.100 | 0.100 | 0.100 | 0.100 |
| final_amr_visible_B | 0.100 | 0.100 | 0.100 | 0.100 | 0.100 |

#### Two Antibiotics With Cross-Resistance, Flat PPO

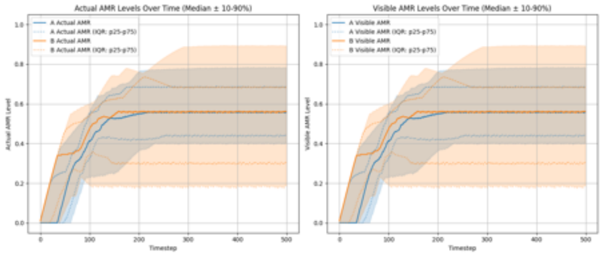

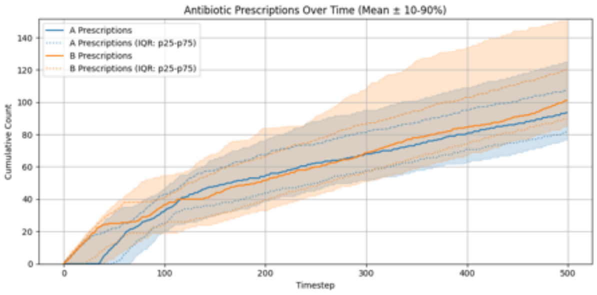

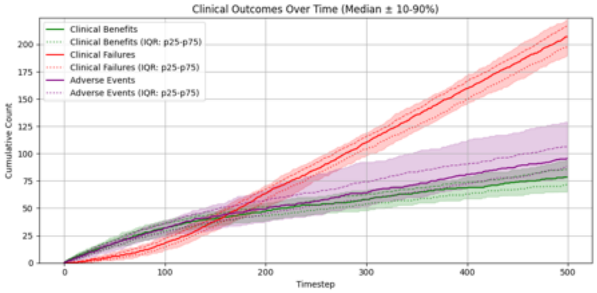

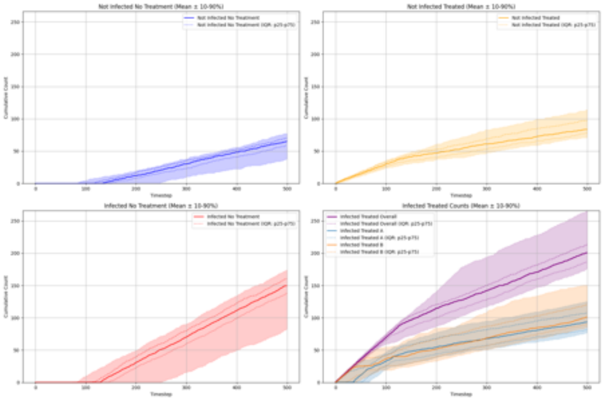

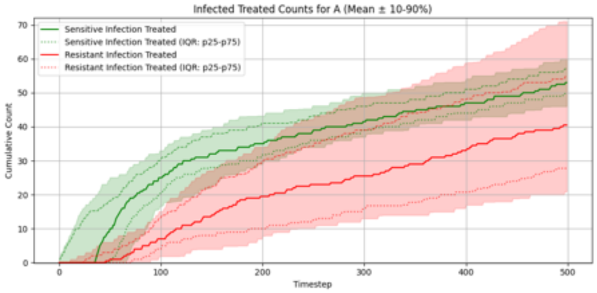

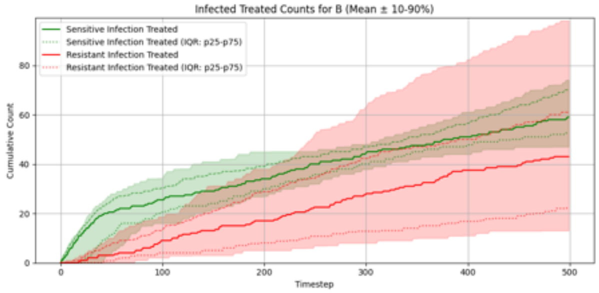

#### Summary Statistics (Two Antibiotics With Cross-Resistance, Flat PPO)

| metric | p10 | p25 | p50 | p75 | p90 |
|---|---|---|---|---|---|
| overall__total__reward | -230.728 | -204.649 | -187.646 | -167.373 | -153.619 |
| overall__count__clinical__benefits | 65.000 | 71.500 | 78.500 | 85.500 | 94.000 |
| overall__count__clinical__failures | 189.900 | 197.750 | 207.000 | 217.000 | 223.200 |
| overall__count__adverse__events | 78.000 | 88.000 | 95.500 | 106.250 | 129.000 |
| overall__not__infected__no__treatment__count | 37.800 | 59.000 | 65.000 | 70.000 | 77.100 |
| overall__not__infected__treated__count | 70.800 | 78.000 | 83.500 | 97.000 | 114.100 |
| overall__infected__no__treatment__count | 82.800 | 137.750 | 149.500 | 160.250 | 174.100 |
| overall__infected__treated__count | 175.800 | 187.000 | 200.500 | 213.500 | 265.000 |
| overall__sensitive__infection__treated__count__per__abx__dict__A | 46.000 | 49.750 | 53.000 | 57.000 | 60.000 |
| overall__sensitive__infection__treated__count__per__abx__dict__B | 47.000 | 52.750 | 59.000 | 70.000 | 74.100 |
| overall__resistant__infection__treated__count__per__abx__dict__A | 20.900 | 27.750 | 40.500 | 55.000 | 71.000 |
| overall__resistant__infection__treated__count__per__abx__dict__B | 13.000 | 22.750 | 43.000 | 61.000 | 98.200 |
| overall__sensitive__infection__treated__count | 98.900 | 103.750 | 112.000 | 123.000 | 129.100 |
| overall__resistant__infection__treated__count | 49.800 | 73.750 | 87.000 | 99.250 | 163.900 |
| overall__abx__prescriptions__count__per__abx__A | 76.700 | 81.500 | 93.500 | 107.250 | 125.000 |
| overall__abx__prescriptions__count__per__abx__B | 84.900 | 89.500 | 101.000 | 120.250 | 150.500 |
| overall__abx__prescriptions__count | 175.800 | 187.000 | 200.500 | 213.500 | 265.000 |
| final__amr__actual__A | 0.402 | 0.439 | 0.558 | 0.685 | 0.783 |
| final__amr__visible__A | 0.402 | 0.439 | 0.558 | 0.685 | 0.783 |
| final__amr__actual__B | 0.184 | 0.300 | 0.561 | 0.683 | 0.893 |
| final__amr__visible__B | 0.184 | 0.300 | 0.561 | 0.683 | 0.893 |

##### Two Antibiotics With Cross-Resistance, Hierarchical PPO

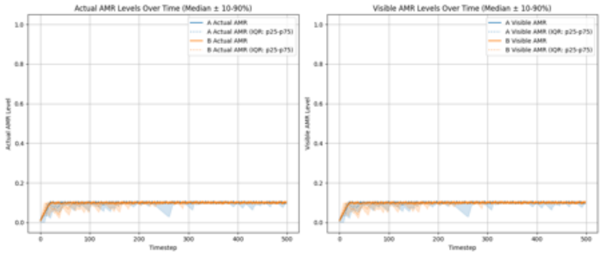

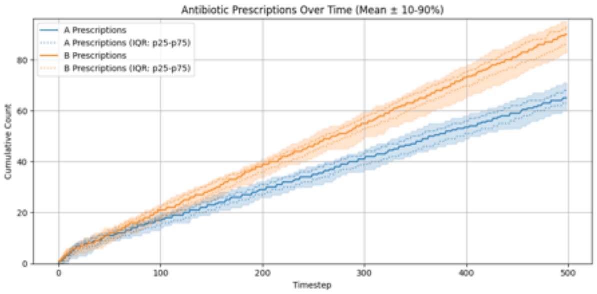

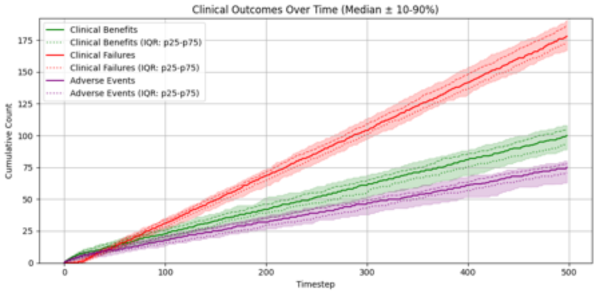

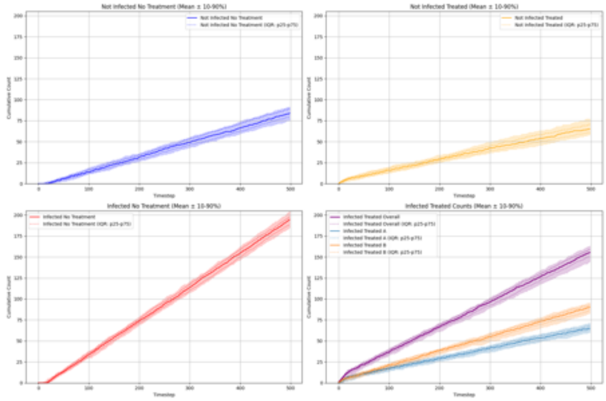

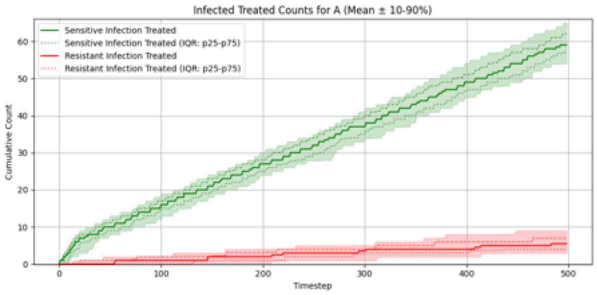

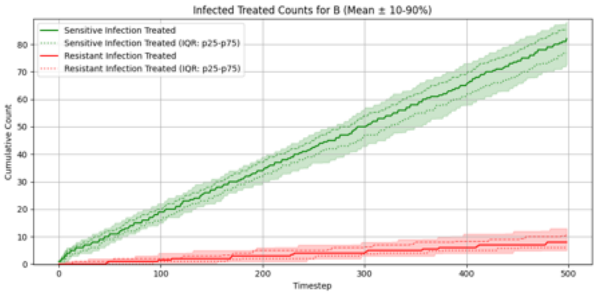

##### Summary Statistics (Two Antibiotics With Cross-Resistance, Hierarchical PPO)

| metric | p10 | p25 | p50 | p75 | p90 |
|---|---|---|---|---|---|
| overall__total__reward | -143.544 | -135.424 | -126.348 | -112.474 | -105.649 |
| overall__count__clinical__benefits | 89.000 | 92.750 | 100.000 | 104.250 | 108.100 |
| overall__count__clinical__failures | 166.900 | 172.000 | 178.000 | 186.000 | 191.000 |
| overall__count__adverse__events | 63.000 | 70.750 | 75.000 | 78.000 | 80.000 |
| overall__not__infected__no__treatment__count | 74.900 | 79.000 | 84.000 | 89.000 | 91.200 |
| overall__not__infected__treated__count | 58.000 | 61.000 | 65.000 | 71.250 | 77.100 |
| overall__infected__no__treatment__count | 184.800 | 189.000 | 194.500 | 198.000 | 204.000 |
| overall__infected__treated__count | 144.900 | 150.750 | 155.500 | 159.000 | 163.000 |
| overall__sensitive__infection__treated__count__per__abx__dict__A | 54.000 | 57.000 | 59.000 | 62.000 | 65.000 |
| overall__sensitive__infection__treated__count__per__abx__dict__B | 72.800 | 76.750 | 82.000 | 85.250 | 88.200 |
| overall__resistant__infection__treated__count__per__abx__dict__A | 3.000 | 4.000 | 5.500 | 7.000 | 9.000 |
| overall__resistant__infection__treated__count__per__abx__dict__B | 5.000 | 6.000 | 8.000 | 11.000 | 13.000 |
| overall__sensitive__infection__treated__count | 130.000 | 136.000 | 140.500 | 147.000 | 150.100 |
| overall__resistant__infection__treated__count | 9.000 | 11.000 | 14.000 | 16.250 | 20.100 |
| overall__abx__prescriptions__count__per__abx__A | 60.000 | 63.000 | 65.000 | 68.000 | 71.000 |
| overall__abx__prescriptions__count__per__abx__B | 82.700 | 86.750 | 90.000 | 93.000 | 95.100 |
| overall__abx__prescriptions__count | 144.900 | 150.750 | 155.500 | 159.000 | 163.000 |
| final__amr__actual__A | 0.086 | 0.100 | 0.100 | 0.101 | 0.102 |
| final__amr__visible__A | 0.086 | 0.100 | 0.100 | 0.101 | 0.102 |
| final__amr__actual__B | 0.100 | 0.101 | 0.101 | 0.101 | 0.101 |
| final__amr__visible__B | 0.100 | 0.101 | 0.101 | 0.101 | 0.101 |

### Experiment Set 2

#### Single Antibiotic, Flat PPO

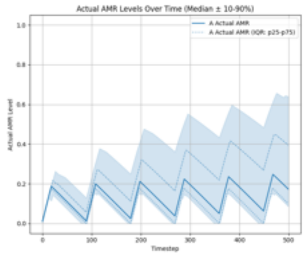

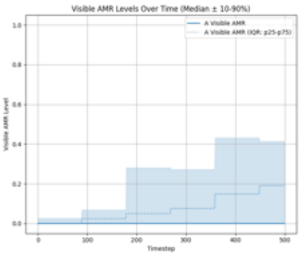

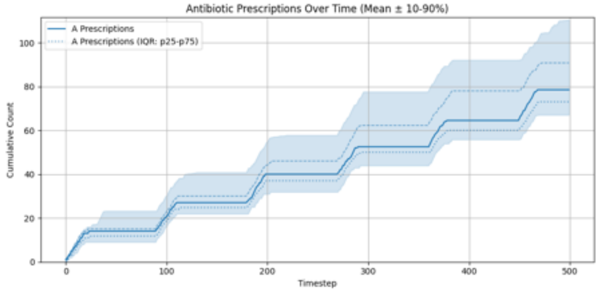

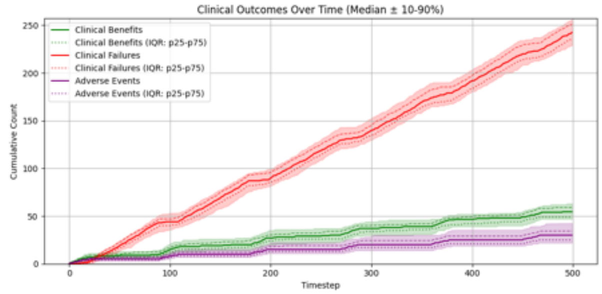

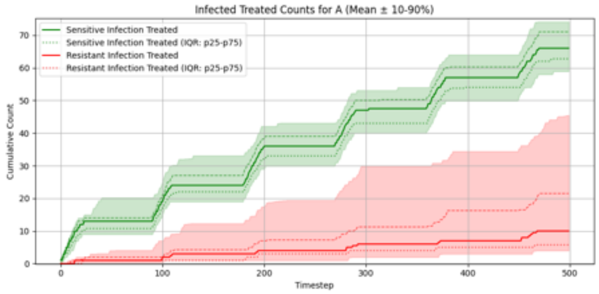

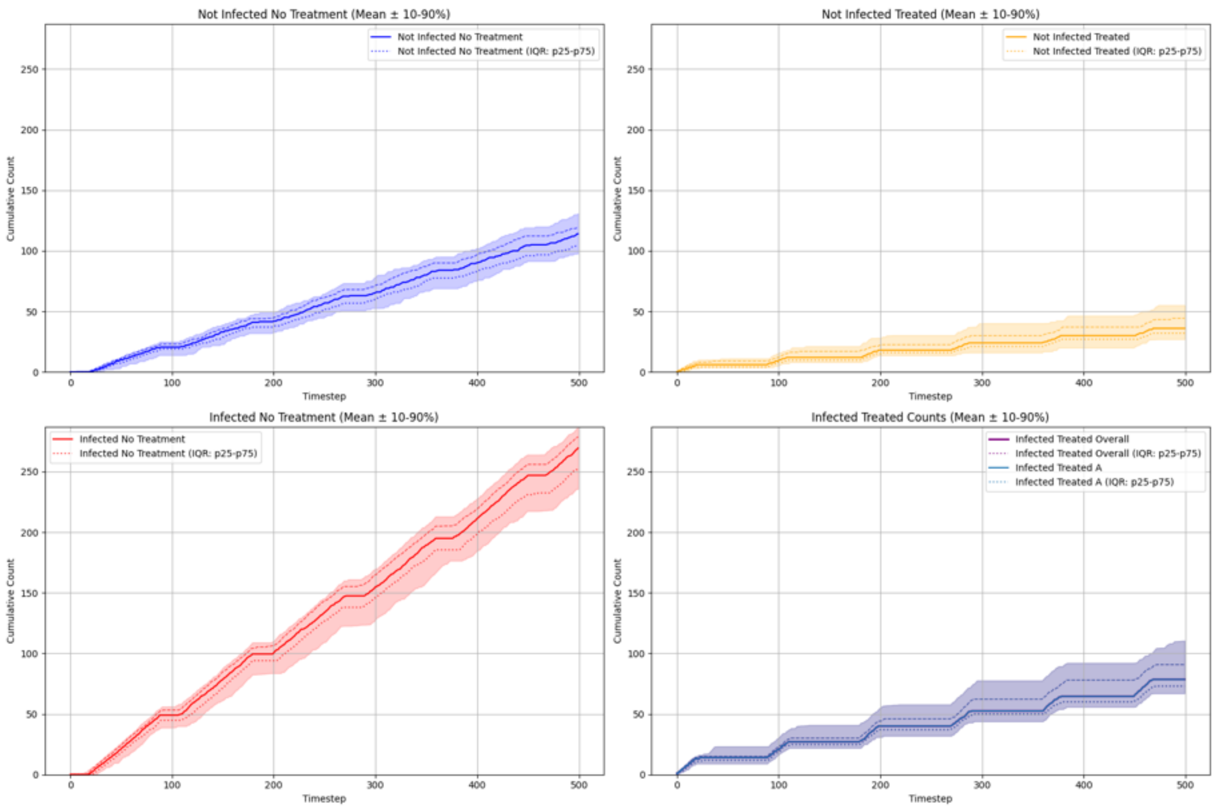

#### Summary Statistics (Single Antibiotic, Flat PPO)

| metric | p10 | p25 | p50 | p75 | p90 |
|---|---|---|---|---|---|
| overall_total_reward | -233.510 | -223.624 | -213.966 | -201.157 | -192.484 |
| overall_count_clinical_benefits | 47.000 | 49.000 | 54.500 | 59.000 | 63.000 |
| overall_count_clinical_failures | 228.800 | 235.750 | 242.500 | 251.000 | 256.100 |
| overall_count_adverse_events | 21.900 | 25.000 | 30.000 | 34.250 | 42.400 |
| overall_not_infected_no_treatment_count | 97.700 | 104.000 | 114.000 | 120.000 | 131.000 |
| overall_not_infected_treated_count | 27.000 | 32.000 | 36.000 | 44.250 | 55.100 |
| overall_infected_no_treatment_count | 235.700 | 253.000 | 269.500 | 279.000 | 286.100 |
| overall_infected_treated_count | 67.000 | 73.000 | 78.500 | 90.750 | 110.500 |
| overall_sensitive_infection_treated_count_per_abx_dict_A | 58.900 | 62.750 | 66.000 | 71.000 | 74.100 |
| overall_resistant_infection_treated_count_per_abx_dict_A | 4.000 | 5.750 | 10.000 | 21.500 | 45.500 |
| overall_sensitive_infection_treated_count | 58.900 | 62.750 | 66.000 | 71.000 | 74.100 |
| overall_resistant_infection_treated_count | 4.000 | 5.750 | 10.000 | 21.500 | 45.500 |
| overall_abx_prescriptions_count_per_abx_A | 67.000 | 73.000 | 78.500 | 90.750 | 110.500 |
| overall_abx_prescriptions_count | 67.000 | 73.000 | 78.500 | 90.750 | 110.500 |
| final_amr_actual_A | 0.097 | 0.111 | 0.184 | 0.404 | 0.643 |
| final_amr_visible_A | 0.000 | 0.000 | 0.000 | 0.191 | 0.413 |

#### Single Antibiotic, Recurrent PPO

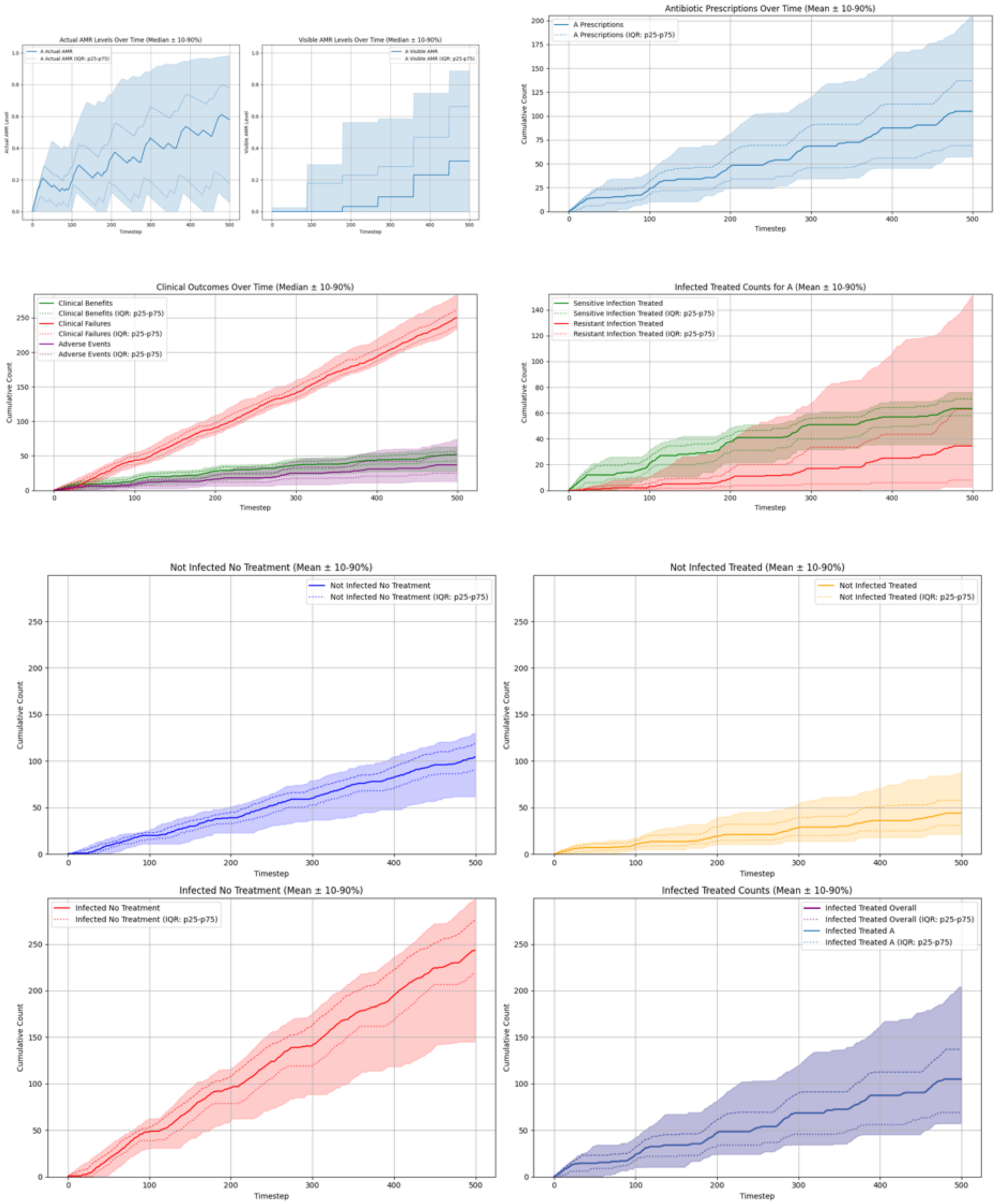

#### Summary Statistics (Single Antibiotic, Recurrent PPO)

| metric | p10 | p25 | p50 | p75 | p90 |
|---|---|---|---|---|---|
| overall__total__reward | -287.299 | -262.082 | -225.992 | -212.025 | -199.866 |
| overall__count__clinical__benefits | 27.800 | 42.500 | 52.000 | 56.000 | 63.100 |
| overall__count__clinical__failures | 233.000 | 237.000 | 250.000 | 262.250 | 283.100 |
| overall__count__adverse__events | 13.100 | 25.000 | 37.000 | 51.000 | 74.000 |
| overall__not__infected__no__treatment__count | 61.900 | 90.500 | 104.500 | 119.000 | 130.400 |
| overall__not__infected__treated__count | 21.300 | 30.750 | 44.000 | 57.750 | 88.100 |
| overall__infected__no__treatment__count | 145.900 | 219.250 | 243.500 | 276.000 | 298.600 |
| overall__infected__treated__count | 57.300 | 69.000 | 105.000 | 137.000 | 204.200 |
| overall__sensitive__infection__treated__count__per__abx__dict__A | 35.700 | 58.000 | 63.500 | 71.000 | 76.200 |
| overall__resistant__infection__treated__count__per__abx__dict__A | 2.700 | 8.000 | 34.500 | 62.750 | 151.200 |
| overall__sensitive__infection__treated__count | 35.700 | 58.000 | 63.500 | 71.000 | 76.200 |
| overall__resistant__infection__treated__count | 2.700 | 8.000 | 34.500 | 62.750 | 151.200 |
| overall__abx__prescriptions__count__per__abx__A | 57.300 | 69.000 | 105.000 | 137.000 | 204.200 |
| overall__abx__prescriptions__count | 57.300 | 69.000 | 105.000 | 137.000 | 204.200 |
| final__amr__actual__A | 0.066 | 0.187 | 0.587 | 0.786 | 0.981 |
| final__amr__visible__A | 0.000 | 0.000 | 0.319 | 0.662 | 0.888 |

##### Single Antibiotic, Hierarchical PPO

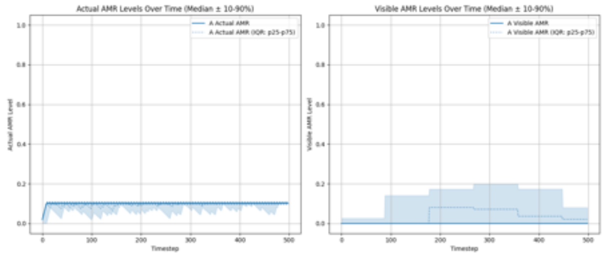

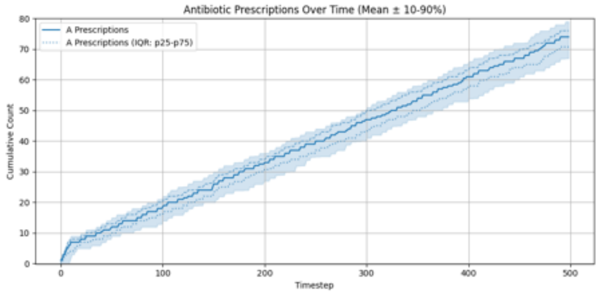

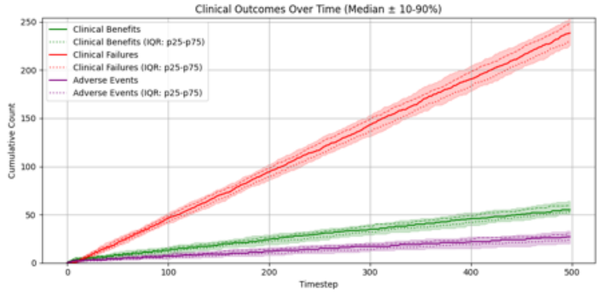

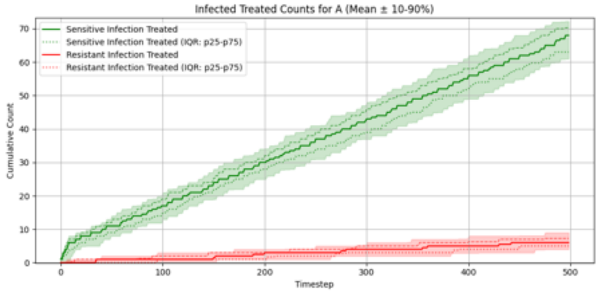

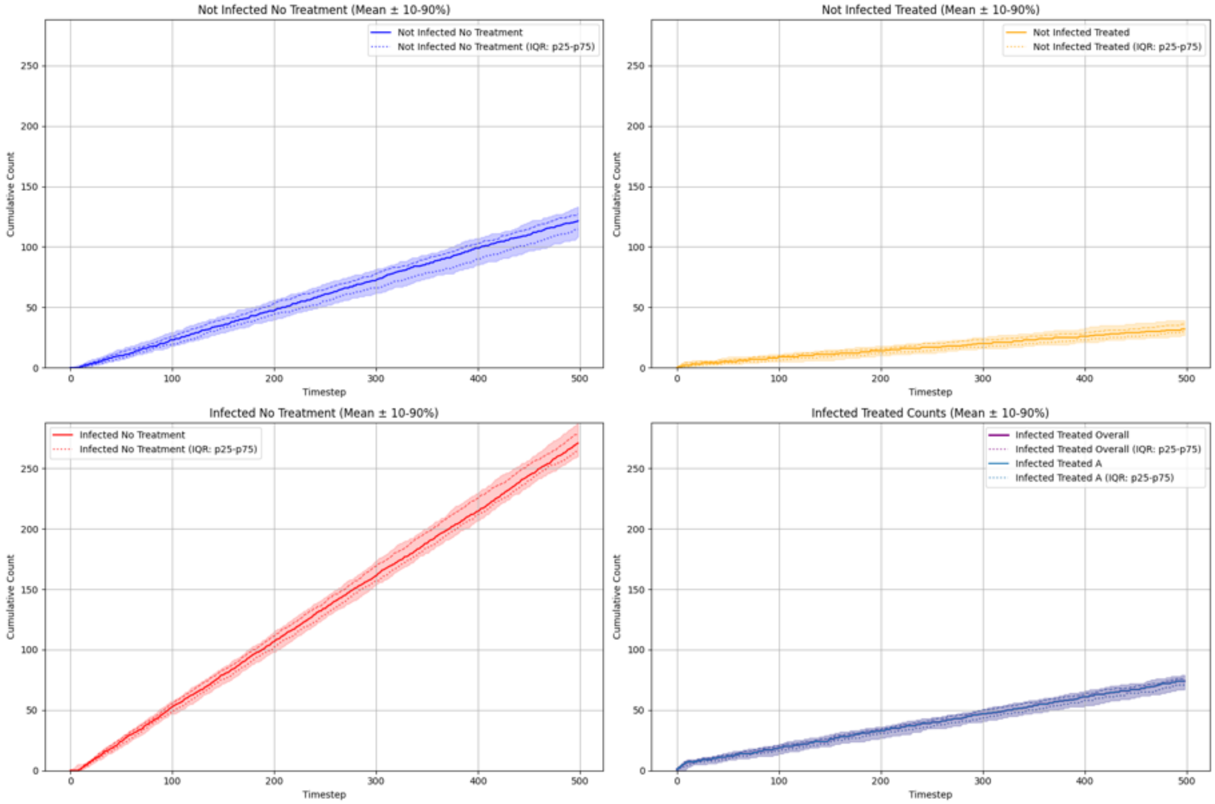

#### Summary Statistics (Single Antibiotic, Hierarchical PPO)

| metric | p10 | p25 | p50 | p75 | p90 |
|---|---|---|---|---|---|
| overall_total_reward | -219.858 | -213.905 | -205.253 | -195.357 | -186.419 |
| overall_count_clinical_benefits | 50.900 | 53.000 | 55.000 | 59.250 | 64.000 |
| overall_count_clinical_failures | 225.900 | 231.000 | 238.500 | 248.250 | 253.100 |
| overall_count_adverse_events | 20.000 | 23.750 | 27.000 | 30.000 | 33.000 |
| overall_not_infected_no_treatment_count | 108.600 | 114.500 | 121.500 | 127.000 | 133.100 |
| overall_not_infected_treated_count | 26.900 | 29.750 | 32.000 | 36.000 | 39.000 |
| overall_infected_no_treatment_count | 260.000 | 265.000 | 271.000 | 279.000 | 287.000 |
| overall_infected_treated_count | 67.000 | 70.750 | 74.000 | 76.000 | 79.000 |
| overall_sensitive_infection_treated_count_per_abx_dict_A | 61.000 | 63.000 | 68.000 | 70.250 | 72.200 |
| overall_resistant_infection_treated_count_per_abx_dict_A | 4.000 | 5.000 | 6.000 | 7.250 | 9.000 |
| overall_sensitive_infection_treated_count | 61.000 | 63.000 | 68.000 | 70.250 | 72.200 |
| overall_resistant_infection_treated_count | 4.000 | 5.000 | 6.000 | 7.250 | 9.000 |
| overall_abx_prescriptions_count_per_abx_A | 67.000 | 70.750 | 74.000 | 76.000 | 79.000 |
| overall_abx_prescriptions_count | 67.000 | 70.750 | 74.000 | 76.000 | 79.000 |
| final_amr_actual_A | 0.101 | 0.102 | 0.102 | 0.102 | 0.102 |
| final_amr_visible_A | 0.000 | 0.000 | 0.000 | 0.021 | 0.080 |

#### Single Antibiotic, Hierarchical Recurrent PPO

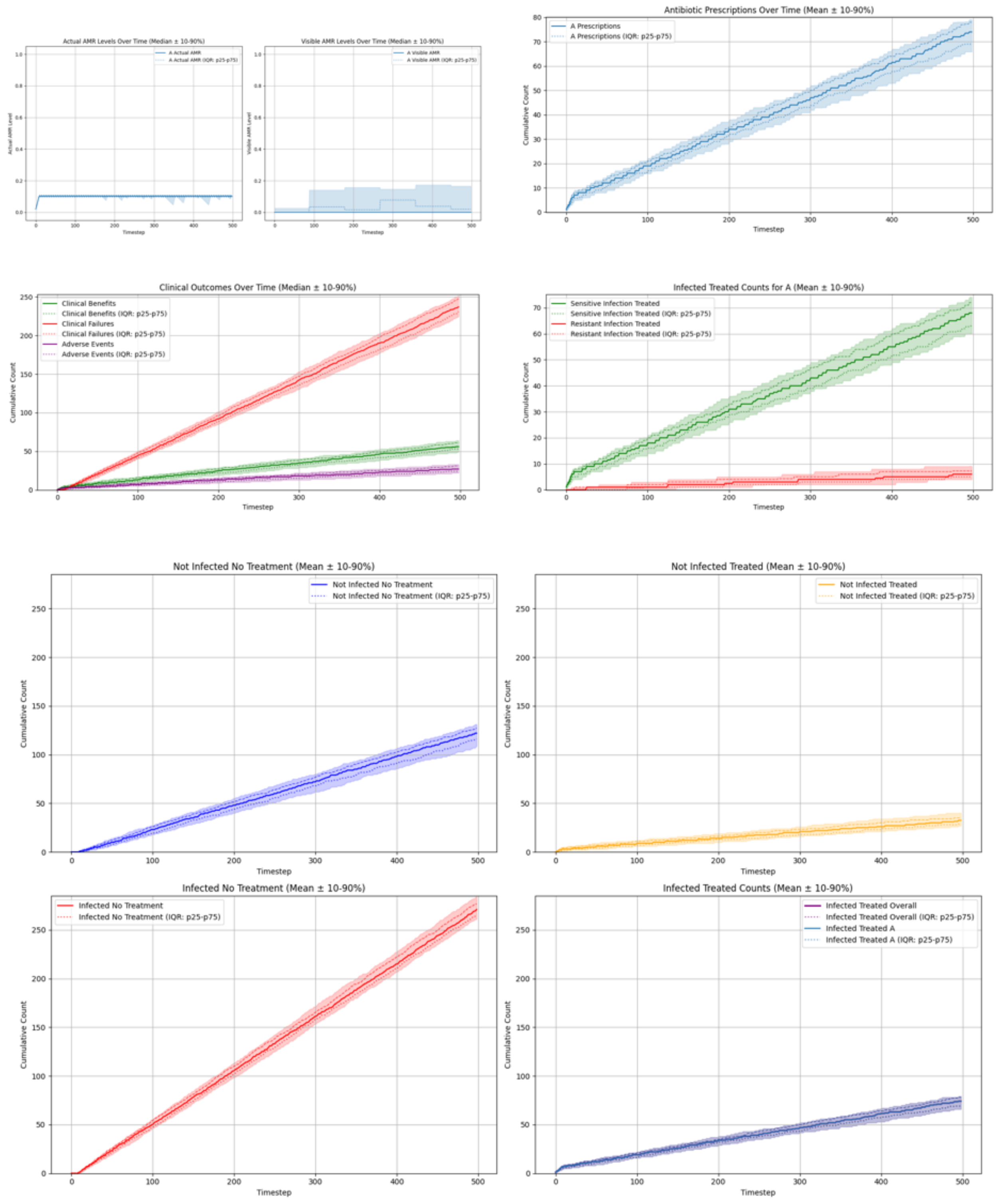

#### Summary Statistics (Single Antibiotic, Hierarchical Recurrent PPO)

| metric | p10 | p25 | p50 | p75 | p90 |
|---|---|---|---|---|---|
| overall_total_reward | -221.696 | -212.506 | -202.861 | -194.007 | -184.516 |
| overall_count_clinical_benefits | 49.000 | 53.000 | 56.000 | 61.000 | 64.000 |
| overall_count_clinical_failures | 224.900 | 230.750 | 237.000 | 247.250 | 252.100 |
| overall_count_adverse_events | 22.000 | 24.000 | 27.000 | 31.000 | 33.000 |
| overall_not_infected_no_treatment_count | 108.900 | 116.000 | 122.000 | 127.000 | 131.100 |
| overall_not_infected_treated_count | 26.900 | 28.750 | 32.500 | 35.000 | 40.100 |
| overall_infected_no_treatment_count | 261.000 | 265.750 | 271.000 | 276.750 | 284.100 |
| overall_infected_treated_count | 65.900 | 69.000 | 74.000 | 78.000 | 79.100 |
| overall_sensitive_infection_treated_count_per_abx_dict_A | 60.000 | 63.000 | 68.000 | 72.000 | 74.100 |
| overall_resistant_infection_treated_count_per_abx_dict_A | 4.000 | 5.000 | 6.000 | 7.250 | 9.100 |
| overall_sensitive_infection_treated_count | 60.000 | 63.000 | 68.000 | 72.000 | 74.100 |
| overall_resistant_infection_treated_count | 4.000 | 5.000 | 6.000 | 7.250 | 9.100 |
| overall_abx_prescriptions_count_per_abx_A | 65.900 | 69.000 | 74.000 | 78.000 | 79.100 |
| overall_abx_prescriptions_count | 65.900 | 69.000 | 74.000 | 78.000 | 79.100 |
| final_amr_actual_A | 0.085 | 0.102 | 0.102 | 0.102 | 0.102 |
| final_amr_visible_A | 0.000 | 0.000 | 0.000 | 0.019 | 0.166 |

##### Two Antibiotics No Cross-Resistance, Flat PPO

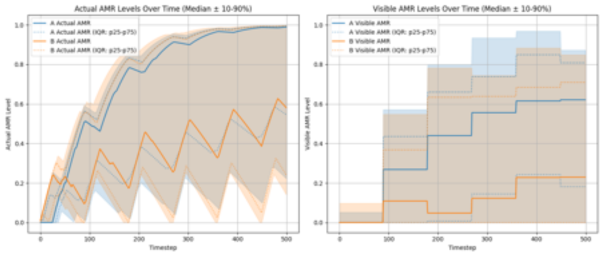

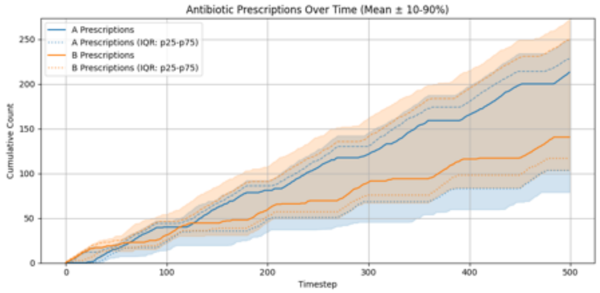

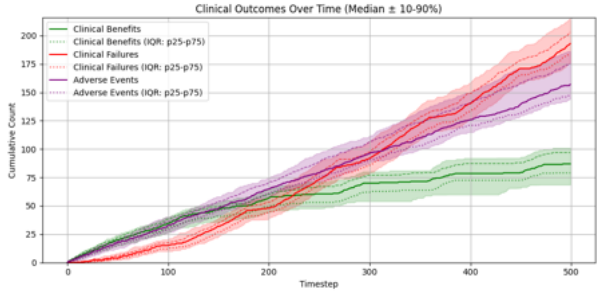

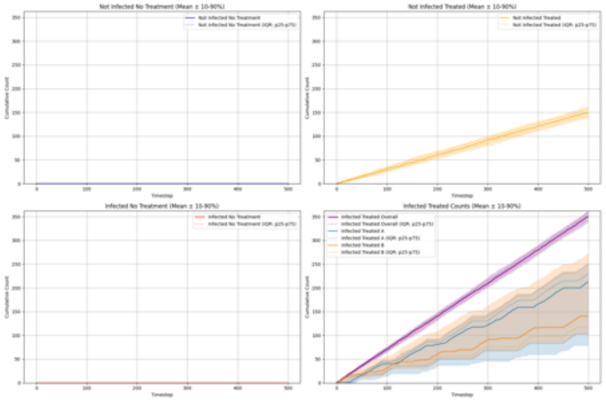

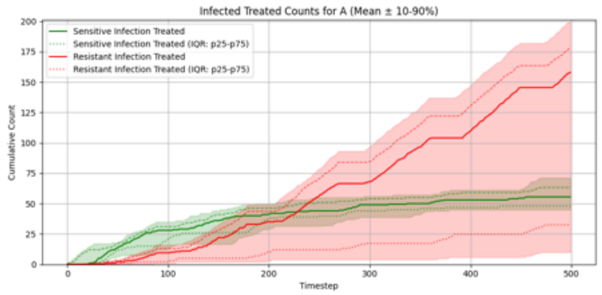

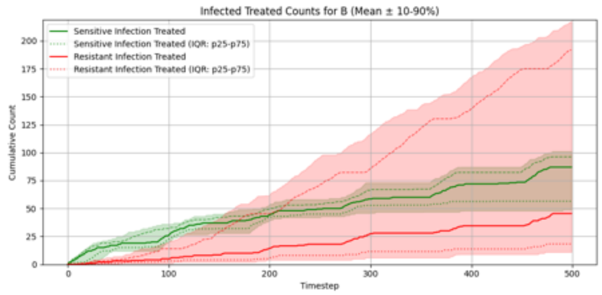

#### Summary Statistics (Two Antibiotics No Cross-Resistance, Flat PPO)

| metric | p10 | p25 | p50 | p75 | p90 |
|---|---|---|---|---|---|
| overall__total__reward | -245.382 | -219.736 | -203.199 | -189.750 | -175.880 |
| overall__count__clinical__benefits | 68.600 | 79.000 | 87.000 | 97.000 | 101.000 |
| overall__count__clinical__failures | 175.900 | 184.000 | 193.000 | 203.000 | 214.800 |
| overall__count__adverse__events | 144.000 | 147.000 | 157.000 | 175.250 | 186.200 |
| overall__not__infected__no__treatment__count | 0.000 | 0.000 | 0.000 | 0.000 | 0.000 |
| overall__not__infected__treated__count | 138.800 | 142.750 | 150.000 | 156.000 | 162.100 |
| overall__infected__no__treatment__count | 0.000 | 0.000 | 0.000 | 0.000 | 0.000 |
| overall__infected__treated__count | 337.900 | 344.000 | 350.000 | 357.250 | 361.200 |
| overall__sensitive__infection__treated__count__per__abx__dict__A | 44.900 | 48.000 | 55.500 | 63.250 | 71.200 |
| overall__sensitive__infection__treated__count__per__abx__dict__B | 48.900 | 56.500 | 87.000 | 96.250 | 101.100 |
| overall__resistant__infection__treated__count__per__abx__dict__A | 9.900 | 32.500 | 158.000 | 178.250 | 200.100 |
| overall__resistant__infection__treated__count__per__abx__dict__B | 11.000 | 18.250 | 45.500 | 191.750 | 217.700 |
| overall__sensitive__infection__treated__count | 111.500 | 126.750 | 138.000 | 148.250 | 154.100 |
| overall__resistant__infection__treated__count | 194.700 | 201.000 | 214.000 | 224.250 | 235.100 |
| overall__abx__prescriptions__count__per__abx__A | 78.900 | 103.250 | 213.000 | 228.250 | 250.300 |
| overall__abx__prescriptions__count__per__abx__B | 103.000 | 116.750 | 140.500 | 249.000 | 272.100 |
| overall__abx__prescriptions__count | 337.900 | 344.000 | 350.000 | 357.250 | 361.200 |
| final__amr__actual__A | 0.234 | 0.554 | 0.989 | 0.994 | 0.996 |
| final__amr__visible__A | 0.000 | 0.182 | 0.621 | 0.809 | 0.872 |
| final__amr__actual__B | 0.161 | 0.260 | 0.594 | 0.994 | 0.998 |
| final__amr__visible__B | 0.000 | 0.000 | 0.230 | 0.709 | 0.849 |

#### Two Antibiotics No Cross-Resistance, Recurrent PPO

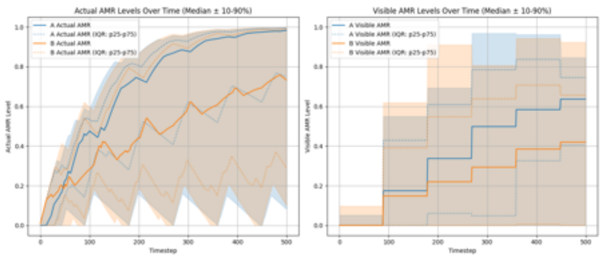

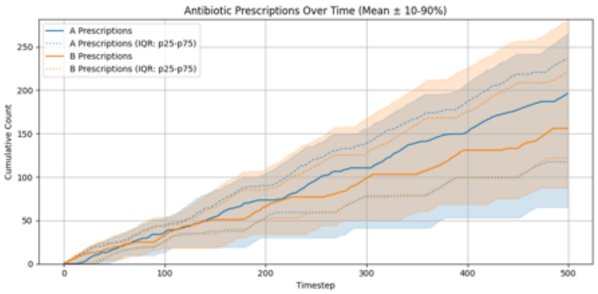

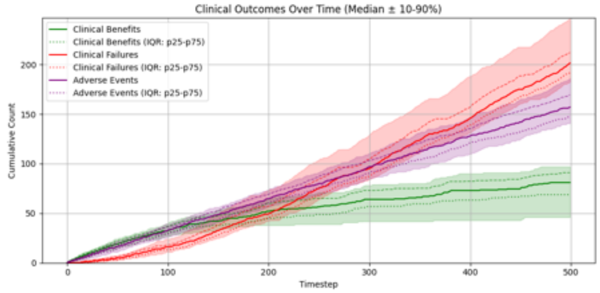

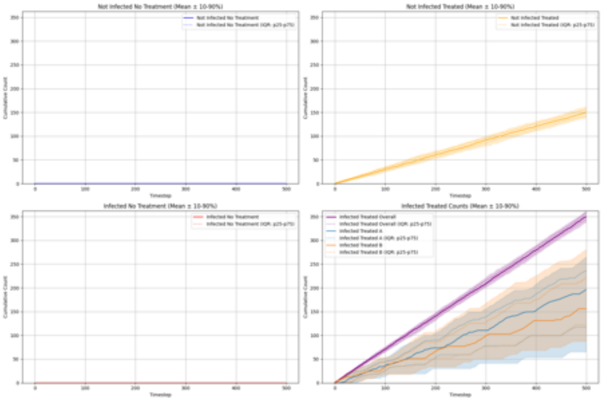

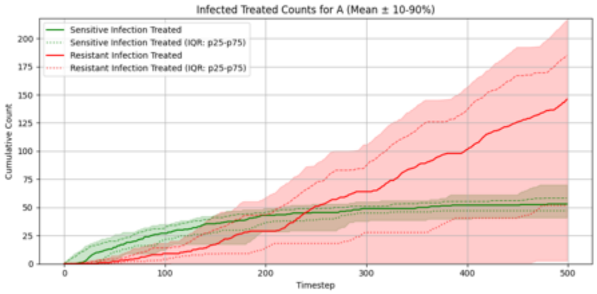

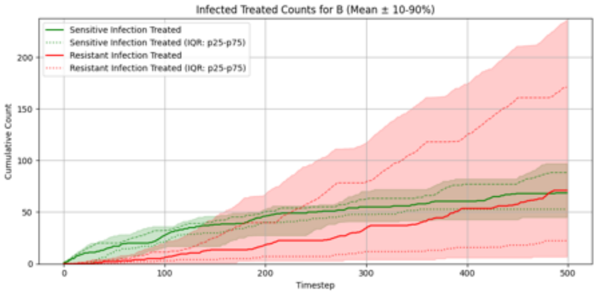

#### Summary Statistics (Two Antibiotics No Cross-Resistance, Recurrent PPO)

| metric | p10 | p25 | p50 | p75 | p90 |
|---|---|---|---|---|---|
| overall_total_reward | -299.362 | -241.815 | -215.781 | -204.134 | -185.579 |
| overall_count_clinical_benefits | 46.000 | 68.750 | 81.000 | 91.000 | 97.100 |
| overall_count_clinical_failures | 182.700 | 192.000 | 201.500 | 212.250 | 246.000 |
| overall_count_adverse_events | 140.800 | 147.750 | 157.000 | 169.250 | 186.100 |
| overall_not_infected_no_treatment_count | 0.000 | 0.000 | 0.000 | 0.000 | 0.000 |
| overall_not_infected_treated_count | 138.800 | 142.750 | 150.000 | 156.000 | 162.100 |
| overall_infected_no_treatment_count | 0.000 | 0.000 | 0.000 | 0.000 | 0.000 |
| overall_infected_treated_count | 337.900 | 344.000 | 349.500 | 357.000 | 361.200 |
| overall_sensitive_infection_treated_count_per_abx_dict_A | 40.900 | 47.000 | 53.000 | 58.250 | 70.100 |
| overall_sensitive_infection_treated_count_per_abx_dict_B | 45.000 | 52.750 | 68.500 | 88.250 | 97.100 |
| overall_resistant_infection_treated_count_per_abx_dict_A | 2.700 | 51.750 | 146.000 | 185.000 | 216.700 |
| overall_resistant_infection_treated_count_per_abx_dict_B | 6.800 | 22.250 | 71.000 | 171.000 | 236.400 |
| overall_sensitive_infection_treated_count | 77.600 | 111.000 | 129.000 | 140.000 | 150.100 |
| overall_resistant_infection_treated_count | 203.000 | 210.750 | 222.000 | 234.250 | 272.400 |
| overall_abx_prescriptions_count_per_abx_A | 65.200 | 117.500 | 196.000 | 236.250 | 265.000 |
| overall_abx_prescriptions_count_per_abx_B | 87.300 | 121.750 | 156.000 | 221.750 | 280.500 |
| overall_abx_prescriptions_count | 337.900 | 344.000 | 349.500 | 357.000 | 361.200 |
| final_amr_actual_A | 0.093 | 0.753 | 0.982 | 0.994 | 0.997 |
| final_amr_visible_A | 0.000 | 0.405 | 0.636 | 0.745 | 0.846 |
| final_amr_actual_B | 0.120 | 0.308 | 0.742 | 0.987 | 0.998 |
| final_amr_visible_B | 0.000 | 0.000 | 0.419 | 0.657 | 0.924 |

#### Two Antibiotics No Cross-Resistance, Hierarchical PPO

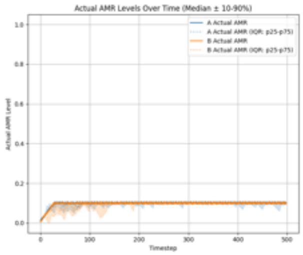

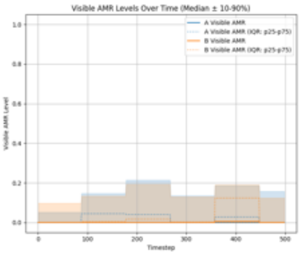

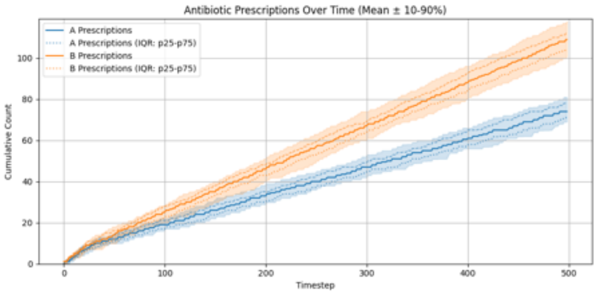

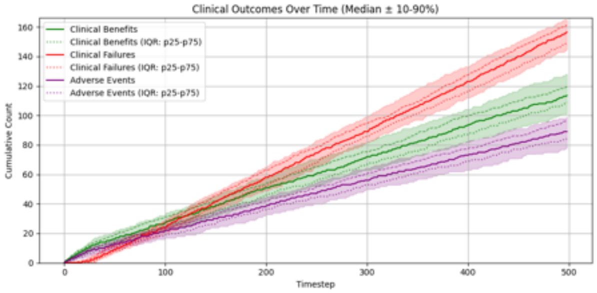

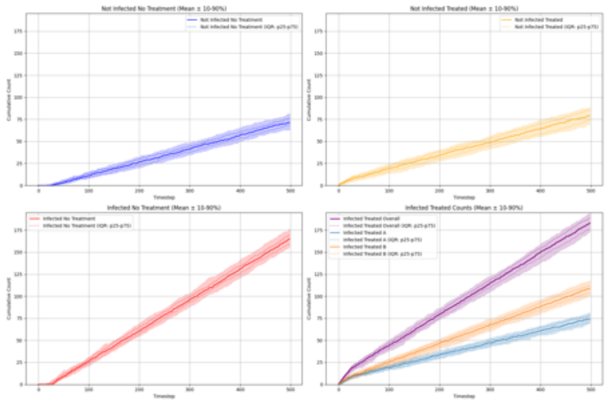

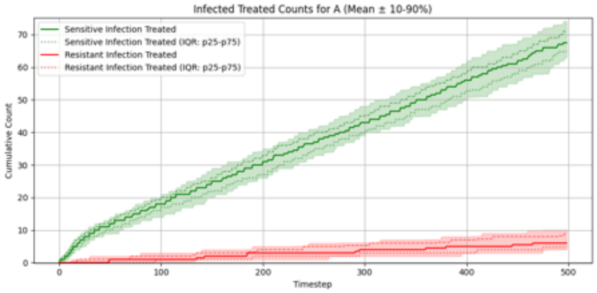

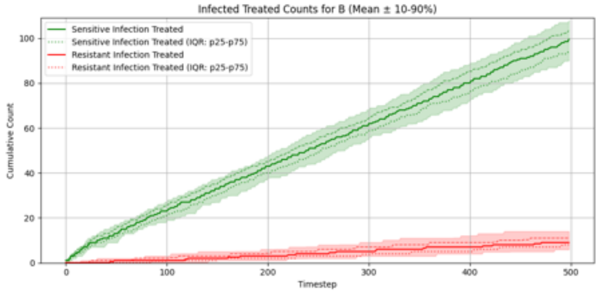

#### Summary Statistics (Two Antibiotics No Cross-Resistance, Hierarchical PPO)

| metric | p10 | p25 | p50 | p75 | p90 |
|---|---|---|---|---|---|
| overall_total_reward | -112.084 | -102.412 | -97.289 | -82.617 | -74.104 |
| overall_count_clinical_benefits | 101.000 | 109.000 | 113.500 | 119.250 | 128.000 |
| overall_count_clinical_failures | 144.900 | 149.000 | 156.500 | 161.000 | 165.100 |
| overall_count_adverse_events | 77.900 | 84.000 | 89.000 | 96.000 | 99.000 |
| overall_not_infected_no_treatment_count | 63.000 | 69.000 | 71.500 | 76.000 | 81.000 |
| overall_not_infected_treated_count | 68.900 | 74.000 | 79.000 | 86.000 | 88.100 |
| overall_infected_no_treatment_count | 156.000 | 160.000 | 165.000 | 169.250 | 176.100 |
| overall_infected_treated_count | 173.700 | 177.000 | 183.000 | 189.000 | 194.000 |
| overall_sensitive_infection_treated_count_per_abx_dict_A | 62.900 | 64.750 | 67.500 | 71.000 | 74.100 |
| overall_sensitive_infection_treated_count_per_abx_dict_B | 90.000 | 94.000 | 99.500 | 103.250 | 108.000 |
| overall_resistant_infection_treated_count_per_abx_dict_A | 4.000 | 4.750 | 6.000 | 9.000 | 10.100 |
| overall_resistant_infection_treated_count_per_abx_dict_B | 5.900 | 8.000 | 9.000 | 11.250 | 14.000 |
| overall_sensitive_infection_treated_count | 156.900 | 163.250 | 166.000 | 171.250 | 179.000 |
| overall_resistant_infection_treated_count | 11.000 | 13.000 | 16.000 | 19.000 | 22.100 |
| overall_abx_prescriptions_count_per_abx_A | 69.000 | 71.000 | 74.000 | 78.000 | 81.000 |
| overall_abx_prescriptions_count_per_abx_B | 100.900 | 103.750 | 109.000 | 112.000 | 118.000 |
| overall_abx_prescriptions_count | 173.700 | 177.000 | 183.000 | 189.000 | 194.000 |
| final_amr_actual_A | 0.088 | 0.101 | 0.102 | 0.102 | 0.102 |
| final_amr_visible_A | 0.000 | 0.000 | 0.000 | 0.000 | 0.157 |
| final_amr_actual_B | 0.100 | 0.100 | 0.100 | 0.100 | 0.100 |
| final_amr_visible_B | 0.000 | 0.000 | 0.000 | 0.000 | 0.124 |

#### Two Antibiotics No Cross-Resistance, Hierarchical Recurrent PPO

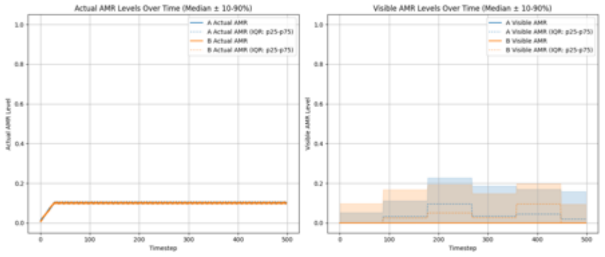

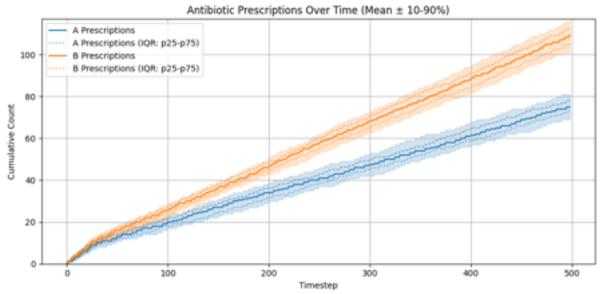

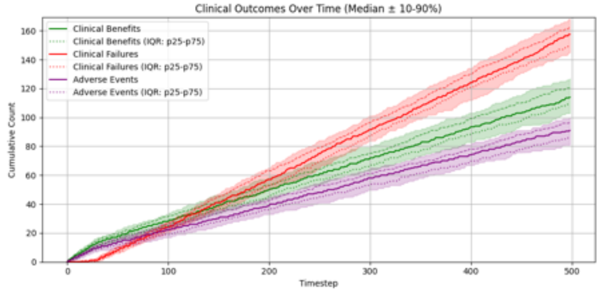

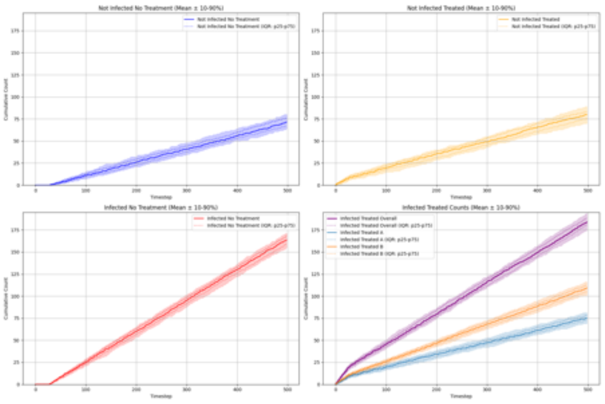

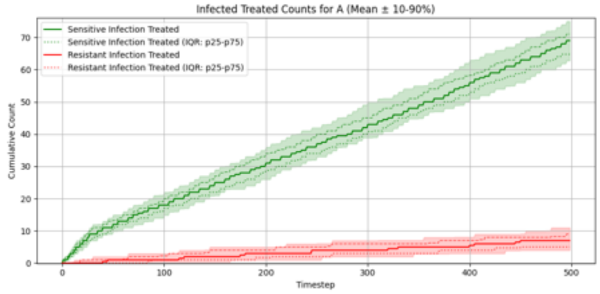

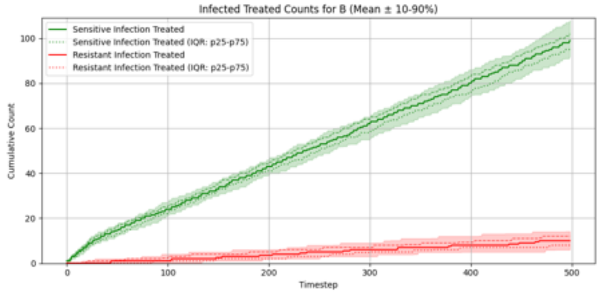

#### Summary Statistics (Two Antibiotics No Cross-Resistance, Hierarchical Recurrent PPO)

| metric | p10 | p25 | p50 | p75 | p90 |
|---|---|---|---|---|---|
| overall_total_reward | -112.858 | -105.868 | -98.544 | -83.052 | -76.855 |
| overall_count_clinical_benefits | 102.900 | 109.000 | 114.000 | 120.250 | 127.000 |
| overall_count_clinical_failures | 144.900 | 150.000 | 157.500 | 162.000 | 168.200 |
| overall_count_adverse_events | 81.000 | 86.000 | 91.000 | 97.000 | 100.000 |
| overall_not_infected_no_treatment_count | 63.900 | 66.750 | 71.500 | 76.000 | 80.100 |
| overall_not_infected_treated_count | 70.000 | 76.000 | 80.000 | 83.250 | 89.000 |
| overall_infected_no_treatment_count | 154.900 | 159.000 | 163.500 | 168.250 | 171.100 |
| overall_infected_treated_count | 175.000 | 180.750 | 184.000 | 188.000 | 194.000 |
| overall_sensitive_infection_treated_count_per_abx_dict_A | 62.800 | 64.750 | 69.000 | 71.000 | 75.000 |
| overall_sensitive_infection_treated_count_per_abx_dict_B | 92.000 | 95.000 | 99.000 | 102.000 | 108.000 |
| overall_resistant_infection_treated_count_per_abx_dict_A | 4.000 | 5.000 | 7.000 | 9.000 | 11.000 |
| overall_resistant_infection_treated_count_per_abx_dict_B | 6.000 | 8.000 | 10.000 | 12.000 | 14.100 |
| overall_sensitive_infection_treated_count | 156.900 | 163.000 | 168.000 | 171.000 | 176.100 |
| overall_resistant_infection_treated_count | 11.900 | 14.750 | 17.500 | 20.000 | 23.100 |
| overall_abx_prescriptions_count_per_abx_A | 69.000 | 73.000 | 75.000 | 78.000 | 81.100 |
| overall_abx_prescriptions_count_per_abx_B | 101.000 | 105.750 | 109.500 | 112.250 | 116.100 |
| overall_abx_prescriptions_count | 175.000 | 180.750 | 184.000 | 188.000 | 194.000 |
| final_amr_actual_A | 0.102 | 0.102 | 0.102 | 0.102 | 0.102 |
| final_amr_visible_A | 0.000 | 0.000 | 0.000 | 0.019 | 0.159 |
| final_amr_actual_B | 0.100 | 0.100 | 0.100 | 0.100 | 0.100 |
| final_amr_visible_B | 0.000 | 0.000 | 0.000 | 0.000 | 0.094 |

#### Two Antibiotics With Cross-Resistance, Flat PPO

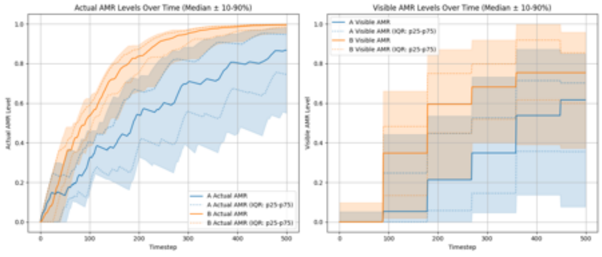

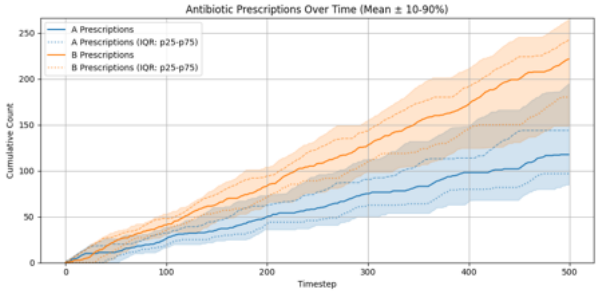

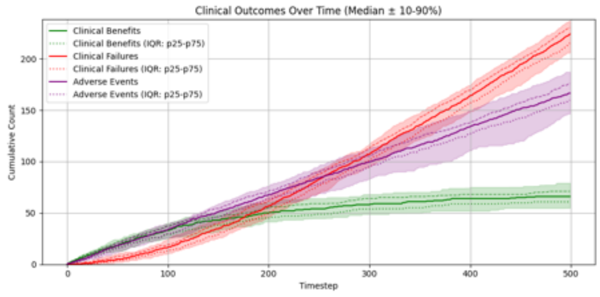

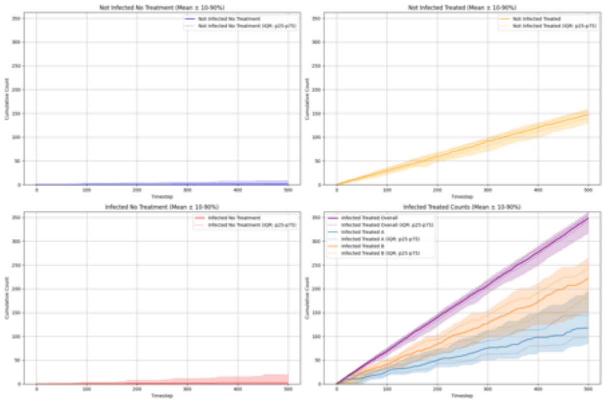

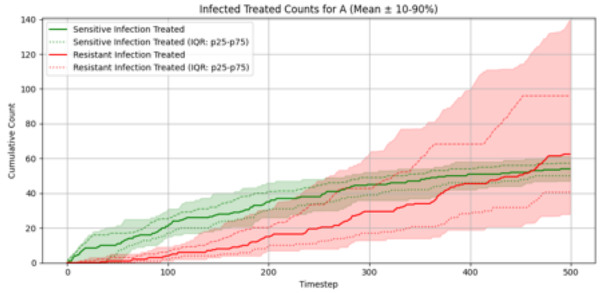

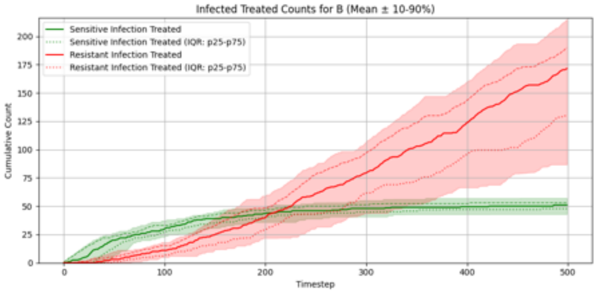

#### Summary Statistics (Two Antibiotics With Cross-Resistance, Flat PPO)

| metric | p10 | p25 | p50 | p75 | p90 |
|---|---|---|---|---|---|
| overall_total_reward | -275.331 | -265.121 | -252.424 | -237.636 | -226.897 |
| overall_count_clinical_benefits | 54.800 | 60.750 | 66.000 | 71.000 | 79.100 |
| overall_count_clinical_failures | 205.800 | 215.250 | 224.000 | 230.250 | 237.200 |
| overall_count_adverse_events | 146.500 | 159.750 | 166.500 | 176.000 | 187.200 |
| overall_not_infected_no_treatment_count | 0.000 | 0.000 | 0.000 | 1.250 | 8.300 |
| overall_not_infected_treated_count | 129.600 | 139.000 | 147.000 | 155.250 | 160.200 |
| overall_infected_no_treatment_count | 0.000 | 0.000 | 0.000 | 3.000 | 19.700 |
| overall_infected_treated_count | 316.900 | 340.750 | 348.000 | 355.000 | 361.000 |
| overall_sensitive_infection_treated_count_per_abx_dict_A | 47.000 | 50.000 | 54.000 | 57.250 | 61.200 |
| overall_sensitive_infection_treated_count_per_abx_dict_B | 43.000 | 47.750 | 51.000 | 53.000 | 57.100 |
| overall_resistant_infection_treated_count_per_abx_dict_A | 27.900 | 40.750 | 62.500 | 96.000 | 139.800 |
| overall_resistant_infection_treated_count_per_abx_dict_B | 86.800 | 129.750 | 171.500 | 189.750 | 215.200 |
| overall_sensitive_infection_treated_count | 89.800 | 98.000 | 103.500 | 109.250 | 115.100 |
| overall_resistant_infection_treated_count | 215.000 | 233.750 | 245.500 | 250.250 | 257.100 |
| overall_abx_prescriptions_count_per_abx_A | 84.700 | 96.750 | 117.500 | 143.750 | 194.800 |
| overall_abx_prescriptions_count_per_abx_B | 148.700 | 180.000 | 221.500 | 242.250 | 264.900 |
| overall_abx_prescriptions_count | 316.900 | 340.750 | 348.000 | 355.000 | 361.000 |
| final_amr_actual_A | 0.554 | 0.748 | 0.865 | 0.947 | 0.981 |
| final_amr_visible_A | 0.077 | 0.355 | 0.617 | 0.702 | 0.850 |
| final_amr_actual_B | 0.955 | 0.985 | 0.996 | 0.998 | 0.999 |
| final_amr_visible_B | 0.373 | 0.599 | 0.754 | 0.854 | 0.959 |

#### Two Antibiotics With Cross-Resistance, Recurrent PPO

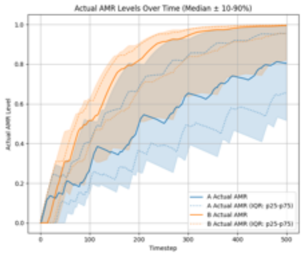

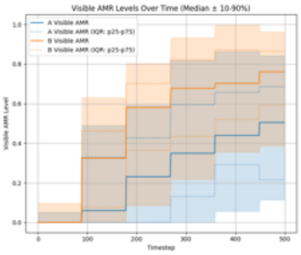

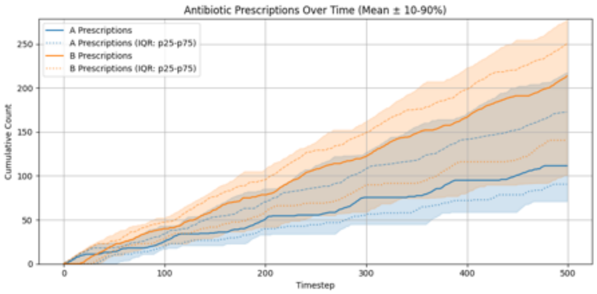

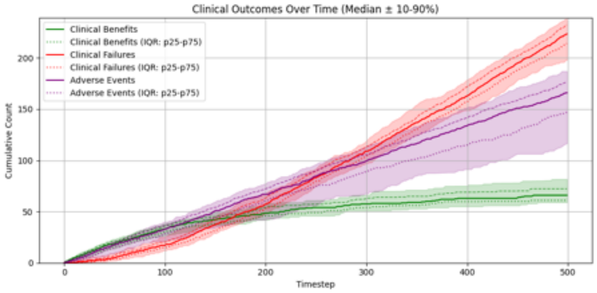

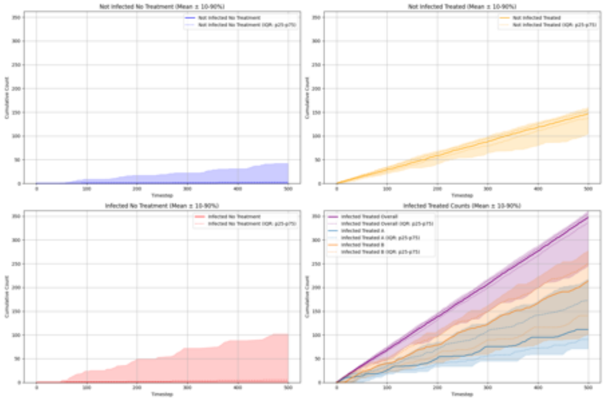

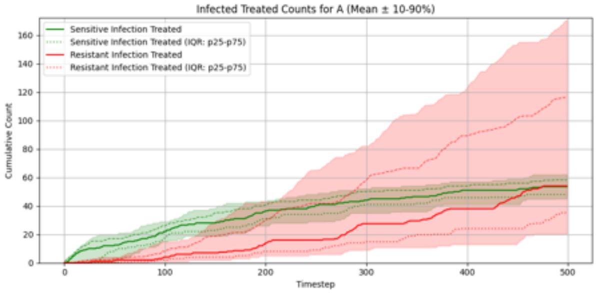

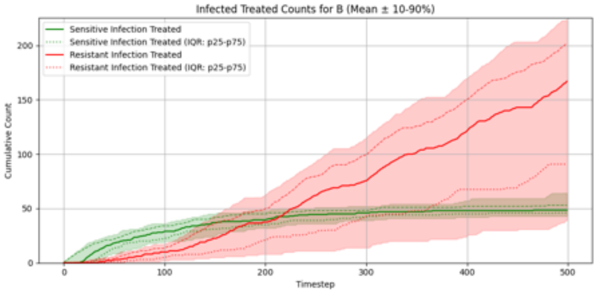

#### Summary Statistics (Two Antibiotics With Cross-Resistance, Recurrent PPO)

| metric | p10 | p25 | p50 | p75 | p90 |
|---|---|---|---|---|---|
| overall_total_reward | -274.038 | -267.750 | -255.277 | -241.974 | -187.639 |
| overall_count_clinical_benefits | 59.000 | 61.000 | 66.000 | 72.250 | 81.700 |
| overall_count_clinical_failures | 198.000 | 214.250 | 223.500 | 232.250 | 238.100 |
| overall_count_adverse_events | 116.500 | 146.750 | 166.000 | 176.250 | 187.100 |
| overall_not_infected_no_treatment_count | 0.000 | 0.000 | 0.000 | 2.500 | 42.200 |
| overall_not_infected_treated_count | 105.100 | 137.000 | 147.000 | 154.000 | 160.000 |
| overall_infected_no_treatment_count | 0.000 | 0.000 | 0.000 | 5.000 | 102.100 |
| overall_infected_treated_count | 244.900 | 336.250 | 347.000 | 353.500 | 361.000 |
| overall_sensitive_infection_treated_count_per_abx_dict_A | 45.000 | 48.000 | 53.500 | 58.250 | 62.100 |
| overall_sensitive_infection_treated_count_per_abx_dict_B | 42.900 | 46.000 | 48.500 | 53.000 | 64.200 |
| overall_resistant_infection_treated_count_per_abx_dict_A | 20.400 | 35.250 | 54.000 | 116.750 | 171.100 |
| overall_resistant_infection_treated_count_per_abx_dict_B | 38.900 | 90.750 | 167.000 | 201.500 | 224.300 |
| overall_sensitive_infection_treated_count | 91.900 | 97.000 | 103.000 | 108.250 | 126.000 |
| overall_resistant_infection_treated_count | 130.400 | 231.000 | 246.500 | 253.750 | 261.300 |
| overall_abx_prescriptions_count_per_abx_A | 71.000 | 90.500 | 111.500 | 172.500 | 217.700 |
| overall_abx_prescriptions_count_per_abx_B | 101.100 | 140.500 | 213.500 | 249.750 | 277.400 |
| overall_abx_prescriptions_count | 244.900 | 336.250 | 347.000 | 353.500 | 361.000 |
| final_amr_actual_A | 0.521 | 0.653 | 0.806 | 0.955 | 0.993 |
| final_amr_visible_A | 0.114 | 0.216 | 0.505 | 0.686 | 0.843 |
| final_amr_actual_B | 0.819 | 0.954 | 0.995 | 0.998 | 0.999 |
| final_amr_visible_B | 0.387 | 0.593 | 0.761 | 0.865 | 0.963 |

#### Two Antibiotics With Cross-Resistance, Hierarchical PPO

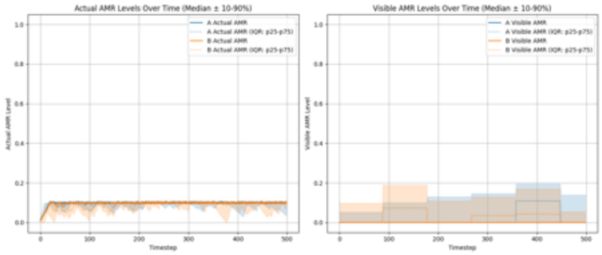

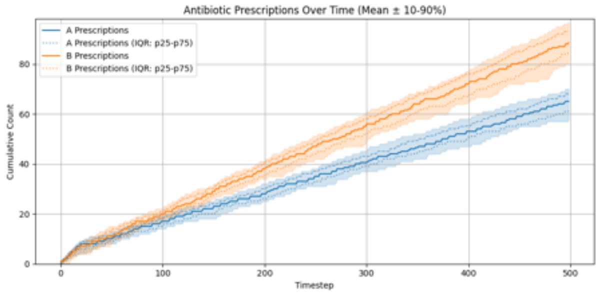

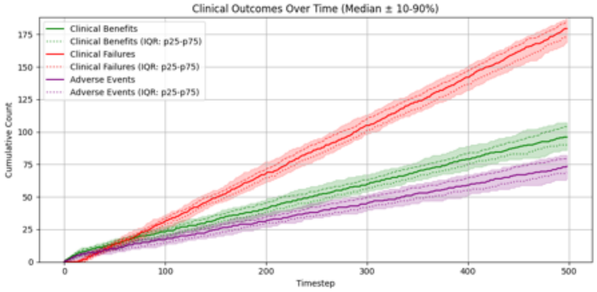

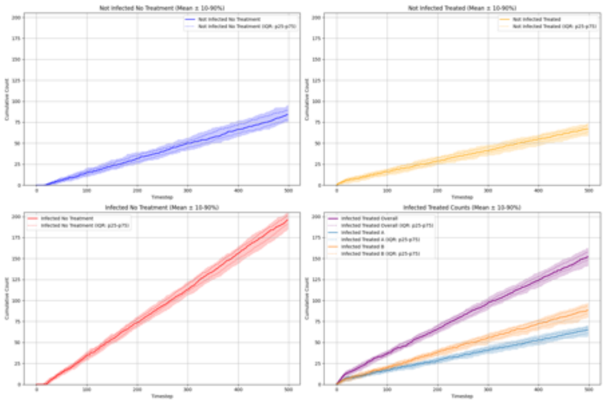

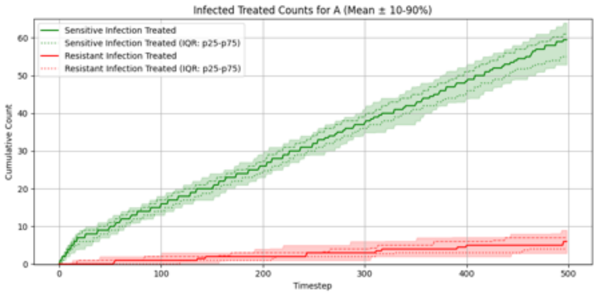

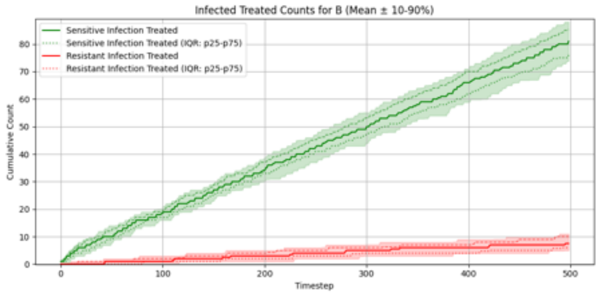

#### Summary Statistics (Two Antibiotics With Cross-Resistance, Hierarchical PPO)

| metric | p10 | p25 | p50 | p75 | p90 |
|---|---|---|---|---|---|
| overall__total__reward | -141.785 | -134.697 | -126.114 | -115.900 | -107.514 |
| overall__count__clinical__benefits | 86.000 | 90.000 | 96.000 | 104.000 | 107.200 |
| overall__count__clinical__failures | 168.800 | 173.750 | 179.500 | 185.000 | 187.100 |
| overall__count__adverse__events | 63.900 | 68.750 | 73.500 | 80.000 | 82.100 |
| overall__not__infected__no__treatment__count | 75.000 | 78.750 | 84.000 | 89.250 | 95.100 |
| overall__not__infected__treated__count | 59.000 | 63.750 | 67.000 | 71.250 | 74.000 |
| overall__infected__no__treatment__count | 184.900 | 191.000 | 196.000 | 201.250 | 204.100 |
| overall__infected__treated__count | 142.000 | 148.000 | 152.000 | 158.000 | 162.200 |
| overall__sensitive__infection__treated__count__per__abx__dict__A | 52.900 | 55.000 | 59.500 | 61.000 | 64.000 |
| overall__sensitive__infection__treated__count__per__abx__dict__B | 73.800 | 76.000 | 81.000 | 85.250 | 88.100 |
| overall__resistant__infection__treated__count__per__abx__dict__A | 2.900 | 4.000 | 6.000 | 7.000 | 9.000 |
| overall__resistant__infection__treated__count__per__abx__dict__B | 5.000 | 6.000 | 7.500 | 10.000 | 11.000 |
| overall__sensitive__infection__treated__count | 131.000 | 133.000 | 140.000 | 143.000 | 148.100 |
| overall__resistant__infection__treated__count | 9.000 | 10.750 | 13.500 | 16.000 | 18.000 |
| overall__abx__prescriptions__count__per__abx__A | 57.900 | 61.000 | 65.000 | 68.000 | 70.000 |
| overall__abx__prescriptions__count__per__abx__B | 79.900 | 84.750 | 88.500 | 93.000 | 97.000 |
| overall__abx__prescriptions__count | 142.000 | 148.000 | 152.000 | 158.000 | 162.200 |
| final__amr__actual__A | 0.044 | 0.100 | 0.100 | 0.101 | 0.102 |
| final__amr__visible__A | 0.000 | 0.000 | 0.000 | 0.000 | 0.139 |
| final__amr__actual__B | 0.083 | 0.100 | 0.101 | 0.101 | 0.101 |
| final__amr__visible__B | 0.000 | 0.000 | 0.000 | 0.000 | 0.054 |

#### Two Antibiotics With Cross-Resistance, Hierarchical Recurrent PPO

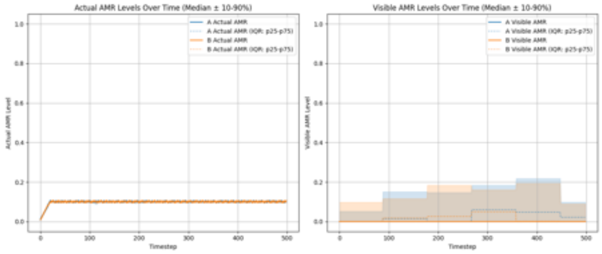

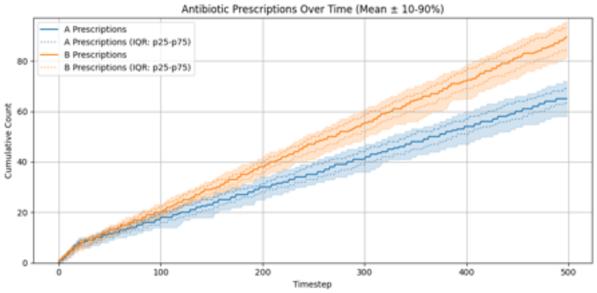

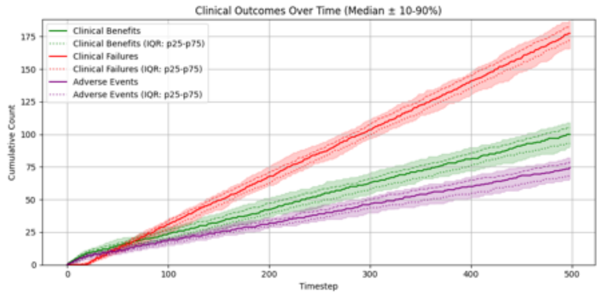

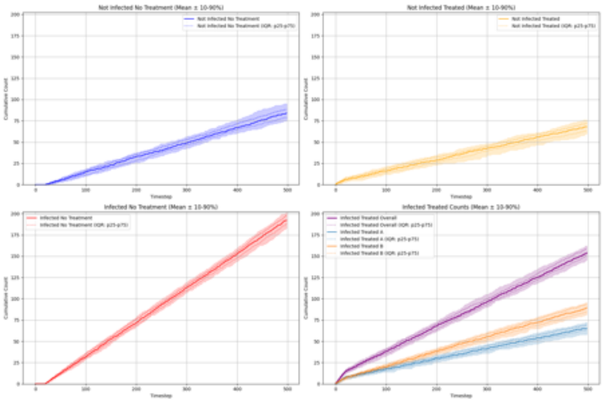

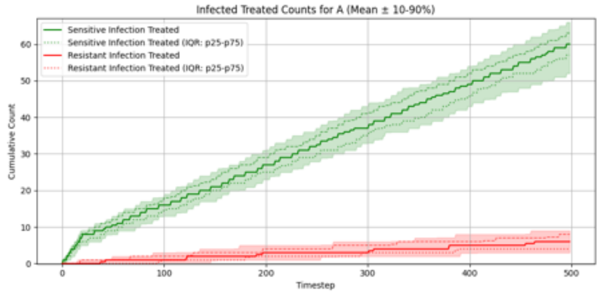

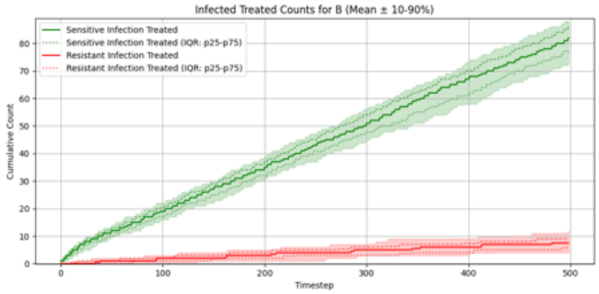

##### Summary Statistics (Two Antibiotics With Cross-Resistance, Hierarchical Recurrent PPO)

| metric | p10 | p25 | p50 | p75 | p90 |
|---|---|---|---|---|---|
| overall_total_reward | -138.822 | -128.776 | -119.819 | -116.246 | -104.535 |
| overall_count_clinical_benefits | 90.800 | 93.000 | 100.000 | 105.000 | 109.000 |
| overall_count_clinical_failures | 165.800 | 172.000 | 177.500 | 182.500 | 187.100 |
| overall_count_adverse_events | 66.000 | 68.000 | 74.500 | 79.000 | 82.100 |
| overall_not_infected_no_treatment_count | 74.800 | 78.750 | 84.000 | 88.250 | 95.100 |
| overall_not_infected_treated_count | 59.700 | 63.000 | 68.000 | 72.000 | 76.100 |
| overall_infected_no_treatment_count | 182.800 | 188.000 | 192.000 | 198.000 | 201.200 |
| overall_infected_treated_count | 145.900 | 150.750 | 154.000 | 159.250 | 162.100 |
| overall_sensitive_infection_treated_count_per_abx_dict_A | 52.000 | 57.000 | 60.000 | 63.000 | 66.000 |
| overall_sensitive_infection_treated_count_per_abx_dict_B | 72.900 | 78.000 | 82.000 | 86.000 | 88.000 |
| overall_resistant_infection_treated_count_per_abx_dict_A | 3.000 | 4.000 | 6.000 | 8.000 | 9.000 |
| overall_resistant_infection_treated_count_per_abx_dict_B | 4.000 | 5.750 | 7.500 | 9.000 | 12.000 |
| overall_sensitive_infection_treated_count | 131.000 | 135.750 | 142.000 | 146.000 | 151.000 |
| overall_resistant_infection_treated_count | 9.000 | 11.000 | 14.000 | 16.000 | 18.100 |
| overall_abx_prescriptions_count_per_abx_A | 58.000 | 63.750 | 65.000 | 69.000 | 72.000 |
| overall_abx_prescriptions_count_per_abx_B | 81.900 | 85.000 | 89.500 | 93.000 | 96.000 |
| overall_abx_prescriptions_count | 145.900 | 150.750 | 154.000 | 159.250 | 162.100 |
| final_amr_actual_A | 0.099 | 0.100 | 0.100 | 0.100 | 0.100 |
| final_amr_visible_A | 0.000 | 0.000 | 0.000 | 0.022 | 0.098 |
| final_amr_actual_B | 0.101 | 0.101 | 0.101 | 0.101 | 0.101 |
| final_amr_visible_B | 0.000 | 0.000 | 0.000 | 0.000 | 0.090 |

#### Experiment Set 3

##### Single Antibiotic, Accurate Risk Stratification

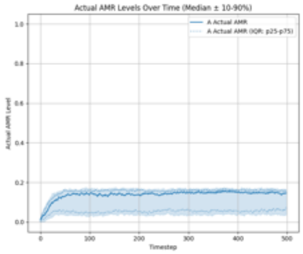

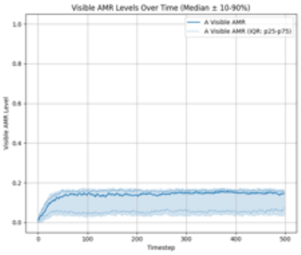

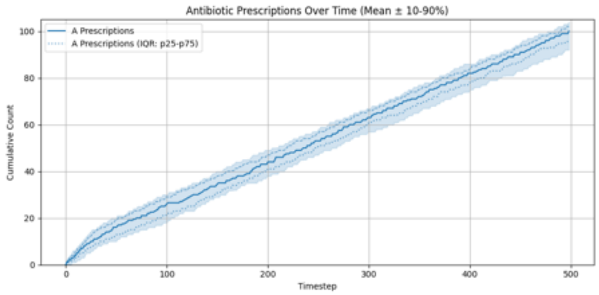

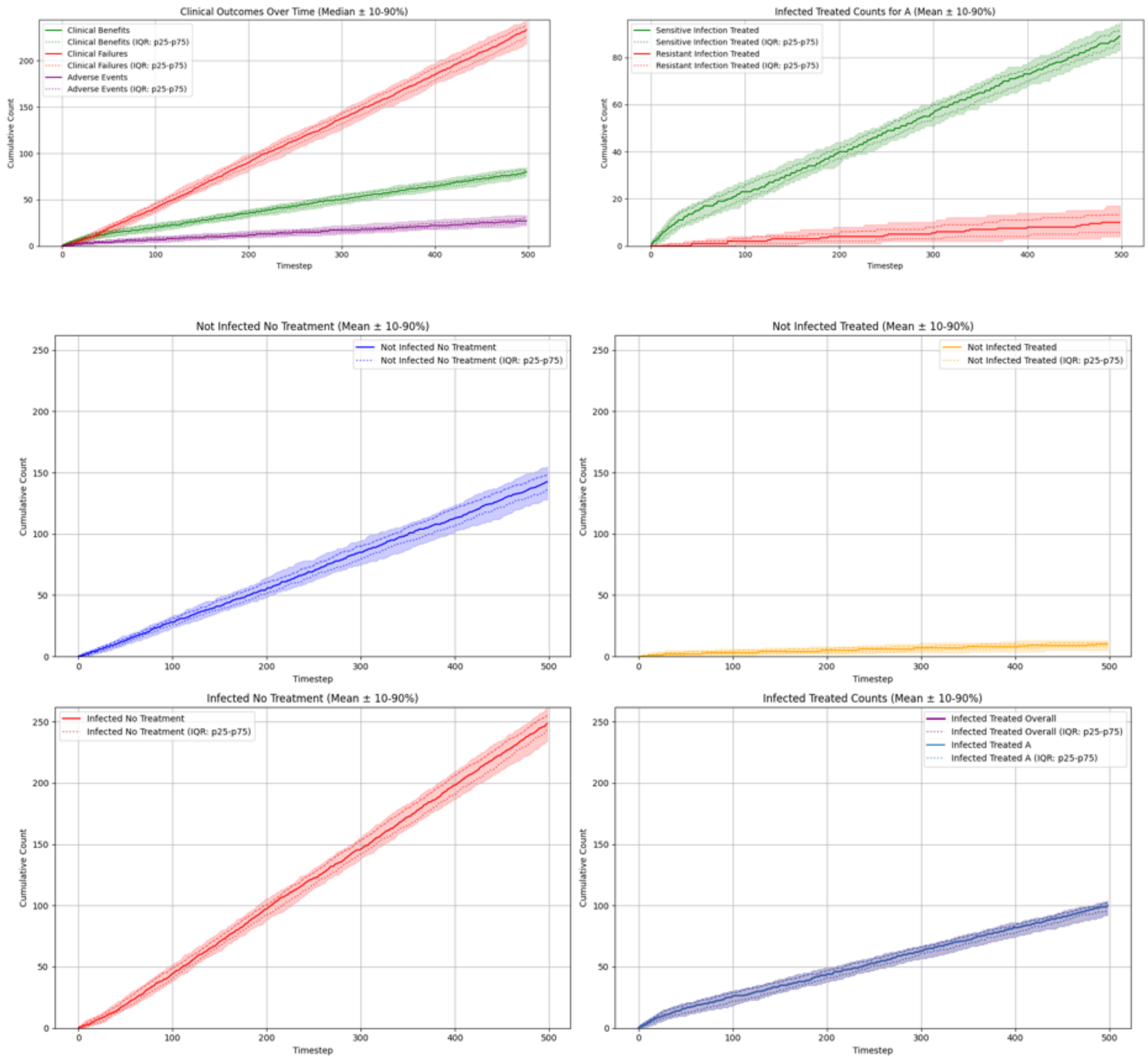

#### Summary Statistics (Single Antibiotic, Accurate Risk Stratification)

| metric | p10 | p25 | p50 | p75 | p90 |
|---|---|---|---|---|---|
| overall_total_reward | -197.287 | -185.317 | -179.264 | -169.768 | -158.447 |
| overall_count_clinical_benefits | 74.900 | 77.000 | 80.000 | 82.000 | 85.000 |
| overall_count_clinical_failures | 218.700 | 225.500 | 233.500 | 238.250 | 243.100 |
| overall_count_adverse_events | 22.000 | 24.000 | 27.000 | 30.000 | 33.000 |
| overall_not_infected_no_treatment_count | 128.000 | 135.750 | 142.500 | 148.000 | 155.200 |
| overall_not_infected_treated_count | 5.000 | 8.000 | 10.000 | 11.000 | 13.000 |
| overall_infected_no_treatment_count | 233.900 | 242.750 | 248.500 | 255.250 | 261.000 |
| overall_infected_treated_count | 92.000 | 96.000 | 100.000 | 102.000 | 104.000 |
| overall_sensitive_infection_treated_count_per_abx_dict_A | 83.000 | 86.000 | 89.000 | 91.250 | 95.000 |
| overall_resistant_infection_treated_count_per_abx_dict_A | 3.900 | 5.750 | 10.000 | 13.250 | 17.000 |
| overall_sensitive_infection_treated_count | 83.000 | 86.000 | 89.000 | 91.250 | 95.000 |
| overall_resistant_infection_treated_count | 3.900 | 5.750 | 10.000 | 13.250 | 17.000 |
| overall_abx_prescriptions_count_per_abx_A | 92.000 | 96.000 | 100.000 | 102.000 | 104.000 |
| overall_abx_prescriptions_count | 92.000 | 96.000 | 100.000 | 102.000 | 104.000 |
| final_amr_actual_A | 0.034 | 0.062 | 0.146 | 0.156 | 0.165 |
| final_amr_visible_A | 0.034 | 0.062 | 0.146 | 0.156 | 0.165 |

#### Single Antibiotic, Exaggerated Risk Stratification

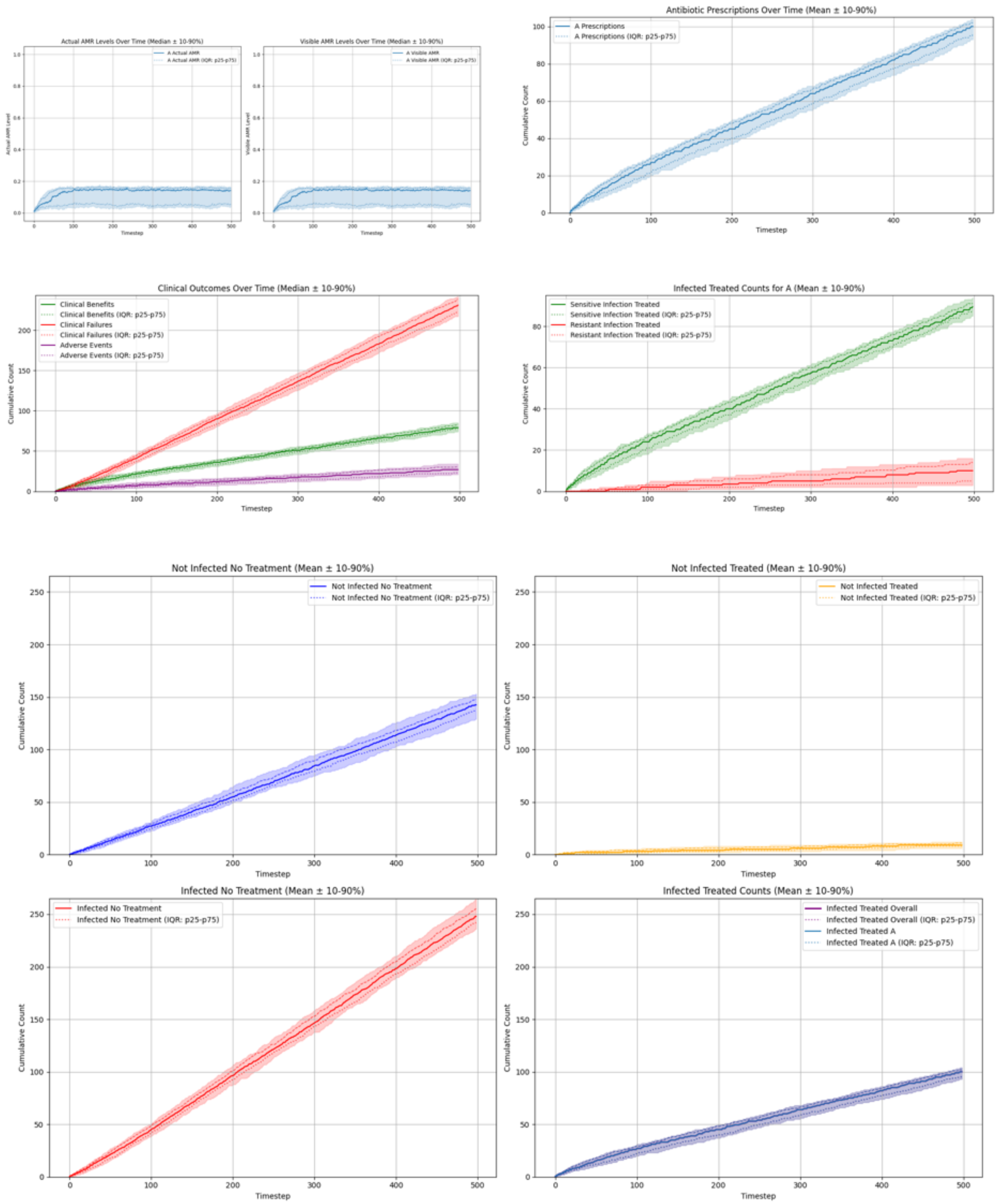

#### Summary Statistics (Single Antibiotic, Exaggerated Risk Stratification)

| metric | p10 | p25 | p50 | p75 | p90 |
|---|---|---|---|---|---|
| overall_total_reward | -188.528 | -181.568 | -174.766 | -165.119 | -158.988 |
| overall_count_clinical_benefits | 74.000 | 77.000 | 79.000 | 83.000 | 85.100 |
| overall_count_clinical_failures | 217.000 | 222.750 | 231.000 | 238.000 | 242.100 |
| overall_count_adverse_events | 20.900 | 23.000 | 27.000 | 30.250 | 34.100 |
| overall_not_infected_no_treatment_count | 129.000 | 137.500 | 142.500 | 148.000 | 153.100 |
| overall_not_infected_treated_count | 6.000 | 8.000 | 9.000 | 11.000 | 12.000 |
| overall_infected_no_treatment_count | 236.000 | 242.000 | 248.000 | 255.000 | 264.100 |
| overall_infected_treated_count | 93.000 | 95.750 | 100.000 | 102.000 | 104.000 |
| overall_sensitive_infection_treated_count_per_abx_dict_A | 85.000 | 87.750 | 89.500 | 91.000 | 94.000 |
| overall_resistant_infection_treated_count_per_abx_dict_A | 2.900 | 5.000 | 10.000 | 14.000 | 16.000 |
| overall_sensitive_infection_treated_count | 85.000 | 87.750 | 89.500 | 91.000 | 94.000 |
| overall_resistant_infection_treated_count | 2.900 | 5.000 | 10.000 | 14.000 | 16.000 |
| overall_abx_prescriptions_count_per_abx_A | 93.000 | 95.750 | 100.000 | 102.000 | 104.000 |
| overall_abx_prescriptions_count | 93.000 | 95.750 | 100.000 | 102.000 | 104.000 |
| final_amr_actual_A | 0.038 | 0.052 | 0.143 | 0.152 | 0.161 |
| final_amr_visible_A | 0.038 | 0.052 | 0.143 | 0.152 | 0.161 |

#### Single Antibiotic, Compressed Risk Stratification

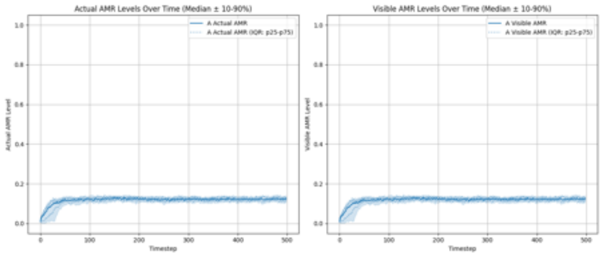

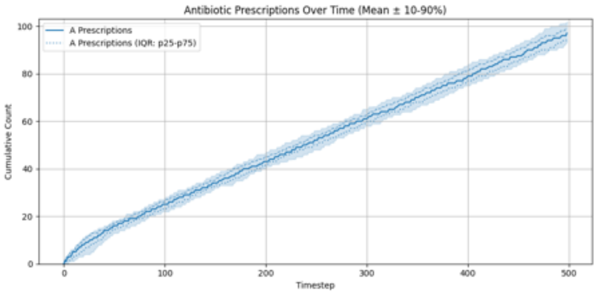

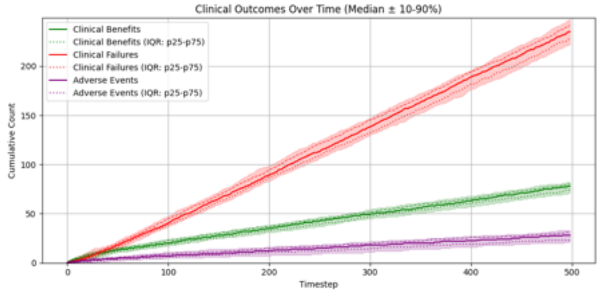

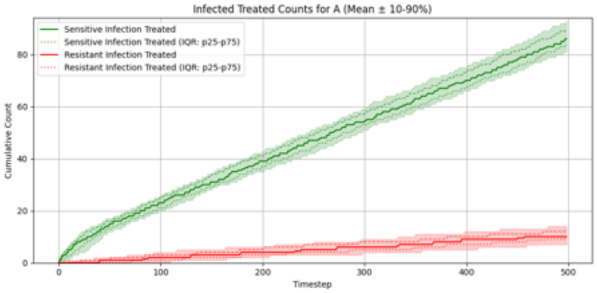

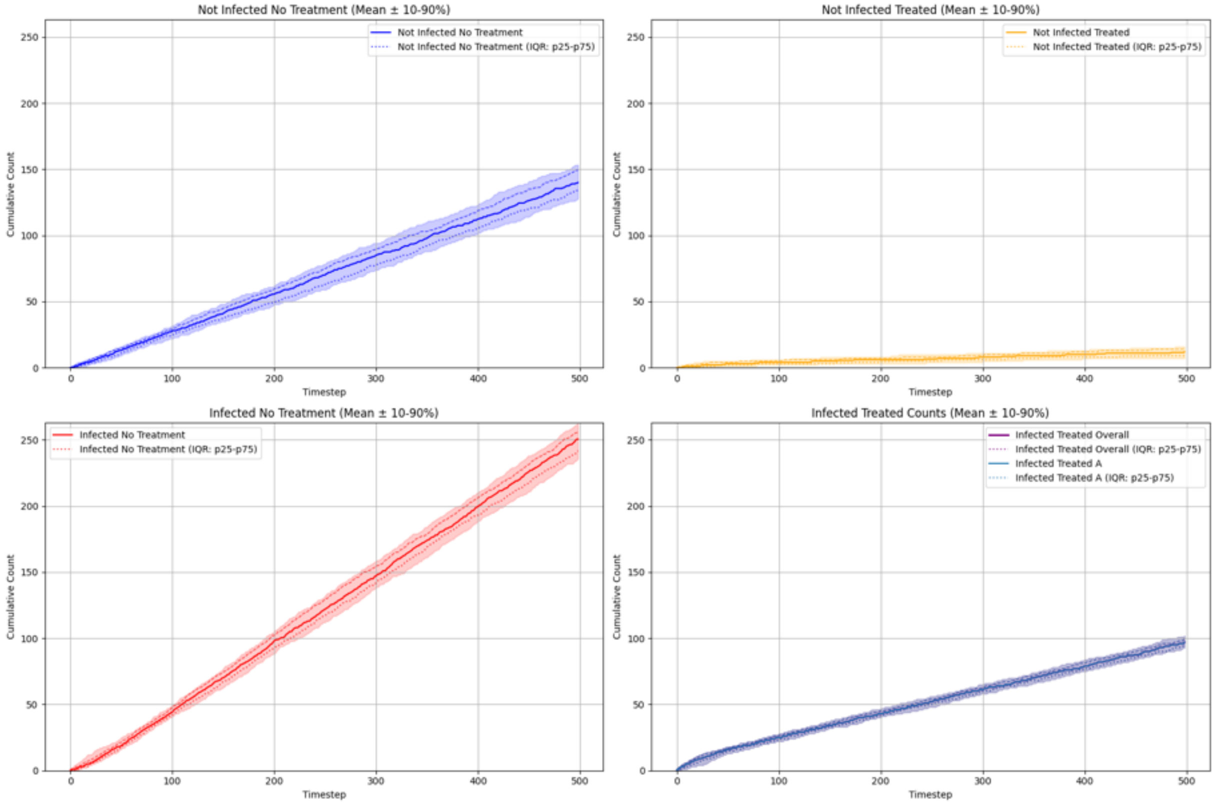

#### Summary Statistics (Single Antibiotic, Compressed Risk Stratification)

| metric | p10 | p25 | p50 | p75 | p90 |
|---|---|---|---|---|---|
| overall_total_reward | -202.726 | -193.917 | -184.367 | -177.617 | -171.446 |
| overall_count_clinical_benefits | 71.000 | 74.000 | 78.000 | 80.000 | 82.000 |
| overall_count_clinical_failures | 221.900 | 227.750 | 235.000 | 241.000 | 248.000 |
| overall_count_adverse_events | 20.900 | 23.750 | 28.000 | 31.000 | 33.000 |
| overall_not_infected_no_treatment_count | 127.900 | 134.000 | 140.000 | 149.250 | 153.400 |
| overall_not_infected_treated_count | 7.000 | 9.000 | 12.000 | 14.250 | 16.000 |
| overall_infected_no_treatment_count | 235.800 | 241.750 | 250.500 | 256.250 | 262.100 |
| overall_infected_treated_count | 92.900 | 94.000 | 97.000 | 99.000 | 102.000 |
| overall_sensitive_infection_treated_count_per_abx_dict_A | 81.000 | 83.750 | 86.000 | 89.250 | 93.000 |
| overall_resistant_infection_treated_count_per_abx_dict_A | 7.000 | 9.000 | 10.000 | 12.000 | 14.000 |
| overall_sensitive_infection_treated_count | 81.000 | 83.750 | 86.000 | 89.250 | 93.000 |
| overall_resistant_infection_treated_count | 7.000 | 9.000 | 10.000 | 12.000 | 14.000 |
| overall_abx_prescriptions_count_per_abx_A | 92.900 | 94.000 | 97.000 | 99.000 | 102.000 |
| overall_abx_prescriptions_count | 92.900 | 94.000 | 97.000 | 99.000 | 102.000 |
| final_amr_actual_A | 0.106 | 0.112 | 0.123 | 0.129 | 0.133 |
| final_amr_visible_A | 0.106 | 0.112 | 0.123 | 0.129 | 0.133 |

##### Two Antibiotics No Cross-Resistance, Accurate Risk Stratification

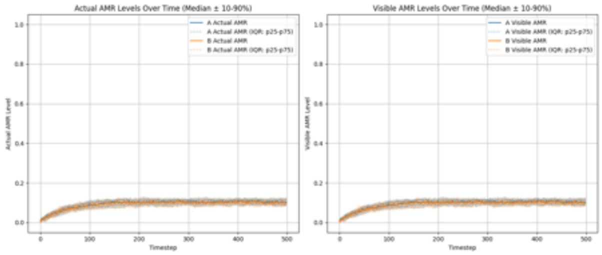

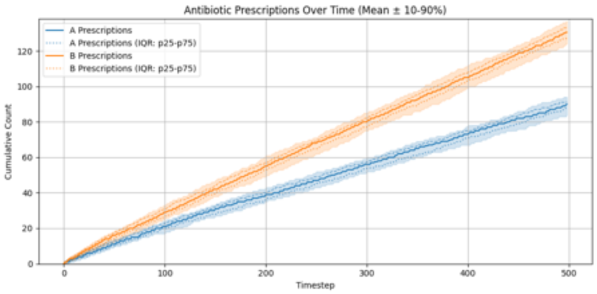

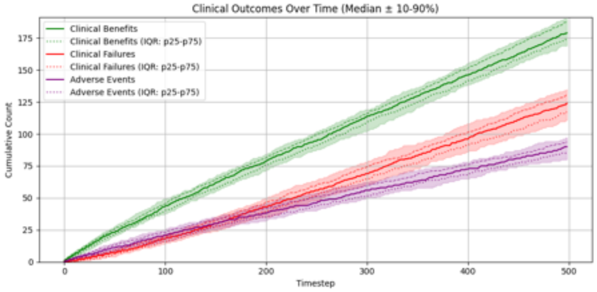

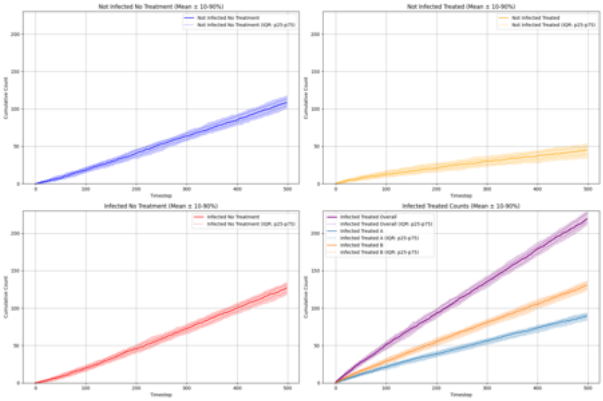

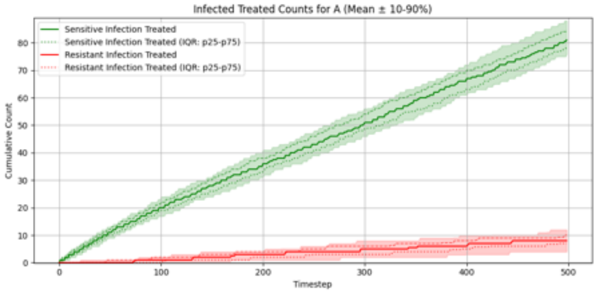

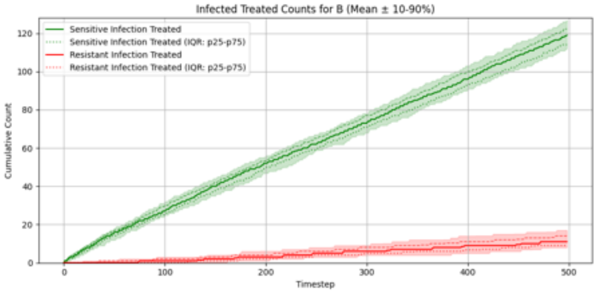

##### Summary Statistics (Two Antibiotics No Cross-Resistance, Accurate Risk Stratification)

| metric | p10 | p25 | p50 | p75 | p90 |
|---|---|---|---|---|---|
| overall__total__reward | -6.047 | 5.574 | 14.773 | 23.348 | 30.404 |
| overall__count__clinical__benefits | 169.000 | 174.000 | 179.000 | 188.000 | 190.100 |
| overall__count__clinical__failures | 110.800 | 117.000 | 124.000 | 131.000 | 135.000 |
| overall__count__adverse__events | 80.000 | 85.000 | 90.000 | 94.000 | 97.100 |
| overall__not__infected__no__treatment__count | 100.800 | 104.000 | 108.000 | 113.250 | 118.000 |
| overall__not__infected__treated__count | 34.000 | 39.750 | 45.000 | 49.000 | 54.000 |
| overall__infected__no__treatment__count | 118.000 | 121.000 | 127.000 | 131.250 | 134.000 |
| overall__infected__treated__count | 211.000 | 214.000 | 219.000 | 227.000 | 229.000 |
| overall__sensitive__infection__treated__count__per__abx__dict__A | 75.900 | 78.000 | 81.000 | 84.250 | 88.000 |
| overall__sensitive__infection__treated__count__per__abx__dict__B | 111.900 | 114.000 | 119.000 | 122.250 | 127.000 |
| overall__resistant__infection__treated__count__per__abx__dict__A | 4.000 | 7.000 | 8.000 | 10.000 | 12.000 |
| overall__resistant__infection__treated__count__per__abx__dict__B | 8.000 | 9.000 | 11.000 | 14.000 | 17.000 |
| overall__sensitive__infection__treated__count | 188.900 | 192.750 | 198.000 | 206.250 | 212.100 |
| overall__resistant__infection__treated__count | 13.900 | 16.000 | 19.000 | 23.000 | 26.300 |
| overall__abx__prescriptions__count__per__abx__A | 83.900 | 88.000 | 90.000 | 91.250 | 94.000 |
| overall__abx__prescriptions__count__per__abx__B | 124.000 | 127.000 | 130.500 | 133.250 | 137.000 |
| overall__abx__prescriptions__count | 211.000 | 214.000 | 219.000 | 227.000 | 229.000 |
| final__amr__actual__A | 0.087 | 0.091 | 0.102 | 0.112 | 0.123 |
| final__amr__visible__A | 0.087 | 0.091 | 0.102 | 0.112 | 0.123 |
| final__amr__actual__B | 0.086 | 0.090 | 0.101 | 0.112 | 0.122 |
| final__amr__visible__B | 0.086 | 0.090 | 0.101 | 0.112 | 0.122 |

#### Two Antibiotics No Cross-Resistance, Exaggerated Risk Stratification

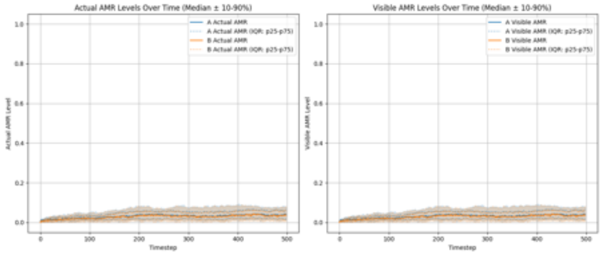

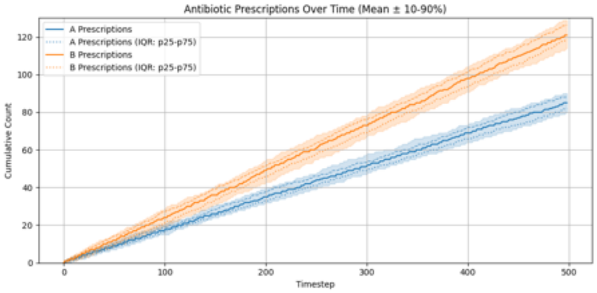

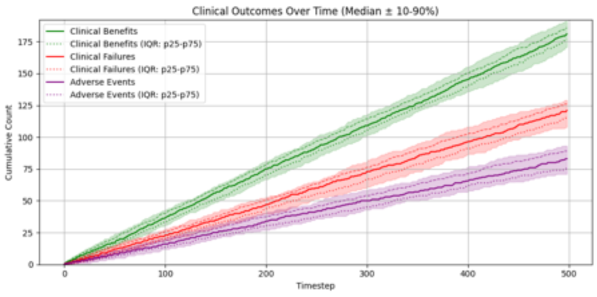

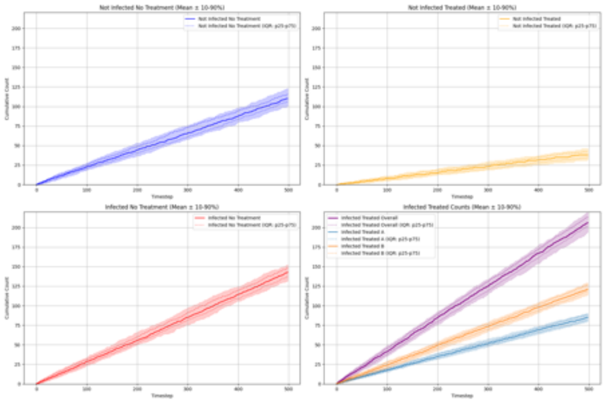

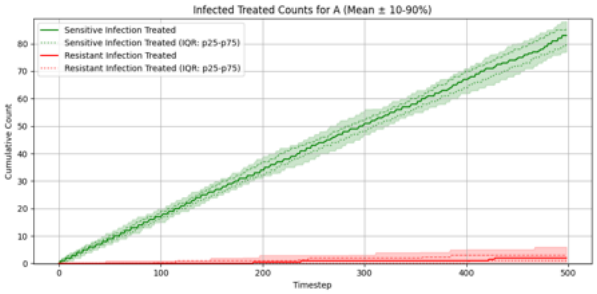

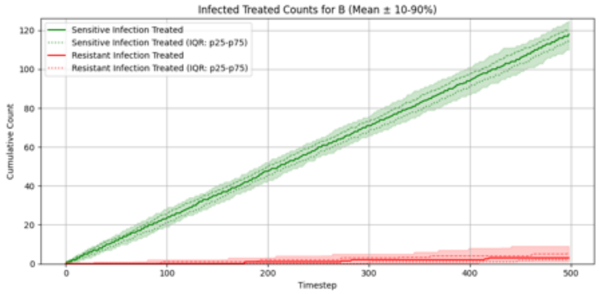

##### Summary Statistics (Two Antibiotics No Cross-Resistance, Exaggerated Risk Stratification)

| metric | p10 | p25 | p50 | p75 | p90 |
|---|---|---|---|---|---|
| overall_total_reward | 15.610 | 22.488 | 33.437 | 41.030 | 48.718 |
| overall_count_clinical_benefits | 170.800 | 176.750 | 181.000 | 186.000 | 191.100 |
| overall_count_clinical_failures | 107.900 | 115.000 | 121.000 | 126.000 | 129.000 |
| overall_count_adverse_events | 71.000 | 75.750 | 83.000 | 89.000 | 93.100 |
| overall_not_infected_no_treatment_count | 100.800 | 106.000 | 110.000 | 116.000 | 123.000 |
| overall_not_infected_treated_count | 31.900 | 34.750 | 38.000 | 42.000 | 46.000 |
| overall_infected_no_treatment_count | 131.900 | 137.000 | 143.000 | 148.000 | 152.100 |
| overall_infected_treated_count | 194.000 | 202.000 | 206.000 | 213.250 | 219.100 |
| overall_sensitive_infection_treated_count_per_abx_dict_A | 77.000 | 80.000 | 83.000 | 85.250 | 88.100 |
| overall_sensitive_infection_treated_count_per_abx_dict_B | 110.000 | 114.750 | 118.000 | 120.250 | 125.100 |
| overall_resistant_infection_treated_count_per_abx_dict_A | 0.000 | 1.000 | 2.000 | 3.000 | 6.000 |
| overall_resistant_infection_treated_count_per_abx_dict_B | 1.000 | 2.000 | 3.000 | 5.250 | 9.000 |
| overall_sensitive_infection_treated_count | 189.000 | 196.000 | 200.000 | 205.250 | 210.000 |
| overall_resistant_infection_treated_count | 2.000 | 3.000 | 5.000 | 9.000 | 14.000 |
| overall_abx_prescriptions_count_per_abx_A | 79.900 | 82.000 | 85.000 | 88.250 | 90.100 |
| overall_abx_prescriptions_count_per_abx_B | 114.000 | 118.750 | 121.000 | 126.250 | 129.100 |
| overall_abx_prescriptions_count | 194.000 | 202.000 | 206.000 | 213.250 | 219.100 |
| final_amr_actual_A | 0.012 | 0.018 | 0.040 | 0.055 | 0.079 |
| final_amr_visible_A | 0.012 | 0.018 | 0.040 | 0.055 | 0.079 |
| final_amr_actual_B | 0.006 | 0.015 | 0.036 | 0.057 | 0.076 |
| final_amr_visible_B | 0.006 | 0.015 | 0.036 | 0.057 | 0.076 |

##### Two Antibiotics No Cross-Resistance, Compressed Risk Stratification

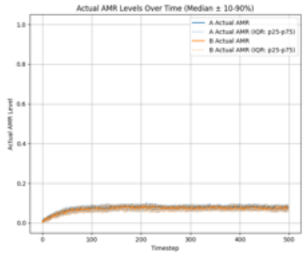

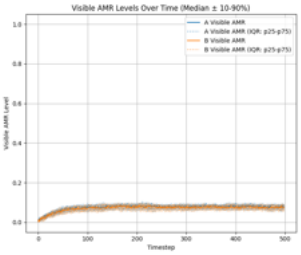

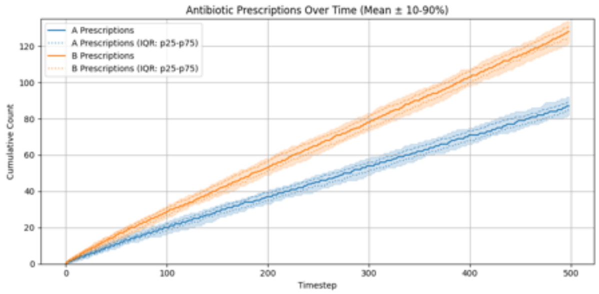

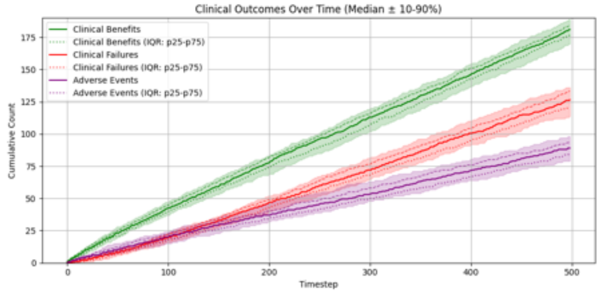

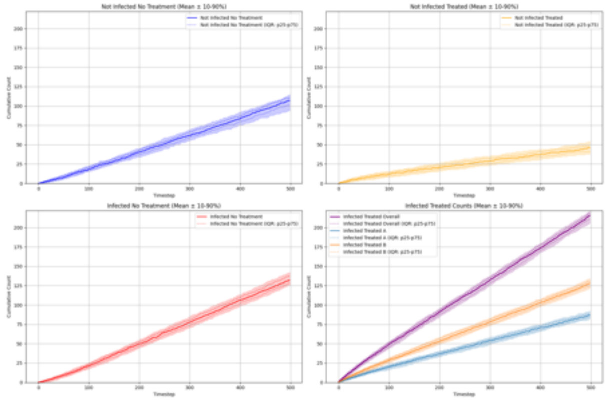

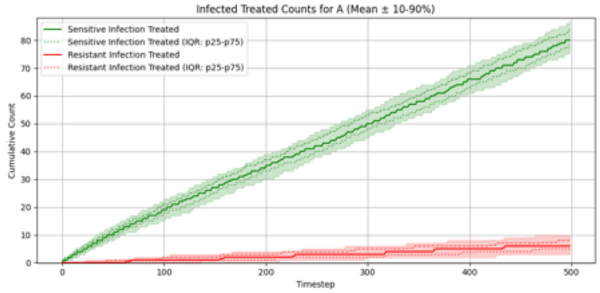

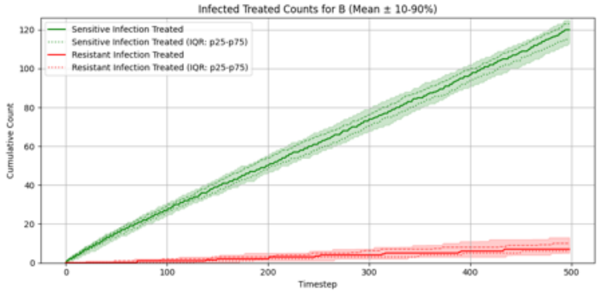

#### Summary Statistics (Two Antibiotics No Cross-Resistance, Compressed Risk Stratification)

| metric | p10 | p25 | p50 | p75 | p90 |
|---|---|---|---|---|---|
| overall_total_reward | -10.275 | 3.024 | 9.925 | 18.101 | 24.764 |
| overall_count_clinical_benefits | 169.900 | 176.750 | 181.000 | 184.000 | 189.000 |
| overall_count_clinical_failures | 113.900 | 120.000 | 126.500 | 133.250 | 136.100 |
| overall_count_adverse_events | 79.000 | 84.000 | 89.000 | 93.250 | 98.100 |
| overall_not_infected_no_treatment_count | 94.000 | 101.500 | 107.000 | 111.000 | 115.000 |
| overall_not_infected_treated_count | 38.000 | 41.000 | 46.000 | 48.250 | 54.000 |
| overall_infected_no_treatment_count | 125.000 | 128.000 | 132.000 | 137.500 | 142.100 |
| overall_infected_treated_count | 205.900 | 212.000 | 215.500 | 219.000 | 221.000 |
| overall_sensitive_infection_treated_count_per_abx_dict_A | 76.000 | 78.000 | 80.000 | 84.000 | 87.000 |
| overall_sensitive_infection_treated_count_per_abx_dict_B | 113.000 | 115.000 | 120.000 | 123.250 | 125.100 |
| overall_resistant_infection_treated_count_per_abx_dict_A | 3.000 | 5.000 | 6.000 | 8.000 | 10.000 |
| overall_resistant_infection_treated_count_per_abx_dict_B | 5.000 | 6.000 | 7.000 | 10.000 | 13.000 |
| overall_sensitive_infection_treated_count | 193.000 | 195.750 | 200.000 | 204.250 | 208.100 |
| overall_resistant_infection_treated_count | 10.000 | 12.000 | 14.000 | 16.250 | 20.100 |
| overall_abx_prescriptions_count_per_abx_A | 82.000 | 84.750 | 87.000 | 89.000 | 92.000 |
| overall_abx_prescriptions_count_per_abx_B | 121.900 | 124.000 | 128.000 | 131.000 | 134.000 |
| overall_abx_prescriptions_count | 205.900 | 212.000 | 215.500 | 219.000 | 221.000 |
| final_amr_actual_A | 0.059 | 0.067 | 0.077 | 0.085 | 0.092 |
| final_amr_visible_A | 0.059 | 0.067 | 0.077 | 0.085 | 0.092 |
| final_amr_actual_B | 0.058 | 0.066 | 0.075 | 0.085 | 0.089 |
| final_amr_visible_B | 0.058 | 0.066 | 0.075 | 0.085 | 0.089 |

##### Two Antibiotics With Cross-Resistance, Accurate Risk Stratification

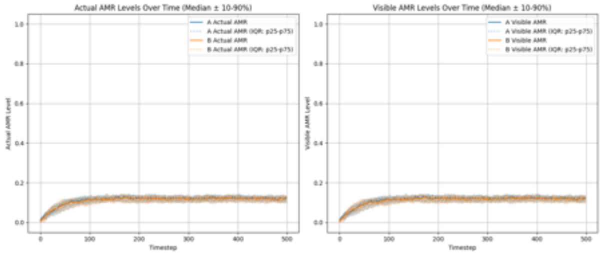

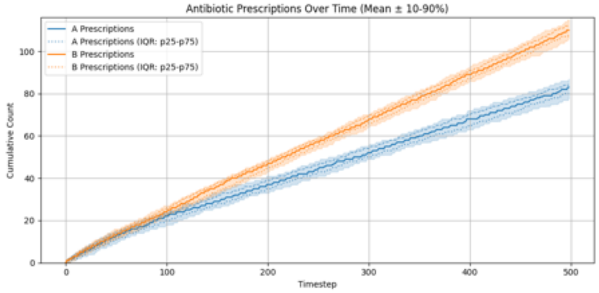

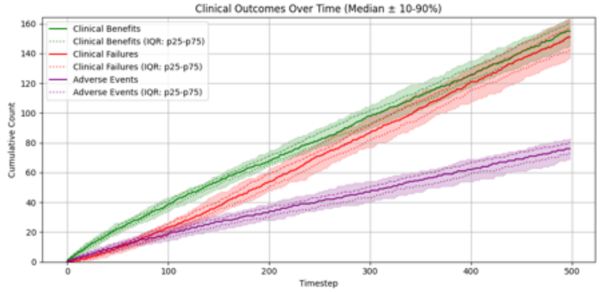

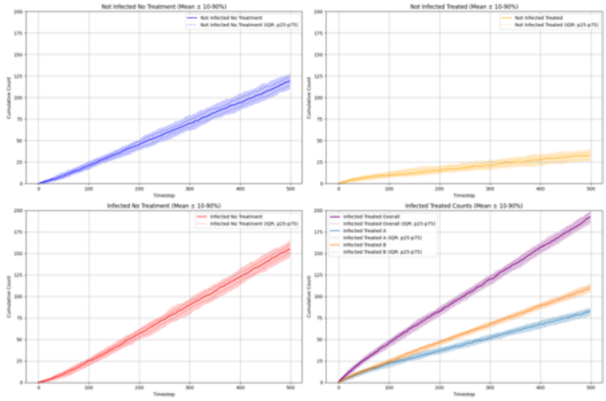

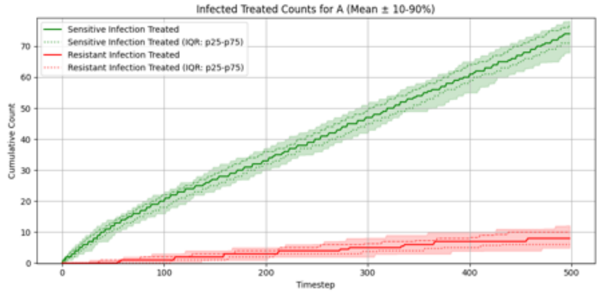

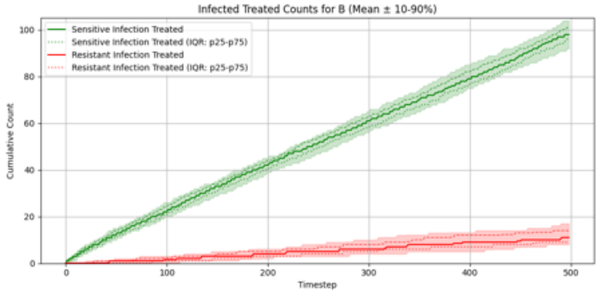

##### Summary Statistics (Two Antibiotics With Cross-Resistance, Accurate Risk Stratification)

| metric | p10 | p25 | p50 | p75 | p90 |
|---|---|---|---|---|---|
| overall__total__reward | -48.300 | -42.049 | -32.900 | -25.449 | -13.250 |
| overall__count__clinical__benefits | 145.000 | 152.000 | 155.000 | 160.250 | 163.200 |
| overall__count__clinical__failures | 136.900 | 142.000 | 151.000 | 157.000 | 163.100 |
| overall__count__adverse__events | 69.000 | 73.000 | 76.000 | 80.000 | 83.000 |
| overall__not__infected__no__treatment__count | 110.800 | 114.000 | 119.000 | 123.250 | 128.000 |
| overall__not__infected__treated__count | 26.900 | 28.750 | 33.000 | 36.000 | 40.100 |
| overall__infected__no__treatment__count | 146.000 | 150.750 | 155.000 | 160.250 | 164.100 |
| overall__infected__treated__count | 184.900 | 188.000 | 192.500 | 197.000 | 199.000 |
| overall__sensitive__infection__treated__count__per__abx__dict__A | 67.900 | 71.750 | 74.000 | 77.000 | 78.100 |
| overall__sensitive__infection__treated__count__per__abx__dict__B | 92.000 | 96.000 | 98.000 | 101.000 | 104.100 |
| overall__resistant__infection__treated__count__per__abx__dict__A | 4.900 | 6.000 | 8.000 | 10.000 | 12.100 |
| overall__resistant__infection__treated__count__per__abx__dict__B | 8.000 | 9.000 | 11.000 | 14.000 | 17.000 |
| overall__sensitive__infection__treated__count | 165.900 | 167.750 | 172.000 | 177.250 | 180.100 |
| overall__resistant__infection__treated__count | 14.900 | 16.000 | 19.000 | 23.000 | 26.000 |
| overall__abx__prescriptions__count__per__abx__A | 77.800 | 80.000 | 83.000 | 84.250 | 87.000 |
| overall__abx__prescriptions__count__per__abx__B | 105.900 | 108.000 | 110.000 | 112.250 | 115.000 |
| overall__abx__prescriptions__count | 184.900 | 188.000 | 192.500 | 197.000 | 199.000 |
| final__amr__actual__A | 0.105 | 0.111 | 0.120 | 0.129 | 0.135 |
| final__amr__visible__A | 0.105 | 0.111 | 0.120 | 0.129 | 0.135 |
| final__amr__actual__B | 0.102 | 0.111 | 0.118 | 0.128 | 0.136 |
| final__amr__visible__B | 0.102 | 0.111 | 0.118 | 0.128 | 0.136 |

##### Two Antibiotics With Cross-Resistance, Exaggerated Risk Stratification

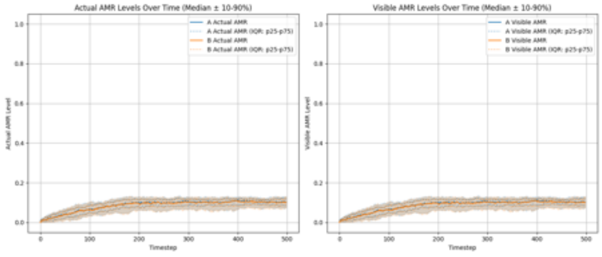

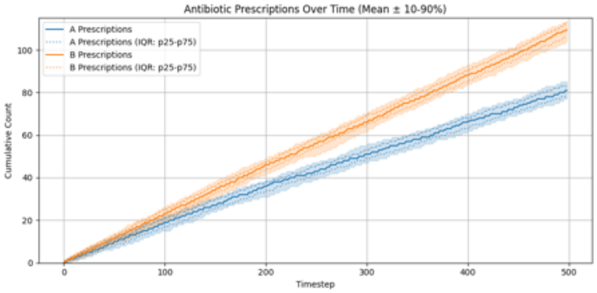

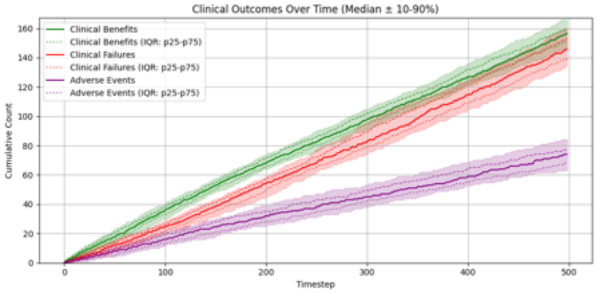

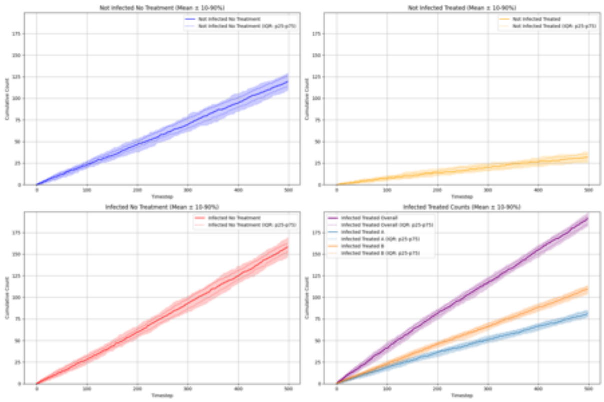

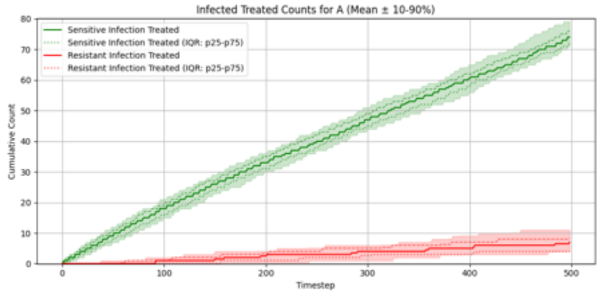

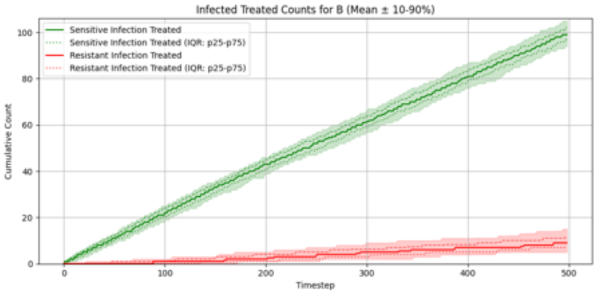

##### Summary Statistics (Two Antibiotics With Cross-Resistance, Exaggerated Risk Stratification)

| metric | p10 | p25 | p50 | p75 | p90 |
|---|---|---|---|---|---|
| overall_total_reward | -38.200 | -31.423 | -19.549 | -7.823 | 3.333 |
| overall_count_clinical_benefits | 147.900 | 151.750 | 156.500 | 160.250 | 166.100 |
| overall_count_clinical_failures | 134.800 | 139.750 | 146.000 | 153.000 | 160.100 |
| overall_count_adverse_events | 62.900 | 68.000 | 74.000 | 77.000 | 84.100 |
| overall_not_infected_no_treatment_count | 109.800 | 114.000 | 119.500 | 126.000 | 129.000 |
| overall_not_infected_treated_count | 25.000 | 28.000 | 32.000 | 35.000 | 38.000 |
| overall_infected_no_treatment_count | 147.900 | 152.750 | 158.000 | 162.250 | 169.000 |
| overall_infected_treated_count | 183.000 | 185.750 | 191.000 | 193.250 | 198.000 |
| overall_sensitive_infection_treated_count_per_abx_dict_A | 71.000 | 71.750 | 74.000 | 76.000 | 79.100 |
| overall_sensitive_infection_treated_count_per_abx_dict_B | 94.000 | 97.000 | 99.000 | 102.000 | 105.000 |
| overall_resistant_infection_treated_count_per_abx_dict_A | 4.000 | 4.000 | 7.000 | 8.250 | 11.000 |
| overall_resistant_infection_treated_count_per_abx_dict_B | 5.000 | 7.000 | 9.000 | 11.250 | 15.000 |
| overall_sensitive_infection_treated_count | 164.000 | 168.750 | 173.000 | 178.000 | 181.100 |
| overall_resistant_infection_treated_count | 9.900 | 13.000 | 16.000 | 20.000 | 23.000 |
| overall_abx_prescriptions_count_per_abx_A | 77.000 | 78.000 | 81.000 | 83.000 | 85.100 |
| overall_abx_prescriptions_count_per_abx_B | 104.000 | 106.000 | 109.500 | 113.000 | 114.000 |
| overall_abx_prescriptions_count | 183.000 | 185.750 | 191.000 | 193.250 | 198.000 |
| final_amr_actual_A | 0.076 | 0.085 | 0.106 | 0.122 | 0.131 |
| final_amr_visible_A | 0.076 | 0.085 | 0.106 | 0.122 | 0.131 |
| final_amr_actual_B | 0.077 | 0.085 | 0.101 | 0.117 | 0.128 |
| final_amr_visible_B | 0.077 | 0.085 | 0.101 | 0.117 | 0.128 |

##### Two Antibiotics With Cross-Resistance, Compressed Risk Stratification

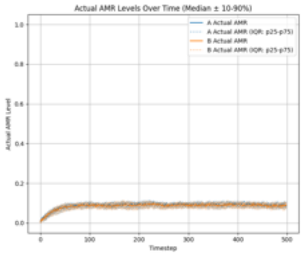

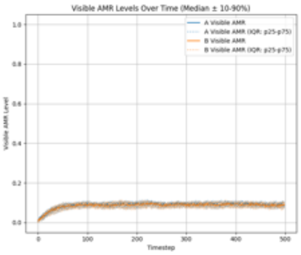

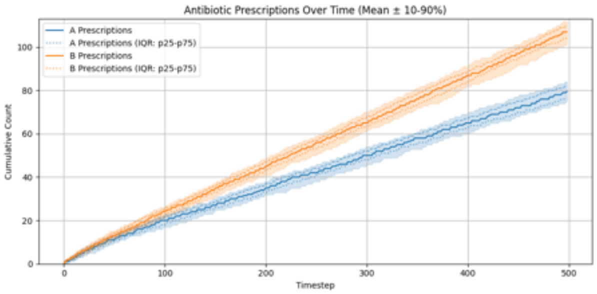

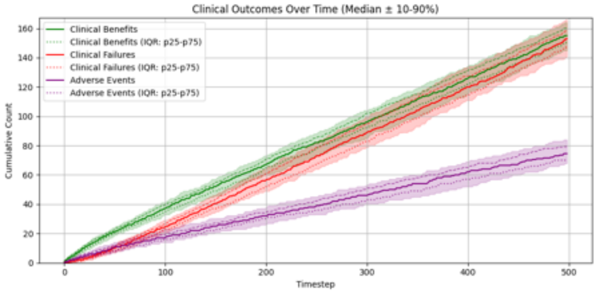

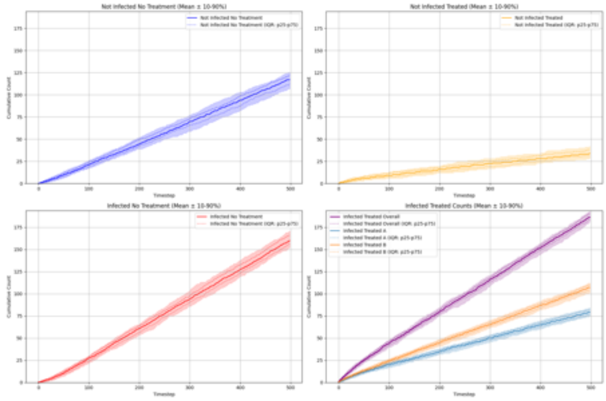

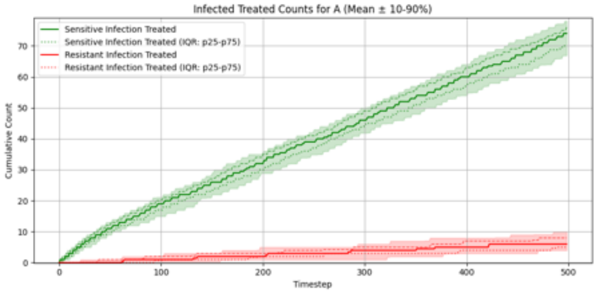

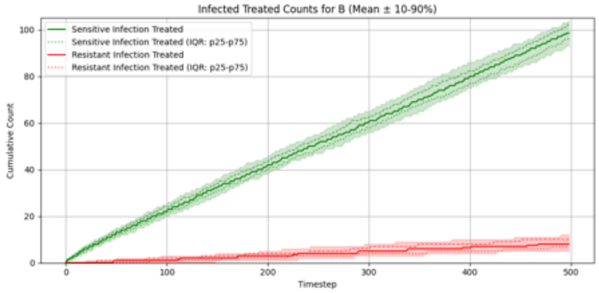

#### Summary Statistics (Two Antibiotics With Cross-Resistance, Compressed Risk Stratification)

| metric | p10 | p25 | p50 | p75 | p90 |
|---|---|---|---|---|---|
| overall_total_reward | -60.379 | -48.073 | -37.998 | -28.647 | -21.271 |
| overall_count_clinical_benefits | 144.900 | 151.000 | 155.000 | 160.000 | 164.000 |
| overall_count_clinical_failures | 140.900 | 147.000 | 153.000 | 159.000 | 166.100 |
| overall_count_adverse_events | 67.900 | 70.000 | 74.500 | 79.250 | 84.000 |
| overall_not_infected_no_treatment_count | 106.800 | 112.500 | 117.000 | 122.000 | 125.100 |
| overall_not_infected_treated_count | 28.900 | 31.000 | 34.000 | 38.000 | 42.000 |
| overall_infected_no_treatment_count | 152.000 | 156.000 | 160.000 | 166.000 | 171.000 |
| overall_infected_treated_count | 181.800 | 184.000 | 186.500 | 190.250 | 193.000 |
| overall_sensitive_infection_treated_count_per_abx_dict_A | 66.900 | 70.000 | 74.000 | 76.000 | 78.000 |
| overall_sensitive_infection_treated_count_per_abx_dict_B | 93.000 | 96.000 | 98.500 | 102.000 | 104.000 |
| overall_resistant_infection_treated_count_per_abx_dict_A | 4.000 | 5.000 | 6.000 | 8.000 | 10.000 |
| overall_resistant_infection_treated_count_per_abx_dict_B | 5.000 | 6.750 | 8.000 | 10.000 | 12.100 |
| overall_sensitive_infection_treated_count | 163.900 | 168.000 | 172.000 | 177.000 | 179.000 |
| overall_resistant_infection_treated_count | 11.000 | 12.000 | 14.500 | 17.000 | 19.100 |
| overall_abx_prescriptions_count_per_abx_A | 74.900 | 76.000 | 79.500 | 82.000 | 84.100 |
| overall_abx_prescriptions_count_per_abx_B | 101.000 | 104.000 | 107.000 | 110.000 | 112.000 |
| overall_abx_prescriptions_count | 181.800 | 184.000 | 186.500 | 190.250 | 193.000 |
| final_amr_actual_A | 0.074 | 0.081 | 0.090 | 0.100 | 0.105 |
| final_amr_visible_A | 0.074 | 0.081 | 0.090 | 0.100 | 0.105 |
| final_amr_actual_B | 0.069 | 0.076 | 0.088 | 0.098 | 0.104 |
| final_amr_visible_B | 0.069 | 0.076 | 0.088 | 0.098 | 0.104 |

### Experiment Set 4

#### Single Antibiotic, Hierarchical PPO

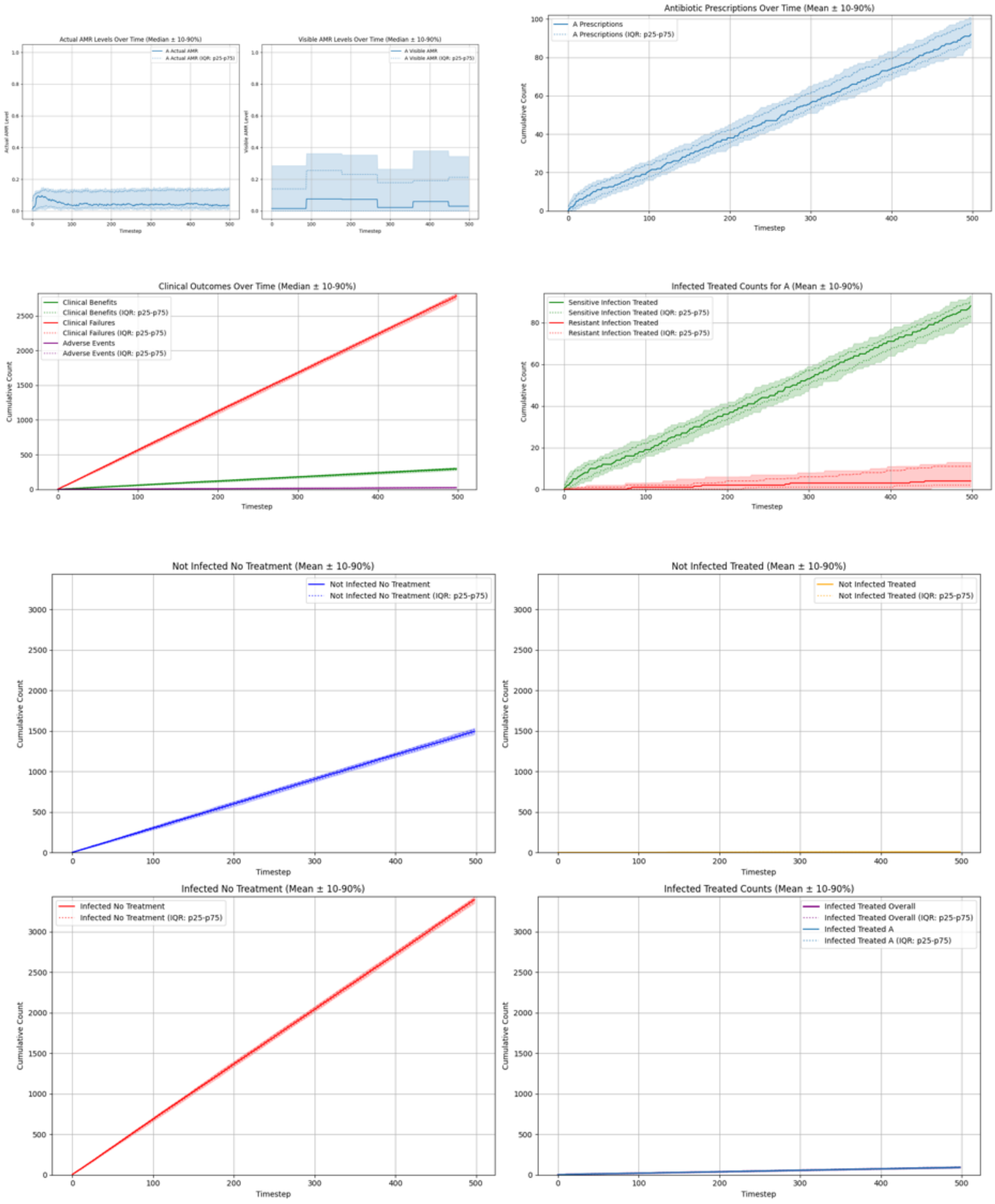

##### Summary Statistics (Single Antibiotic, Hierarchical PPO)

| metric | p10 | p25 | p50 | p75 | p90 |
|---|---|---|---|---|---|
| overall_total_reward | -272.970 | -271.459 | -268.621 | -266.258 | -263.439 |
| overall_count_clinical_benefits | 275.000 | 284.250 | 296.000 | 305.000 | 317.100 |
| overall_count_clinical_failures | 2739.900 | 2764.500 | 2784.500 | 2803.250 | 2822.300 |
| overall_count_adverse_events | 19.900 | 22.000 | 25.000 | 29.250 | 31.100 |
| overall_not_infected_no_treatment_count | 1459.800 | 1481.250 | 1496.500 | 1522.000 | 1535.500 |
| overall_not_infected_treated_count | 4.000 | 5.750 | 8.000 | 10.250 | 13.000 |
| overall_infected_no_treatment_count | 3348.800 | 3370.250 | 3394.000 | 3412.000 | 3433.100 |
| overall_infected_treated_count | 84.900 | 88.000 | 92.000 | 98.000 | 101.000 |
| overall_sensitive_infection_treated_count_per_abx_dict_A | 80.000 | 83.000 | 88.000 | 90.000 | 93.000 |
| overall_resistant_infection_treated_count_per_abx_dict_A | 1.000 | 2.000 | 4.000 | 11.000 | 13.000 |
| overall_sensitive_infection_treated_count | 80.000 | 83.000 | 88.000 | 90.000 | 93.000 |
| overall_resistant_infection_treated_count | 1.000 | 2.000 | 4.000 | 11.000 | 13.000 |
| overall_abx_prescriptions_count_per_abx_A | 84.900 | 88.000 | 92.000 | 98.000 | 101.000 |
| overall_abx_prescriptions_count | 84.900 | 88.000 | 92.000 | 98.000 | 101.000 |
| final_amr_actual_A | 0.011 | 0.021 | 0.033 | 0.135 | 0.144 |
| final_amr_visible_A | 0.000 | 0.000 | 0.030 | 0.212 | 0.345 |

##### Single Antibiotic, Hierarchical Recurrent PPO

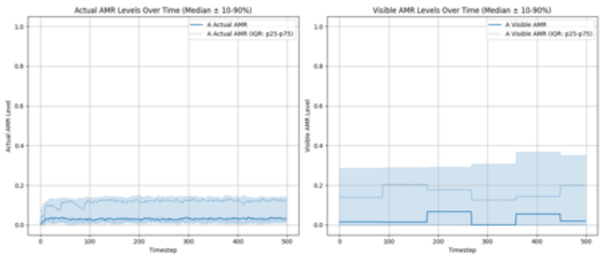

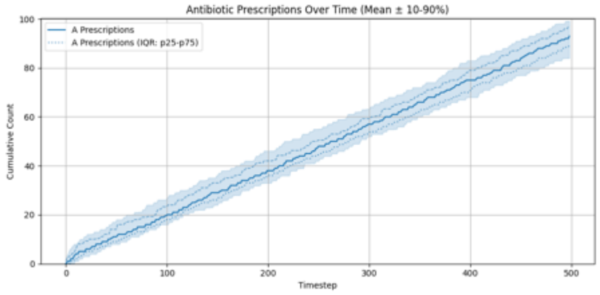

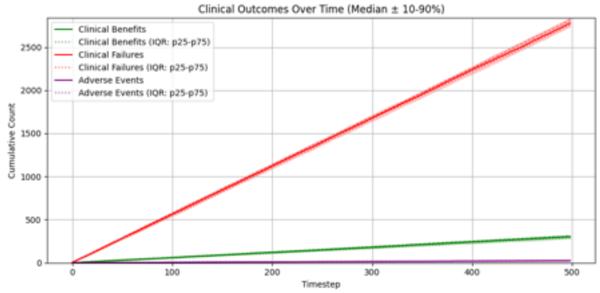

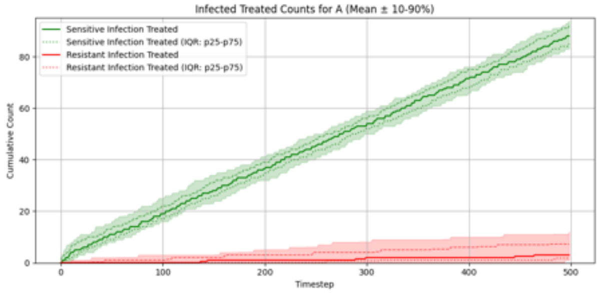

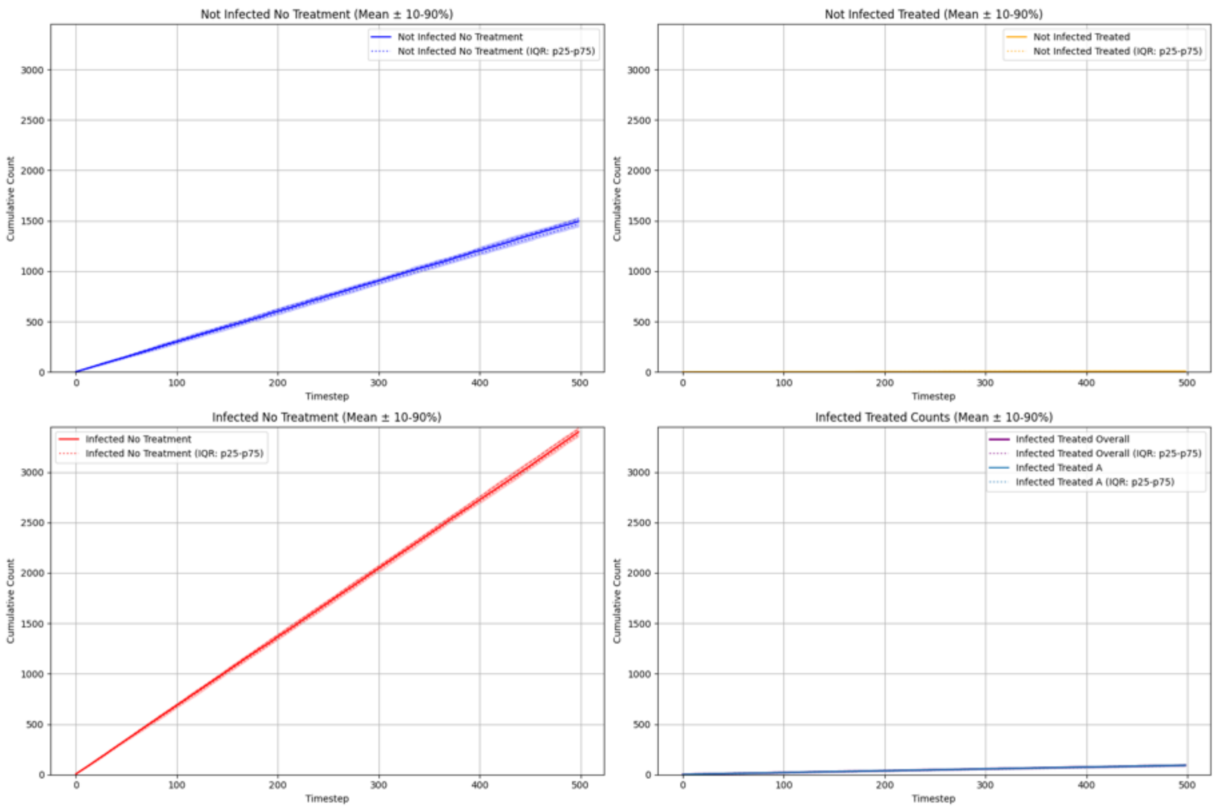

#### Summary Statistics (Single Antibiotic, Hierarchical Recurrent PPO)

| metric | p10 | p25 | p50 | p75 | p90 |
|---|---|---|---|---|---|
| overall_total_reward | -276.008 | -270.600 | -267.444 | -264.670 | -262.630 |
| overall_count_clinical_benefits | 282.100 | 289.000 | 302.500 | 314.000 | 320.100 |
| overall_count_clinical_failures | 2737.800 | 2758.000 | 2779.500 | 2815.750 | 2837.200 |
| overall_count_adverse_events | 18.900 | 21.000 | 25.000 | 28.000 | 32.100 |
| overall_not_infected_no_treatment_count | 1443.000 | 1466.750 | 1495.500 | 1516.250 | 1536.200 |
| overall_not_infected_treated_count | 3.000 | 5.000 | 7.000 | 10.000 | 12.000 |
| overall_infected_no_treatment_count | 3354.800 | 3369.750 | 3398.500 | 3425.250 | 3449.100 |
| overall_infected_treated_count | 84.000 | 89.000 | 93.000 | 97.000 | 99.200 |
| overall_sensitive_infection_treated_count_per_abx_dict_A | 83.000 | 85.000 | 88.000 | 92.000 | 94.000 |
| overall_resistant_infection_treated_count_per_abx_dict_A | 1.000 | 2.000 | 3.000 | 7.250 | 12.000 |
| overall_sensitive_infection_treated_count | 83.000 | 85.000 | 88.000 | 92.000 | 94.000 |
| overall_resistant_infection_treated_count | 1.000 | 2.000 | 3.000 | 7.250 | 12.000 |
| overall_abx_prescriptions_count_per_abx_A | 84.000 | 89.000 | 93.000 | 97.000 | 99.200 |
| overall_abx_prescriptions_count | 84.000 | 89.000 | 93.000 | 97.000 | 99.200 |
| final_amr_actual_A | 0.011 | 0.022 | 0.029 | 0.125 | 0.138 |
| final_amr_visible_A | 0.000 | 0.000 | 0.019 | 0.199 | 0.351 |

##### Two Antibiotics No Cross-Resistance, Hierarchical PPO

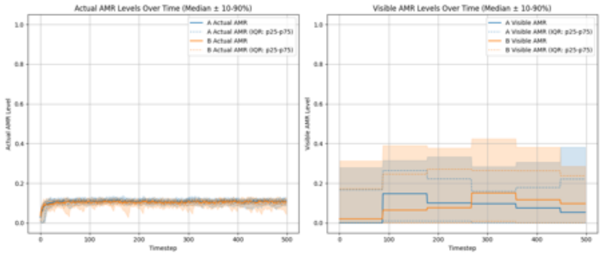

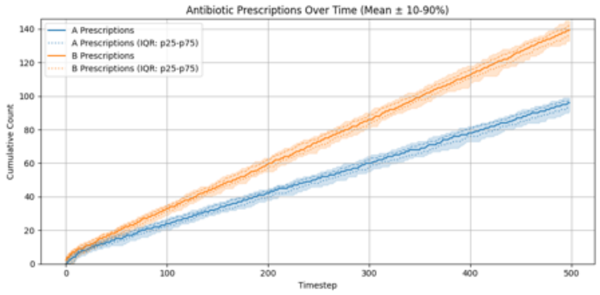

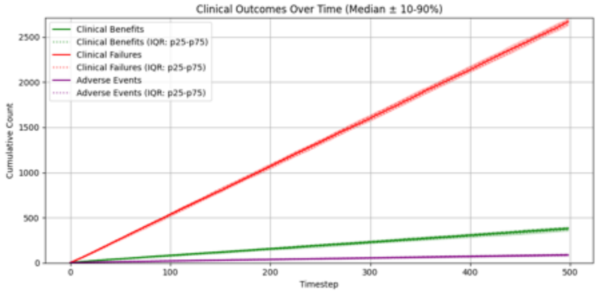

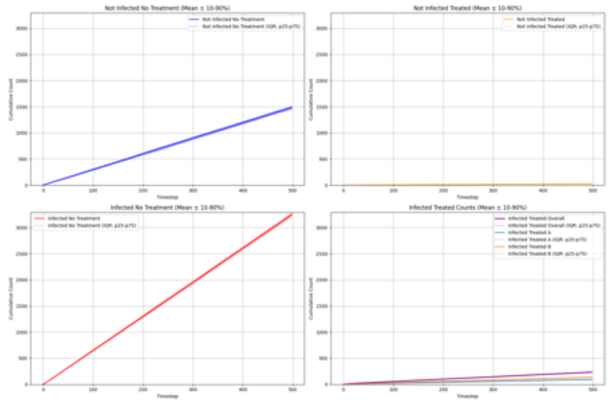

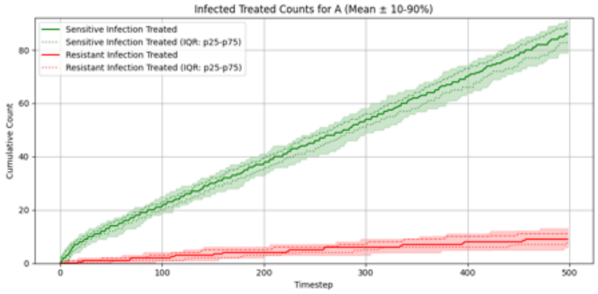

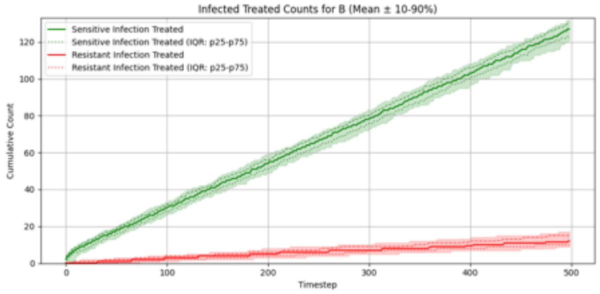

##### Summary Statistics (Two Antibiotics No Cross-Resistance, Hierarchical PPO)

| metric | p10 | p25 | p50 | p75 | p90 |
|---|---|---|---|---|---|
| overall_total_reward | -254.854 | -251.545 | -249.370 | -246.534 | -243.885 |
| overall_count_clinical_benefits | 353.700 | 366.750 | 379.000 | 391.250 | 398.000 |
| overall_count_clinical_failures | 2629.500 | 2644.000 | 2668.500 | 2685.250 | 2709.600 |
| overall_count_adverse_events | 77.000 | 81.750 | 87.500 | 94.250 | 99.000 |
| overall_not_infected_no_treatment_count | 1449.000 | 1470.750 | 1483.000 | 1503.000 | 1515.200 |
| overall_not_infected_treated_count | 19.000 | 22.000 | 26.500 | 29.000 | 31.100 |
| overall_infected_no_treatment_count | 3214.800 | 3225.000 | 3248.500 | 3260.500 | 3291.400 |
| overall_infected_treated_count | 227.700 | 232.000 | 234.000 | 237.250 | 242.000 |
| overall_sensitive_infection_treated_count_per_abx_dict_A | 79.000 | 82.750 | 86.000 | 89.000 | 91.000 |
| overall_sensitive_infection_treated_count_per_abx_dict_B | 120.900 | 123.000 | 127.000 | 130.000 | 132.000 |
| overall_resistant_infection_treated_count_per_abx_dict_A | 5.900 | 7.750 | 9.000 | 11.250 | 13.000 |
| overall_resistant_infection_treated_count_per_abx_dict_B | 8.900 | 10.000 | 12.000 | 15.000 | 17.000 |
| overall_sensitive_infection_treated_count | 203.900 | 208.000 | 213.500 | 216.000 | 220.100 |
| overall_resistant_infection_treated_count | 16.000 | 18.750 | 22.000 | 26.000 | 27.100 |
| overall_abx_prescriptions_count_per_abx_A | 90.900 | 93.000 | 96.000 | 98.000 | 99.000 |
| overall_abx_prescriptions_count_per_abx_B | 132.900 | 136.000 | 139.500 | 142.000 | 145.000 |
| overall_abx_prescriptions_count | 227.700 | 232.000 | 234.000 | 237.250 | 242.000 |
| final_amr_actual_A | 0.086 | 0.098 | 0.110 | 0.118 | 0.122 |
| final_amr_visible_A | 0.000 | 0.000 | 0.053 | 0.221 | 0.382 |
| final_amr_actual_B | 0.062 | 0.097 | 0.107 | 0.114 | 0.119 |
| final_amr_visible_B | 0.000 | 0.000 | 0.098 | 0.237 | 0.285 |

#### Two Antibiotics No Cross-Resistance, Hierarchical Recurrent PPO

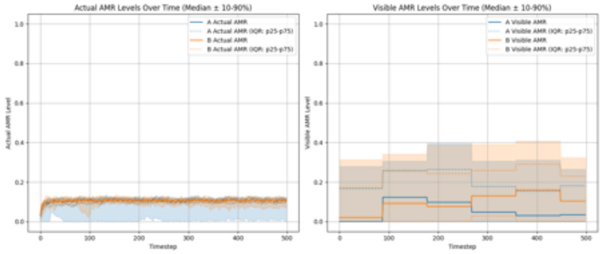

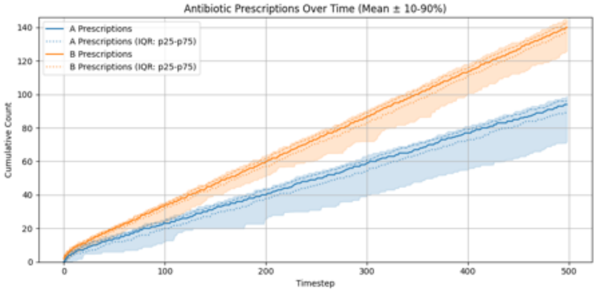

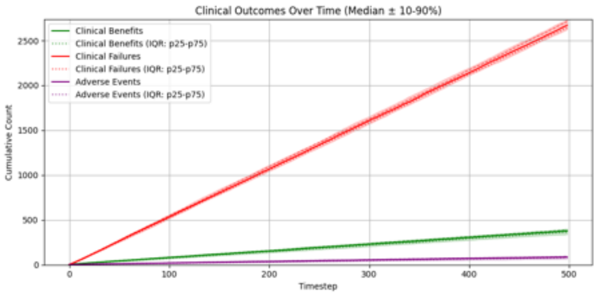

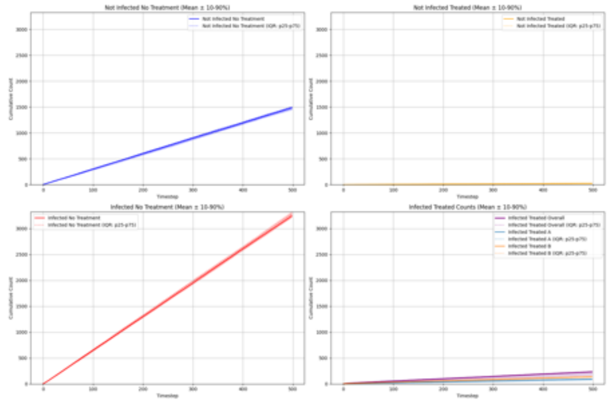

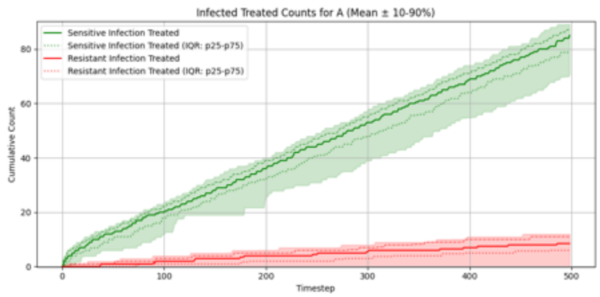

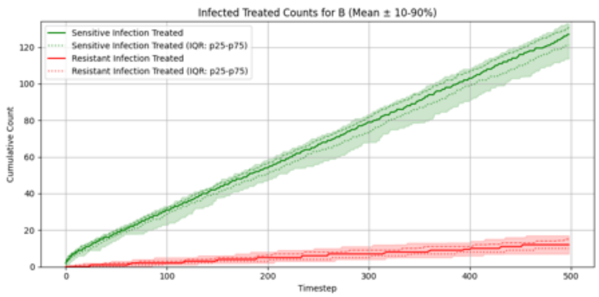

##### Summary Statistics (Two Antibiotics No Cross-Resistance, Hierarchical Recurrent PPO)

| metric | p10 | p25 | p50 | p75 | p90 |
|---|---|---|---|---|---|
| overall_total_reward | -260.594 | -254.915 | -249.771 | -246.572 | -242.876 |
| overall_count_clinical_benefits | 339.000 | 364.750 | 376.000 | 388.250 | 400.100 |
| overall_count_clinical_failures | 2623.800 | 2640.500 | 2670.500 | 2710.250 | 2733.500 |
| overall_count_adverse_events | 66.900 | 79.000 | 86.500 | 92.000 | 97.100 |
| overall_not_infected_no_treatment_count | 1446.800 | 1470.750 | 1486.000 | 1503.250 | 1511.300 |
| overall_not_infected_treated_count | 15.000 | 19.750 | 25.500 | 29.000 | 32.000 |
| overall_infected_no_treatment_count | 3219.000 | 3225.750 | 3249.000 | 3283.500 | 3324.500 |
| overall_infected_treated_count | 168.700 | 225.750 | 234.000 | 238.000 | 242.000 |
| overall_sensitive_infection_treated_count_per_abx_dict_A | 70.800 | 79.000 | 85.000 | 87.000 | 89.100 |
| overall_sensitive_infection_treated_count_per_abx_dict_B | 113.600 | 121.000 | 127.000 | 131.000 | 133.000 |
| overall_resistant_infection_treated_count_per_abx_dict_A | 0.000 | 6.000 | 8.500 | 11.000 | 12.000 |
| overall_resistant_infection_treated_count_per_abx_dict_B | 7.000 | 10.000 | 12.000 | 15.000 | 17.000 |
| overall_sensitive_infection_treated_count | 160.800 | 204.000 | 211.000 | 216.250 | 220.000 |
| overall_resistant_infection_treated_count | 12.500 | 18.750 | 22.000 | 25.000 | 26.100 |
| overall_abx_prescriptions_count_per_abx_A | 71.900 | 89.000 | 94.000 | 96.250 | 99.000 |
| overall_abx_prescriptions_count_per_abx_B | 125.800 | 136.750 | 140.000 | 143.000 | 145.000 |
| overall_abx_prescriptions_count | 168.700 | 225.750 | 234.000 | 238.000 | 242.000 |
| final_amr_actual_A | 0.007 | 0.099 | 0.109 | 0.116 | 0.122 |
| final_amr_visible_A | 0.000 | 0.000 | 0.034 | 0.181 | 0.265 |
| final_amr_actual_B | 0.077 | 0.100 | 0.109 | 0.115 | 0.119 |
| final_amr_visible_B | 0.000 | 0.000 | 0.104 | 0.230 | 0.324 |

##### Two Antibiotics With Cross-Resistance, Hierarchical PPO

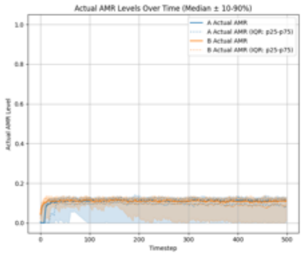

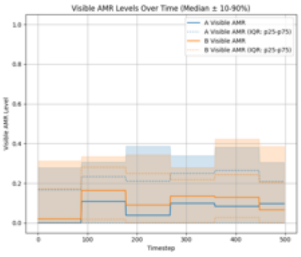

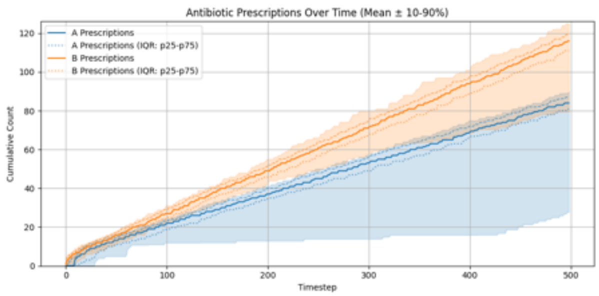

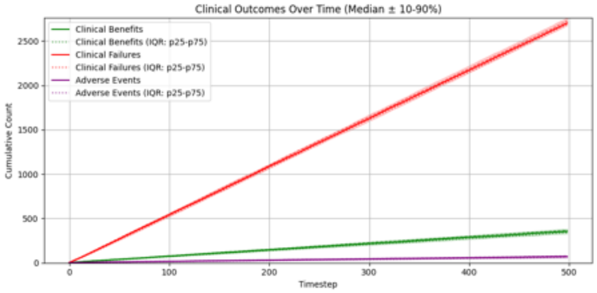

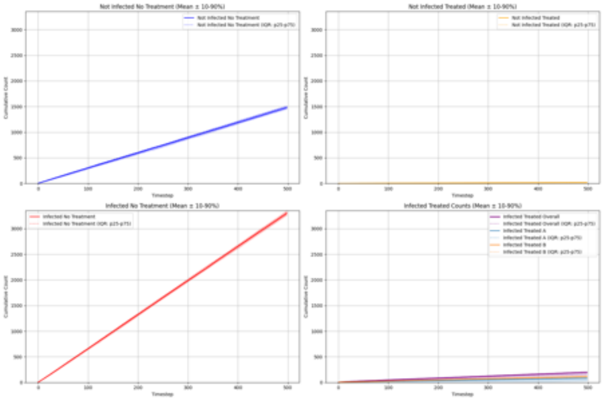

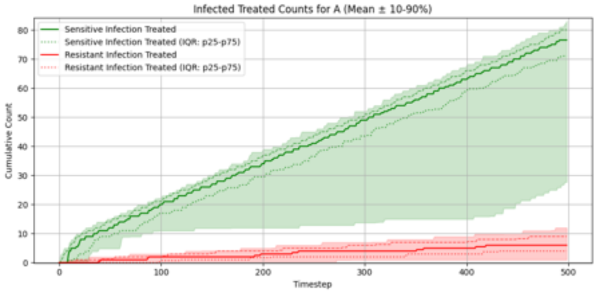

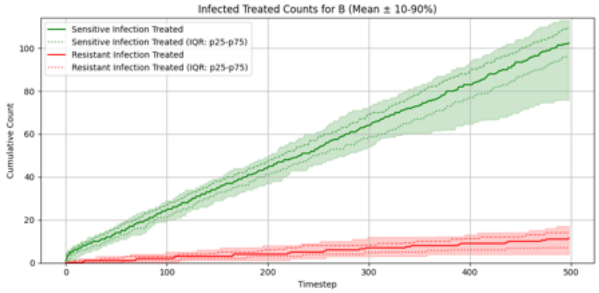

#### Summary Statistics (Two Antibiotics With Cross-Resistance, Hierarchical PPO)

| metric | p10 | p25 | p50 | p75 | p90 |
|---|---|---|---|---|---|
| overall_total_reward | -262.979 | -259.141 | -255.465 | -252.847 | -250.688 |
| overall_count_clinical_benefits | 327.900 | 341.500 | 354.500 | 366.000 | 381.100 |
| overall_count_clinical_failures | 2674.900 | 2688.000 | 2703.000 | 2726.500 | 2759.500 |
| overall_count_adverse_events | 55.000 | 64.500 | 71.500 | 77.000 | 82.000 |
| overall_not_infected_no_treatment_count | 1450.600 | 1467.750 | 1480.500 | 1504.000 | 1515.000 |
| overall_not_infected_treated_count | 12.900 | 15.000 | 18.500 | 21.250 | 23.000 |
| overall_infected_no_treatment_count | 3258.800 | 3281.000 | 3298.500 | 3317.500 | 3352.100 |
| overall_infected_treated_count | 145.900 | 164.000 | 199.000 | 204.000 | 206.000 |
| overall_sensitive_infection_treated_count_per_abx_dict_A | 27.700 | 71.000 | 76.500 | 80.000 | 83.100 |
| overall_sensitive_infection_treated_count_per_abx_dict_B | 75.900 | 96.000 | 102.500 | 110.000 | 113.100 |
| overall_resistant_infection_treated_count_per_abx_dict_A | 1.000 | 4.000 | 6.000 | 9.000 | 12.000 |
| overall_resistant_infection_treated_count_per_abx_dict_B | 3.900 | 7.000 | 11.500 | 14.000 | 17.000 |
| overall_sensitive_infection_treated_count | 136.900 | 152.000 | 176.500 | 184.250 | 188.100 |
| overall_resistant_infection_treated_count | 8.900 | 12.000 | 17.000 | 23.000 | 25.300 |
| overall_abx_prescriptions_count_per_abx_A | 27.700 | 81.000 | 84.000 | 87.000 | 90.000 |
| overall_abx_prescriptions_count_per_abx_B | 79.500 | 111.000 | 116.000 | 119.250 | 125.100 |
| overall_abx_prescriptions_count | 145.900 | 164.000 | 199.000 | 204.000 | 206.000 |
| final_amr_actual_A | 0.004 | 0.085 | 0.114 | 0.121 | 0.131 |
| final_amr_visible_A | 0.000 | 0.000 | 0.097 | 0.209 | 0.305 |
| final_amr_actual_B | 0.009 | 0.085 | 0.112 | 0.121 | 0.126 |
| final_amr_visible_B | 0.000 | 0.000 | 0.066 | 0.204 | 0.386 |

#### Two Antibiotics With Cross-Resistance, Hierarchical Recurrent PPO

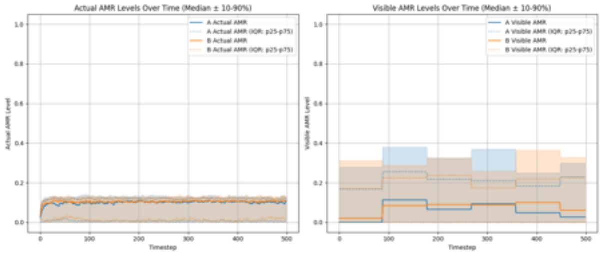

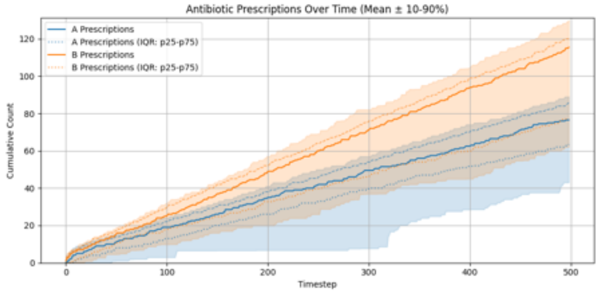

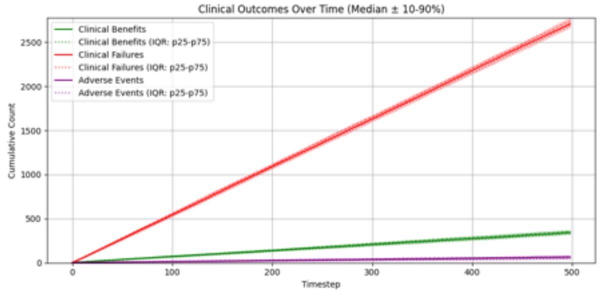

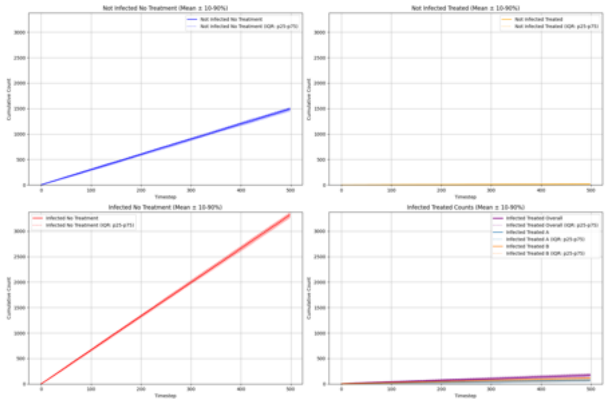

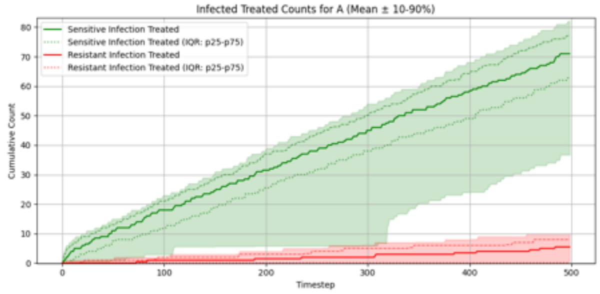

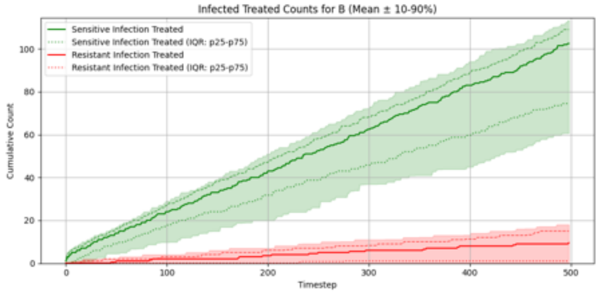

#### Summary Statistics (Two Antibiotics With Cross-Resistance, Hierarchical Recurrent PPO)

| metric | p10 | p25 | p50 | p75 | p90 |
|---|---|---|---|---|---|
| overall_total_reward | -264.887 | -260.714 | -256.961 | -253.896 | -252.106 |
| overall_count_clinical_benefits | 323.700 | 329.750 | 343.500 | 354.750 | 369.100 |
| overall_count_clinical_failures | 2668.000 | 2692.000 | 2710.500 | 2741.250 | 2773.600 |
| overall_count_adverse_events | 43.000 | 50.000 | 66.000 | 73.250 | 78.200 |
| overall_not_infected_no_treatment_count | 1450.600 | 1475.750 | 1489.500 | 1505.250 | 1525.100 |
| overall_not_infected_treated_count | 9.900 | 13.750 | 17.000 | 20.250 | 22.300 |
| overall_infected_no_treatment_count | 3264.700 | 3279.000 | 3314.000 | 3336.500 | 3373.800 |
| overall_infected_treated_count | 135.900 | 142.750 | 172.500 | 200.250 | 205.100 |
| overall_sensitive_infection_treated_count_per_abx_dict_A | 36.700 | 62.750 | 71.000 | 77.250 | 82.100 |
| overall_sensitive_infection_treated_count_per_abx_dict_B | 60.700 | 75.000 | 102.500 | 109.000 | 113.400 |
| overall_resistant_infection_treated_count_per_abx_dict_A | 0.000 | 0.000 | 5.500 | 8.000 | 10.000 |
| overall_resistant_infection_treated_count_per_abx_dict_B | 0.000 | 1.000 | 9.500 | 15.000 | 18.000 |
| overall_sensitive_infection_treated_count | 123.900 | 134.000 | 157.500 | 176.750 | 187.000 |
| overall_resistant_infection_treated_count | 0.000 | 6.500 | 16.000 | 19.250 | 24.000 |
| overall_abx_prescriptions_count_per_abx_A | 43.000 | 63.500 | 76.500 | 86.000 | 89.100 |
| overall_abx_prescriptions_count_per_abx_B | 62.000 | 77.000 | 115.500 | 120.000 | 130.200 |
| overall_abx_prescriptions_count | 135.900 | 142.750 | 172.500 | 200.250 | 205.100 |
| final_amr_actual_A | 0.000 | 0.005 | 0.099 | 0.120 | 0.129 |
| final_amr_visible_A | 0.000 | 0.000 | 0.026 | 0.229 | 0.299 |
| final_amr_actual_B | 0.002 | 0.020 | 0.109 | 0.123 | 0.133 |
| final_amr_visible_B | 0.000 | 0.000 | 0.060 | 0.225 | 0.328 |

### Fixed Prescribing Rules

#### Experiment Set 1

##### Single Antibiotic, Expected Reward Greedy

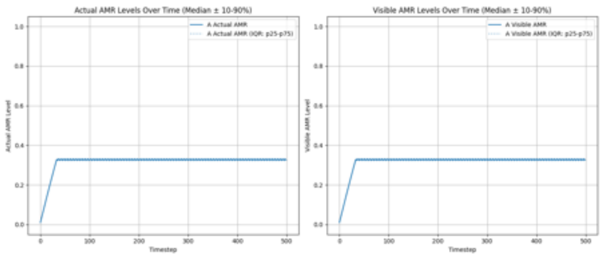

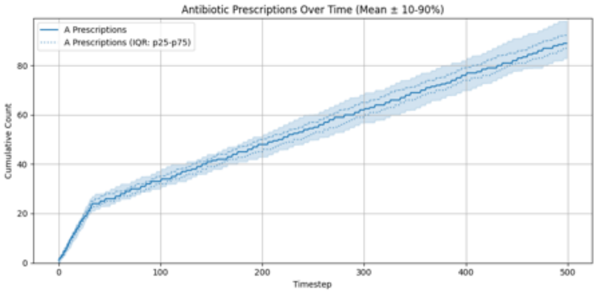

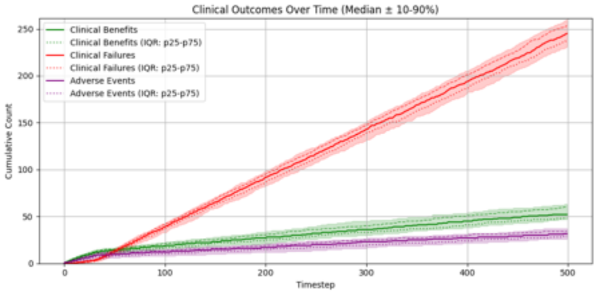

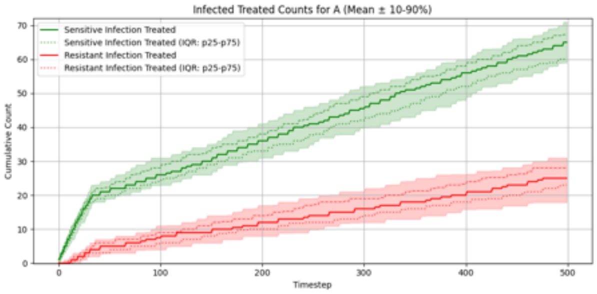

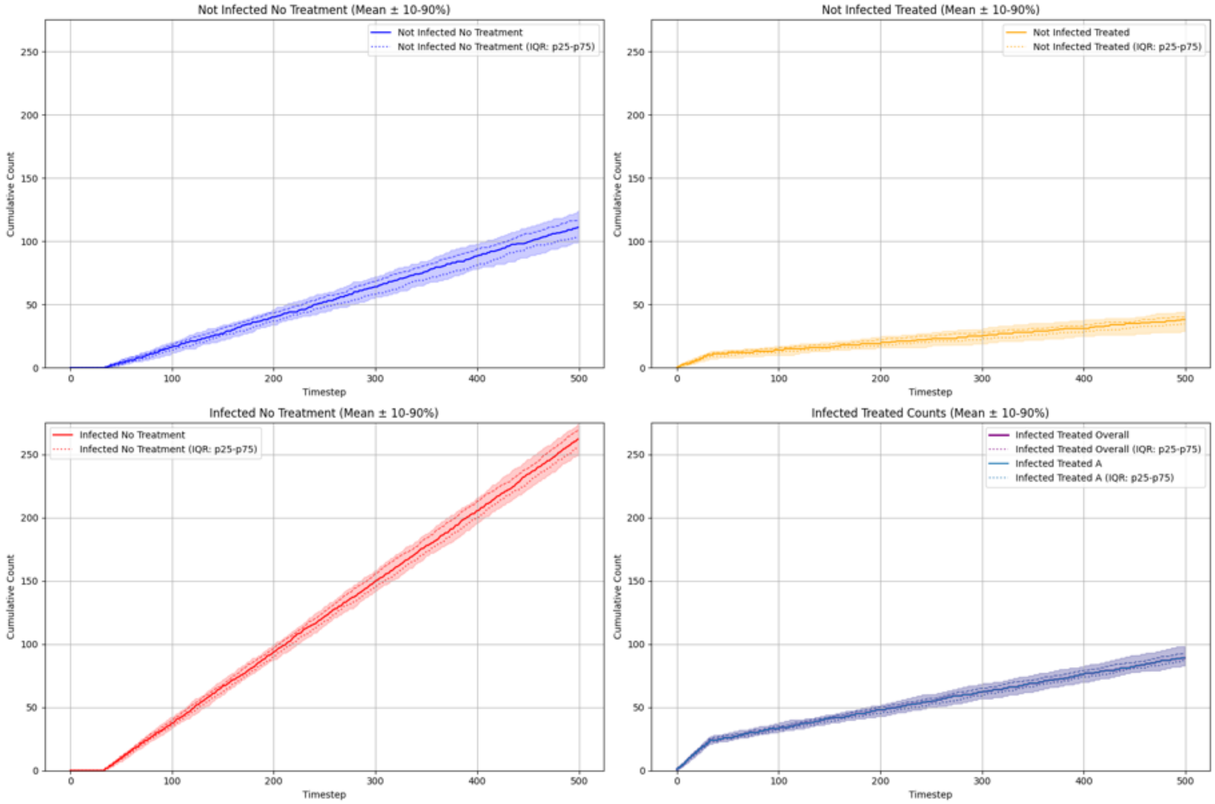

#### Summary Statistics (Single Antibiotic, Expected Reward Greedy)

| metric | p10 | p25 | p50 | p75 | p90 |
|---|---|---|---|---|---|
| overall_total_reward | -231.180 | -226.650 | -218.500 | -206.100 | -195.840 |
| overall_count_clinical_benefits | 47.000 | 48.000 | 52.000 | 60.000 | 63.100 |
| overall_count_clinical_failures | 231.000 | 238.000 | 245.000 | 253.500 | 260.100 |
| overall_count_adverse_events | 26.000 | 29.000 | 31.500 | 34.000 | 37.000 |
| overall_not_infected_no_treatment_count | 98.900 | 103.750 | 111.000 | 117.000 | 124.100 |
| overall_not_infected_treated_count | 28.900 | 34.750 | 38.000 | 40.000 | 44.100 |
| overall_infected_no_treatment_count | 248.900 | 256.000 | 262.000 | 269.250 | 274.100 |
| overall_infected_treated_count | 82.900 | 87.000 | 89.000 | 92.250 | 98.100 |
| overall_sensitive_infection_treated_count_per_abx_dict_A | 59.000 | 60.000 | 65.000 | 67.250 | 71.000 |
| overall_resistant_infection_treated_count_per_abx_dict_A | 17.900 | 23.000 | 25.000 | 28.000 | 31.000 |
| overall_sensitive_infection_treated_count | 59.000 | 60.000 | 65.000 | 67.250 | 71.000 |
| overall_resistant_infection_treated_count | 17.900 | 23.000 | 25.000 | 28.000 | 31.000 |
| overall_abx_prescriptions_count_per_abx_A | 82.900 | 87.000 | 89.000 | 92.250 | 98.100 |
| overall_abx_prescriptions_count | 82.900 | 87.000 | 89.000 | 92.250 | 98.100 |
| final_amr_actual_A | 0.327 | 0.327 | 0.327 | 0.327 | 0.327 |
| final_amr_visible_A | 0.327 | 0.327 | 0.327 | 0.327 | 0.327 |

#### Single Antibiotic, Expected Reward Lowest AMR

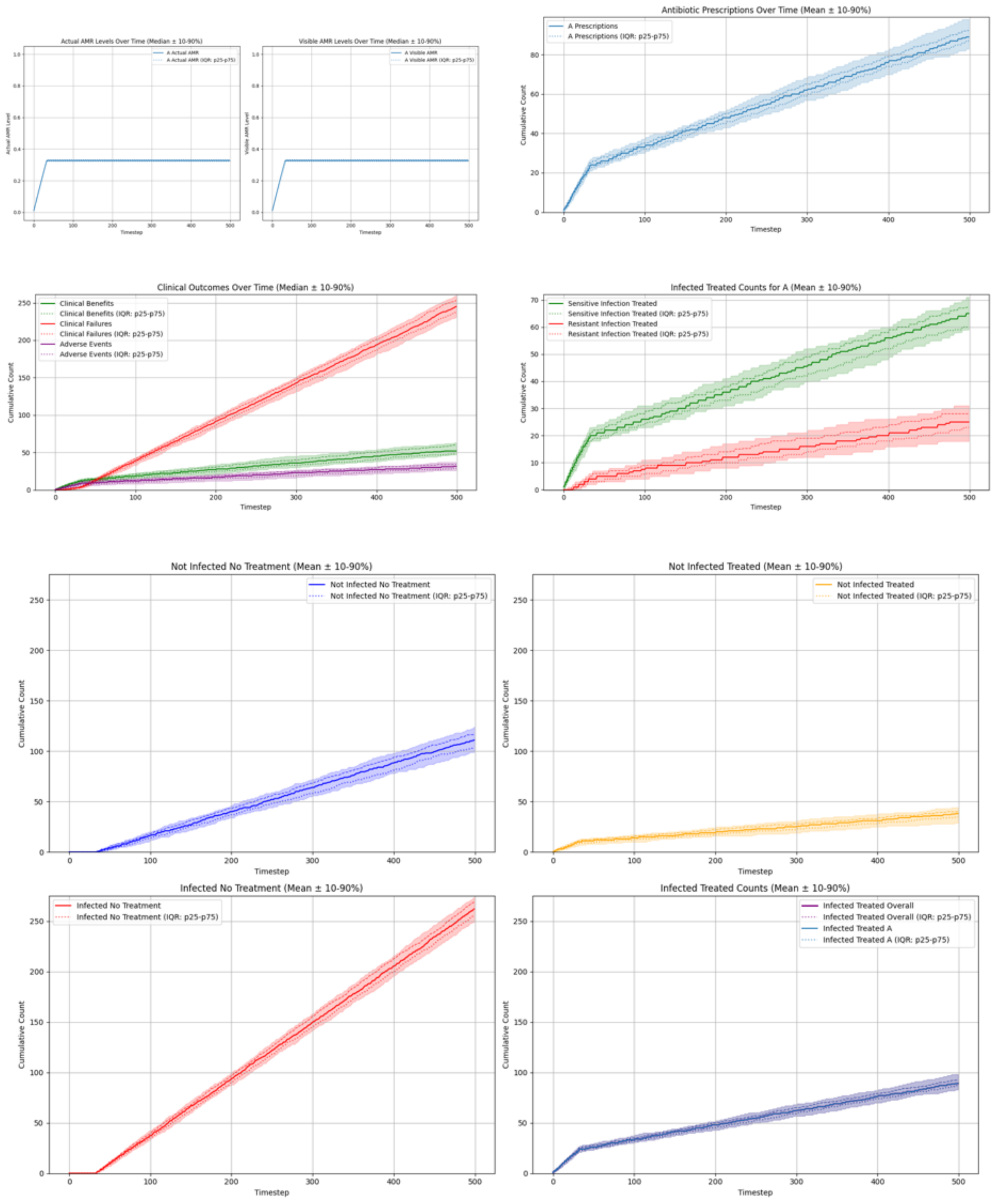

#### Summary Statistics (Single Antibiotic, Expected Reward Lowest AMR)

| metric | p10 | p25 | p50 | p75 | p90 |
|---|---|---|---|---|---|
| overall_total_reward | -231.180 | -226.650 | -218.500 | -206.100 | -195.840 |
| overall_count_clinical_benefits | 47.000 | 48.000 | 52.000 | 60.000 | 63.100 |
| overall_count_clinical_failures | 231.000 | 238.000 | 245.000 | 253.500 | 260.100 |
| overall_count_adverse_events | 26.000 | 29.000 | 31.500 | 34.000 | 37.000 |
| overall_not_infected_no_treatment_count | 98.900 | 103.750 | 111.000 | 117.000 | 124.100 |
| overall_not_infected_treated_count | 28.900 | 34.750 | 38.000 | 40.000 | 44.100 |
| overall_infected_no_treatment_count | 248.900 | 256.000 | 262.000 | 269.250 | 274.100 |
| overall_infected_treated_count | 82.900 | 87.000 | 89.000 | 92.250 | 98.100 |
| overall_sensitive_infection_treated_count_per_abx_dict_A | 59.000 | 60.000 | 65.000 | 67.250 | 71.000 |
| overall_resistant_infection_treated_count_per_abx_dict_A | 17.900 | 23.000 | 25.000 | 28.000 | 31.000 |
| overall_sensitive_infection_treated_count | 59.000 | 60.000 | 65.000 | 67.250 | 71.000 |
| overall_resistant_infection_treated_count | 17.900 | 23.000 | 25.000 | 28.000 | 31.000 |
| overall_abx_prescriptions_count_per_abx_A | 82.900 | 87.000 | 89.000 | 92.250 | 98.100 |
| overall_abx_prescriptions_count | 82.900 | 87.000 | 89.000 | 92.250 | 98.100 |
| final_amr_actual_A | 0.327 | 0.327 | 0.327 | 0.327 | 0.327 |
| final_amr_visible_A | 0.327 | 0.327 | 0.327 | 0.327 | 0.327 |

##### Two Antibiotics No Cross-Resistance, Expected Reward Greedy

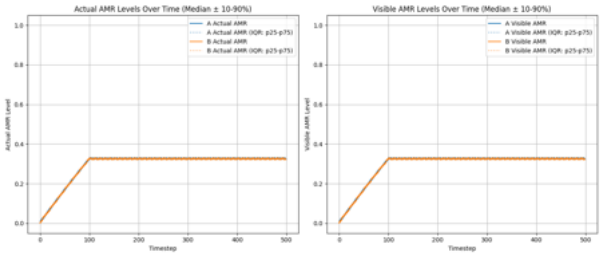

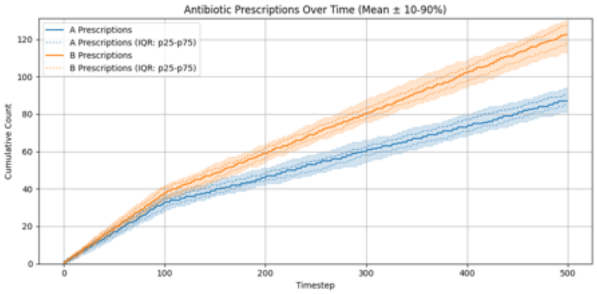

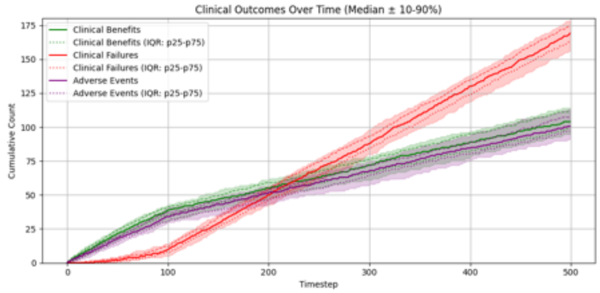

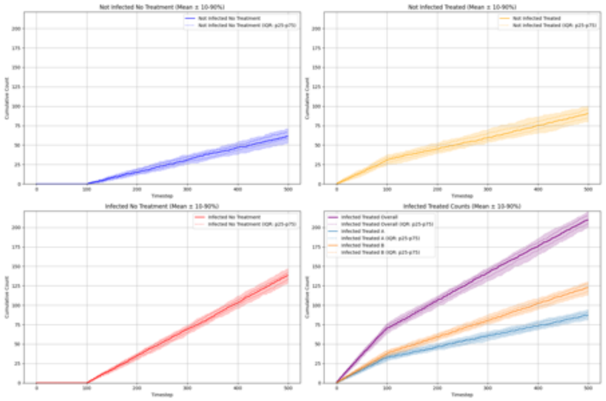

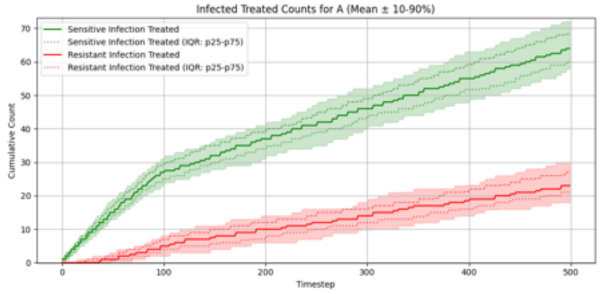

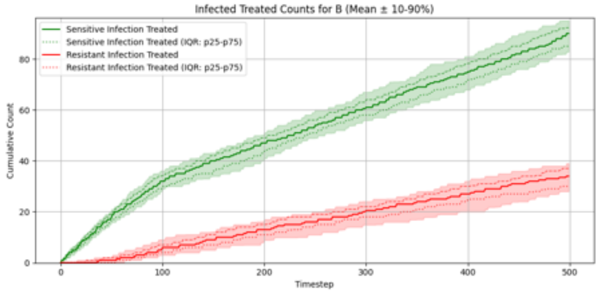

#### Summary Statistics (Two Antibiotics No Cross-Resistance, Expected Reward Greedy)

| metric | p10 | p25 | p50 | p75 | p90 |
|---|---|---|---|---|---|
| overall_total_reward | -142.180 | -135.300 | -123.750 | -114.100 | -108.360 |
| overall_count_clinical_benefits | 94.800 | 98.500 | 104.000 | 111.250 | 114.100 |
| overall_count_clinical_failures | 156.000 | 163.000 | 169.000 | 174.250 | 179.000 |
| overall_count_adverse_events | 90.900 | 97.000 | 100.500 | 107.250 | 113.000 |
| overall_not_infected_no_treatment_count | 52.900 | 58.000 | 61.500 | 66.250 | 71.100 |
| overall_not_infected_treated_count | 79.900 | 85.000 | 90.500 | 95.250 | 100.100 |
| overall_infected_no_treatment_count | 128.900 | 133.750 | 138.500 | 142.000 | 147.100 |
| overall_infected_treated_count | 199.900 | 204.750 | 209.500 | 215.000 | 220.100 |
| overall_sensitive_infection_treated_count_per_abx_dict_A | 57.900 | 60.000 | 64.000 | 68.250 | 72.100 |
| overall_sensitive_infection_treated_count_per_abx_dict_B | 82.800 | 85.000 | 90.000 | 92.250 | 95.100 |
| overall_resistant_infection_treated_count_per_abx_dict_A | 18.000 | 21.000 | 23.000 | 27.000 | 30.000 |
| overall_resistant_infection_treated_count_per_abx_dict_B | 28.000 | 30.000 | 34.000 | 37.000 | 39.000 |
| overall_sensitive_infection_treated_count | 143.900 | 147.750 | 153.500 | 158.250 | 162.100 |
| overall_resistant_infection_treated_count | 46.900 | 51.750 | 57.000 | 62.000 | 66.100 |
| overall_abx_prescriptions_count_per_abx_A | 81.000 | 85.000 | 87.000 | 90.250 | 94.000 |
| overall_abx_prescriptions_count_per_abx_B | 112.900 | 117.750 | 122.500 | 127.250 | 130.100 |
| overall_abx_prescriptions_count | 199.900 | 204.750 | 209.500 | 215.000 | 220.100 |
| final_amr_actual_A | 0.327 | 0.327 | 0.327 | 0.327 | 0.327 |
| final_amr_visible_A | 0.327 | 0.327 | 0.327 | 0.327 | 0.327 |
| final_amr_actual_B | 0.326 | 0.326 | 0.326 | 0.326 | 0.326 |
| final_amr_visible_B | 0.326 | 0.326 | 0.326 | 0.326 | 0.326 |

#### Two Antibiotics No Cross-Resistance, Expected Reward Lowest AMR

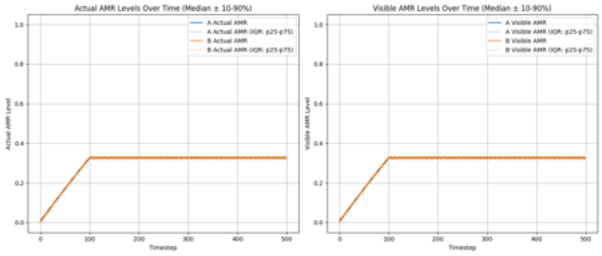

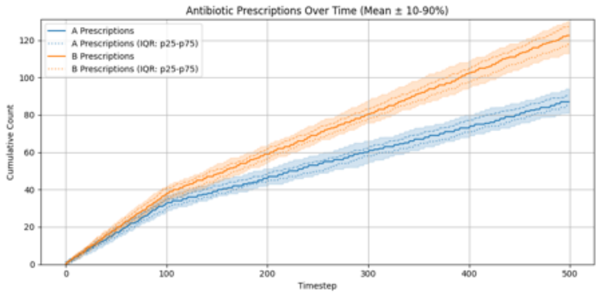

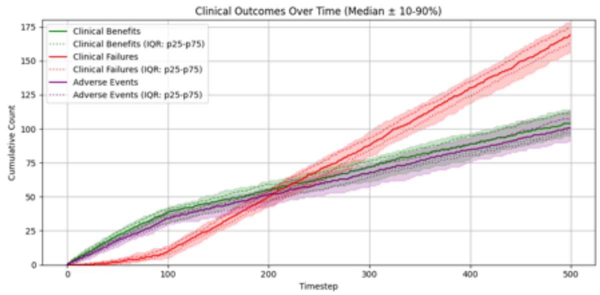

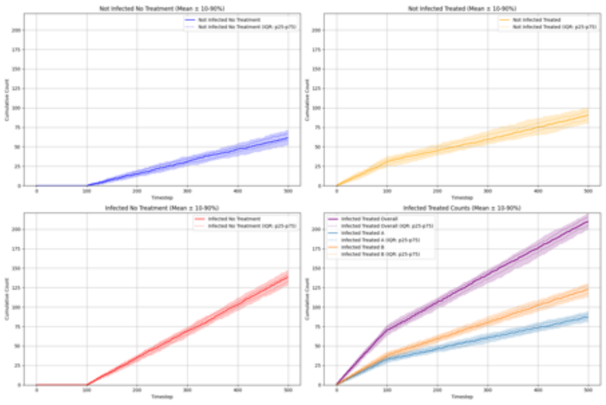

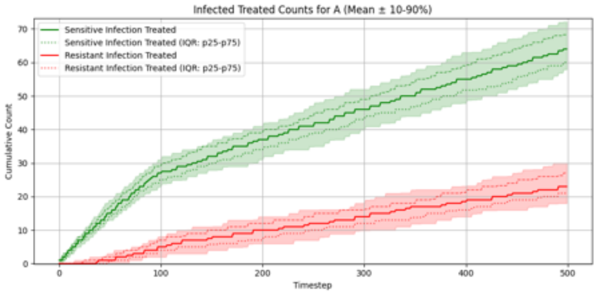

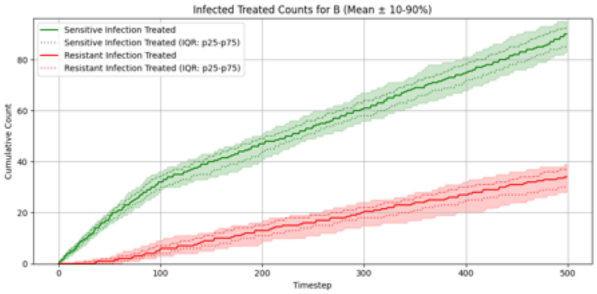

##### Summary Statistics (Two Antibiotics No Cross-Resistance, Expected Reward Lowest AMR)

| metric | p10 | p25 | p50 | p75 | p90 |
|---|---|---|---|---|---|
| overall_total_reward | -142.180 | -135.300 | -123.750 | -114.100 | -108.360 |
| overall_count_clinical_benefits | 94.800 | 98.500 | 104.000 | 111.250 | 114.100 |
| overall_count_clinical_failures | 156.000 | 163.000 | 169.000 | 174.250 | 179.000 |
| overall_count_adverse_events | 90.900 | 97.000 | 100.500 | 107.250 | 113.000 |
| overall_not_infected_no_treatment_count | 52.900 | 58.000 | 61.500 | 66.250 | 71.100 |
| overall_not_infected_treated_count | 79.900 | 85.000 | 90.500 | 95.250 | 100.100 |
| overall_infected_no_treatment_count | 128.900 | 133.750 | 138.500 | 142.000 | 147.100 |
| overall_infected_treated_count | 199.900 | 204.750 | 209.500 | 215.000 | 220.100 |
| overall_sensitive_infection_treated_count_per_abx_dict_A | 57.900 | 60.000 | 64.000 | 68.250 | 72.100 |
| overall_sensitive_infection_treated_count_per_abx_dict_B | 82.800 | 85.000 | 90.000 | 92.250 | 95.100 |
| overall_resistant_infection_treated_count_per_abx_dict_A | 18.000 | 21.000 | 23.000 | 27.000 | 30.000 |
| overall_resistant_infection_treated_count_per_abx_dict_B | 28.000 | 30.000 | 34.000 | 37.000 | 39.000 |
| overall_sensitive_infection_treated_count | 143.900 | 147.750 | 153.500 | 158.250 | 162.100 |
| overall_resistant_infection_treated_count | 46.900 | 51.750 | 57.000 | 62.000 | 66.100 |
| overall_abx_prescriptions_count_per_abx_A | 81.000 | 85.000 | 87.000 | 90.250 | 94.000 |
| overall_abx_prescriptions_count_per_abx_B | 112.900 | 117.750 | 122.500 | 127.250 | 130.100 |
| overall_abx_prescriptions_count | 199.900 | 204.750 | 209.500 | 215.000 | 220.100 |
| final_amr_actual_A | 0.327 | 0.327 | 0.327 | 0.327 | 0.327 |
| final_amr_visible_A | 0.327 | 0.327 | 0.327 | 0.327 | 0.327 |
| final_amr_actual_B | 0.326 | 0.326 | 0.326 | 0.326 | 0.326 |
| final_amr_visible_B | 0.326 | 0.326 | 0.326 | 0.326 | 0.326 |

##### Two Antibiotics With Cross-Resistance, Expected Reward Greedy

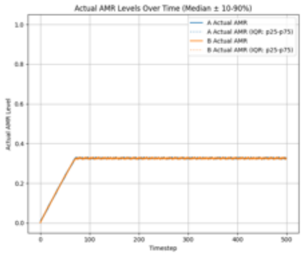

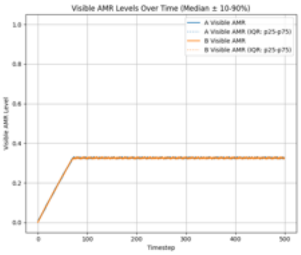

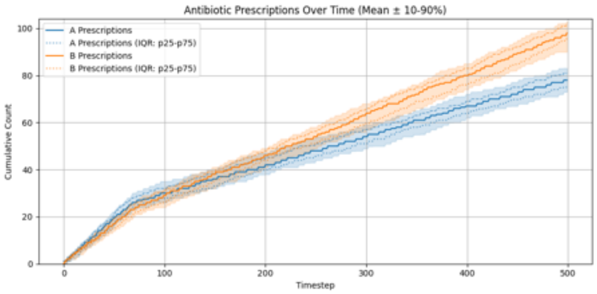

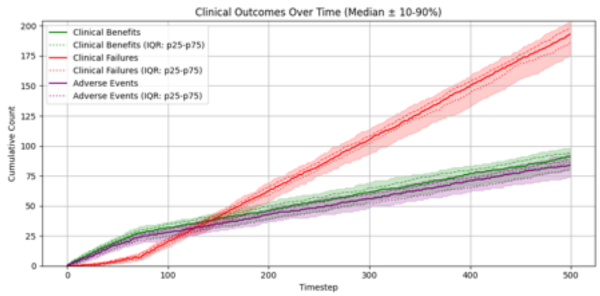

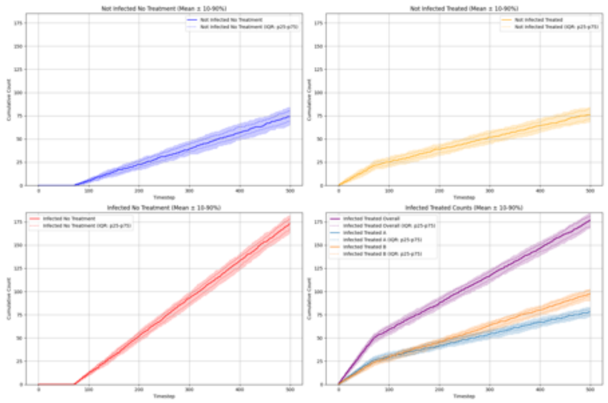

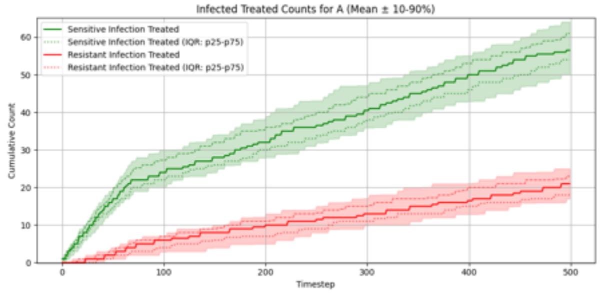

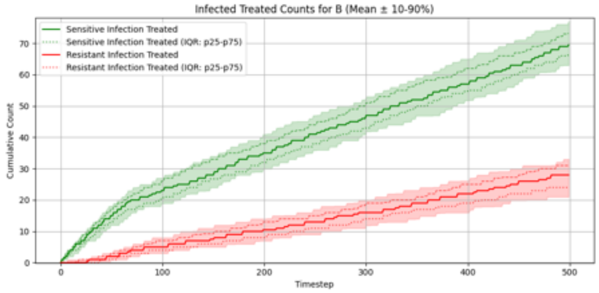

#### Summary Statistics (Two Antibiotics With Cross-Resistance, Expected Reward Greedy)

| metric | p10 | p25 | p50 | p75 | p90 |
|---|---|---|---|---|---|
| overall_total_reward | -167.040 | -160.325 | -154.350 | -140.875 | -132.010 |
| overall_count_clinical_benefits | 79.900 | 86.000 | 91.000 | 94.000 | 98.000 |
| overall_count_clinical_failures | 175.600 | 185.750 | 193.000 | 198.000 | 203.100 |
| overall_count_adverse_events | 73.900 | 80.500 | 84.000 | 88.000 | 93.000 |
| overall_not_infected_no_treatment_count | 65.000 | 69.000 | 74.000 | 80.000 | 84.000 |
| overall_not_infected_treated_count | 68.900 | 71.750 | 76.000 | 82.250 | 84.100 |
| overall_infected_no_treatment_count | 163.000 | 167.000 | 173.000 | 178.000 | 182.000 |
| overall_infected_treated_count | 168.900 | 170.750 | 177.000 | 181.250 | 184.100 |
| overall_sensitive_infection_treated_count_per_abx_dict_A | 50.000 | 54.000 | 56.500 | 61.000 | 64.100 |
| overall_sensitive_infection_treated_count_per_abx_dict_B | 63.000 | 66.750 | 69.500 | 73.250 | 77.100 |
| overall_resistant_infection_treated_count_per_abx_dict_A | 17.000 | 18.000 | 21.000 | 23.000 | 25.000 |
| overall_resistant_infection_treated_count_per_abx_dict_B | 21.000 | 24.000 | 28.000 | 31.000 | 33.100 |
| overall_sensitive_infection_treated_count | 118.900 | 121.000 | 128.000 | 131.250 | 135.100 |
| overall_resistant_infection_treated_count | 42.000 | 44.000 | 49.000 | 53.000 | 55.100 |
| overall_abx_prescriptions_count_per_abx_A | 73.000 | 75.000 | 78.000 | 81.000 | 83.100 |
| overall_abx_prescriptions_count_per_abx_B | 90.000 | 95.750 | 98.000 | 102.000 | 103.000 |
| overall_abx_prescriptions_count | 168.900 | 170.750 | 177.000 | 181.250 | 184.100 |
| final_amr_actual_A | 0.326 | 0.326 | 0.326 | 0.326 | 0.326 |
| final_amr_visible_A | 0.326 | 0.326 | 0.326 | 0.326 | 0.326 |
| final_amr_actual_B | 0.327 | 0.327 | 0.327 | 0.327 | 0.327 |
| final_amr_visible_B | 0.327 | 0.327 | 0.327 | 0.327 | 0.327 |

##### Two Antibiotics With Cross-Resistance, Expected Reward Lowest AMR

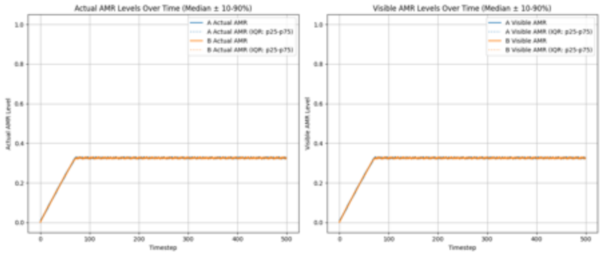

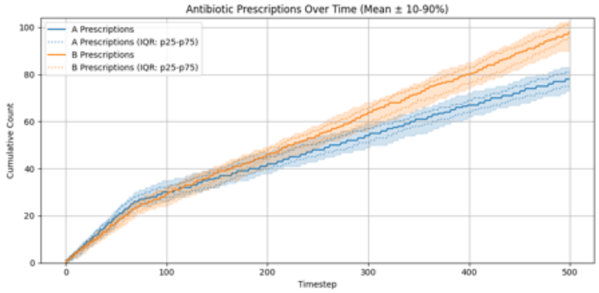

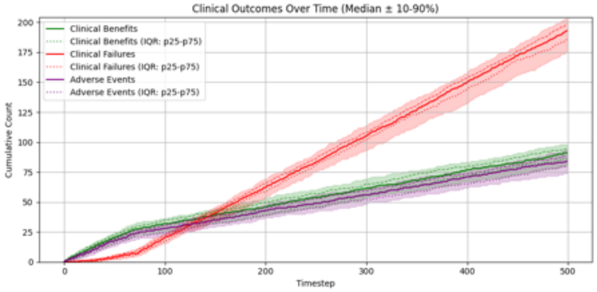

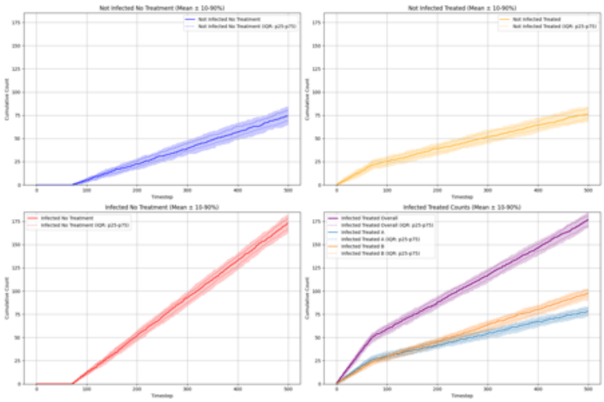

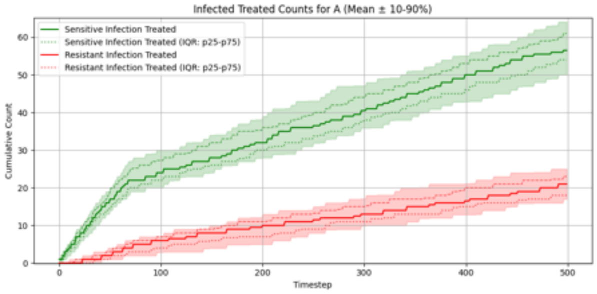

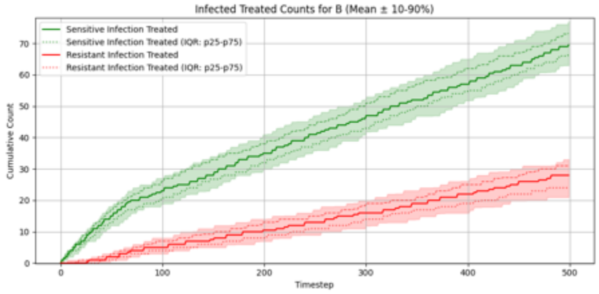

##### Summary Statistics (Two Antibiotics With Cross-Resistance, Expected Reward Lowest AMR)

| metric | p10 | p25 | p50 | p75 | p90 |
|---|---|---|---|---|---|
| overall_total_reward | -167.040 | -160.325 | -154.350 | -140.875 | -132.010 |
| overall_count_clinical_benefits | 79.900 | 86.000 | 91.000 | 94.000 | 98.000 |
| overall_count_clinical_failures | 175.600 | 185.750 | 193.000 | 198.000 | 203.100 |
| overall_count_adverse_events | 73.900 | 80.500 | 84.000 | 88.000 | 93.000 |
| overall_not_infected_no_treatment_count | 65.000 | 69.000 | 74.000 | 80.000 | 84.000 |
| overall_not_infected_treated_count | 68.900 | 71.750 | 76.000 | 82.250 | 84.100 |
| overall_infected_no_treatment_count | 163.000 | 167.000 | 173.000 | 178.000 | 182.000 |
| overall_infected_treated_count | 168.900 | 170.750 | 177.000 | 181.250 | 184.100 |
| overall_sensitive_infection_treated_count_per_abx_dict_A | 50.000 | 54.000 | 56.500 | 61.000 | 64.100 |
| overall_sensitive_infection_treated_count_per_abx_dict_B | 63.000 | 66.750 | 69.500 | 73.250 | 77.100 |
| overall_resistant_infection_treated_count_per_abx_dict_A | 17.000 | 18.000 | 21.000 | 23.000 | 25.000 |
| overall_resistant_infection_treated_count_per_abx_dict_B | 21.000 | 24.000 | 28.000 | 31.000 | 33.100 |
| overall_sensitive_infection_treated_count | 118.900 | 121.000 | 128.000 | 131.250 | 135.100 |
| overall_resistant_infection_treated_count | 42.000 | 44.000 | 49.000 | 53.000 | 55.100 |
| overall_abx_prescriptions_count_per_abx_A | 73.000 | 75.000 | 78.000 | 81.000 | 83.100 |
| overall_abx_prescriptions_count_per_abx_B | 90.000 | 95.750 | 98.000 | 102.000 | 103.000 |
| overall_abx_prescriptions_count | 168.900 | 170.750 | 177.000 | 181.250 | 184.100 |
| final_amr_actual_A | 0.326 | 0.326 | 0.326 | 0.326 | 0.326 |
| final_amr_visible_A | 0.326 | 0.326 | 0.326 | 0.326 | 0.326 |
| final_amr_actual_B | 0.327 | 0.327 | 0.327 | 0.327 | 0.327 |
| final_amr_visible_B | 0.327 | 0.327 | 0.327 | 0.327 | 0.327 |

### Experiment Set 2

#### Single Antibiotic, Expected Reward Greedy

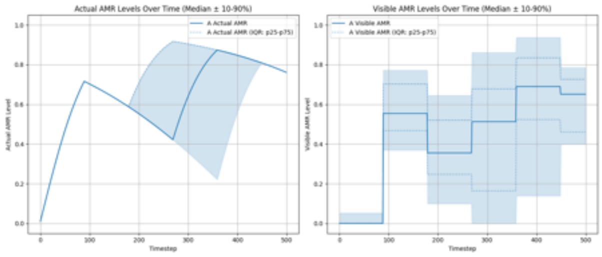

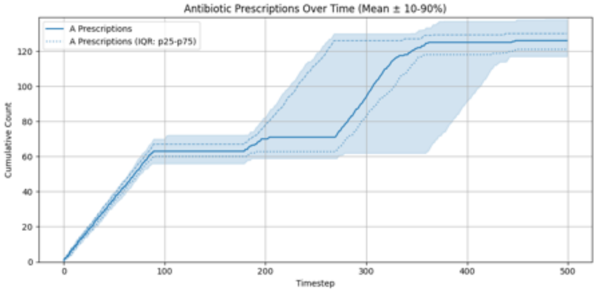

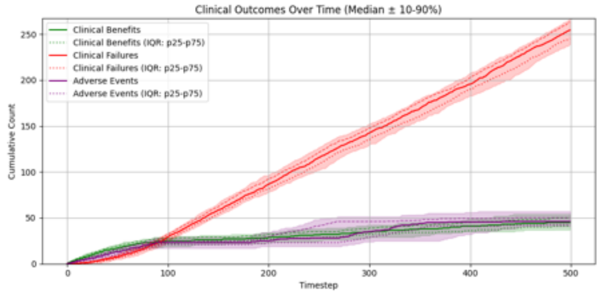

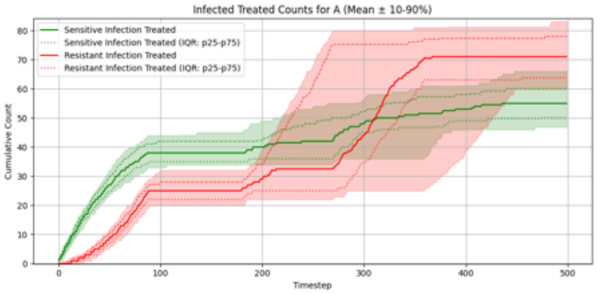

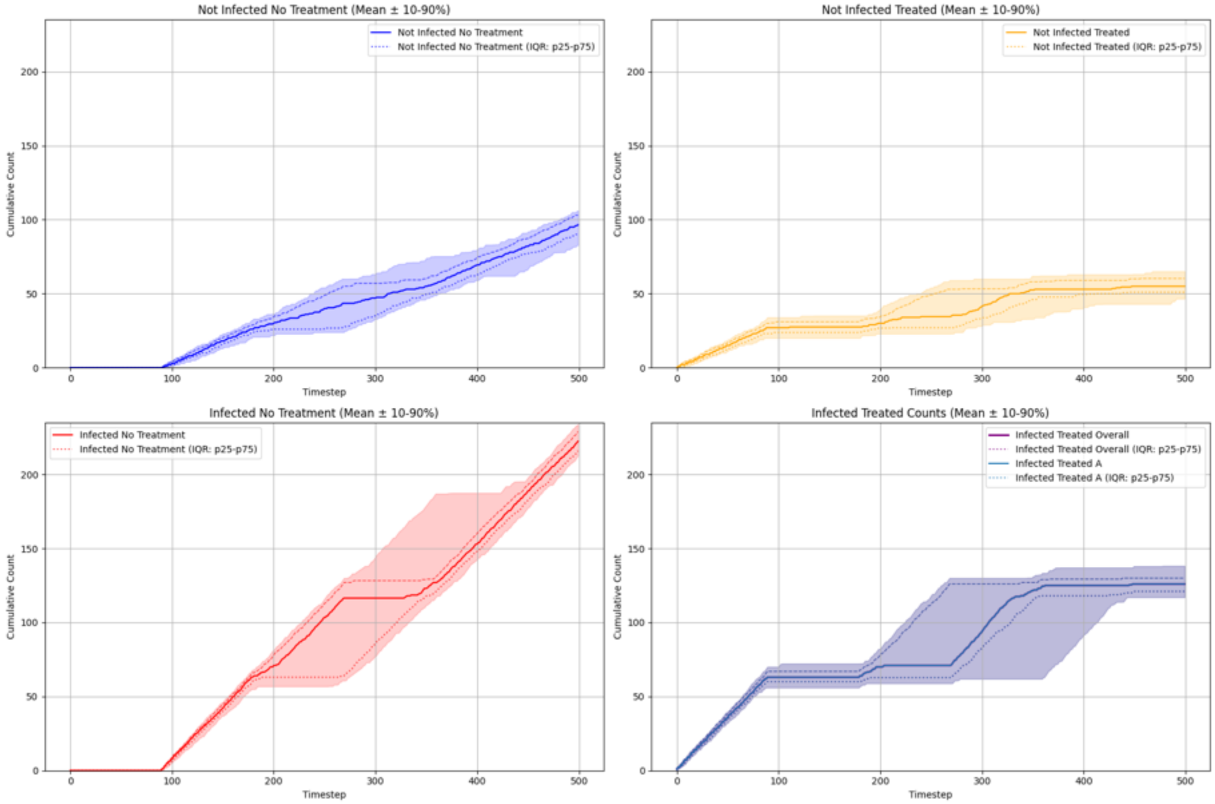

#### Summary Statistics (Single Antibiotic, Expected Reward Greedy)

| metric | p10 | p25 | p50 | p75 | p90 |
|---|---|---|---|---|---|
| overall_total_reward | -268.280 | -257.850 | -249.000 | -234.750 | -225.520 |
| overall_count_clinical_benefits | 37.000 | 41.000 | 45.000 | 50.250 | 54.100 |
| overall_count_clinical_failures | 239.000 | 245.000 | 254.500 | 263.000 | 266.100 |
| overall_count_adverse_events | 40.800 | 42.750 | 46.000 | 50.000 | 57.000 |
| overall_not_infected_no_treatment_count | 82.700 | 90.000 | 96.500 | 103.000 | 106.100 |
| overall_not_infected_treated_count | 46.600 | 51.000 | 55.000 | 60.250 | 65.100 |
| overall_infected_no_treatment_count | 212.600 | 216.000 | 222.500 | 229.000 | 234.100 |
| overall_infected_treated_count | 117.000 | 121.000 | 126.000 | 130.000 | 138.200 |
| overall_sensitive_infection_treated_count_per_abx_dict_A | 46.900 | 50.000 | 55.000 | 60.000 | 66.000 |
| overall_resistant_infection_treated_count_per_abx_dict_A | 59.900 | 63.750 | 71.000 | 78.000 | 83.100 |
| overall_sensitive_infection_treated_count | 46.900 | 50.000 | 55.000 | 60.000 | 66.000 |
| overall_resistant_infection_treated_count | 59.900 | 63.750 | 71.000 | 78.000 | 83.100 |
| overall_abx_prescriptions_count_per_abx_A | 117.000 | 121.000 | 126.000 | 130.000 | 138.200 |
| overall_abx_prescriptions_count | 117.000 | 121.000 | 126.000 | 130.000 | 138.200 |
| final_amr_actual_A | 0.766 | 0.766 | 0.766 | 0.766 | 0.766 |
| final_amr_visible_A | 0.399 | 0.460 | 0.651 | 0.726 | 0.786 |

**Single Antibiotic, Expected Reward Lowest AMR**

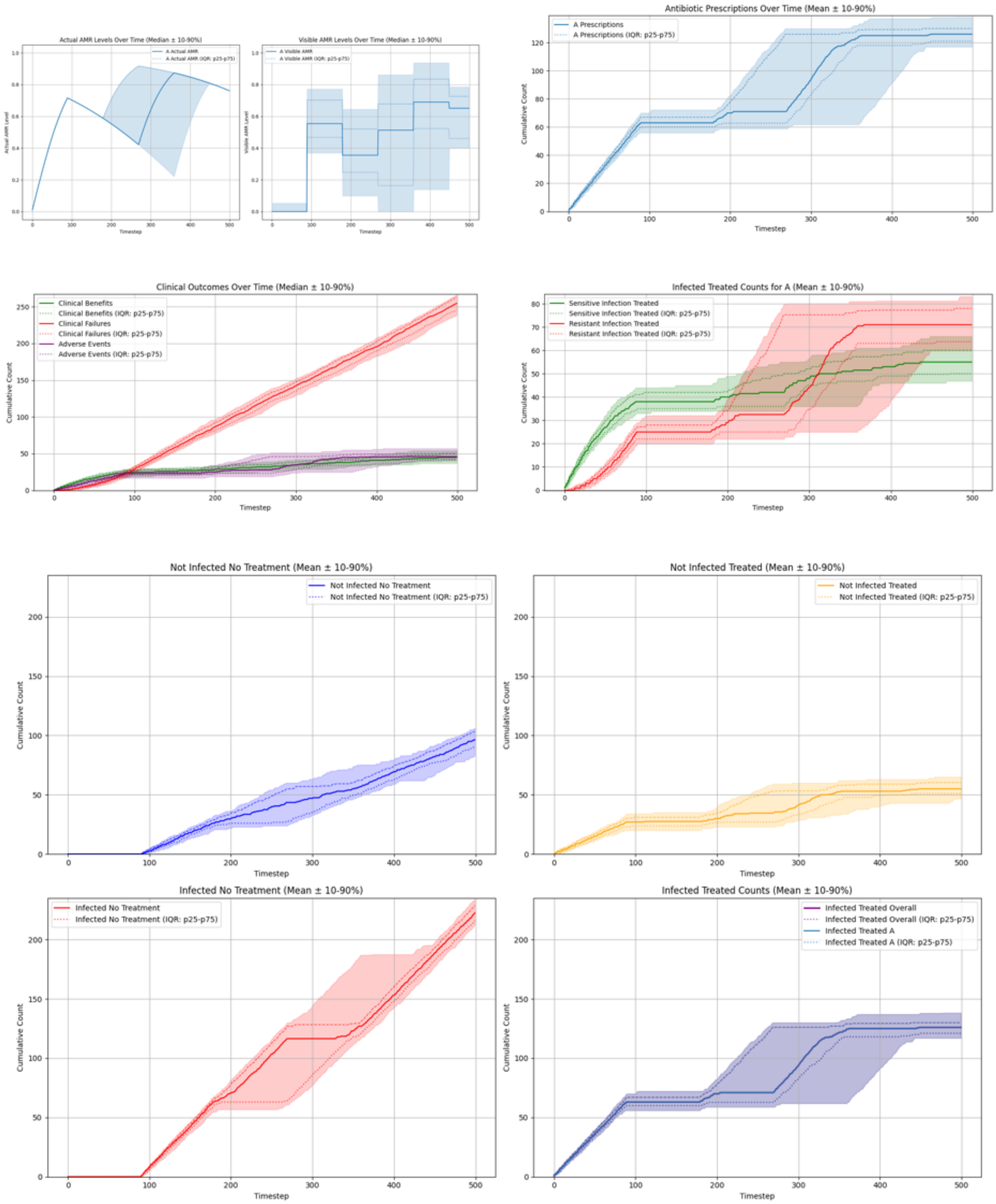

**Summary Statistics (Single Antibiotic, Expected Reward Lowest AMR)**

| metric | p10 | p25 | p50 | p75 | p90 |
|---|---|---|---|---|---|
| overall_total_reward | -268.280 | -257.850 | -249.000 | -234.750 | -225.520 |
| overall_count_clinical_benefits | 37.000 | 41.000 | 45.000 | 50.250 | 54.100 |
| overall_count_clinical_failures | 239.000 | 245.000 | 254.500 | 263.000 | 266.100 |
| overall_count_adverse_events | 40.800 | 42.750 | 46.000 | 50.000 | 57.000 |
| overall_not_infected_no_treatment_count | 82.700 | 90.000 | 96.500 | 103.000 | 106.100 |
| overall_not_infected_treated_count | 46.600 | 51.000 | 55.000 | 60.250 | 65.100 |
| overall_infected_no_treatment_count | 212.600 | 216.000 | 222.500 | 229.000 | 234.100 |
| overall_infected_treated_count | 117.000 | 121.000 | 126.000 | 130.000 | 138.200 |
| overall_sensitive_infection_treated_count_per_abx_dict_A | 46.900 | 50.000 | 55.000 | 60.000 | 66.000 |
| overall_resistant_infection_treated_count_per_abx_dict_A | 59.900 | 63.750 | 71.000 | 78.000 | 83.100 |
| overall_sensitive_infection_treated_count | 46.900 | 50.000 | 55.000 | 60.000 | 66.000 |
| overall_resistant_infection_treated_count | 59.900 | 63.750 | 71.000 | 78.000 | 83.100 |
| overall_abx_prescriptions_count_per_abx_A | 117.000 | 121.000 | 126.000 | 130.000 | 138.200 |
| overall_abx_prescriptions_count | 117.000 | 121.000 | 126.000 | 130.000 | 138.200 |
| final_amr_actual_A | 0.766 | 0.766 | 0.766 | 0.766 | 0.766 |
| final_amr_visible_A | 0.399 | 0.460 | 0.651 | 0.726 | 0.786 |

#### Two Antibiotics No Cross-Resistance, Expected Reward Greedy

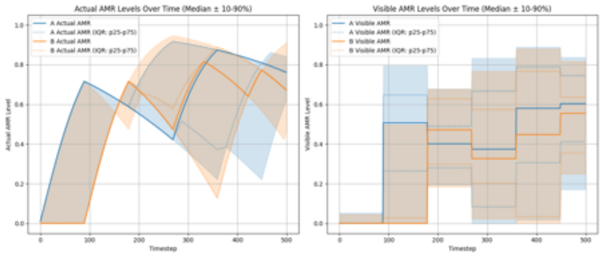

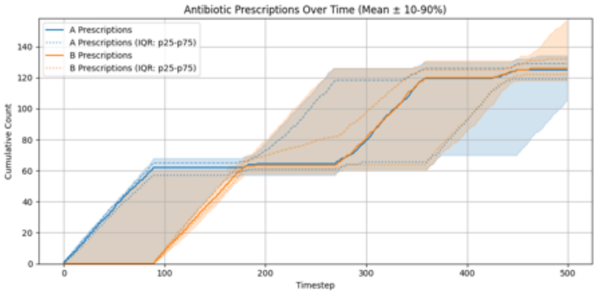

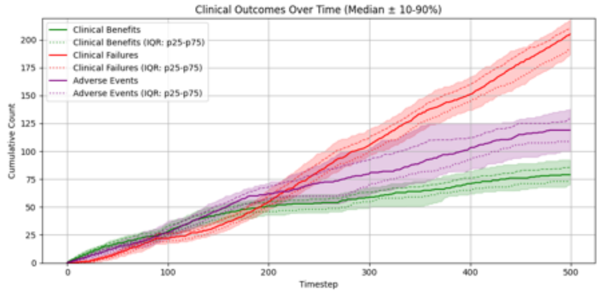

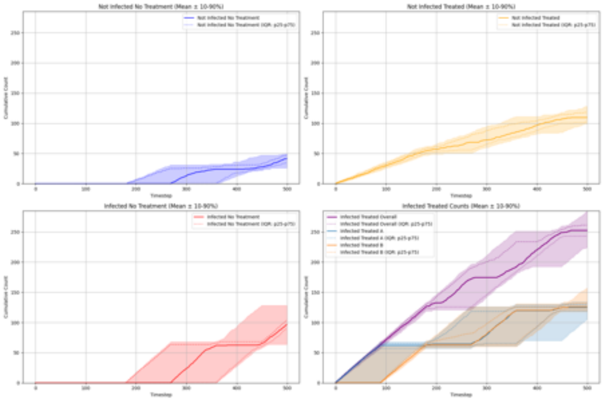

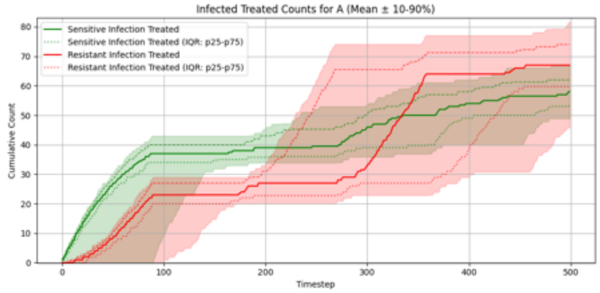

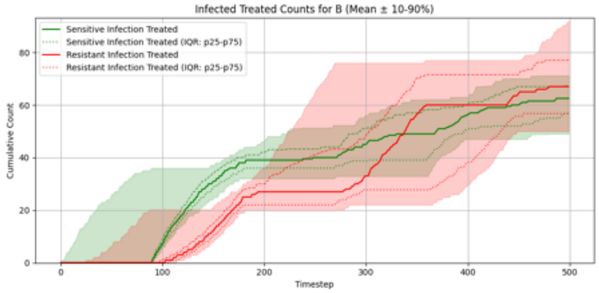

#### Summary Statistics (Two Antibiotics No Cross-Resistance, Expected Reward Greedy)

| metric | p10 | p25 | p50 | p75 | p90 |
|---|---|---|---|---|---|
| overall__total__reward | -221.530 | -210.650 | -197.150 | -180.725 | -170.750 |
| overall__count__clinical__benefits | 70.000 | 73.000 | 79.000 | 85.500 | 92.300 |
| overall__count__clinical__failures | 186.700 | 191.750 | 205.000 | 210.250 | 218.100 |
| overall__count__adverse__events | 99.800 | 108.750 | 119.000 | 129.000 | 138.000 |
| overall__not__infected__no__treatment__count | 26.000 | 34.750 | 41.500 | 47.000 | 50.000 |
| overall__not__infected__treated__count | 98.900 | 102.000 | 109.500 | 118.250 | 128.100 |
| overall__infected__no__treatment__count | 63.800 | 89.750 | 96.500 | 104.000 | 128.200 |
| overall__infected__treated__count | 223.900 | 243.000 | 252.500 | 261.250 | 284.000 |
| overall__sensitive__infection__treated__count__per__abx__dict__A | 48.900 | 53.750 | 58.000 | 62.000 | 67.000 |
| overall__sensitive__infection__treated__count__per__abx__dict__B | 49.000 | 56.750 | 62.500 | 67.250 | 71.200 |
| overall__resistant__infection__treated__count__per__abx__dict__A | 45.800 | 59.750 | 67.000 | 74.000 | 82.000 |
| overall__resistant__infection__treated__count__per__abx__dict__B | 50.000 | 56.750 | 67.000 | 77.250 | 92.100 |
| overall__sensitive__infection__treated__count | 105.800 | 111.000 | 119.000 | 127.000 | 133.000 |
| overall__resistant__infection__treated__count | 105.700 | 117.750 | 131.000 | 150.000 | 165.000 |
| overall__abx__prescriptions__count__per__abx__A | 104.900 | 119.000 | 125.000 | 129.000 | 134.200 |
| overall__abx__prescriptions__count__per__abx__B | 118.000 | 122.000 | 126.000 | 132.000 | 157.100 |
| overall__abx__prescriptions__count | 223.900 | 243.000 | 252.500 | 261.250 | 284.000 |
| final__amr__actual__A | 0.591 | 0.766 | 0.766 | 0.766 | 0.844 |
| final__amr__visible__A | 0.171 | 0.412 | 0.604 | 0.745 | 0.837 |
| final__amr__actual__B | 0.425 | 0.682 | 0.682 | 0.682 | 0.902 |
| final__amr__visible__B | 0.249 | 0.355 | 0.555 | 0.636 | 0.817 |

#### Two Antibiotics No Cross-Resistance, Expected Reward Lowest AMR

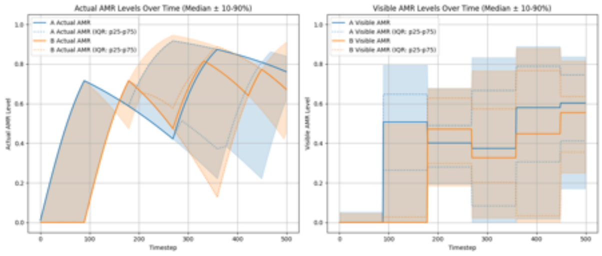

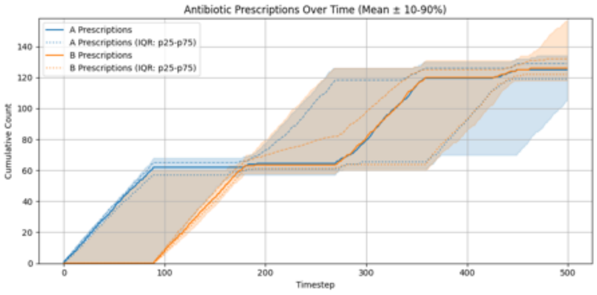

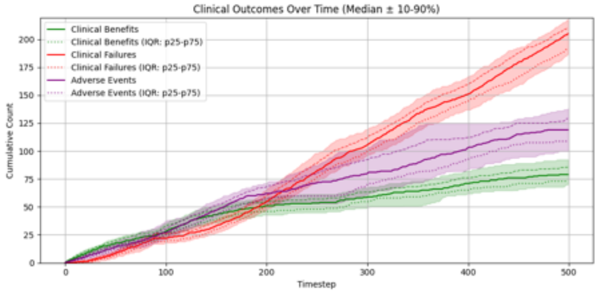

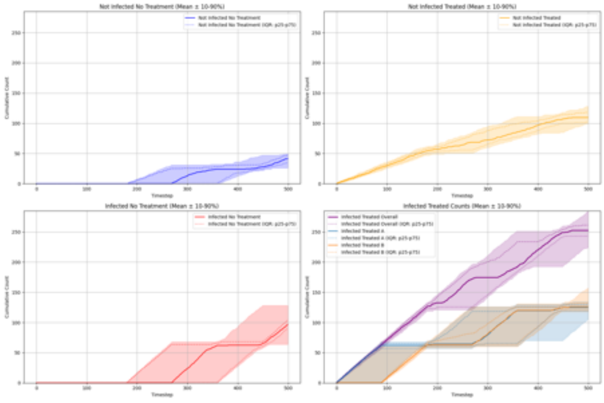

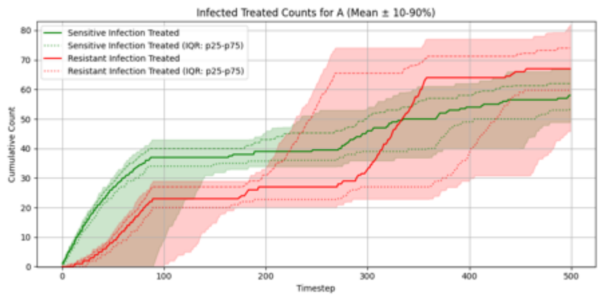

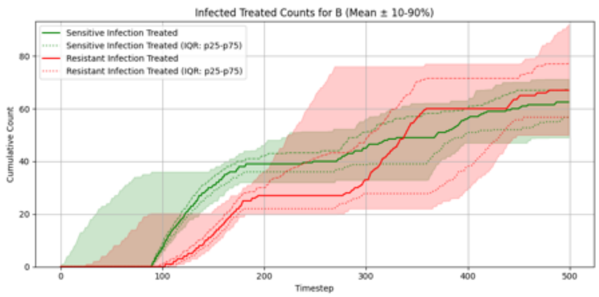

#### Summary Statistics (Two Antibiotics No Cross-Resistance, Expected Reward Lowest AMR)

| metric | p10 | p25 | p50 | p75 | p90 |
|---|---|---|---|---|---|
| overall_total_reward | -221.530 | -210.650 | -197.150 | -180.725 | -170.750 |
| overall_count_clinical_benefits | 70.000 | 73.000 | 79.000 | 85.500 | 92.300 |
| overall_count_clinical_failures | 186.700 | 191.750 | 205.000 | 210.250 | 218.100 |
| overall_count_adverse_events | 99.800 | 108.750 | 119.000 | 129.000 | 138.000 |
| overall_not_infected_no_treatment_count | 26.000 | 34.750 | 41.500 | 47.000 | 50.000 |
| overall_not_infected_treated_count | 98.900 | 102.000 | 109.500 | 118.250 | 128.100 |
| overall_infected_no_treatment_count | 63.800 | 89.750 | 96.500 | 104.000 | 128.200 |
| overall_infected_treated_count | 223.900 | 243.000 | 252.500 | 261.250 | 284.000 |
| overall_sensitive_infection_treated_count_per_abx_dict_A | 48.900 | 53.750 | 58.000 | 62.000 | 67.000 |
| overall_sensitive_infection_treated_count_per_abx_dict_B | 49.000 | 56.750 | 62.500 | 67.250 | 71.200 |
| overall_resistant_infection_treated_count_per_abx_dict_A | 45.800 | 59.750 | 67.000 | 74.000 | 82.000 |
| overall_resistant_infection_treated_count_per_abx_dict_B | 50.000 | 56.750 | 67.000 | 77.250 | 92.100 |
| overall_sensitive_infection_treated_count | 105.800 | 111.000 | 119.000 | 127.000 | 133.000 |
| overall_resistant_infection_treated_count | 105.700 | 117.750 | 131.000 | 150.000 | 165.000 |
| overall_abx_prescriptions_count_per_abx_A | 104.900 | 119.000 | 125.000 | 129.000 | 134.200 |
| overall_abx_prescriptions_count_per_abx_B | 118.000 | 122.000 | 126.000 | 132.000 | 157.100 |
| overall_abx_prescriptions_count | 223.900 | 243.000 | 252.500 | 261.250 | 284.000 |
| final_amr_actual_A | 0.591 | 0.766 | 0.766 | 0.766 | 0.844 |
| final_amr_visible_A | 0.171 | 0.412 | 0.604 | 0.745 | 0.837 |
| final_amr_actual_B | 0.425 | 0.682 | 0.682 | 0.682 | 0.902 |
| final_amr_visible_B | 0.249 | 0.355 | 0.555 | 0.636 | 0.817 |

#### Two Antibiotics With Cross-Resistance, Expected Reward Greedy

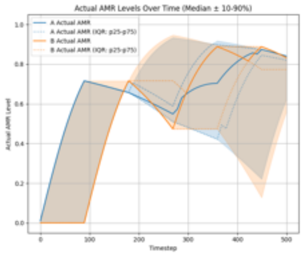

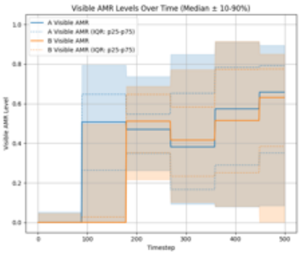

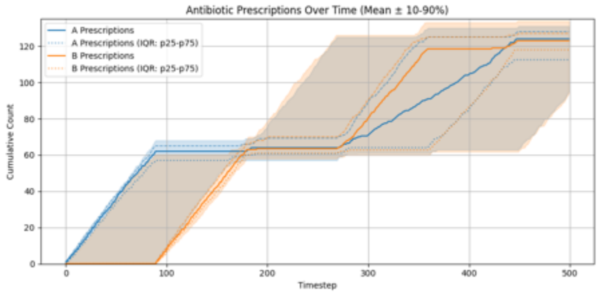

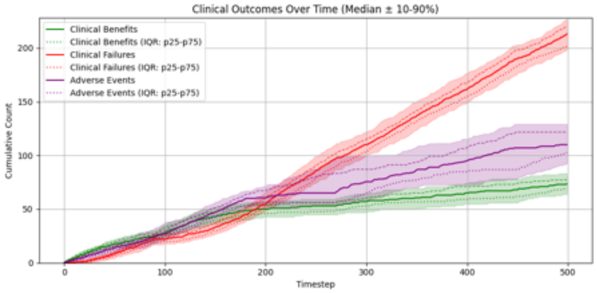

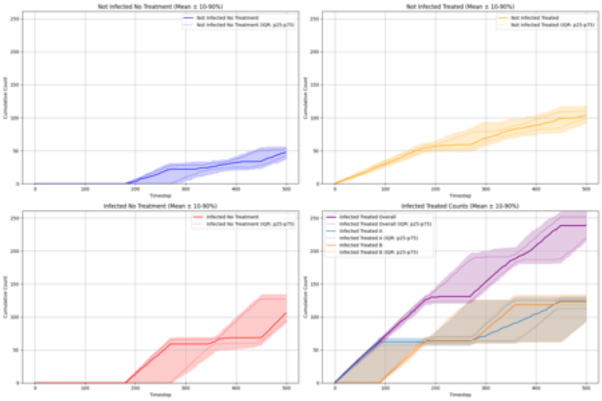

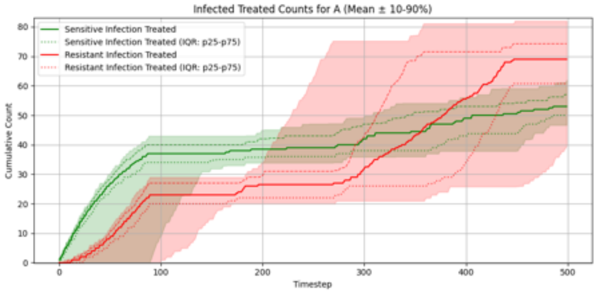

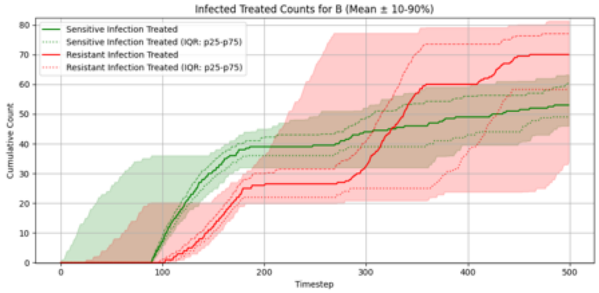

#### Summary Statistics (Two Antibiotics With Cross-Resistance, Expected Reward Greedy)

| metric | p10 | p25 | p50 | p75 | p90 |
|---|---|---|---|---|---|
| overall_total_reward | -234.470 | -217.975 | -204.400 | -194.625 | -180.330 |
| overall_count_clinical_benefits | 64.000 | 68.750 | 73.500 | 77.250 | 83.200 |
| overall_count_clinical_failures | 198.800 | 201.750 | 213.000 | 220.250 | 227.000 |
| overall_count_adverse_events | 92.800 | 102.750 | 110.000 | 121.750 | 129.100 |
| overall_not_infected_no_treatment_count | 36.900 | 42.000 | 47.500 | 53.000 | 56.000 |
| overall_not_infected_treated_count | 91.900 | 98.000 | 103.500 | 110.000 | 118.100 |
| overall_infected_no_treatment_count | 91.000 | 95.000 | 106.000 | 127.250 | 134.100 |
| overall_infected_treated_count | 215.900 | 219.000 | 239.500 | 252.000 | 260.000 |
| overall_sensitive_infection_treated_count_per_abx_dict_A | 46.700 | 50.000 | 53.000 | 57.000 | 62.000 |
| overall_sensitive_infection_treated_count_per_abx_dict_B | 46.000 | 49.000 | 53.000 | 60.250 | 63.200 |
| overall_resistant_infection_treated_count_per_abx_dict_A | 39.800 | 60.750 | 69.000 | 74.250 | 82.000 |
| overall_resistant_infection_treated_count_per_abx_dict_B | 34.000 | 58.250 | 70.000 | 77.000 | 81.300 |
| overall_sensitive_infection_treated_count | 96.800 | 100.750 | 108.000 | 113.250 | 119.100 |
| overall_resistant_infection_treated_count | 101.000 | 108.750 | 132.000 | 147.000 | 155.200 |
| overall_abx_prescriptions_count_per_abx_A | 94.900 | 112.500 | 124.000 | 128.000 | 131.000 |
| overall_abx_prescriptions_count_per_abx_B | 93.900 | 118.000 | 123.000 | 127.000 | 134.000 |
| overall_abx_prescriptions_count | 215.900 | 219.000 | 239.500 | 252.000 | 260.000 |
| final_amr_actual_A | 0.591 | 0.823 | 0.844 | 0.844 | 0.844 |
| final_amr_visible_A | 0.085 | 0.351 | 0.658 | 0.792 | 0.895 |
| final_amr_actual_B | 0.525 | 0.773 | 0.839 | 0.839 | 0.839 |
| final_amr_visible_B | 0.000 | 0.384 | 0.631 | 0.775 | 0.851 |

##### Two Antibiotics With Cross-Resistance, Expected Reward Lowest AMR

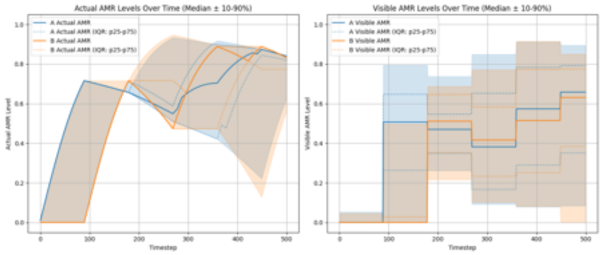

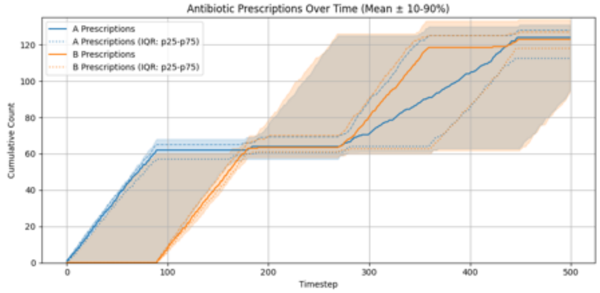

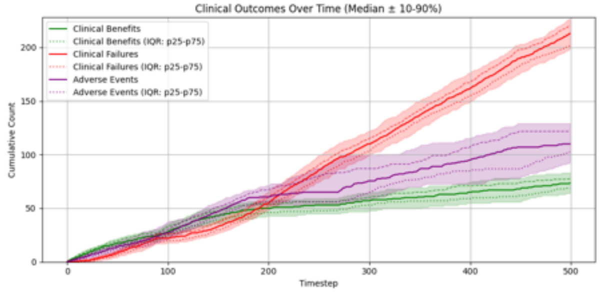

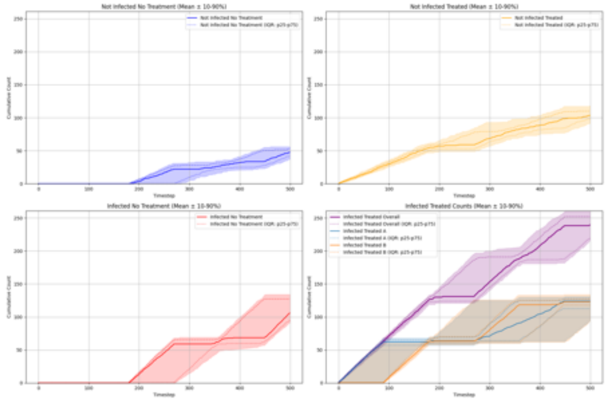

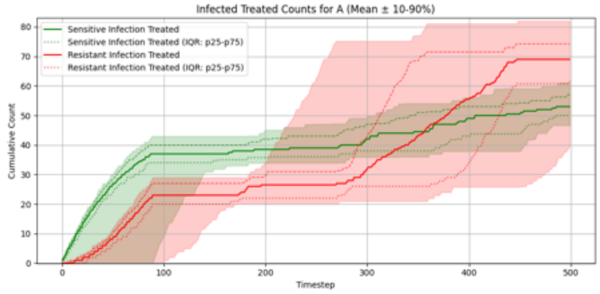

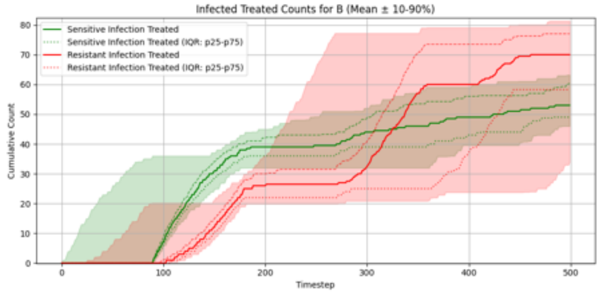

##### Summary Statistics (Two Antibiotics With Cross-Resistance, Expected Reward Lowest AMR)

| metric | p10 | p25 | p50 | p75 | p90 |
|---|---|---|---|---|---|
| overall__total__reward | -234.470 | -217.975 | -204.400 | -194.625 | -180.330 |
| overall__count__clinical__benefits | 64.000 | 68.750 | 73.500 | 77.250 | 83.200 |
| overall__count__clinical__failures | 198.800 | 201.750 | 213.000 | 220.250 | 227.000 |
| overall__count__adverse__events | 92.800 | 102.750 | 110.000 | 121.750 | 129.100 |
| overall__not__infected__no__treatment__count | 36.900 | 42.000 | 47.500 | 53.000 | 56.000 |
| overall__not__infected__treated__count | 91.900 | 98.000 | 103.500 | 110.000 | 118.100 |
| overall__infected__no__treatment__count | 91.000 | 95.000 | 106.000 | 127.250 | 134.100 |
| overall__infected__treated__count | 215.900 | 219.000 | 239.500 | 252.000 | 260.000 |
| overall__sensitive__infection__treated__count__per__abx__dict__A | 46.700 | 50.000 | 53.000 | 57.000 | 62.000 |
| overall__sensitive__infection__treated__count__per__abx__dict__B | 46.000 | 49.000 | 53.000 | 60.250 | 63.200 |
| overall__resistant__infection__treated__count__per__abx__dict__A | 39.800 | 60.750 | 69.000 | 74.250 | 82.000 |
| overall__resistant__infection__treated__count__per__abx__dict__B | 34.000 | 58.250 | 70.000 | 77.000 | 81.300 |
| overall__sensitive__infection__treated__count | 96.800 | 100.750 | 108.000 | 113.250 | 119.100 |
| overall__resistant__infection__treated__count | 101.000 | 108.750 | 132.000 | 147.000 | 155.200 |
| overall__abx__prescriptions__count__per__abx__A | 94.900 | 112.500 | 124.000 | 128.000 | 131.000 |
| overall__abx__prescriptions__count__per__abx__B | 93.900 | 118.000 | 123.000 | 127.000 | 134.000 |
| overall__abx__prescriptions__count | 215.900 | 219.000 | 239.500 | 252.000 | 260.000 |
| final__amr__actual__A | 0.591 | 0.823 | 0.844 | 0.844 | 0.844 |
| final__amr__visible__A | 0.085 | 0.351 | 0.658 | 0.792 | 0.895 |
| final__amr__actual__B | 0.525 | 0.773 | 0.839 | 0.839 | 0.839 |
| final__amr__visible__B | 0.000 | 0.384 | 0.631 | 0.775 | 0.851 |

### Experiment Set 3

#### Single Antibiotic, Accurate Risk Stratification, Expected Reward Greedy

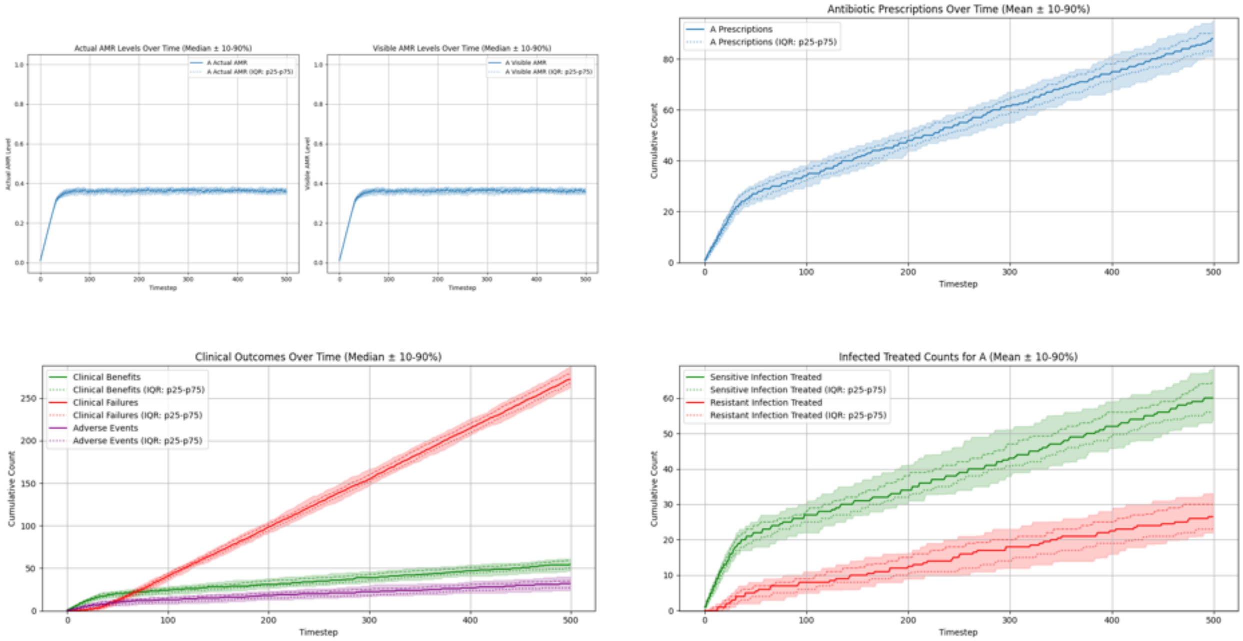

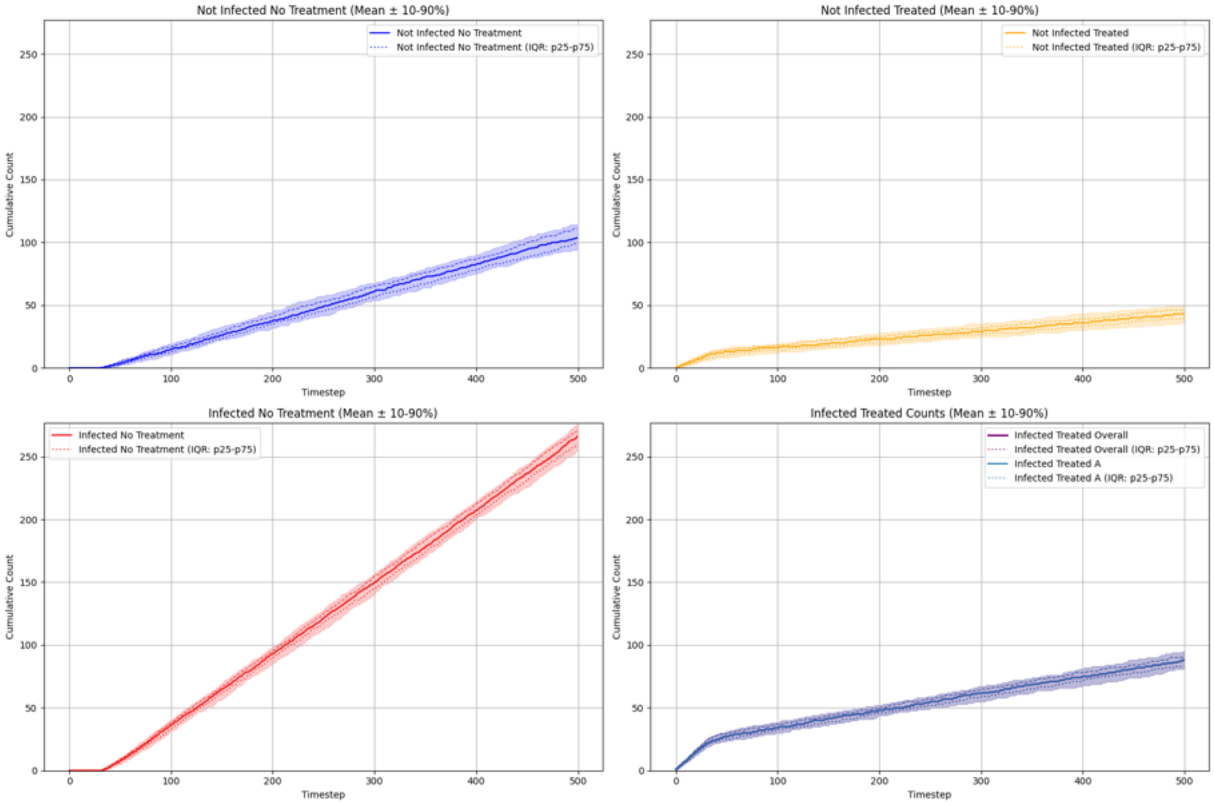

#### Summary Statistics (Single Antibiotic, Accurate Risk Stratification, Expected Reward Greedy)

| metric | p10 | p25 | p50 | p75 | p90 |
|---|---|---|---|---|---|
| overall_total_reward | -287.440 | -278.050 | -269.800 | -260.750 | -251.100 |
| overall_count_clinical_benefits | 46.900 | 50.000 | 54.500 | 59.000 | 61.000 |
| overall_count_clinical_failures | 262.000 | 267.750 | 272.000 | 279.000 | 287.000 |
| overall_count_adverse_events | 23.900 | 27.000 | 32.000 | 35.000 | 40.000 |
| overall_not_infected_no_treatment_count | 94.000 | 99.750 | 103.500 | 111.000 | 114.200 |
| overall_not_infected_treated_count | 35.900 | 40.000 | 43.000 | 46.250 | 50.100 |
| overall_infected_no_treatment_count | 254.900 | 259.750 | 266.000 | 270.250 | 276.000 |
| overall_infected_treated_count | 80.900 | 83.000 | 88.000 | 90.250 | 95.100 |
| overall_sensitive_infection_treated_count_per_abx_dict_A | 53.900 | 56.000 | 60.000 | 64.250 | 68.100 |
| overall_resistant_infection_treated_count_per_abx_dict_A | 22.000 | 23.000 | 26.500 | 30.000 | 33.100 |
| overall_sensitive_infection_treated_count | 53.900 | 56.000 | 60.000 | 64.250 | 68.100 |
| overall_resistant_infection_treated_count | 22.000 | 23.000 | 26.500 | 30.000 | 33.100 |
| overall_abx_prescriptions_count_per_abx_A | 80.900 | 83.000 | 88.000 | 90.250 | 95.100 |
| overall_abx_prescriptions_count | 80.900 | 83.000 | 88.000 | 90.250 | 95.100 |
| final_amr_actual_A | 0.347 | 0.355 | 0.363 | 0.368 | 0.376 |
| final_amr_visible_A | 0.347 | 0.355 | 0.363 | 0.368 | 0.376 |

##### Single Antibiotic, Accurate Risk Stratification, Expected Reward Lowest AMR

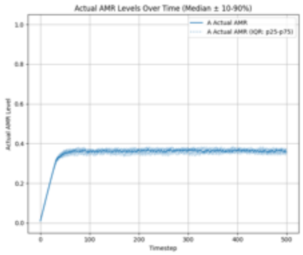

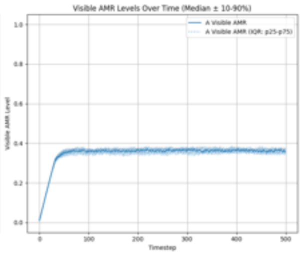

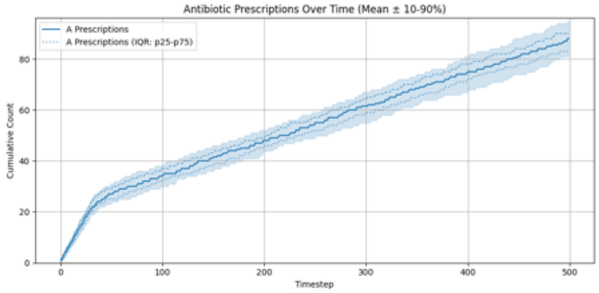

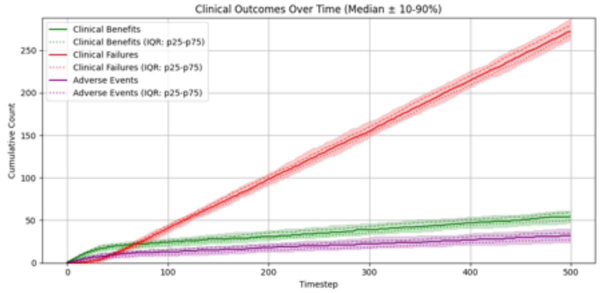

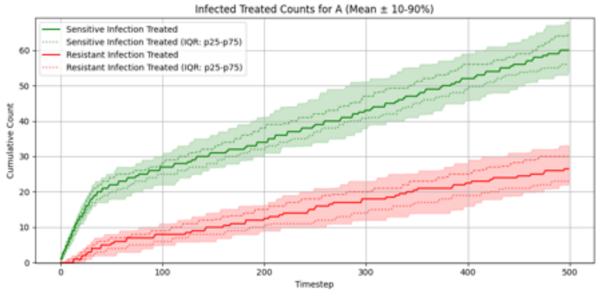

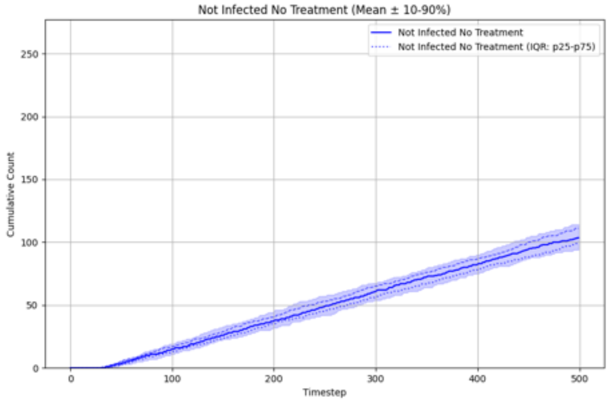

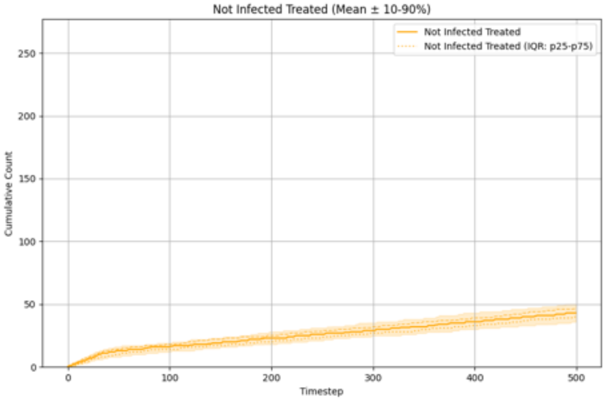

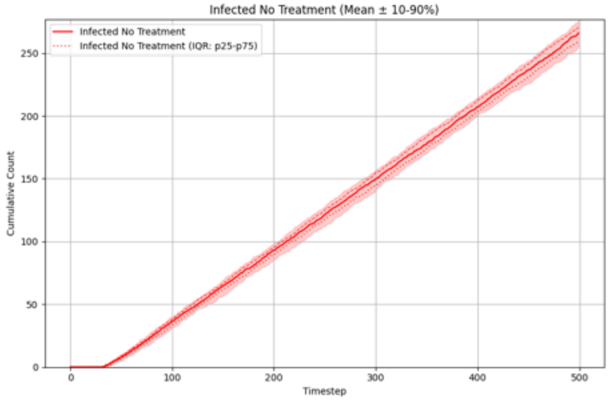

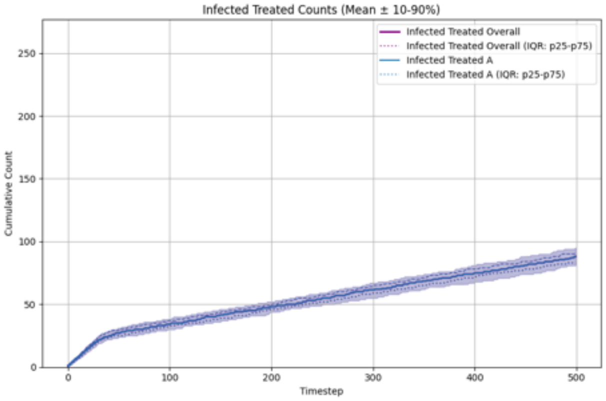

##### Summary Statistics (Single Antibiotic, Accurate Risk Stratification, Expected Reward Lowest AMR)

| metric | p10 | p25 | p50 | p75 | p90 |
|---|---|---|---|---|---|
| overall__total__reward | -287.440 | -278.050 | -269.800 | -260.750 | -251.100 |
| overall__count__clinical__benefits | 46.900 | 50.000 | 54.500 | 59.000 | 61.000 |
| overall__count__clinical__failures | 262.000 | 267.750 | 272.000 | 279.000 | 287.000 |
| overall__count__adverse__events | 23.900 | 27.000 | 32.000 | 35.000 | 40.000 |
| overall__not__infected__no__treatment__count | 94.000 | 99.750 | 103.500 | 111.000 | 114.200 |
| overall__not__infected__treated__count | 35.900 | 40.000 | 43.000 | 46.250 | 50.100 |
| overall__infected__no__treatment__count | 254.900 | 259.750 | 266.000 | 270.250 | 276.000 |
| overall__infected__treated__count | 80.900 | 83.000 | 88.000 | 90.250 | 95.100 |
| overall__sensitive__infection__treated__count__per__abx__dict__A | 53.900 | 56.000 | 60.000 | 64.250 | 68.100 |
| overall__resistant__infection__treated__count__per__abx__dict__A | 22.000 | 23.000 | 26.500 | 30.000 | 33.100 |
| overall__sensitive__infection__treated__count | 53.900 | 56.000 | 60.000 | 64.250 | 68.100 |
| overall__resistant__infection__treated__count | 22.000 | 23.000 | 26.500 | 30.000 | 33.100 |
| overall__abx__prescriptions__count__per__abx__A | 80.900 | 83.000 | 88.000 | 90.250 | 95.100 |
| overall__abx__prescriptions__count | 80.900 | 83.000 | 88.000 | 90.250 | 95.100 |
| final__amr__actual__A | 0.347 | 0.355 | 0.363 | 0.368 | 0.376 |
| final__amr__visible__A | 0.347 | 0.355 | 0.363 | 0.368 | 0.376 |

#### Single Antibiotic, Exaggerated Risk Stratification, Expected Reward Greedy

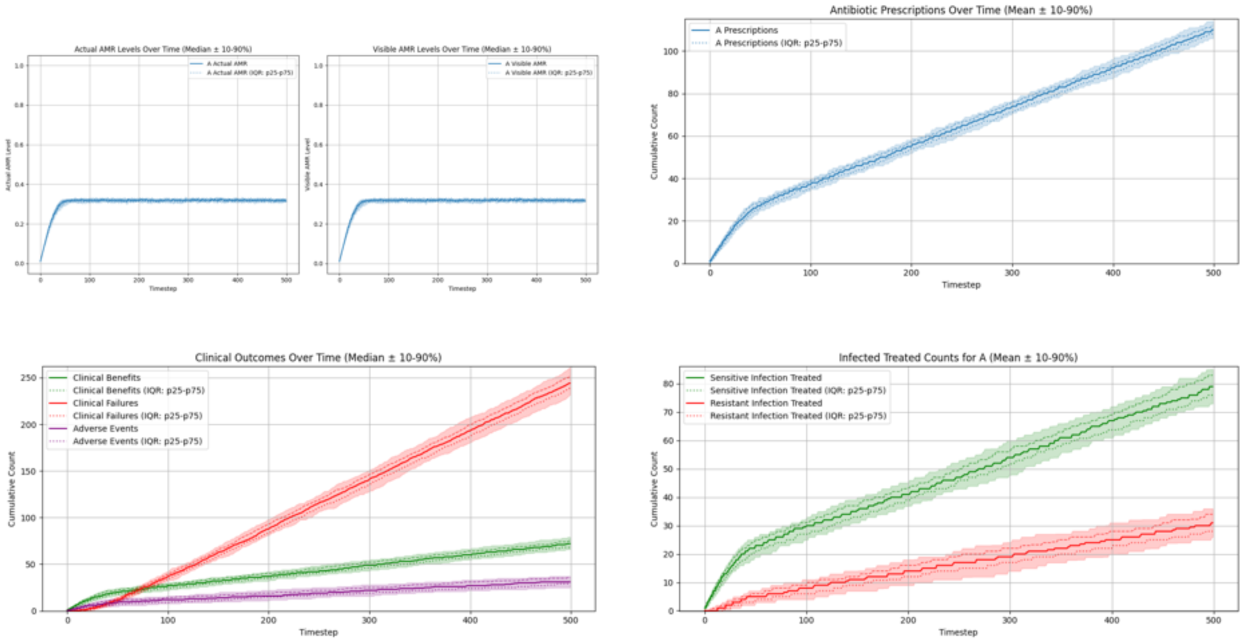

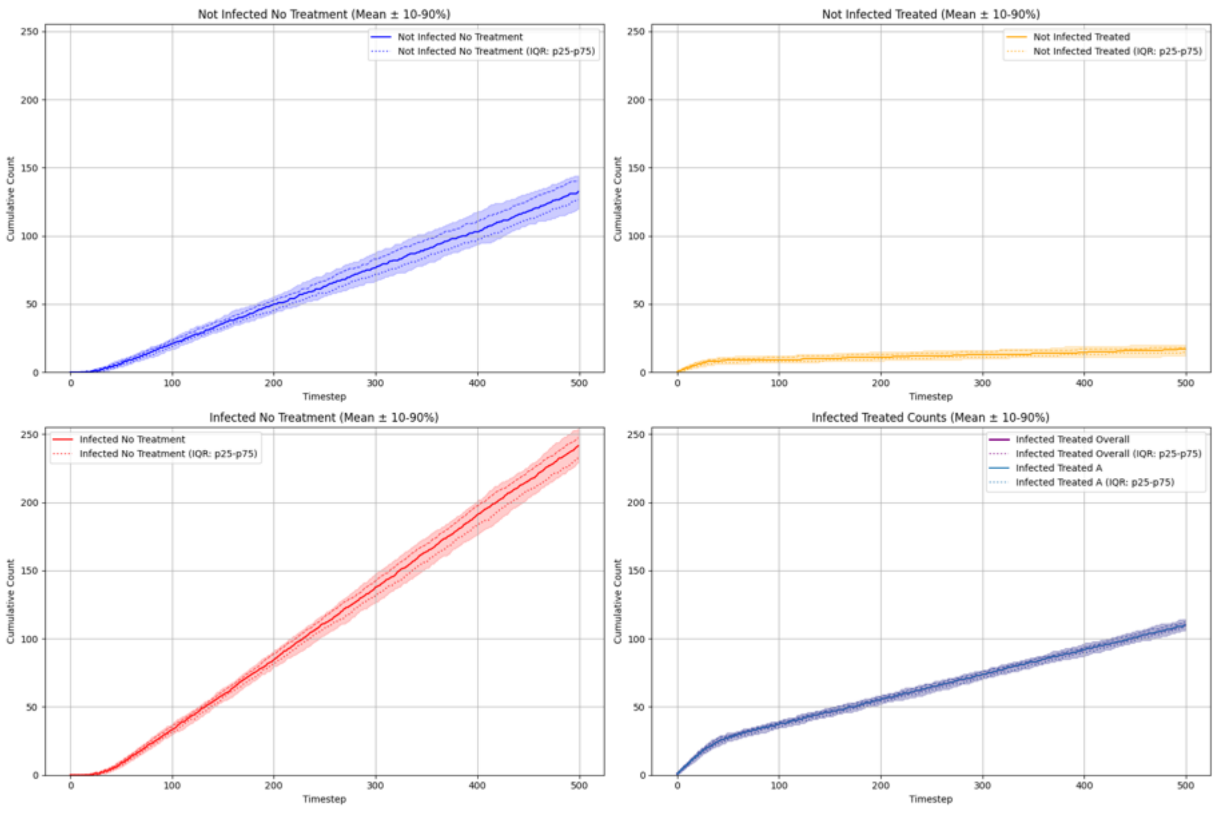

#### Summary Statistics (Single Antibiotic, Exaggerated Risk Stratification, Expected Reward Greedy)

| metric | p10 | p25 | p50 | p75 | p90 |
|---|---|---|---|---|---|
| overall_total_reward | -223.780 | -214.050 | -205.800 | -195.100 | -184.920 |
| overall_count_clinical_benefits | 66.000 | 68.000 | 72.000 | 75.000 | 79.000 |
| overall_count_clinical_failures | 231.700 | 238.750 | 244.000 | 251.250 | 261.000 |
| overall_count_adverse_events | 25.000 | 29.000 | 31.000 | 36.000 | 37.000 |
| overall_not_infected_no_treatment_count | 119.700 | 126.500 | 132.500 | 140.250 | 144.100 |
| overall_not_infected_treated_count | 12.000 | 14.750 | 17.000 | 18.000 | 20.000 |
| overall_infected_no_treatment_count | 229.000 | 233.000 | 241.500 | 247.250 | 254.300 |
| overall_infected_treated_count | 106.000 | 108.000 | 110.000 | 112.000 | 114.100 |
| overall_sensitive_infection_treated_count_per_abx_dict_A | 73.000 | 76.000 | 79.000 | 83.000 | 85.100 |
| overall_resistant_infection_treated_count_per_abx_dict_A | 26.000 | 28.000 | 31.000 | 34.000 | 36.000 |
| overall_sensitive_infection_treated_count | 73.000 | 76.000 | 79.000 | 83.000 | 85.100 |
| overall_resistant_infection_treated_count | 26.000 | 28.000 | 31.000 | 34.000 | 36.000 |
| overall_abx_prescriptions_count_per_abx_A | 106.000 | 108.000 | 110.000 | 112.000 | 114.100 |
| overall_abx_prescriptions_count | 106.000 | 108.000 | 110.000 | 112.000 | 114.100 |
| final_amr_actual_A | 0.312 | 0.315 | 0.319 | 0.322 | 0.325 |
| final_amr_visible_A | 0.312 | 0.315 | 0.319 | 0.322 | 0.325 |

**Single Antibiotic, Exaggerated Risk Stratification, Expected Reward Lowest AMR**

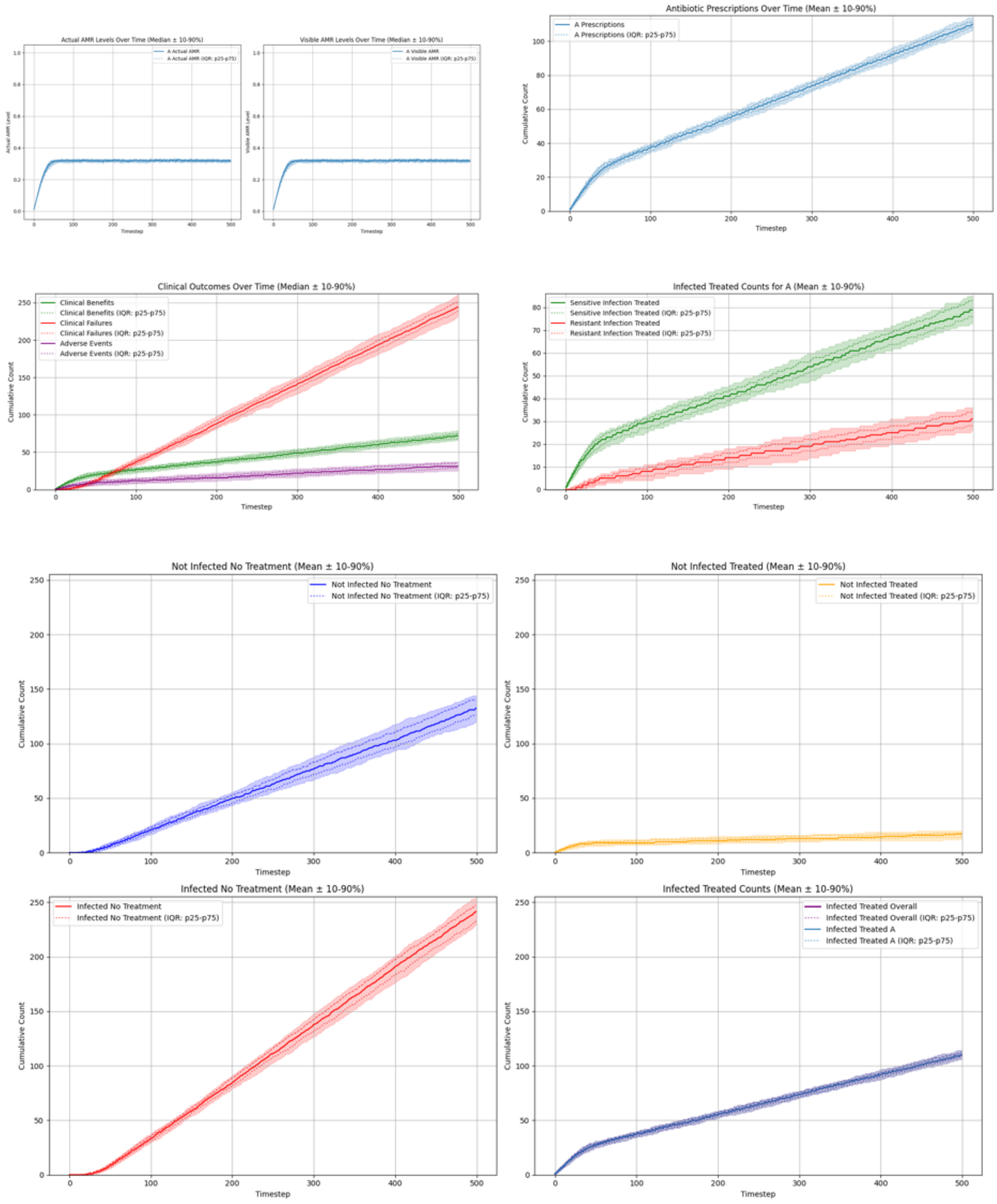

**Summary Statistics (Single Antibiotic, Exaggerated Risk Stratification, Expected Reward Lowest AMR)**

| metric | p10 | p25 | p50 | p75 | p90 |
|---|---|---|---|---|---|
| overall_total_reward | -223.780 | -214.050 | -205.800 | -195.100 | -184.920 |
| overall_count_clinical_benefits | 66.000 | 68.000 | 72.000 | 75.000 | 79.000 |
| overall_count_clinical_failures | 231.700 | 238.750 | 244.000 | 251.250 | 261.000 |
| overall_count_adverse_events | 25.000 | 29.000 | 31.000 | 36.000 | 37.000 |
| overall_not_infected_no_treatment_count | 119.700 | 126.500 | 132.500 | 140.250 | 144.100 |
| overall_not_infected_treated_count | 12.000 | 14.750 | 17.000 | 18.000 | 20.000 |
| overall_infected_no_treatment_count | 229.000 | 233.000 | 241.500 | 247.250 | 254.300 |
| overall_infected_treated_count | 106.000 | 108.000 | 110.000 | 112.000 | 114.100 |
| overall_sensitive_infection_treated_count_per_abx_dict_A | 73.000 | 76.000 | 79.000 | 83.000 | 85.100 |
| overall_resistant_infection_treated_count_per_abx_dict_A | 26.000 | 28.000 | 31.000 | 34.000 | 36.000 |
| overall_sensitive_infection_treated_count | 73.000 | 76.000 | 79.000 | 83.000 | 85.100 |
| overall_resistant_infection_treated_count | 26.000 | 28.000 | 31.000 | 34.000 | 36.000 |
| overall_abx_prescriptions_count_per_abx_A | 106.000 | 108.000 | 110.000 | 112.000 | 114.100 |
| overall_abx_prescriptions_count | 106.000 | 108.000 | 110.000 | 112.000 | 114.100 |
| final_amr_actual_A | 0.312 | 0.315 | 0.319 | 0.322 | 0.325 |
| final_amr_visible_A | 0.312 | 0.315 | 0.319 | 0.322 | 0.325 |

##### Single Antibiotic, Compressed Risk Stratification, Expected Reward Greedy

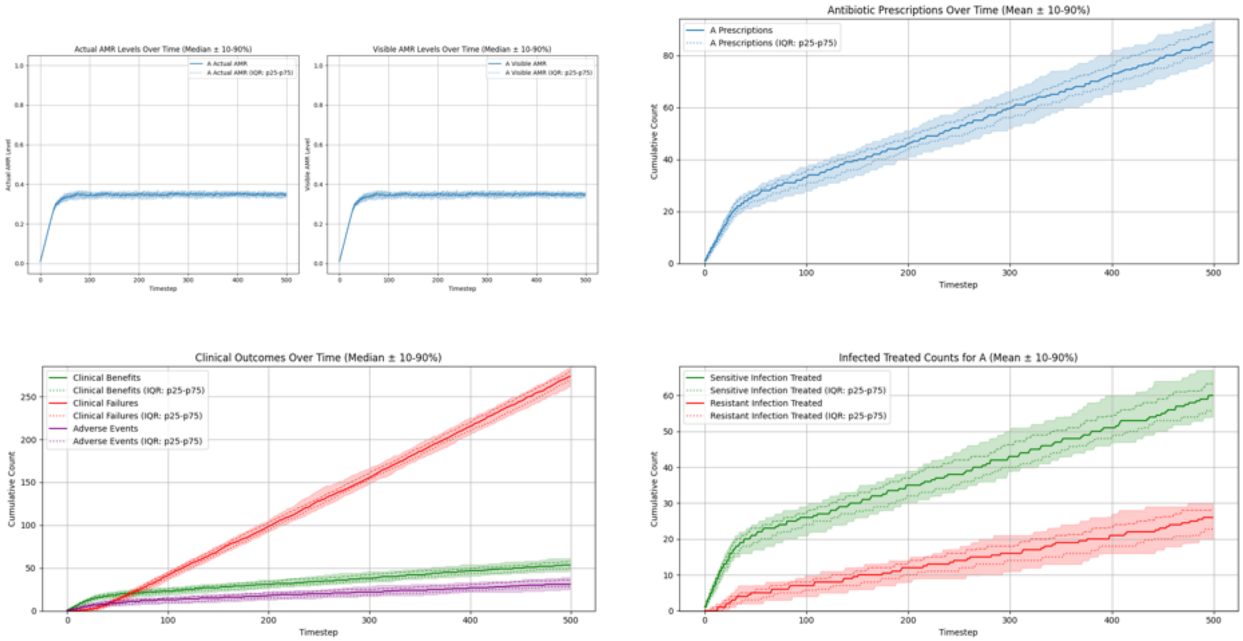

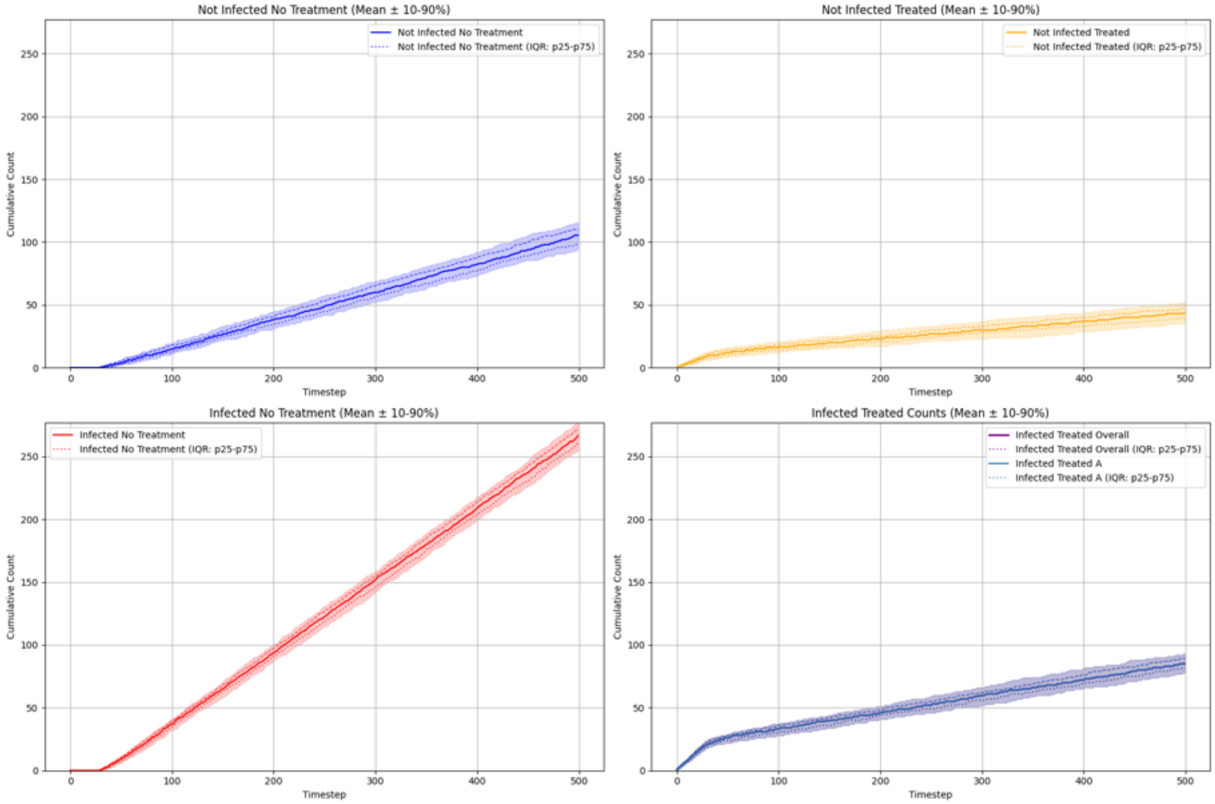

#### Summary Statistics (Single Antibiotic, Compressed Risk Stratification, Expected Reward Greedy)

| metric | p10 | p25 | p50 | p75 | p90 |
|---|---|---|---|---|---|
| overall_total_reward | -287.420 | -278.850 | -270.800 | -261.750 | -253.060 |
| overall_count_clinical_benefits | 46.900 | 50.000 | 53.500 | 58.000 | 61.100 |
| overall_count_clinical_failures | 262.000 | 269.750 | 273.500 | 279.250 | 284.100 |
| overall_count_adverse_events | 25.000 | 28.000 | 31.000 | 36.000 | 39.100 |
| overall_not_infected_no_treatment_count | 94.800 | 98.750 | 105.500 | 110.000 | 116.000 |
| overall_not_infected_treated_count | 35.000 | 40.000 | 43.500 | 47.000 | 52.100 |
| overall_infected_no_treatment_count | 255.000 | 261.000 | 266.500 | 272.000 | 276.100 |
| overall_infected_treated_count | 77.800 | 81.750 | 85.000 | 89.250 | 93.000 |
| overall_sensitive_infection_treated_count_per_abx_dict_A | 54.000 | 56.000 | 60.000 | 63.250 | 67.100 |
| overall_resistant_infection_treated_count_per_abx_dict_A | 19.900 | 22.750 | 26.000 | 28.250 | 30.000 |
| overall_sensitive_infection_treated_count | 54.000 | 56.000 | 60.000 | 63.250 | 67.100 |
| overall_resistant_infection_treated_count | 19.900 | 22.750 | 26.000 | 28.250 | 30.000 |
| overall_abx_prescriptions_count_per_abx_A | 77.800 | 81.750 | 85.000 | 89.250 | 93.000 |
| overall_abx_prescriptions_count | 77.800 | 81.750 | 85.000 | 89.250 | 93.000 |
| final_amr_actual_A | 0.334 | 0.339 | 0.347 | 0.354 | 0.361 |
| final_amr_visible_A | 0.334 | 0.339 | 0.347 | 0.354 | 0.361 |

##### Single Antibiotic, Compressed Risk Stratification, Expected Reward Lowest AMR

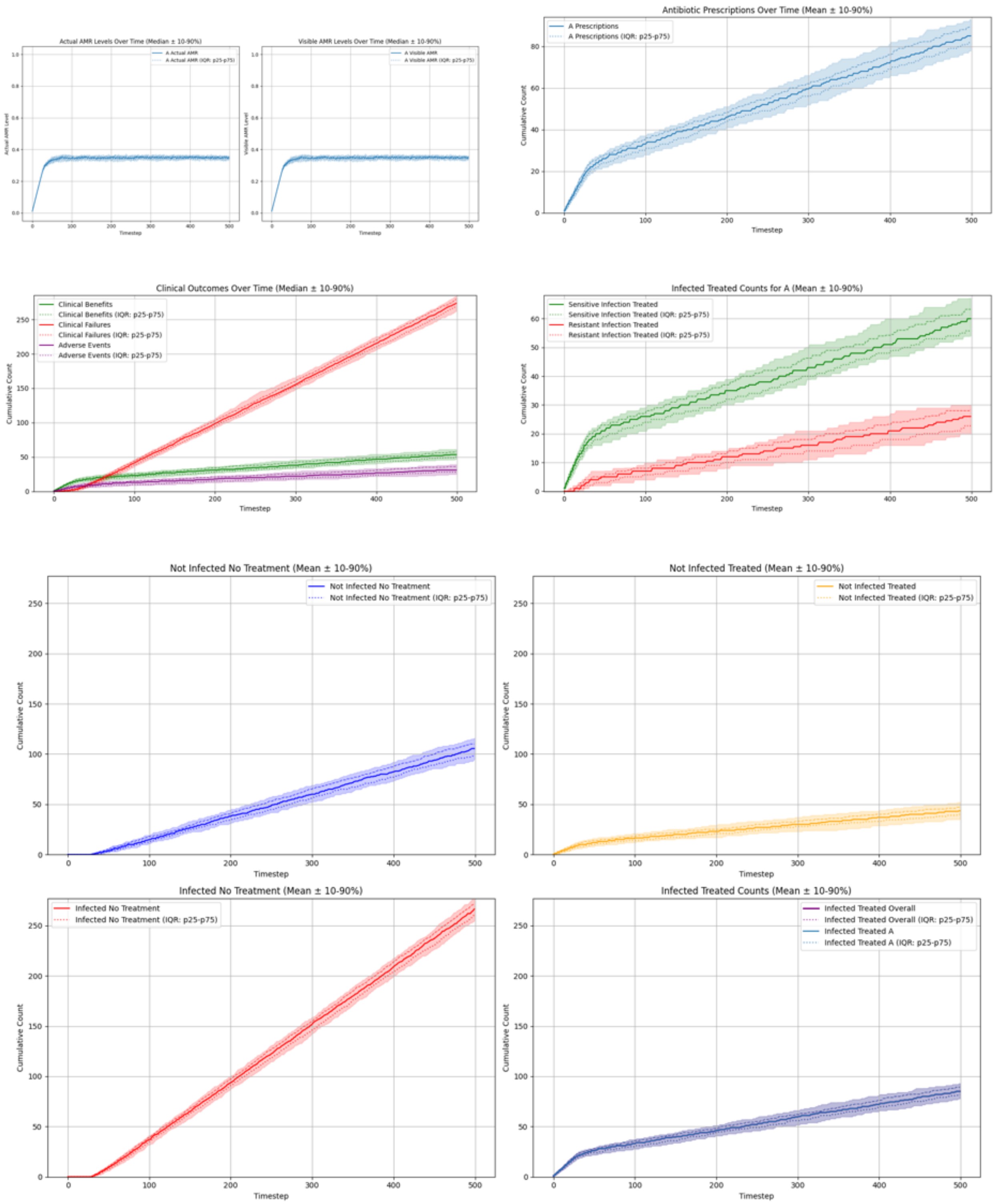

##### Summary Statistics (Single Antibiotic, Compressed Risk Stratification, Expected Reward Lowest AMR)

| metric | p10 | p25 | p50 | p75 | p90 |
|---|---|---|---|---|---|
| overall_total_reward | -287.420 | -278.850 | -270.800 | -261.750 | -253.060 |
| overall_count_clinical_benefits | 46.900 | 50.000 | 53.500 | 58.000 | 61.100 |
| overall_count_clinical_failures | 262.000 | 269.750 | 273.500 | 279.250 | 284.100 |
| overall_count_adverse_events | 25.000 | 28.000 | 31.000 | 36.000 | 39.100 |
| overall_not_infected_no_treatment_count | 94.800 | 98.750 | 105.500 | 110.000 | 116.000 |
| overall_not_infected_treated_count | 35.000 | 40.000 | 43.500 | 47.000 | 52.100 |
| overall_infected_no_treatment_count | 255.000 | 261.000 | 266.500 | 272.000 | 276.100 |
| overall_infected_treated_count | 77.800 | 81.750 | 85.000 | 89.250 | 93.000 |
| overall_sensitive_infection_treated_count_per_abx_dict_A | 54.000 | 56.000 | 60.000 | 63.250 | 67.100 |
| overall_resistant_infection_treated_count_per_abx_dict_A | 19.900 | 22.750 | 26.000 | 28.250 | 30.000 |
| overall_sensitive_infection_treated_count | 54.000 | 56.000 | 60.000 | 63.250 | 67.100 |
| overall_resistant_infection_treated_count | 19.900 | 22.750 | 26.000 | 28.250 | 30.000 |
| overall_abx_prescriptions_count_per_abx_A | 77.800 | 81.750 | 85.000 | 89.250 | 93.000 |
| overall_abx_prescriptions_count | 77.800 | 81.750 | 85.000 | 89.250 | 93.000 |
| final_amr_actual_A | 0.334 | 0.339 | 0.347 | 0.354 | 0.361 |
| final_amr_visible_A | 0.334 | 0.339 | 0.347 | 0.354 | 0.361 |

##### Two Antibiotics No Cross-Resistance, Accurate Risk Stratification, Expected Reward Greedy

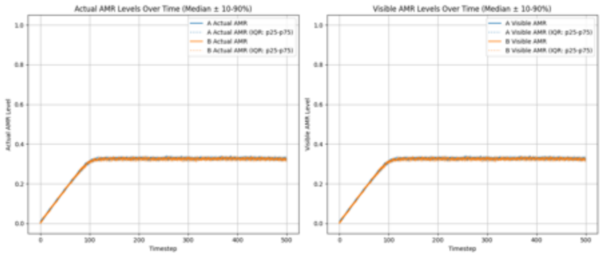

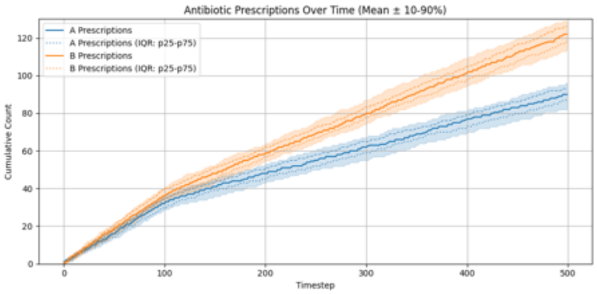

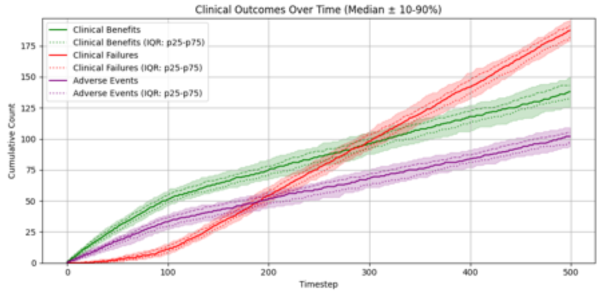

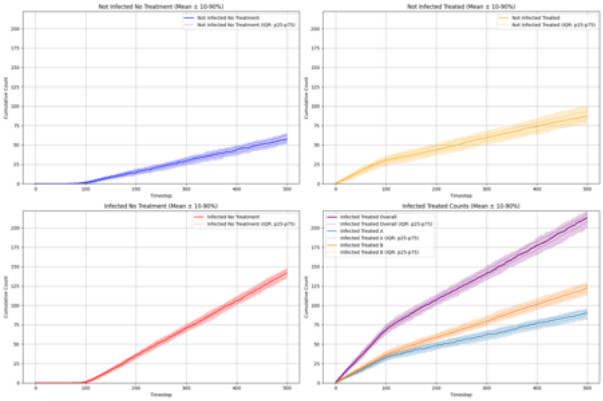

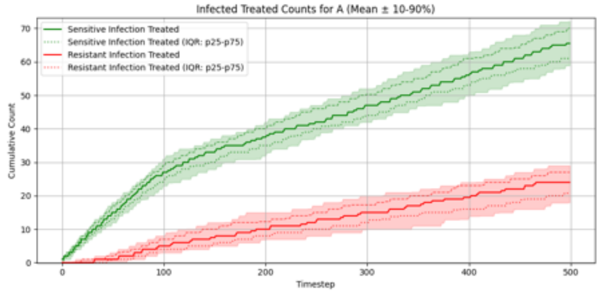

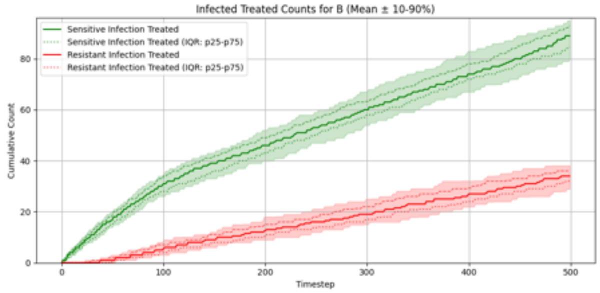

##### Summary Statistics (Two Antibiotics No Cross-Resistance, Accurate Risk Stratification, Expected Reward Greedy)

| metric | p10 | p25 | p50 | p75 | p90 |
|---|---|---|---|---|---|
| overall_total_reward | -146.960 | -136.800 | -129.400 | -119.050 | -114.090 |
| overall_count_clinical_benefits | 125.800 | 133.000 | 138.000 | 143.000 | 150.100 |
| overall_count_clinical_failures | 178.900 | 181.750 | 187.500 | 191.250 | 196.100 |
| overall_count_adverse_events | 93.800 | 96.750 | 102.000 | 105.000 | 109.100 |
| overall_not_infected_no_treatment_count | 52.000 | 54.000 | 57.500 | 62.250 | 65.000 |
| overall_not_infected_treated_count | 78.800 | 82.750 | 87.000 | 93.250 | 100.000 |
| overall_infected_no_treatment_count | 135.000 | 138.000 | 142.000 | 145.250 | 148.000 |
| overall_infected_treated_count | 200.000 | 208.000 | 213.000 | 218.000 | 221.100 |
| overall_sensitive_infection_treated_count_per_abx_dict_A | 58.900 | 61.000 | 65.500 | 70.000 | 72.100 |
| overall_sensitive_infection_treated_count_per_abx_dict_B | 79.900 | 84.500 | 89.000 | 93.000 | 95.000 |
| overall_resistant_infection_treated_count_per_abx_dict_A | 18.000 | 20.750 | 24.000 | 27.000 | 29.000 |
| overall_resistant_infection_treated_count_per_abx_dict_B | 28.900 | 32.000 | 34.000 | 36.000 | 38.100 |
| overall_sensitive_infection_treated_count | 144.900 | 148.000 | 154.500 | 160.250 | 165.100 |
| overall_resistant_infection_treated_count | 49.900 | 52.000 | 58.500 | 63.000 | 66.100 |
| overall_abx_prescriptions_count_per_abx_A | 82.000 | 87.000 | 90.000 | 93.000 | 96.000 |
| overall_abx_prescriptions_count_per_abx_B | 113.900 | 118.000 | 122.000 | 126.250 | 129.100 |
| overall_abx_prescriptions_count | 200.000 | 208.000 | 213.000 | 218.000 | 221.100 |
| final_amr_actual_A | 0.320 | 0.322 | 0.326 | 0.330 | 0.335 |
| final_amr_visible_A | 0.320 | 0.322 | 0.326 | 0.330 | 0.335 |
| final_amr_actual_B | 0.316 | 0.318 | 0.322 | 0.328 | 0.332 |
| final_amr_visible_B | 0.316 | 0.318 | 0.322 | 0.328 | 0.332 |

##### Two Antibiotics No Cross-Resistance, Accurate Risk Stratification, Expected Reward Lowest AMR

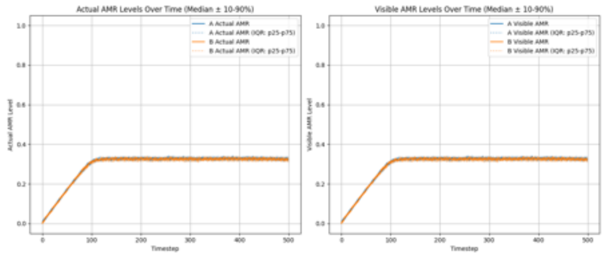

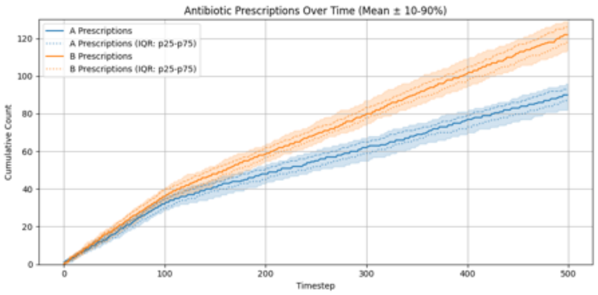

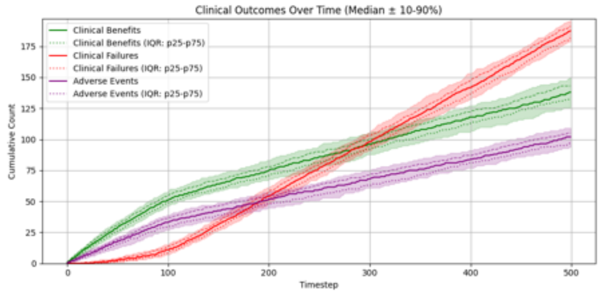

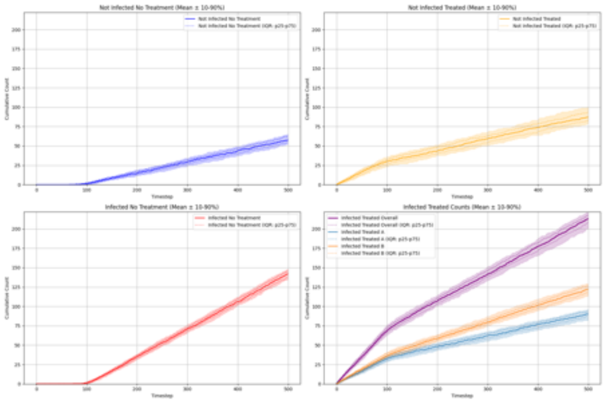

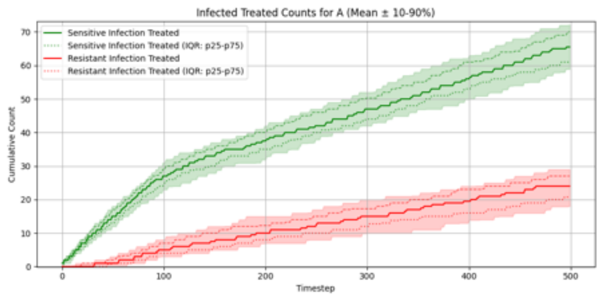

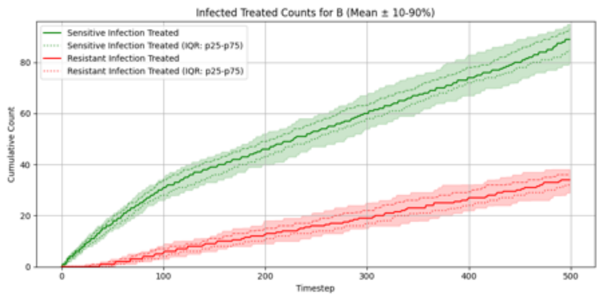

##### Summary Statistics (Two Antibiotics No Cross-Resistance, Accurate Risk Stratification, Expected Reward Lowest AMR)

| metric | p10 | p25 | p50 | p75 | p90 |
|---|---|---|---|---|---|
| overall_total_reward | -146.960 | -136.800 | -129.400 | -119.050 | -114.090 |
| overall_count_clinical_benefits | 125.800 | 133.000 | 138.000 | 143.000 | 150.100 |
| overall_count_clinical_failures | 178.900 | 181.750 | 187.500 | 191.250 | 196.100 |
| overall_count_adverse_events | 93.800 | 96.750 | 102.000 | 105.000 | 109.100 |
| overall_not_infected_no_treatment_count | 52.000 | 54.000 | 57.500 | 62.250 | 65.000 |
| overall_not_infected_treated_count | 78.800 | 82.750 | 87.000 | 93.250 | 100.000 |
| overall_infected_no_treatment_count | 135.000 | 138.000 | 142.000 | 145.250 | 148.000 |
| overall_infected_treated_count | 200.000 | 208.000 | 213.000 | 218.000 | 221.100 |
| overall_sensitive_infection_treated_count_per_abx_dict_A | 58.900 | 61.000 | 65.500 | 70.000 | 72.100 |
| overall_sensitive_infection_treated_count_per_abx_dict_B | 79.900 | 84.500 | 89.000 | 93.000 | 95.000 |
| overall_resistant_infection_treated_count_per_abx_dict_A | 18.000 | 20.750 | 24.000 | 27.000 | 29.000 |
| overall_resistant_infection_treated_count_per_abx_dict_B | 28.900 | 32.000 | 34.000 | 36.000 | 38.100 |
| overall_sensitive_infection_treated_count | 144.900 | 148.000 | 154.500 | 160.250 | 165.100 |
| overall_resistant_infection_treated_count | 49.900 | 52.000 | 58.500 | 63.000 | 66.100 |
| overall_abx_prescriptions_count_per_abx_A | 82.000 | 87.000 | 90.000 | 93.000 | 96.000 |
| overall_abx_prescriptions_count_per_abx_B | 113.900 | 118.000 | 122.000 | 126.250 | 129.100 |
| overall_abx_prescriptions_count | 200.000 | 208.000 | 213.000 | 218.000 | 221.100 |
| final_amr_actual_A | 0.320 | 0.322 | 0.326 | 0.330 | 0.335 |
| final_amr_visible_A | 0.320 | 0.322 | 0.326 | 0.330 | 0.335 |
| final_amr_actual_B | 0.316 | 0.318 | 0.322 | 0.328 | 0.332 |
| final_amr_visible_B | 0.316 | 0.318 | 0.322 | 0.328 | 0.332 |

##### Two Antibiotics No Cross-Resistance, Exaggerated Risk Stratification, Expected Reward Greedy

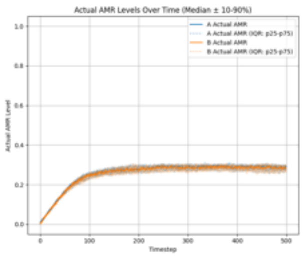

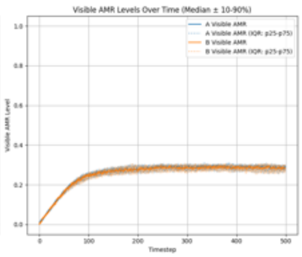

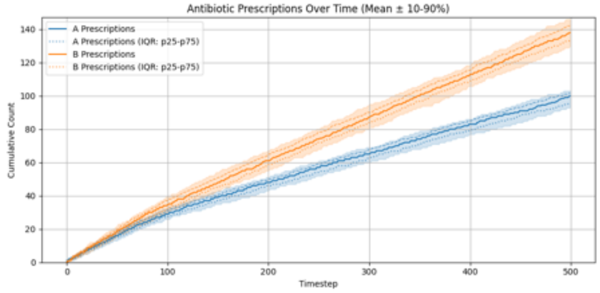

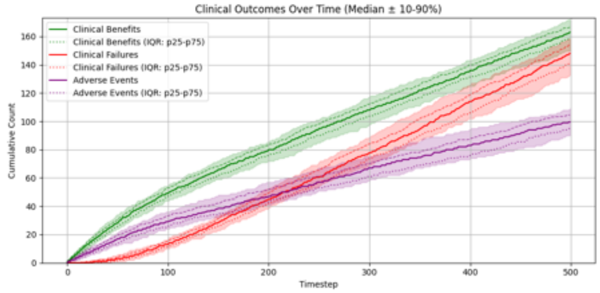

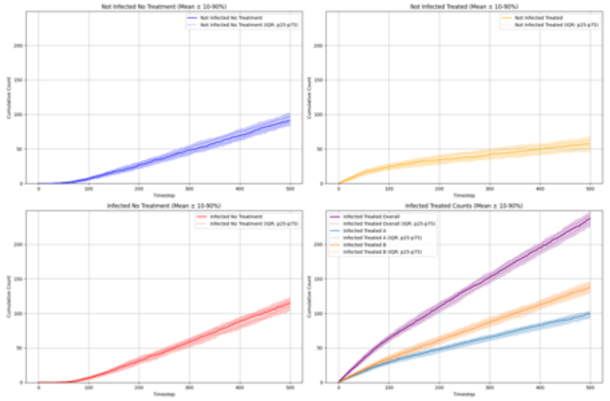

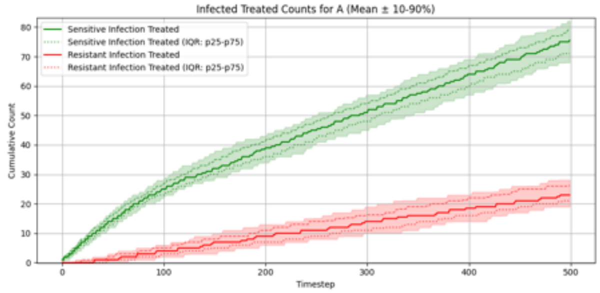

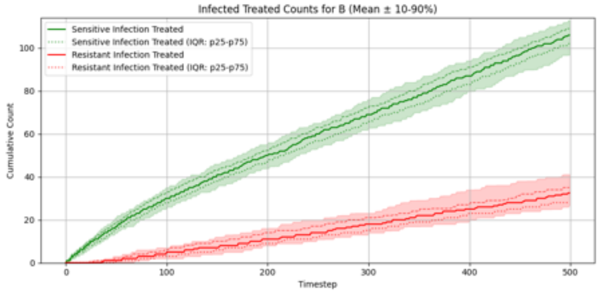

#### Summary Statistics (Two Antibiotics No Cross-Resistance, Exaggerated Risk Stratification, Expected Reward Greedy)

| metric | p10 | p25 | p50 | p75 | p90 |
|---|---|---|---|---|---|
| overall_total_reward | -58.960 | -46.975 | -40.050 | -28.250 | -16.580 |
| overall_count_clinical_benefits | 149.900 | 158.750 | 163.000 | 166.500 | 172.100 |
| overall_count_clinical_failures | 132.900 | 141.000 | 148.000 | 155.000 | 158.400 |
| overall_count_adverse_events | 90.000 | 95.000 | 99.500 | 104.500 | 109.100 |
| overall_not_infected_no_treatment_count | 84.000 | 87.000 | 91.000 | 97.000 | 102.100 |
| overall_not_infected_treated_count | 46.900 | 51.750 | 57.000 | 62.000 | 69.000 |
| overall_infected_no_treatment_count | 104.000 | 109.750 | 115.000 | 118.250 | 123.000 |
| overall_infected_treated_count | 225.000 | 231.000 | 238.000 | 242.000 | 248.100 |
| overall_sensitive_infection_treated_count_per_abx_dict_A | 68.000 | 71.000 | 75.500 | 79.000 | 82.100 |
| overall_sensitive_infection_treated_count_per_abx_dict_B | 96.800 | 102.000 | 106.000 | 109.000 | 113.000 |
| overall_resistant_infection_treated_count_per_abx_dict_A | 19.000 | 21.000 | 23.000 | 26.250 | 28.100 |
| overall_resistant_infection_treated_count_per_abx_dict_B | 26.900 | 28.000 | 32.500 | 35.000 | 41.100 |
| overall_sensitive_infection_treated_count | 168.000 | 176.750 | 181.500 | 186.000 | 192.000 |
| overall_resistant_infection_treated_count | 48.000 | 51.750 | 55.000 | 60.250 | 63.100 |
| overall_abx_prescriptions_count_per_abx_A | 92.900 | 95.750 | 100.000 | 102.000 | 103.100 |
| overall_abx_prescriptions_count_per_abx_B | 129.900 | 133.000 | 138.000 | 142.000 | 146.100 |
| overall_abx_prescriptions_count | 225.000 | 231.000 | 238.000 | 242.000 | 248.100 |
| final_amr_actual_A | 0.265 | 0.279 | 0.287 | 0.295 | 0.299 |
| final_amr_visible_A | 0.265 | 0.279 | 0.287 | 0.295 | 0.299 |
| final_amr_actual_B | 0.264 | 0.275 | 0.281 | 0.293 | 0.298 |
| final_amr_visible_B | 0.264 | 0.275 | 0.281 | 0.293 | 0.298 |

##### Two Antibiotics No Cross-Resistance, Exaggerated Risk Stratification, Expected Reward Lowest AMR

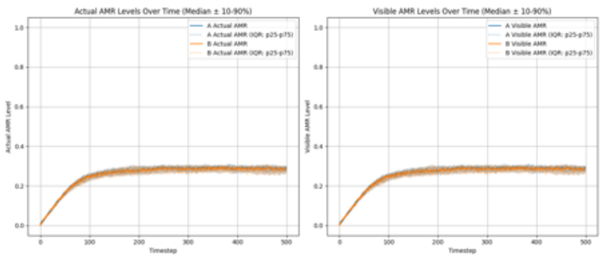

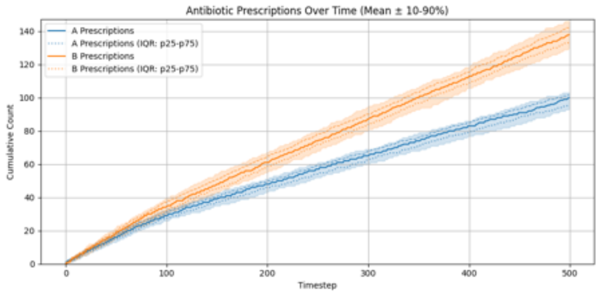

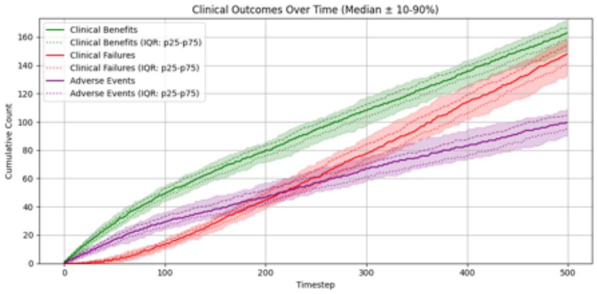

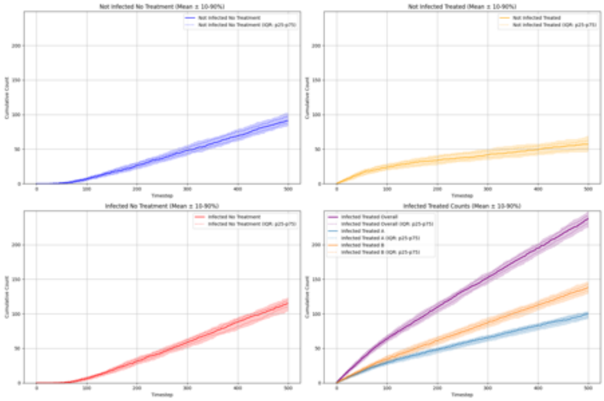

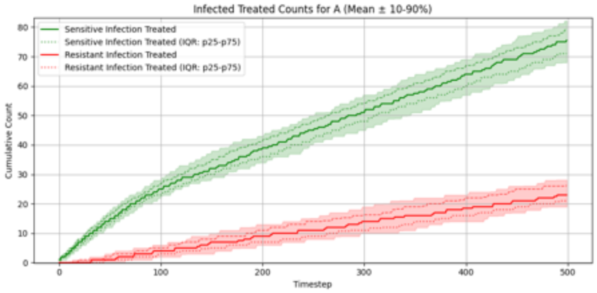

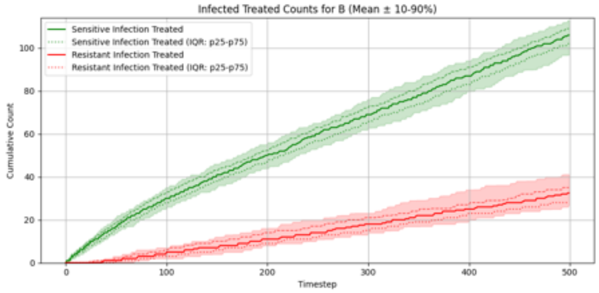

##### Summary Statistics (Two Antibiotics No Cross-Resistance, Exaggerated Risk Stratification, Expected Reward Lowest AMR)

| metric | p10 | p25 | p50 | p75 | p90 |
|---|---|---|---|---|---|
| overall_total_reward | -58.960 | -46.975 | -40.050 | -28.250 | -16.580 |
| overall_count_clinical_benefits | 149.900 | 158.750 | 163.000 | 166.500 | 172.100 |
| overall_count_clinical_failures | 132.900 | 141.000 | 148.000 | 155.000 | 158.400 |
| overall_count_adverse_events | 90.000 | 95.000 | 99.500 | 104.500 | 109.100 |
| overall_not_infected_no_treatment_count | 84.000 | 87.000 | 91.000 | 97.000 | 102.100 |
| overall_not_infected_treated_count | 46.900 | 51.750 | 57.000 | 62.000 | 69.000 |
| overall_infected_no_treatment_count | 104.000 | 109.750 | 115.000 | 118.250 | 123.000 |
| overall_infected_treated_count | 225.000 | 231.000 | 238.000 | 242.000 | 248.100 |
| overall_sensitive_infection_treated_count_per_abx_dict_A | 68.000 | 71.000 | 75.500 | 79.000 | 82.100 |
| overall_sensitive_infection_treated_count_per_abx_dict_B | 96.800 | 102.000 | 106.000 | 109.000 | 113.000 |
| overall_resistant_infection_treated_count_per_abx_dict_A | 19.000 | 21.000 | 23.000 | 26.250 | 28.100 |
| overall_resistant_infection_treated_count_per_abx_dict_B | 26.900 | 28.000 | 32.500 | 35.000 | 41.100 |
| overall_sensitive_infection_treated_count | 168.000 | 176.750 | 181.500 | 186.000 | 192.000 |
| overall_resistant_infection_treated_count | 48.000 | 51.750 | 55.000 | 60.250 | 63.100 |
| overall_abx_prescriptions_count_per_abx_A | 92.900 | 95.750 | 100.000 | 102.000 | 103.100 |
| overall_abx_prescriptions_count_per_abx_B | 129.900 | 133.000 | 138.000 | 142.000 | 146.100 |
| overall_abx_prescriptions_count | 225.000 | 231.000 | 238.000 | 242.000 | 248.100 |
| final_amr_actual_A | 0.265 | 0.279 | 0.287 | 0.295 | 0.299 |
| final_amr_visible_A | 0.265 | 0.279 | 0.287 | 0.295 | 0.299 |
| final_amr_actual_B | 0.264 | 0.275 | 0.281 | 0.293 | 0.298 |
| final_amr_visible_B | 0.264 | 0.275 | 0.281 | 0.293 | 0.298 |

#### Two Antibiotics No Cross-Resistance, Compressed Risk Stratification, Expected Reward Greedy

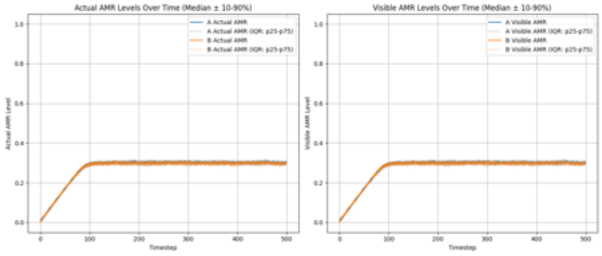

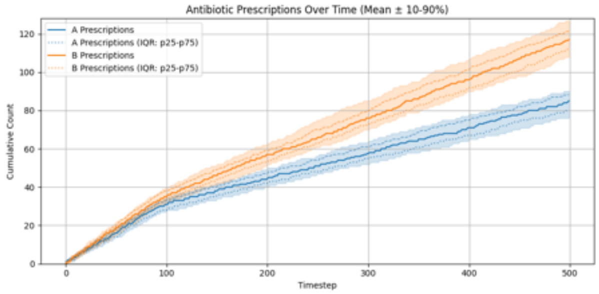

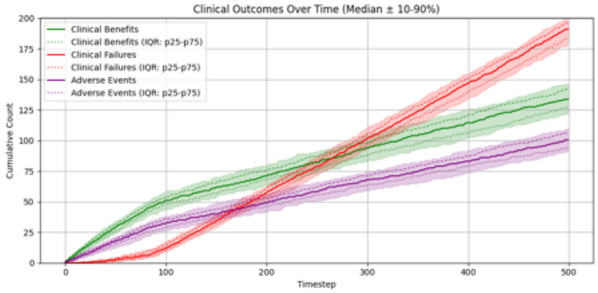

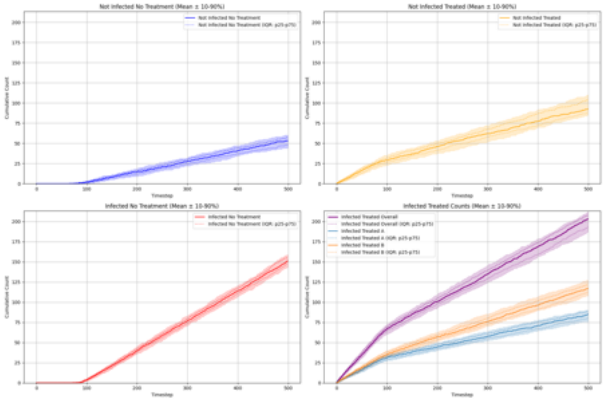

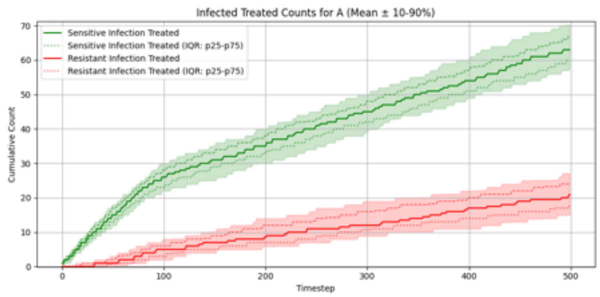

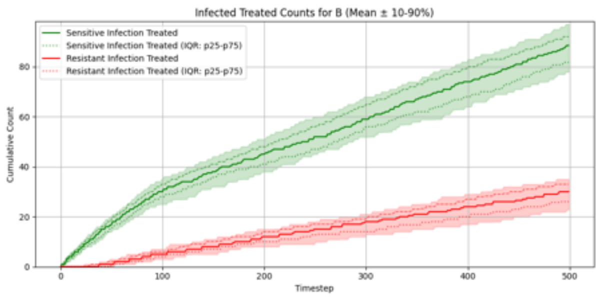

##### Summary Statistics (Two Antibiotics No Cross-Resistance, Compressed Risk Stratification, Expected Reward Greedy)

| metric | p10 | p25 | p50 | p75 | p90 |
|---|---|---|---|---|---|
| overall__total__reward | -156.390 | -145.000 | -138.150 | -129.200 | -117.650 |
| overall__count__clinical__benefits | 121.900 | 127.000 | 134.000 | 142.000 | 146.100 |
| overall__count__clinical__failures | 178.900 | 184.000 | 191.000 | 196.000 | 199.200 |
| overall__count__adverse__events | 91.000 | 94.000 | 100.500 | 106.000 | 110.000 |
| overall__not__infected__no__treatment__count | 44.000 | 48.000 | 53.000 | 57.250 | 60.200 |
| overall__not__infected__treated__count | 84.900 | 88.000 | 92.500 | 104.000 | 110.000 |
| overall__infected__no__treatment__count | 143.000 | 147.000 | 151.000 | 155.250 | 159.000 |
| overall__infected__treated__count | 186.900 | 193.000 | 203.000 | 208.000 | 212.200 |
| overall__sensitive__infection__treated__count__per__abx__dict__A | 57.900 | 60.000 | 63.000 | 67.000 | 70.200 |
| overall__sensitive__infection__treated__count__per__abx__dict__B | 77.900 | 81.750 | 88.500 | 92.000 | 97.100 |
| overall__resistant__infection__treated__count__per__abx__dict__A | 15.000 | 17.750 | 21.000 | 24.000 | 27.100 |
| overall__resistant__infection__treated__count__per__abx__dict__B | 22.800 | 26.000 | 30.000 | 33.000 | 35.000 |
| overall__sensitive__infection__treated__count | 139.000 | 144.750 | 151.500 | 157.250 | 161.100 |
| overall__resistant__infection__treated__count | 42.000 | 45.750 | 50.000 | 55.000 | 58.000 |
| overall__abx__prescriptions__count__per__abx__A | 76.000 | 80.000 | 85.000 | 88.250 | 90.100 |
| overall__abx__prescriptions__count__per__abx__B | 107.800 | 112.000 | 117.000 | 121.250 | 127.000 |
| overall__abx__prescriptions__count | 186.900 | 193.000 | 203.000 | 208.000 | 212.200 |
| final__amr__actual__A | 0.292 | 0.296 | 0.300 | 0.304 | 0.309 |
| final__amr__visible__A | 0.292 | 0.296 | 0.300 | 0.304 | 0.309 |
| final__amr__actual__B | 0.291 | 0.293 | 0.297 | 0.301 | 0.305 |
| final__amr__visible__B | 0.291 | 0.293 | 0.297 | 0.301 | 0.305 |

##### Two Antibiotics No Cross-Resistance, Compressed Risk Stratification, Expected Reward Lowest AMR

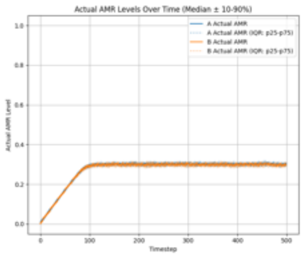

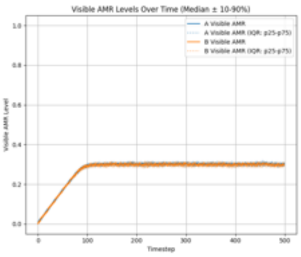

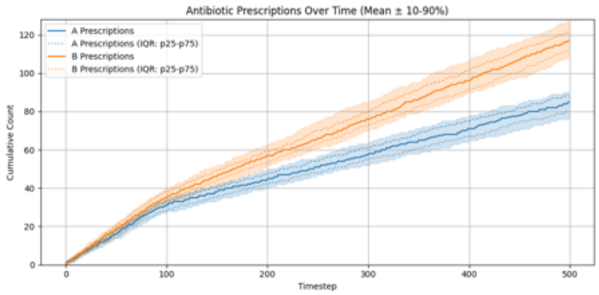

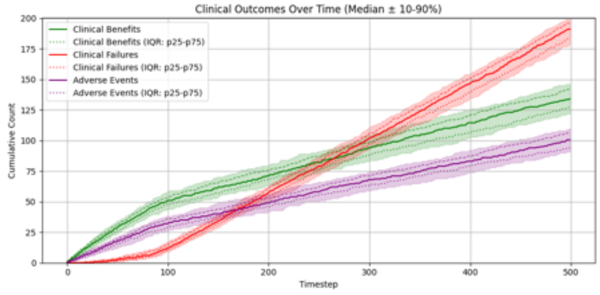

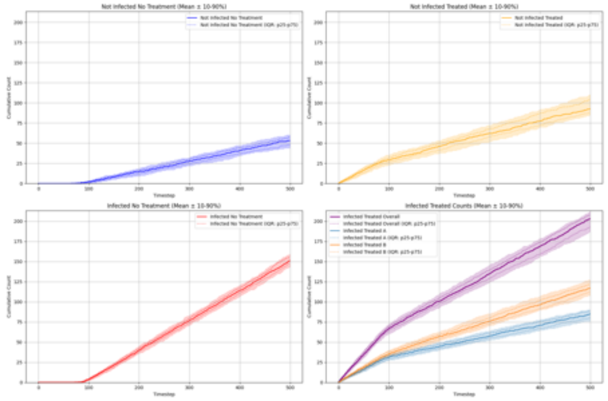

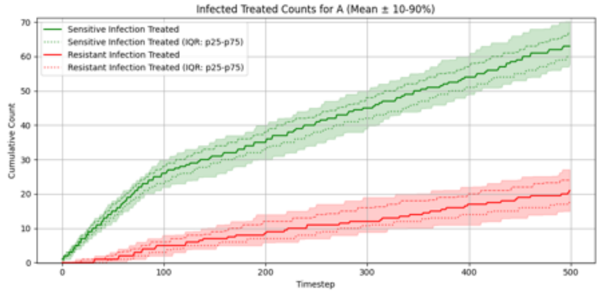

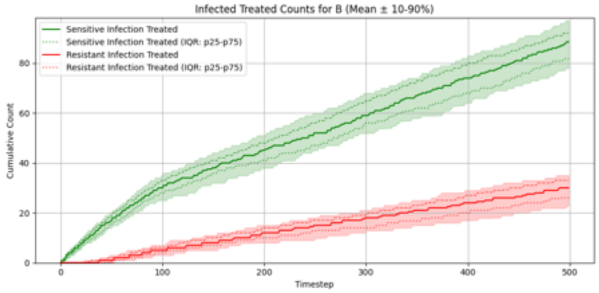

#### Summary Statistics (Two Antibiotics No Cross-Resistance, Compressed Risk Stratification, Expected Reward Lowest AMR)

| metric | p10 | p25 | p50 | p75 | p90 |
|---|---|---|---|---|---|
| overall_total_reward | -156.390 | -145.000 | -138.150 | -129.200 | -117.650 |
| overall_count_clinical_benefits | 121.900 | 127.000 | 134.000 | 142.000 | 146.100 |
| overall_count_clinical_failures | 178.900 | 184.000 | 191.000 | 196.000 | 199.200 |
| overall_count_adverse_events | 91.000 | 94.000 | 100.500 | 106.000 | 110.000 |
| overall_not_infected_no_treatment_count | 44.000 | 48.000 | 53.000 | 57.250 | 60.200 |
| overall_not_infected_treated_count | 84.900 | 88.000 | 92.500 | 104.000 | 110.000 |
| overall_infected_no_treatment_count | 143.000 | 147.000 | 151.000 | 155.250 | 159.000 |
| overall_infected_treated_count | 186.900 | 193.000 | 203.000 | 208.000 | 212.200 |
| overall_sensitive_infection_treated_count_per_abx_dict_A | 57.900 | 60.000 | 63.000 | 67.000 | 70.200 |
| overall_sensitive_infection_treated_count_per_abx_dict_B | 77.900 | 81.750 | 88.500 | 92.000 | 97.100 |
| overall_resistant_infection_treated_count_per_abx_dict_A | 15.000 | 17.750 | 21.000 | 24.000 | 27.100 |
| overall_resistant_infection_treated_count_per_abx_dict_B | 22.800 | 26.000 | 30.000 | 33.000 | 35.000 |
| overall_sensitive_infection_treated_count | 139.000 | 144.750 | 151.500 | 157.250 | 161.100 |
| overall_resistant_infection_treated_count | 42.000 | 45.750 | 50.000 | 55.000 | 58.000 |
| overall_abx_prescriptions_count_per_abx_A | 76.000 | 80.000 | 85.000 | 88.250 | 90.100 |
| overall_abx_prescriptions_count_per_abx_B | 107.800 | 112.000 | 117.000 | 121.250 | 127.000 |
| overall_abx_prescriptions_count | 186.900 | 193.000 | 203.000 | 208.000 | 212.200 |
| final_amr_actual_A | 0.292 | 0.296 | 0.300 | 0.304 | 0.309 |
| final_amr_visible_A | 0.292 | 0.296 | 0.300 | 0.304 | 0.309 |
| final_amr_actual_B | 0.291 | 0.293 | 0.297 | 0.301 | 0.305 |
| final_amr_visible_B | 0.291 | 0.293 | 0.297 | 0.301 | 0.305 |

##### Two Antibiotics With Cross-Resistance, Accurate Risk Stratification, Expected Reward Greedy

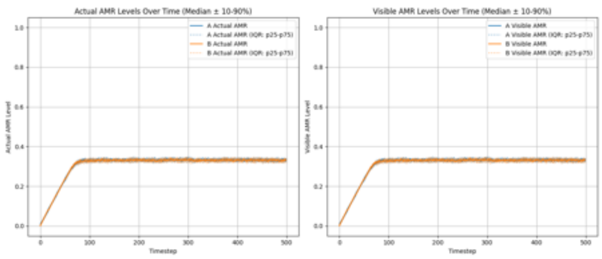

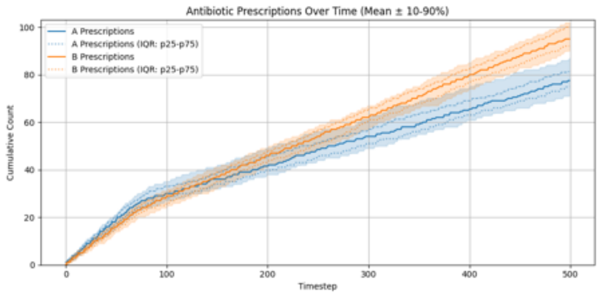

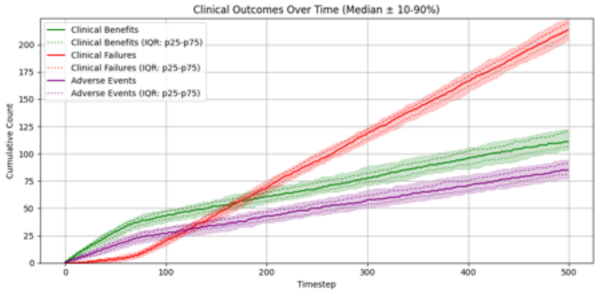

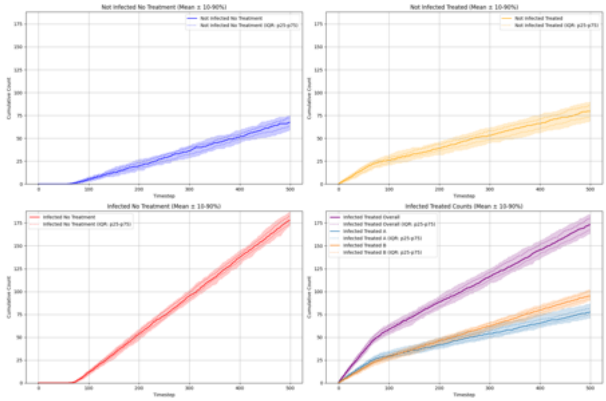

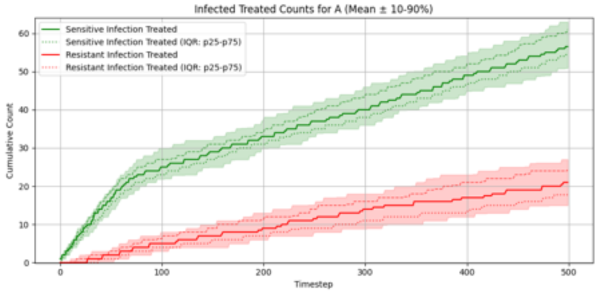

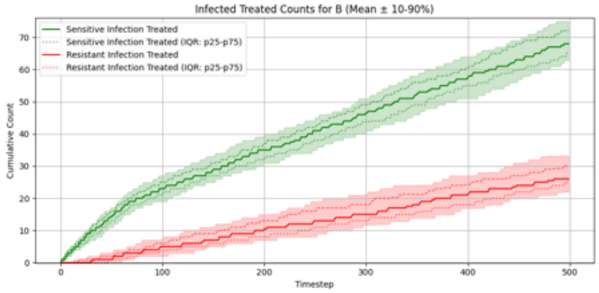

##### Summary Statistics (Two Antibiotics With Cross-Resistance, Accurate Risk Stratification, Expected Reward Greedy)

| metric | p10 | p25 | p50 | p75 | p90 |
|---|---|---|---|---|---|
| overall_total_reward | -193.880 | -183.575 | -175.700 | -167.025 | -159.970 |
| overall_count_clinical_benefits | 103.900 | 106.750 | 111.000 | 120.000 | 122.300 |
| overall_count_clinical_failures | 204.000 | 209.000 | 213.500 | 220.250 | 223.000 |
| overall_count_adverse_events | 77.000 | 81.000 | 85.000 | 91.000 | 94.100 |
| overall_not_infected_no_treatment_count | 58.900 | 63.000 | 67.500 | 72.000 | 75.000 |
| overall_not_infected_treated_count | 69.000 | 73.750 | 79.500 | 86.000 | 90.000 |
| overall_infected_no_treatment_count | 172.000 | 174.750 | 178.000 | 183.000 | 187.100 |
| overall_infected_treated_count | 163.900 | 167.750 | 173.000 | 180.000 | 184.100 |
| overall_sensitive_infection_treated_count_per_abx_dict_A | 50.900 | 54.750 | 56.500 | 61.000 | 63.000 |
| overall_sensitive_infection_treated_count_per_abx_dict_B | 62.900 | 65.000 | 68.000 | 72.000 | 75.000 |
| overall_resistant_infection_treated_count_per_abx_dict_A | 15.000 | 17.750 | 21.000 | 24.250 | 27.000 |
| overall_resistant_infection_treated_count_per_abx_dict_B | 22.000 | 25.000 | 26.000 | 30.000 | 33.200 |
| overall_sensitive_infection_treated_count | 116.900 | 120.000 | 125.000 | 131.000 | 136.100 |
| overall_resistant_infection_treated_count | 41.900 | 44.750 | 49.000 | 52.250 | 56.000 |
| overall_abx_prescriptions_count_per_abx_A | 71.000 | 75.000 | 77.500 | 81.250 | 87.000 |
| overall_abx_prescriptions_count_per_abx_B | 90.000 | 92.000 | 95.000 | 100.000 | 102.000 |
| overall_abx_prescriptions_count | 163.900 | 167.750 | 173.000 | 180.000 | 184.100 |
| final_amr_actual_A | 0.323 | 0.327 | 0.334 | 0.336 | 0.341 |
| final_amr_visible_A | 0.323 | 0.327 | 0.334 | 0.336 | 0.341 |
| final_amr_actual_B | 0.321 | 0.326 | 0.331 | 0.335 | 0.340 |
| final_amr_visible_B | 0.321 | 0.326 | 0.331 | 0.335 | 0.340 |

##### Two Antibiotics With Cross-Resistance, Accurate Risk Stratification, Expected Reward Lowest AMR

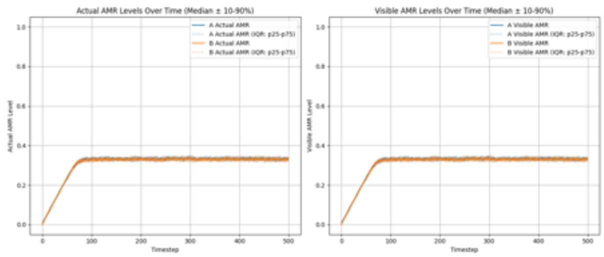

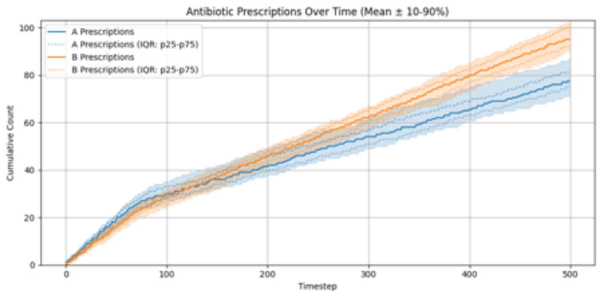

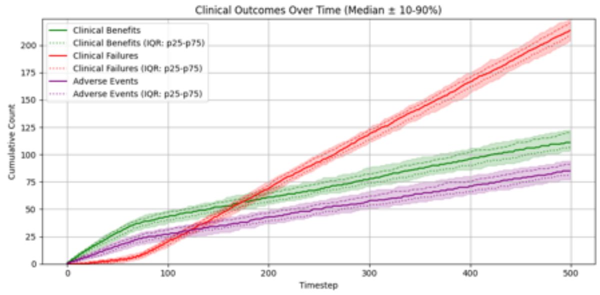

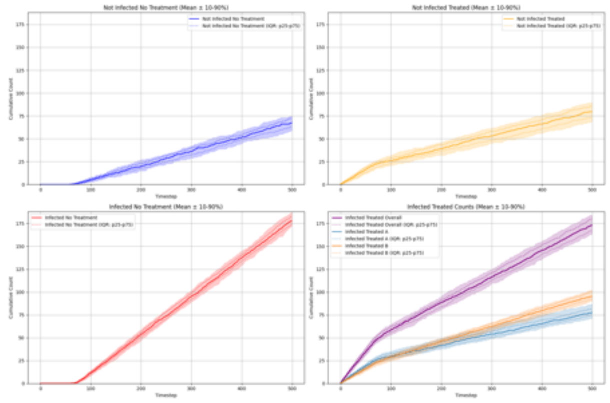

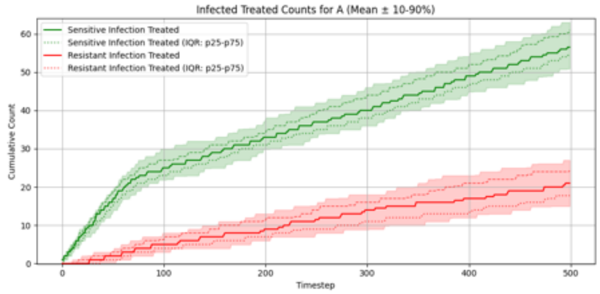

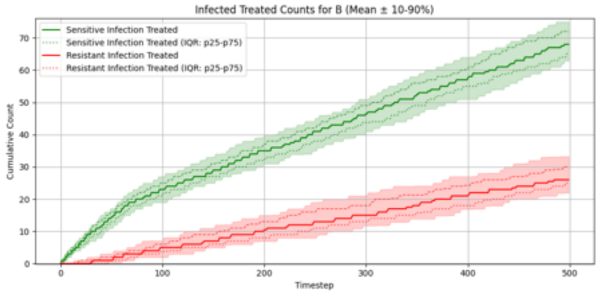

#### Summary Statistics (Two Antibiotics With Cross-Resistance, Accurate Risk Stratification, Expected Reward Lowest AMR)

| metric | p10 | p25 | p50 | p75 | p90 |
|---|---|---|---|---|---|
| overall_total_reward | -193.880 | -183.575 | -175.700 | -167.025 | -159.970 |
| overall_count_clinical_benefits | 103.900 | 106.750 | 111.000 | 120.000 | 122.300 |
| overall_count_clinical_failures | 204.000 | 209.000 | 213.500 | 220.250 | 223.000 |
| overall_count_adverse_events | 77.000 | 81.000 | 85.000 | 91.000 | 94.100 |
| overall_not_infected_no_treatment_count | 58.900 | 63.000 | 67.500 | 72.000 | 75.000 |
| overall_not_infected_treated_count | 69.000 | 73.750 | 79.500 | 86.000 | 90.000 |
| overall_infected_no_treatment_count | 172.000 | 174.750 | 178.000 | 183.000 | 187.100 |
| overall_infected_treated_count | 163.900 | 167.750 | 173.000 | 180.000 | 184.100 |
| overall_sensitive_infection_treated_count_per_abx_dict_A | 50.900 | 54.750 | 56.500 | 61.000 | 63.000 |
| overall_sensitive_infection_treated_count_per_abx_dict_B | 62.900 | 65.000 | 68.000 | 72.000 | 75.000 |
| overall_resistant_infection_treated_count_per_abx_dict_A | 15.000 | 17.750 | 21.000 | 24.250 | 27.000 |
| overall_resistant_infection_treated_count_per_abx_dict_B | 22.000 | 25.000 | 26.000 | 30.000 | 33.200 |
| overall_sensitive_infection_treated_count | 116.900 | 120.000 | 125.000 | 131.000 | 136.100 |
| overall_resistant_infection_treated_count | 41.900 | 44.750 | 49.000 | 52.250 | 56.000 |
| overall_abx_prescriptions_count_per_abx_A | 71.000 | 75.000 | 77.500 | 81.250 | 87.000 |
| overall_abx_prescriptions_count_per_abx_B | 90.000 | 92.000 | 95.000 | 100.000 | 102.000 |
| overall_abx_prescriptions_count | 163.900 | 167.750 | 173.000 | 180.000 | 184.100 |
| final_amr_actual_A | 0.323 | 0.327 | 0.334 | 0.336 | 0.341 |
| final_amr_visible_A | 0.323 | 0.327 | 0.334 | 0.336 | 0.341 |
| final_amr_actual_B | 0.321 | 0.326 | 0.331 | 0.335 | 0.340 |
| final_amr_visible_B | 0.321 | 0.326 | 0.331 | 0.335 | 0.340 |

#### Two Antibiotics With Cross-Resistance, Exaggerated Risk Stratification, Expected Reward Greedy

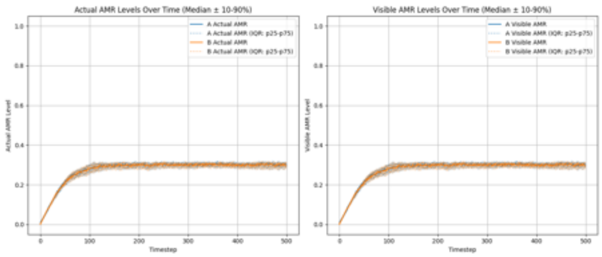

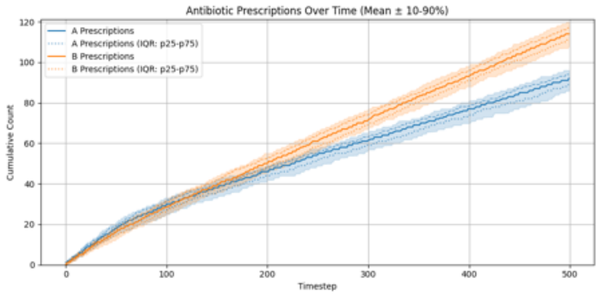

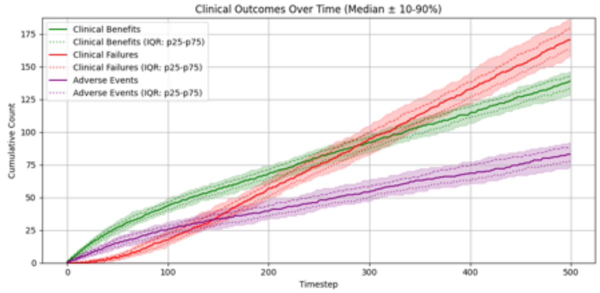

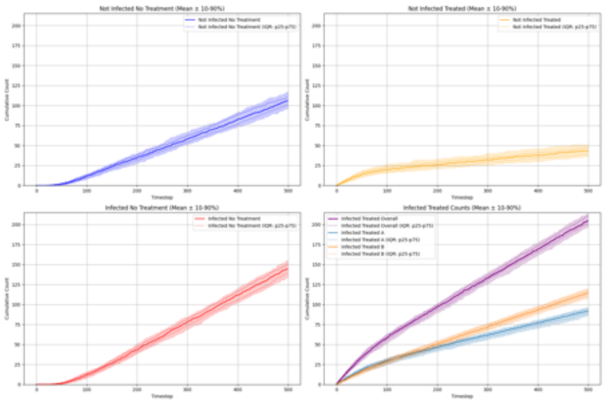

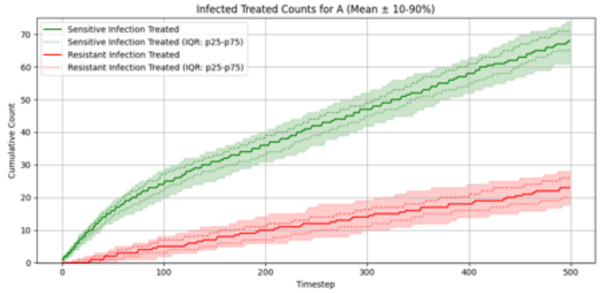

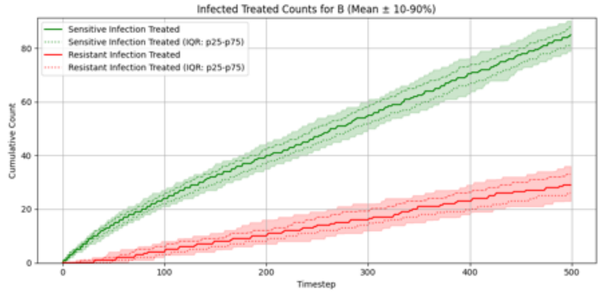

#### Summary Statistics (Two Antibiotics With Cross-Resistance, Exaggerated Risk Stratification, Expected Reward Greedy)

| metric | p10 | p25 | p50 | p75 | p90 |
|---|---|---|---|---|---|
| overall_total_reward | -98.230 | -92.025 | -81.350 | -65.225 | -57.170 |
| overall_count_clinical_benefits | 128.000 | 133.000 | 139.000 | 143.000 | 146.000 |
| overall_count_clinical_failures | 157.800 | 162.750 | 170.500 | 179.000 | 186.200 |
| overall_count_adverse_events | 73.000 | 77.750 | 83.000 | 88.000 | 92.000 |
| overall_not_infected_no_treatment_count | 96.000 | 100.750 | 106.000 | 111.000 | 117.100 |
| overall_not_infected_treated_count | 36.000 | 39.000 | 43.000 | 47.000 | 51.300 |
| overall_infected_no_treatment_count | 133.800 | 139.000 | 144.500 | 150.250 | 155.000 |
| overall_infected_treated_count | 197.000 | 202.000 | 204.500 | 211.000 | 214.100 |
| overall_sensitive_infection_treated_count_per_abx_dict_A | 61.000 | 65.750 | 68.000 | 71.250 | 74.000 |
| overall_sensitive_infection_treated_count_per_abx_dict_B | 78.900 | 81.000 | 85.000 | 88.000 | 90.300 |
| overall_resistant_infection_treated_count_per_abx_dict_A | 17.800 | 20.000 | 23.000 | 26.000 | 28.000 |
| overall_resistant_infection_treated_count_per_abx_dict_B | 23.000 | 26.000 | 29.000 | 33.000 | 36.000 |
| overall_sensitive_infection_treated_count | 143.000 | 148.750 | 154.000 | 158.000 | 161.000 |
| overall_resistant_infection_treated_count | 44.000 | 49.000 | 52.000 | 57.000 | 60.100 |
| overall_abx_prescriptions_count_per_abx_A | 85.900 | 89.000 | 92.000 | 94.000 | 96.000 |
| overall_abx_prescriptions_count_per_abx_B | 108.000 | 111.000 | 114.000 | 117.000 | 120.100 |
| overall_abx_prescriptions_count | 197.000 | 202.000 | 204.500 | 211.000 | 214.100 |
| final_amr_actual_A | 0.282 | 0.294 | 0.303 | 0.308 | 0.311 |
| final_amr_visible_A | 0.282 | 0.294 | 0.303 | 0.308 | 0.311 |
| final_amr_actual_B | 0.281 | 0.292 | 0.301 | 0.307 | 0.310 |
| final_amr_visible_B | 0.281 | 0.292 | 0.301 | 0.307 | 0.310 |

##### Two Antibiotics With Cross-Resistance, Exaggerated Risk Stratification, Expected Reward Lowest AMR

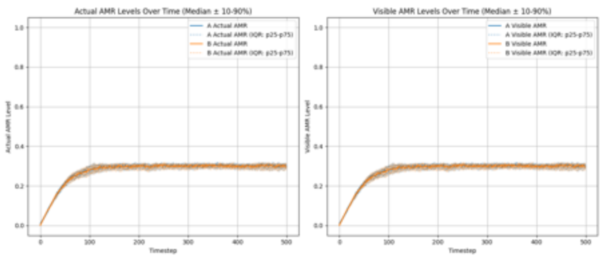

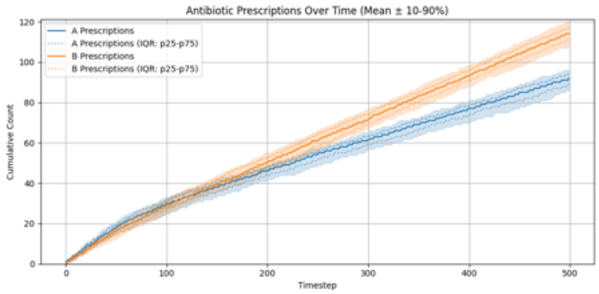

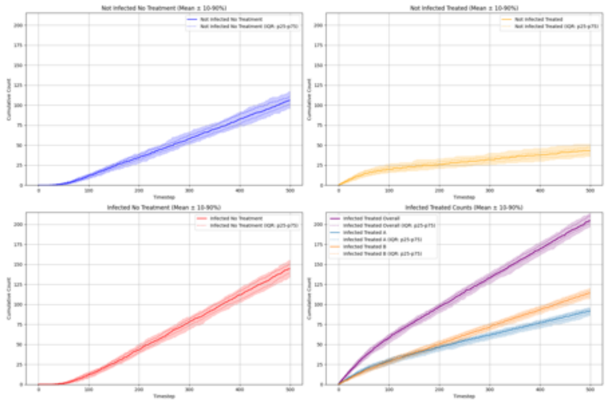

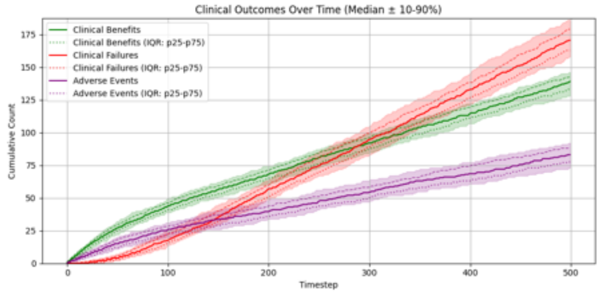

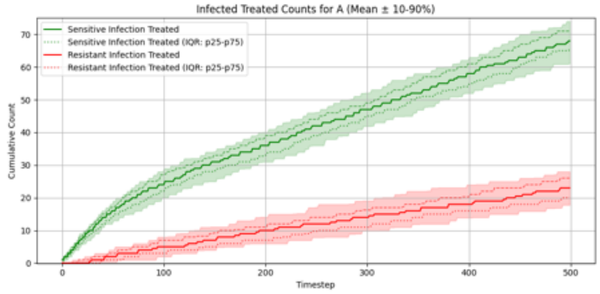

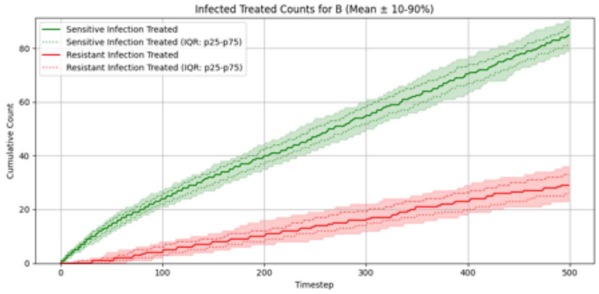

##### Summary Statistics (Two Antibiotics With Cross-Resistance, Exaggerated Risk Stratification, Expected Reward Lowest AMR)

| metric | p10 | p25 | p50 | p75 | p90 |
|---|---|---|---|---|---|
| overall_total_reward | -98.230 | -92.025 | -81.350 | -65.225 | -57.170 |
| overall_count_clinical_benefits | 128.000 | 133.000 | 139.000 | 143.000 | 146.000 |
| overall_count_clinical_failures | 157.800 | 162.750 | 170.500 | 179.000 | 186.200 |
| overall_count_adverse_events | 73.000 | 77.750 | 83.000 | 88.000 | 92.000 |
| overall_not_infected_no_treatment_count | 96.000 | 100.750 | 106.000 | 111.000 | 117.100 |
| overall_not_infected_treated_count | 36.000 | 39.000 | 43.000 | 47.000 | 51.300 |
| overall_infected_no_treatment_count | 133.800 | 139.000 | 144.500 | 150.250 | 155.000 |
| overall_infected_treated_count | 197.000 | 202.000 | 204.500 | 211.000 | 214.100 |
| overall_sensitive_infection_treated_count_per_abx_dict_A | 61.000 | 65.750 | 68.000 | 71.250 | 74.000 |
| overall_sensitive_infection_treated_count_per_abx_dict_B | 78.900 | 81.000 | 85.000 | 88.000 | 90.300 |
| overall_resistant_infection_treated_count_per_abx_dict_A | 17.800 | 20.000 | 23.000 | 26.000 | 28.000 |
| overall_resistant_infection_treated_count_per_abx_dict_B | 23.000 | 26.000 | 29.000 | 33.000 | 36.000 |
| overall_sensitive_infection_treated_count | 143.000 | 148.750 | 154.000 | 158.000 | 161.000 |
| overall_resistant_infection_treated_count | 44.000 | 49.000 | 52.000 | 57.000 | 60.100 |
| overall_abx_prescriptions_count_per_abx_A | 85.900 | 89.000 | 92.000 | 94.000 | 96.000 |
| overall_abx_prescriptions_count_per_abx_B | 108.000 | 111.000 | 114.000 | 117.000 | 120.100 |
| overall_abx_prescriptions_count | 197.000 | 202.000 | 204.500 | 211.000 | 214.100 |
| final_amr_actual_A | 0.282 | 0.294 | 0.303 | 0.308 | 0.311 |
| final_amr_visible_A | 0.282 | 0.294 | 0.303 | 0.308 | 0.311 |
| final_amr_actual_B | 0.281 | 0.292 | 0.301 | 0.307 | 0.310 |
| final_amr_visible_B | 0.281 | 0.292 | 0.301 | 0.307 | 0.310 |

#### Two Antibiotics With Cross-Resistance, Compressed Risk Stratification, Expected Reward Greedy

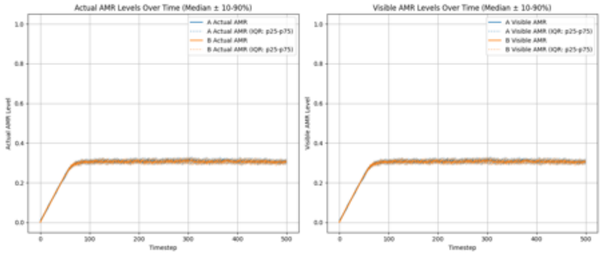

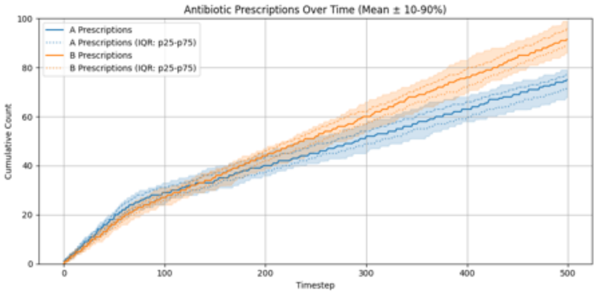

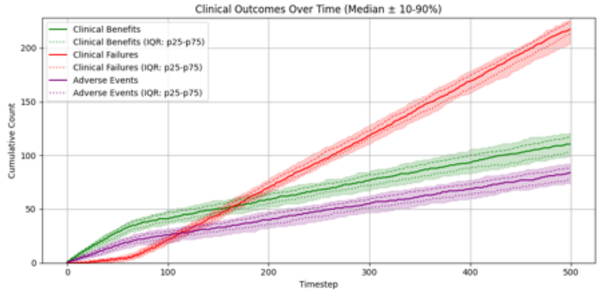

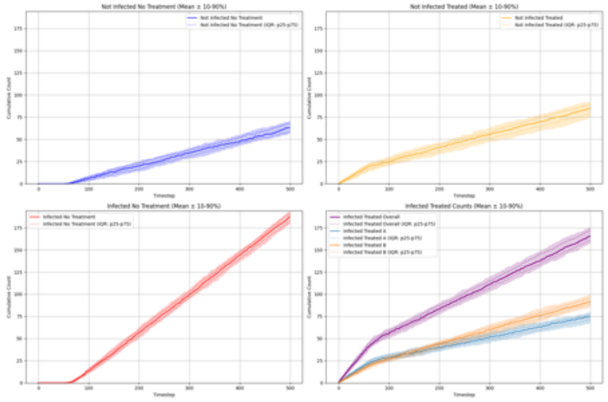

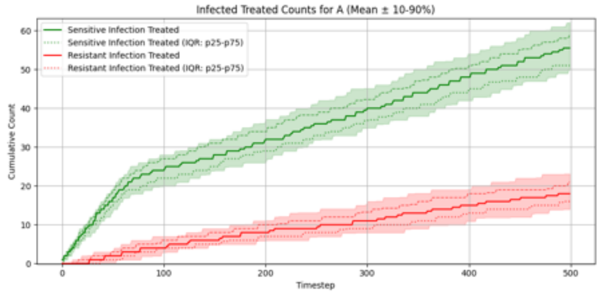

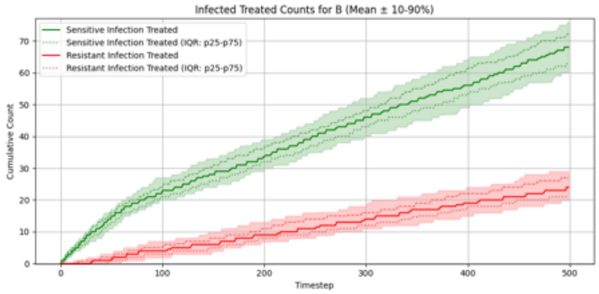

#### Summary Statistics (Two Antibiotics With Cross-Resistance, Compressed Risk Stratification, Expected Reward Greedy)

| metric | p10 | p25 | p50 | p75 | p90 |
|---|---|---|---|---|---|
| overall_total_reward | -202.030 | -192.050 | -185.650 | -169.275 | -161.370 |
| overall_count_clinical_benefits | 99.900 | 102.750 | 110.500 | 117.000 | 121.200 |
| overall_count_clinical_failures | 203.900 | 213.000 | 217.500 | 224.000 | 227.000 |
| overall_count_adverse_events | 74.000 | 77.750 | 84.000 | 88.000 | 92.100 |
| overall_not_infected_no_treatment_count | 56.900 | 58.000 | 63.000 | 68.000 | 71.000 |
| overall_not_infected_treated_count | 74.700 | 78.000 | 85.000 | 89.250 | 92.100 |
| overall_infected_no_treatment_count | 179.900 | 182.000 | 187.500 | 191.000 | 193.100 |
| overall_infected_treated_count | 157.900 | 160.750 | 165.500 | 173.000 | 175.100 |
| overall_sensitive_infection_treated_count_per_abx_dict_A | 49.900 | 51.000 | 55.500 | 59.000 | 62.100 |
| overall_sensitive_infection_treated_count_per_abx_dict_B | 60.000 | 62.750 | 68.000 | 72.250 | 76.000 |
| overall_resistant_infection_treated_count_per_abx_dict_A | 14.000 | 16.000 | 18.000 | 21.000 | 23.100 |
| overall_resistant_infection_treated_count_per_abx_dict_B | 19.900 | 21.000 | 24.000 | 27.000 | 29.000 |
| overall_sensitive_infection_treated_count | 112.000 | 115.750 | 124.000 | 129.250 | 134.000 |
| overall_resistant_infection_treated_count | 34.900 | 39.750 | 43.000 | 46.000 | 50.100 |
| overall_abx_prescriptions_count_per_abx_A | 67.800 | 71.750 | 75.000 | 77.000 | 79.100 |
| overall_abx_prescriptions_count_per_abx_B | 85.900 | 88.750 | 91.500 | 96.000 | 99.000 |
| overall_abx_prescriptions_count | 157.900 | 160.750 | 165.500 | 173.000 | 175.100 |
| final_amr_actual_A | 0.296 | 0.299 | 0.306 | 0.311 | 0.317 |
| final_amr_visible_A | 0.296 | 0.299 | 0.306 | 0.311 | 0.317 |
| final_amr_actual_B | 0.296 | 0.299 | 0.304 | 0.309 | 0.315 |
| final_amr_visible_B | 0.296 | 0.299 | 0.304 | 0.309 | 0.315 |

#### Two Antibiotics With Cross-Resistance, Compressed Risk Stratification, Expected Reward Lowest AMR

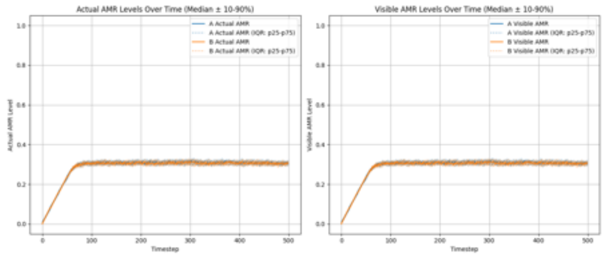

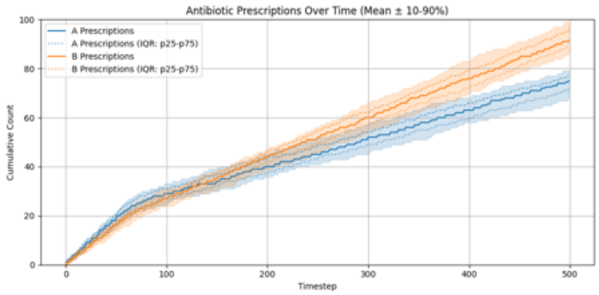

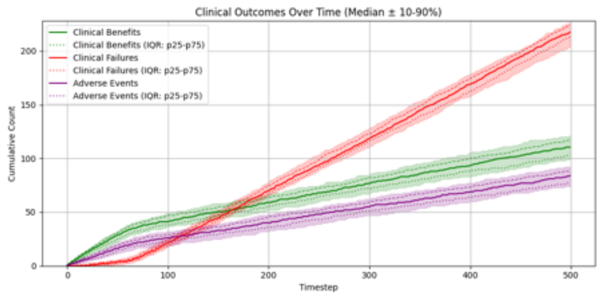

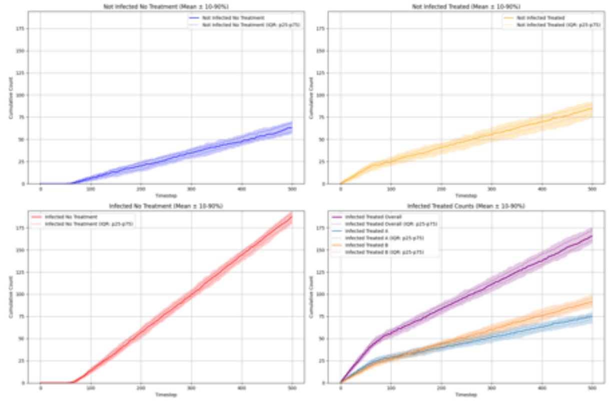

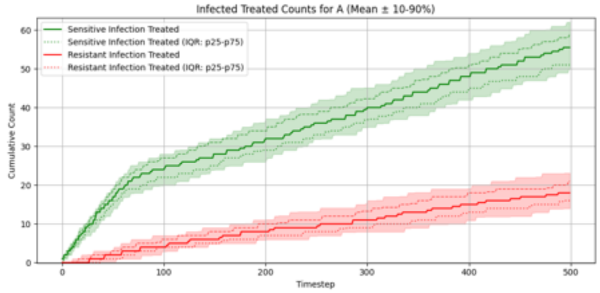

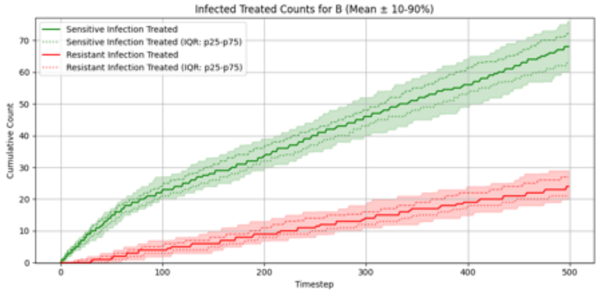

##### Summary Statistics (Two Antibiotics With Cross-Resistance, Compressed Risk Stratification, Expected Reward Lowest AMR)

| metric | p10 | p25 | p50 | p75 | p90 |
|---|---|---|---|---|---|
| overall_total_reward | -202.030 | -192.050 | -185.650 | -169.275 | -161.370 |
| overall_count_clinical_benefits | 99.900 | 102.750 | 110.500 | 117.000 | 121.200 |
| overall_count_clinical_failures | 203.900 | 213.000 | 217.500 | 224.000 | 227.000 |
| overall_count_adverse_events | 74.000 | 77.750 | 84.000 | 88.000 | 92.100 |
| overall_not_infected_no_treatment_count | 56.900 | 58.000 | 63.000 | 68.000 | 71.000 |
| overall_not_infected_treated_count | 74.700 | 78.000 | 85.000 | 89.250 | 92.100 |
| overall_infected_no_treatment_count | 179.900 | 182.000 | 187.500 | 191.000 | 193.100 |
| overall_infected_treated_count | 157.900 | 160.750 | 165.500 | 173.000 | 175.100 |
| overall_sensitive_infection_treated_count_per_abx_dict_A | 49.900 | 51.000 | 55.500 | 59.000 | 62.100 |
| overall_sensitive_infection_treated_count_per_abx_dict_B | 60.000 | 62.750 | 68.000 | 72.250 | 76.000 |
| overall_resistant_infection_treated_count_per_abx_dict_A | 14.000 | 16.000 | 18.000 | 21.000 | 23.100 |
| overall_resistant_infection_treated_count_per_abx_dict_B | 19.900 | 21.000 | 24.000 | 27.000 | 29.000 |
| overall_sensitive_infection_treated_count | 112.000 | 115.750 | 124.000 | 129.250 | 134.000 |
| overall_resistant_infection_treated_count | 34.900 | 39.750 | 43.000 | 46.000 | 50.100 |
| overall_abx_prescriptions_count_per_abx_A | 67.800 | 71.750 | 75.000 | 77.000 | 79.100 |
| overall_abx_prescriptions_count_per_abx_B | 85.900 | 88.750 | 91.500 | 96.000 | 99.000 |
| overall_abx_prescriptions_count | 157.900 | 160.750 | 165.500 | 173.000 | 175.100 |
| final_amr_actual_A | 0.296 | 0.299 | 0.306 | 0.311 | 0.317 |
| final_amr_visible_A | 0.296 | 0.299 | 0.306 | 0.311 | 0.317 |
| final_amr_actual_B | 0.296 | 0.299 | 0.304 | 0.309 | 0.315 |
| final_amr_visible_B | 0.296 | 0.299 | 0.304 | 0.309 | 0.315 |

#### Experiment Set 4

##### Single Antibiotic, Expected Reward Greedy

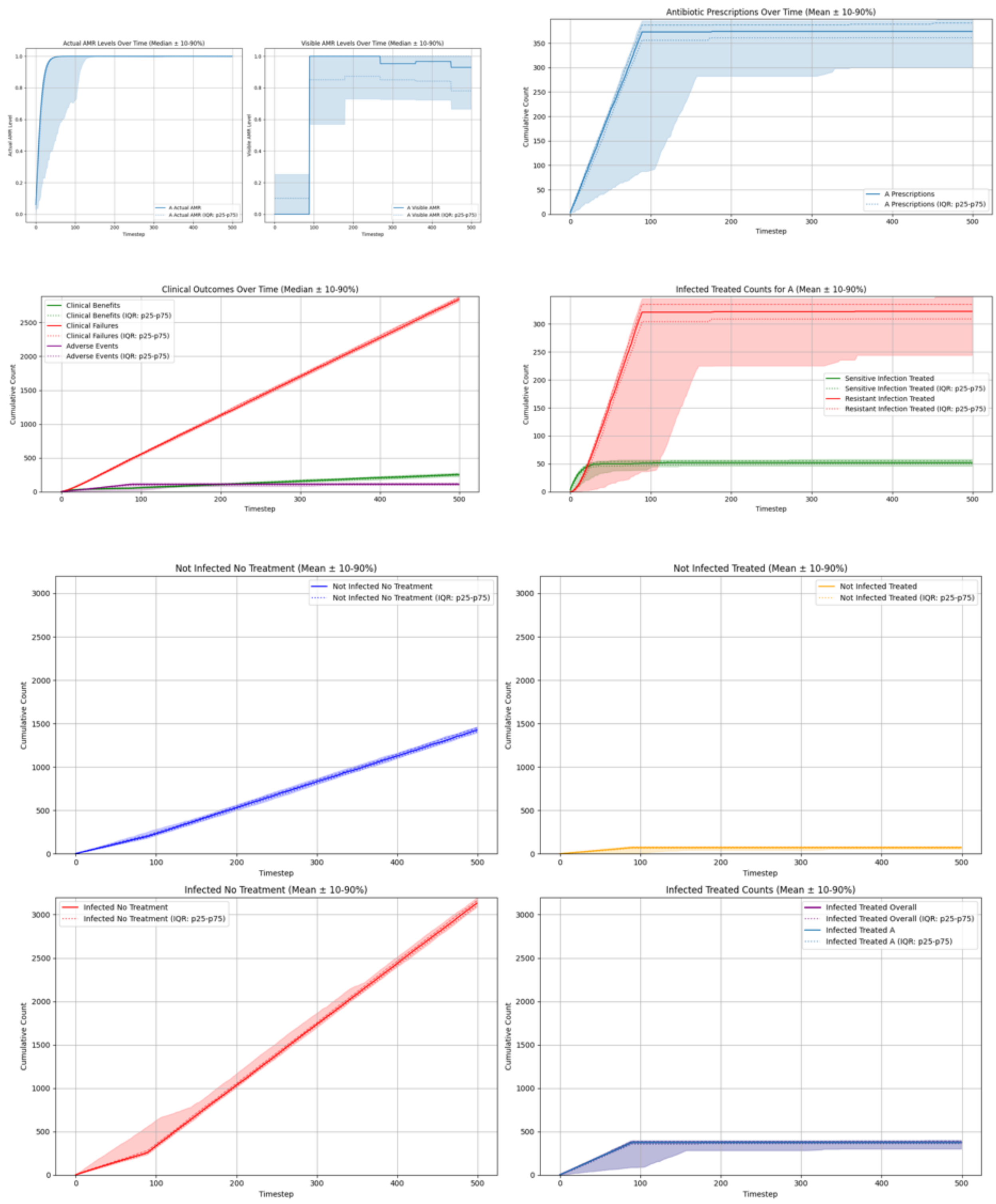

##### Summary Statistics (Single Antibiotic, Expected Reward Greedy)

| metric | p10 | p25 | p50 | p75 | p90 |
|---|---|---|---|---|---|
| overall_total_reward | -294.415 | -291.188 | -287.687 | -284.922 | -282.087 |
| overall_count_clinical_benefits | 224.000 | 243.250 | 252.000 | 264.250 | 280.100 |
| overall_count_clinical_failures | 2811.700 | 2827.000 | 2841.000 | 2863.250 | 2884.500 |
| overall_count_adverse_events | 94.800 | 103.000 | 113.000 | 121.500 | 127.000 |
| overall_not_infected_no_treatment_count | 1393.000 | 1406.750 | 1426.000 | 1448.000 | 1462.700 |
| overall_not_infected_treated_count | 54.800 | 65.000 | 70.500 | 78.250 | 82.200 |
| overall_infected_no_treatment_count | 3089.300 | 3100.750 | 3130.500 | 3156.750 | 3196.800 |
| overall_infected_treated_count | 300.800 | 361.000 | 374.000 | 391.250 | 398.200 |
| overall_sensitive_infection_treated_count_per_abx_dict_A | 46.000 | 49.750 | 52.000 | 55.250 | 58.300 |
| overall_resistant_infection_treated_count_per_abx_dict_A | 244.100 | 309.000 | 322.500 | 335.000 | 348.300 |
| overall_sensitive_infection_treated_count | 46.000 | 49.750 | 52.000 | 55.250 | 58.300 |
| overall_resistant_infection_treated_count | 244.100 | 309.000 | 322.500 | 335.000 | 348.300 |
| overall_abx_prescriptions_count_per_abx_A | 300.800 | 361.000 | 374.000 | 391.250 | 398.200 |
| overall_abx_prescriptions_count | 300.800 | 361.000 | 374.000 | 391.250 | 398.200 |
| final_amr_actual_A | 0.997 | 1.000 | 1.000 | 1.000 | 1.000 |
| final_amr_visible_A | 0.666 | 0.781 | 0.929 | 1.000 | 1.000 |

#### Single Antibiotic, Expected Reward Lowest AMR

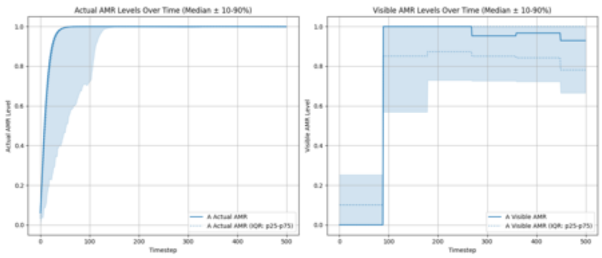

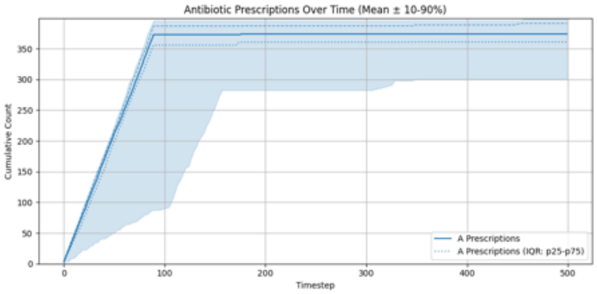

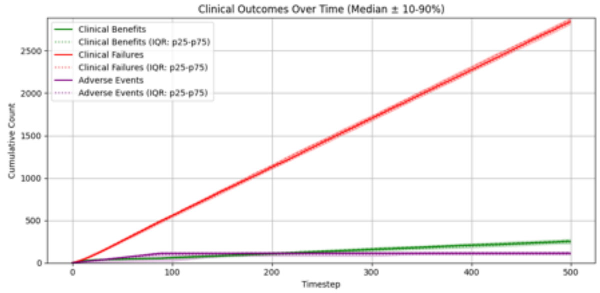

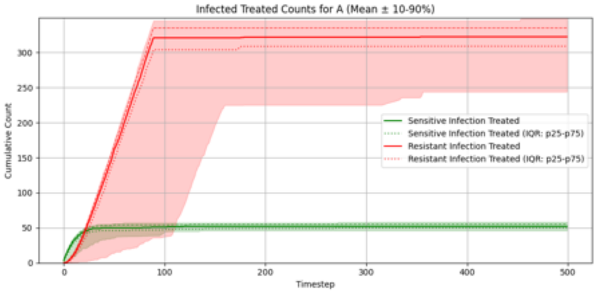

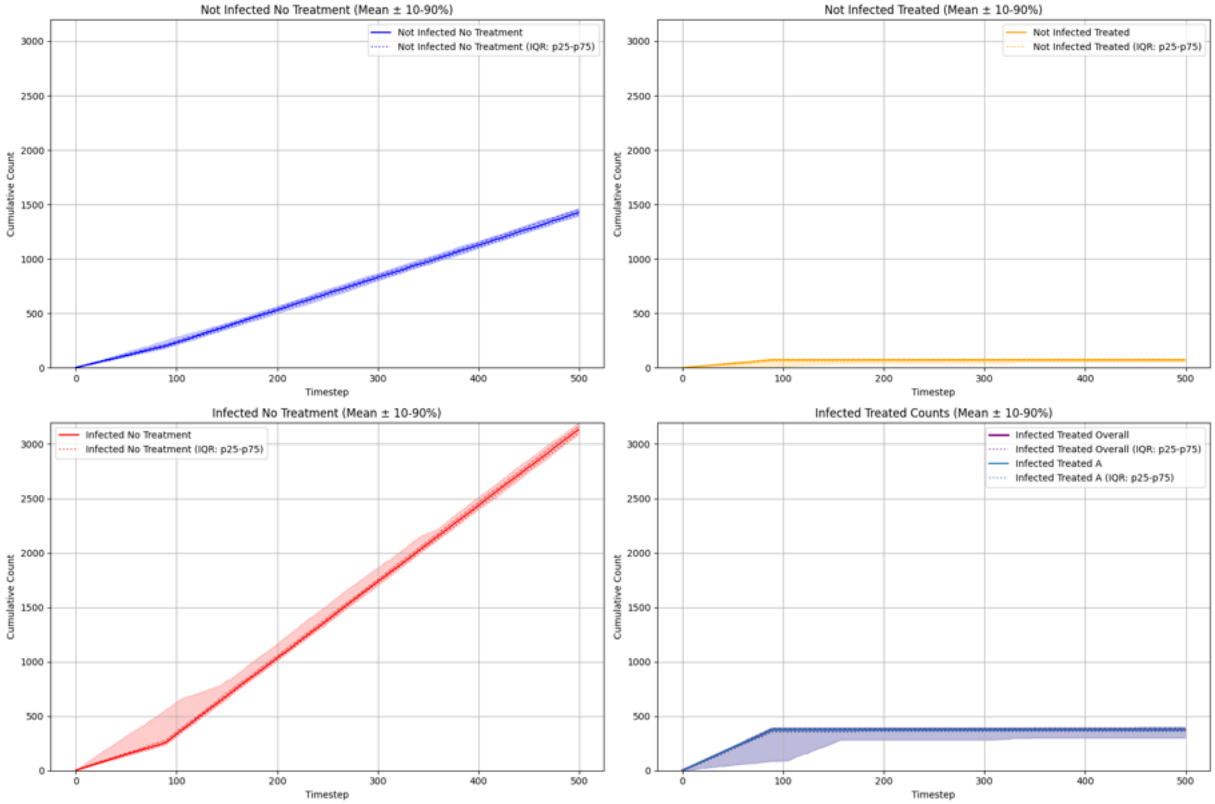

#### Summary Statistics (Single Antibiotic, Expected Reward Lowest AMR)

| metric | p10 | p25 | p50 | p75 | p90 |
|---|---|---|---|---|---|
| overall_total_reward | -294.415 | -291.188 | -287.687 | -284.922 | -282.087 |
| overall_count_clinical_benefits | 224.000 | 243.250 | 252.000 | 264.250 | 280.100 |
| overall_count_clinical_failures | 2811.700 | 2827.000 | 2841.000 | 2863.250 | 2884.500 |
| overall_count_adverse_events | 94.800 | 103.000 | 113.000 | 121.500 | 127.000 |
| overall_not_infected_no_treatment_count | 1393.000 | 1406.750 | 1426.000 | 1448.000 | 1462.700 |
| overall_not_infected_treated_count | 54.800 | 65.000 | 70.500 | 78.250 | 82.200 |
| overall_infected_no_treatment_count | 3089.300 | 3100.750 | 3130.500 | 3156.750 | 3196.800 |
| overall_infected_treated_count | 300.800 | 361.000 | 374.000 | 391.250 | 398.200 |
| overall_sensitive_infection_treated_count_per_abx_dict_A | 46.000 | 49.750 | 52.000 | 55.250 | 58.300 |
| overall_resistant_infection_treated_count_per_abx_dict_A | 244.100 | 309.000 | 322.500 | 335.000 | 348.300 |
| overall_sensitive_infection_treated_count | 46.000 | 49.750 | 52.000 | 55.250 | 58.300 |
| overall_resistant_infection_treated_count | 244.100 | 309.000 | 322.500 | 335.000 | 348.300 |
| overall_abx_prescriptions_count_per_abx_A | 300.800 | 361.000 | 374.000 | 391.250 | 398.200 |
| overall_abx_prescriptions_count | 300.800 | 361.000 | 374.000 | 391.250 | 398.200 |
| final_amr_actual_A | 0.997 | 1.000 | 1.000 | 1.000 | 1.000 |
| final_amr_visible_A | 0.666 | 0.781 | 0.929 | 1.000 | 1.000 |

#### Two Antibiotics No Cross-Resistance, Expected Reward Greedy

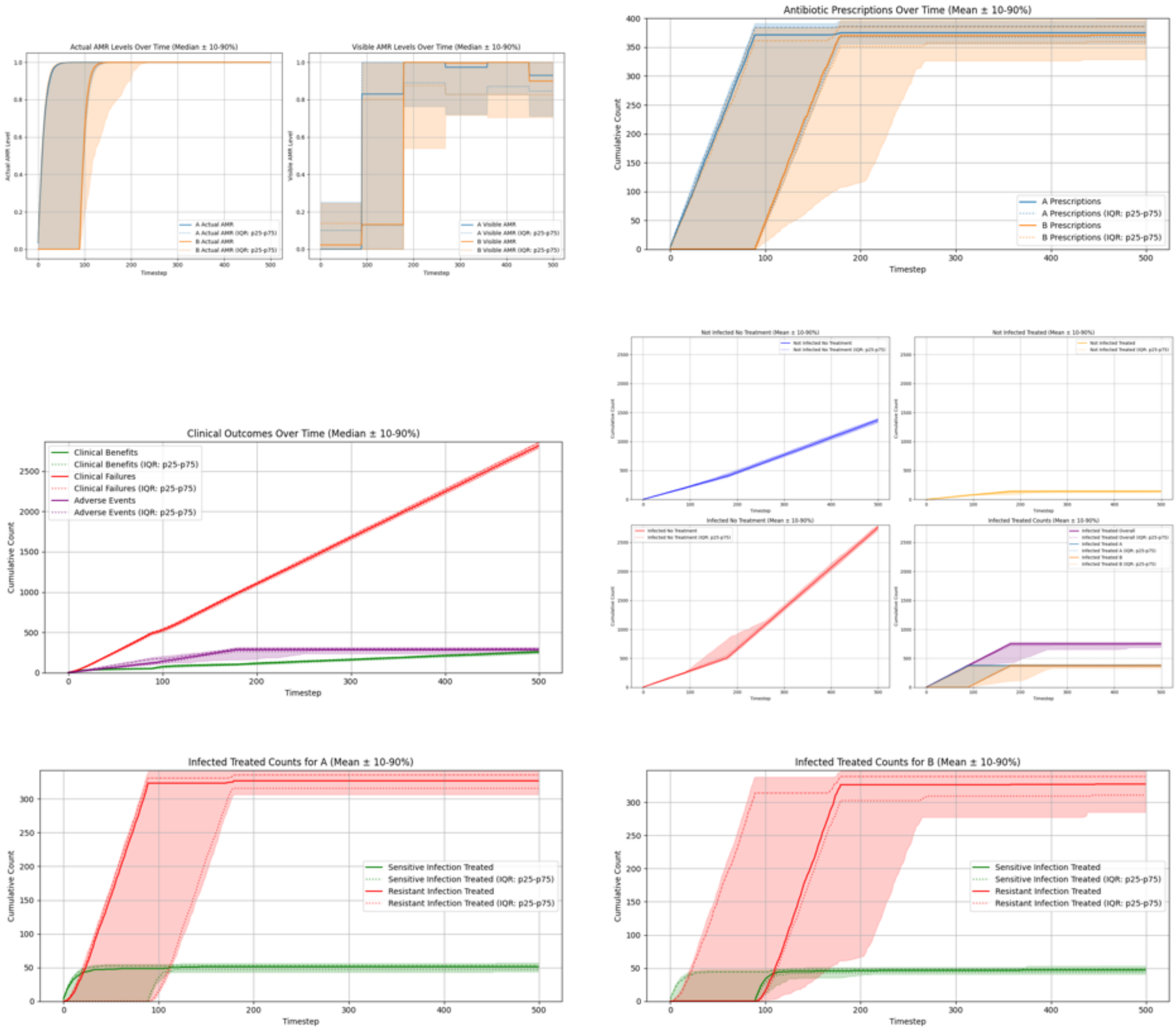

#### Summary Statistics (Two Antibiotics No Cross-Resistance, Expected Reward Greedy)

| metric | p10 | p25 | p50 | p75 | p90 |
|---|---|---|---|---|---|
| overall_total_reward | -299.966 | -296.911 | -293.233 | -290.606 | -286.741 |
| overall_count_clinical_benefits | 236.700 | 248.000 | 257.000 | 267.000 | 272.400 |
| overall_count_clinical_failures | 2786.000 | 2796.000 | 2814.500 | 2848.250 | 2863.000 |
| overall_count_adverse_events | 252.700 | 276.000 | 286.000 | 300.250 | 313.200 |
| overall_not_infected_no_treatment_count | 1326.600 | 1341.000 | 1365.000 | 1386.250 | 1402.800 |
| overall_not_infected_treated_count | 122.000 | 132.750 | 138.500 | 148.250 | 157.100 |
| overall_infected_no_treatment_count | 2708.000 | 2726.000 | 2756.500 | 2774.000 | 2800.800 |
| overall_infected_treated_count | 688.600 | 729.500 | 748.000 | 762.250 | 784.000 |
| overall_sensitive_infection_treated_count_per_abx_dict_A | 43.900 | 47.000 | 51.000 | 53.250 | 57.000 |
| overall_sensitive_infection_treated_count_per_abx_dict_B | 41.000 | 44.000 | 47.000 | 49.000 | 53.100 |
| overall_resistant_infection_treated_count_per_abx_dict_A | 306.800 | 315.750 | 327.000 | 335.500 | 341.400 |
| overall_resistant_infection_treated_count_per_abx_dict_B | 285.300 | 310.750 | 327.500 | 338.750 | 347.100 |
| overall_sensitive_infection_treated_count | 87.900 | 92.000 | 97.000 | 102.250 | 105.100 |
| overall_resistant_infection_treated_count | 592.500 | 633.250 | 652.000 | 667.250 | 685.300 |
| overall_abx_prescriptions_count_per_abx_A | 356.000 | 367.750 | 375.000 | 386.250 | 395.200 |
| overall_abx_prescriptions_count_per_abx_B | 328.900 | 359.750 | 371.000 | 385.250 | 399.100 |
| overall_abx_prescriptions_count | 688.600 | 729.500 | 748.000 | 762.250 | 784.000 |
| final_amr_actual_A | 1.000 | 1.000 | 1.000 | 1.000 | 1.000 |
| final_amr_visible_A | 0.712 | 0.846 | 0.930 | 1.000 | 1.000 |
| final_amr_actual_B | 0.999 | 1.000 | 1.000 | 1.000 | 1.000 |
| final_amr_visible_B | 0.705 | 0.826 | 0.899 | 1.000 | 1.000 |

#### Two Antibiotics No Cross-Resistance, Expected Reward Lowest AMR

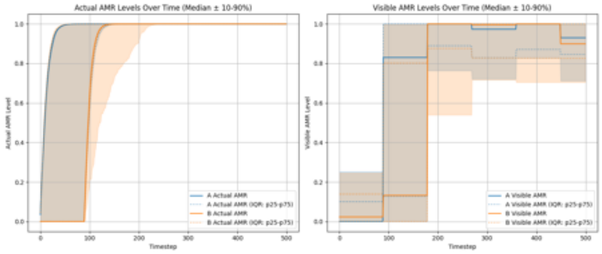

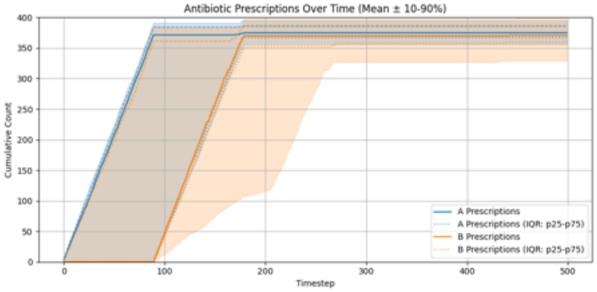

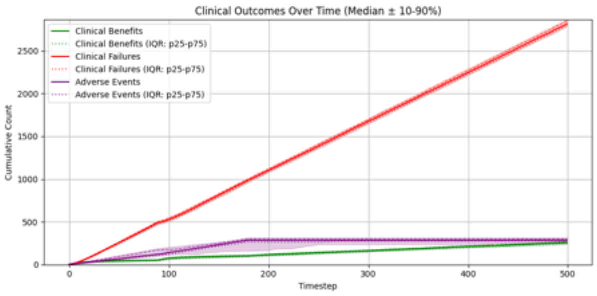

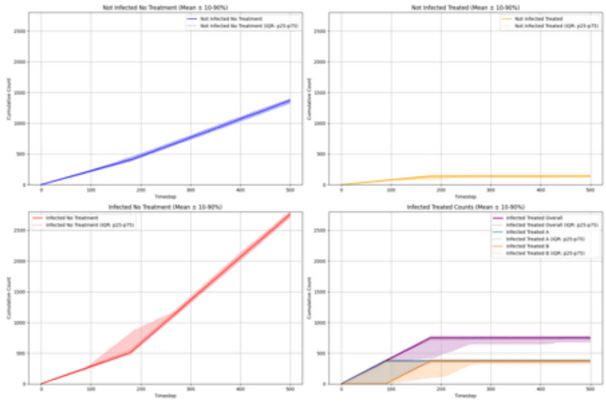

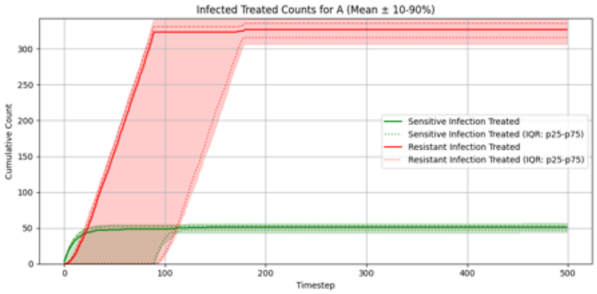

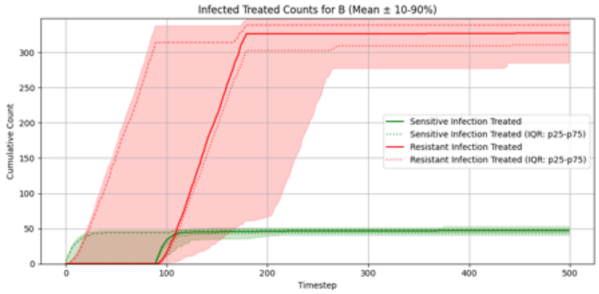

#### Summary Statistics (Two Antibiotics No Cross-Resistance, Expected Reward Lowest AMR)

| metric | p10 | p25 | p50 | p75 | p90 |
|---|---|---|---|---|---|
| overall_total_reward | -299.966 | -296.911 | -293.233 | -290.606 | -286.741 |
| overall_count_clinical_benefits | 236.700 | 248.000 | 257.000 | 267.000 | 272.400 |
| overall_count_clinical_failures | 2786.000 | 2796.000 | 2814.500 | 2848.250 | 2863.000 |
| overall_count_adverse_events | 252.700 | 276.000 | 286.000 | 300.250 | 313.200 |
| overall_not_infected_no_treatment_count | 1326.600 | 1341.000 | 1365.000 | 1386.250 | 1402.800 |
| overall_not_infected_treated_count | 122.000 | 132.750 | 138.500 | 148.250 | 157.100 |
| overall_infected_no_treatment_count | 2708.000 | 2726.000 | 2756.500 | 2774.000 | 2800.800 |
| overall_infected_treated_count | 688.600 | 729.500 | 748.000 | 762.250 | 784.000 |
| overall_sensitive_infection_treated_count_per_abx_dict_A | 43.900 | 47.000 | 51.000 | 53.250 | 57.000 |
| overall_sensitive_infection_treated_count_per_abx_dict_B | 41.000 | 44.000 | 47.000 | 49.000 | 53.100 |
| overall_resistant_infection_treated_count_per_abx_dict_A | 306.800 | 315.750 | 327.000 | 335.500 | 341.400 |
| overall_resistant_infection_treated_count_per_abx_dict_B | 285.300 | 310.750 | 327.500 | 338.750 | 347.100 |
| overall_sensitive_infection_treated_count | 87.900 | 92.000 | 97.000 | 102.250 | 105.100 |
| overall_resistant_infection_treated_count | 592.500 | 633.250 | 652.000 | 667.250 | 685.300 |
| overall_abx_prescriptions_count_per_abx_A | 356.000 | 367.750 | 375.000 | 386.250 | 395.200 |
| overall_abx_prescriptions_count_per_abx_B | 328.900 | 359.750 | 371.000 | 385.250 | 399.100 |
| overall_abx_prescriptions_count | 688.600 | 729.500 | 748.000 | 762.250 | 784.000 |
| final_amr_actual_A | 1.000 | 1.000 | 1.000 | 1.000 | 1.000 |
| final_amr_visible_A | 0.712 | 0.846 | 0.930 | 1.000 | 1.000 |
| final_amr_actual_B | 0.999 | 1.000 | 1.000 | 1.000 | 1.000 |
| final_amr_visible_B | 0.705 | 0.826 | 0.899 | 1.000 | 1.000 |

#### Two Antibiotics With Cross-Resistance, Expected Reward Greedy

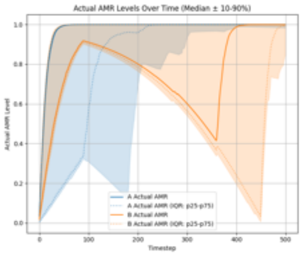

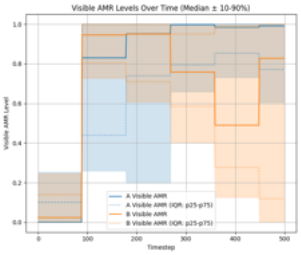

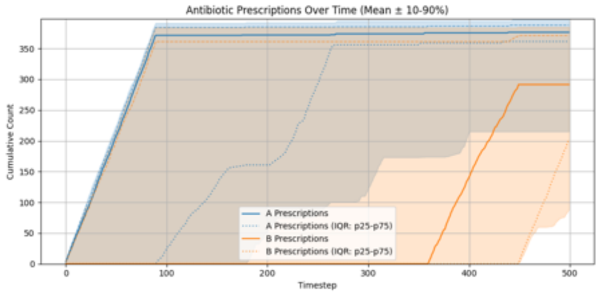

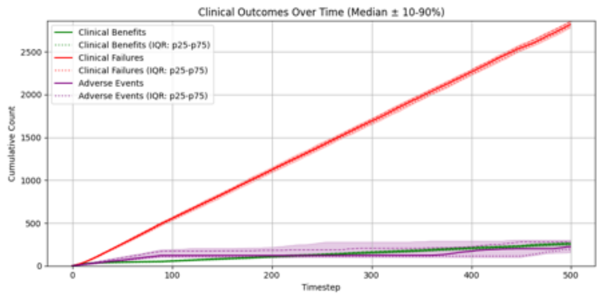

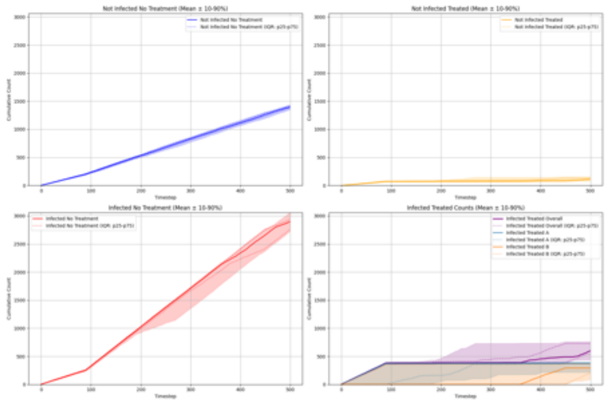

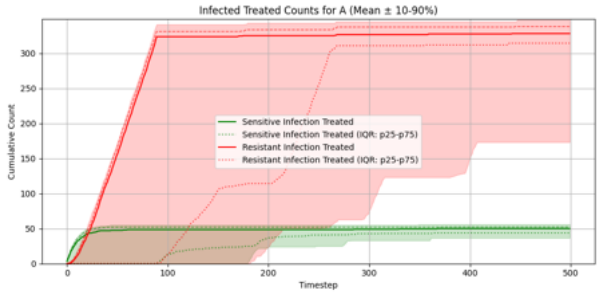

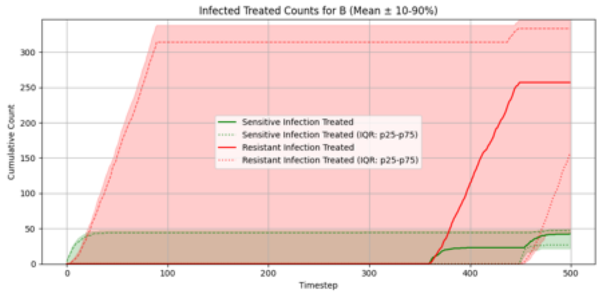

##### Summary Statistics (Two Antibiotics With Cross-Resistance, Expected Reward Greedy)

| metric | p10 | p25 | p50 | p75 | p90 |
|---|---|---|---|---|---|
| overall_total_reward | -299.462 | -294.832 | -289.668 | -285.562 | -283.052 |
| overall_count_clinical_benefits | 236.800 | 247.750 | 257.500 | 267.750 | 278.400 |
| overall_count_clinical_failures | 2781.100 | 2796.250 | 2819.500 | 2846.750 | 2859.400 |
| overall_count_adverse_events | 157.000 | 195.000 | 223.000 | 279.250 | 302.400 |
| overall_not_infected_no_treatment_count | 1348.000 | 1375.750 | 1394.000 | 1418.250 | 1434.300 |
| overall_not_infected_treated_count | 77.800 | 92.750 | 115.000 | 137.000 | 155.000 |
| overall_infected_no_treatment_count | 2722.000 | 2756.000 | 2894.000 | 2944.250 | 3059.500 |
| overall_infected_treated_count | 434.800 | 542.000 | 597.500 | 727.250 | 761.100 |
| overall_sensitive_infection_treated_count_per_abx_dict_A | 36.600 | 43.750 | 50.000 | 52.250 | 56.100 |
| overall_sensitive_infection_treated_count_per_abx_dict_B | 21.800 | 26.750 | 42.500 | 47.000 | 50.100 |
| overall_resistant_infection_treated_count_per_abx_dict_A | 173.100 | 314.250 | 328.000 | 338.000 | 347.500 |
| overall_resistant_infection_treated_count_per_abx_dict_B | 43.900 | 157.250 | 257.000 | 333.250 | 345.200 |
| overall_sensitive_infection_treated_count | 68.000 | 75.750 | 85.000 | 95.250 | 104.100 |
| overall_resistant_infection_treated_count | 365.800 | 453.750 | 506.500 | 641.500 | 683.200 |
| overall_abx_prescriptions_count_per_abx_A | 215.100 | 361.500 | 376.500 | 388.000 | 397.100 |
| overall_abx_prescriptions_count_per_abx_B | 87.700 | 202.250 | 291.500 | 371.250 | 385.100 |
| overall_abx_prescriptions_count | 434.800 | 542.000 | 597.500 | 727.250 | 761.100 |
| final_amr_actual_A | 0.984 | 1.000 | 1.000 | 1.000 | 1.000 |
| final_amr_visible_A | 0.606 | 0.771 | 0.992 | 1.000 | 1.000 |
| final_amr_actual_B | 0.805 | 0.992 | 0.999 | 1.000 | 1.000 |
| final_amr_visible_B | 0.000 | 0.118 | 0.828 | 1.000 | 1.000 |

**Two Antibiotics With Cross-Resistance, Expected Reward Lowest AMR**

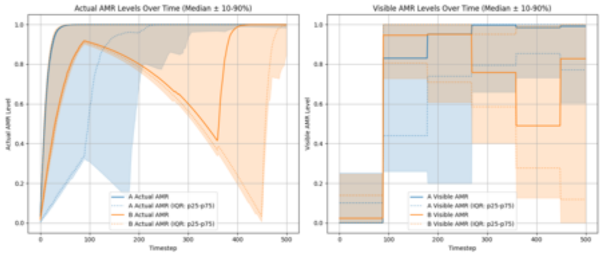

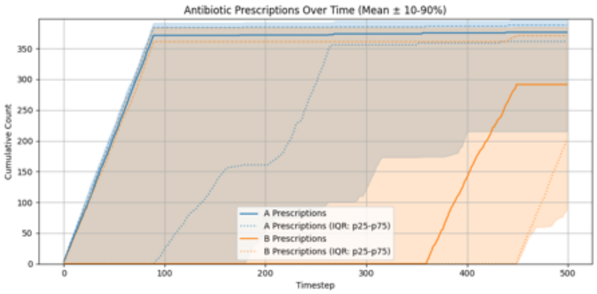

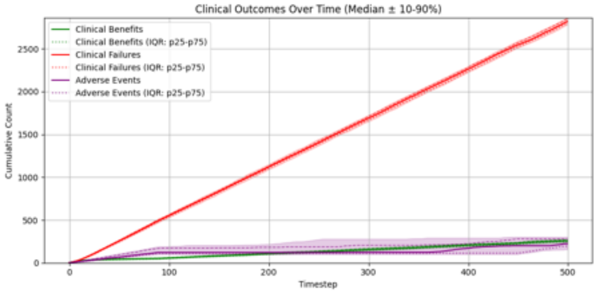

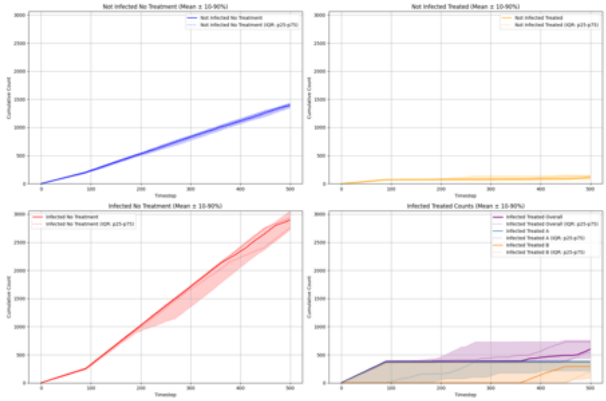

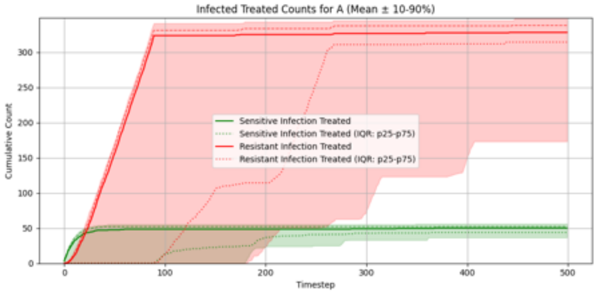

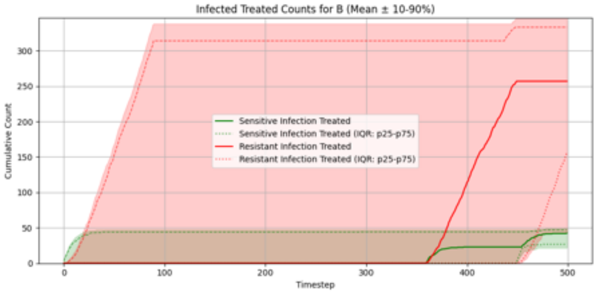

**Summary Statistics (Two Antibiotics With Cross-Resistance, Expected Reward Lowest AMR)**

| metric | p10 | p25 | p50 | p75 | p90 |
|---|---|---|---|---|---|
| overall_total_reward | -299.462 | -294.832 | -289.668 | -285.562 | -283.052 |
| overall_count_clinical_benefits | 236.800 | 247.750 | 257.500 | 267.750 | 278.400 |
| overall_count_clinical_failures | 2781.100 | 2796.250 | 2819.500 | 2846.750 | 2859.400 |
| overall_count_adverse_events | 157.000 | 195.000 | 223.000 | 279.250 | 302.400 |
| overall_not_infected_no_treatment_count | 1348.000 | 1375.750 | 1394.000 | 1418.250 | 1434.300 |
| overall_not_infected_treated_count | 77.800 | 92.750 | 115.000 | 137.000 | 155.000 |
| overall_infected_no_treatment_count | 2722.000 | 2756.000 | 2894.000 | 2944.250 | 3059.500 |
| overall_infected_treated_count | 434.800 | 542.000 | 597.500 | 727.250 | 761.100 |
| overall_sensitive_infection_treated_count_per_abx_dict_A | 36.600 | 43.750 | 50.000 | 52.250 | 56.100 |
| overall_sensitive_infection_treated_count_per_abx_dict_B | 21.800 | 26.750 | 42.500 | 47.000 | 50.100 |
| overall_resistant_infection_treated_count_per_abx_dict_A | 173.100 | 314.250 | 328.000 | 338.000 | 347.500 |
| overall_resistant_infection_treated_count_per_abx_dict_B | 43.900 | 157.250 | 257.000 | 333.250 | 345.200 |
| overall_sensitive_infection_treated_count | 68.000 | 75.750 | 85.000 | 95.250 | 104.100 |
| overall_resistant_infection_treated_count | 365.800 | 453.750 | 506.500 | 641.500 | 683.200 |
| overall_abx_prescriptions_count_per_abx_A | 215.100 | 361.500 | 376.500 | 388.000 | 397.100 |
| overall_abx_prescriptions_count_per_abx_B | 87.700 | 202.250 | 291.500 | 371.250 | 385.100 |
| overall_abx_prescriptions_count | 434.800 | 542.000 | 597.500 | 727.250 | 761.100 |
| final_amr_actual_A | 0.984 | 1.000 | 1.000 | 1.000 | 1.000 |
| final_amr_visible_A | 0.606 | 0.771 | 0.992 | 1.000 | 1.000 |
| final_amr_actual_B | 0.805 | 0.992 | 0.999 | 1.000 | 1.000 |
| final_amr_visible_B | 0.000 | 0.118 | 0.828 | 1.000 | 1.000 |
